# Syngeneic human iPSC Macrophages and Microglia possess distinct neuroinflammatory profiles, responses to HIV infection, and ART treatment

**DOI:** 10.64898/2026.11.17.767399

**Authors:** Oluwatofunmi Oteju, Marzieh Daniali, Taylor Kist, Xiaoke Xu, Katy Emanuel, Meng Niu, Will Dampier, Howard Fox, Peter J. Gaskill

**Affiliations:** Drexel University College of Medicine, Department of Pharmacology and Physiology, Philadelphia, PA, 19102, USA; University of Nebraska, Department of Neurological Sciences, Omaha, NE, 68198, USA; Drexel University College of Medicine, Department of Microbiology and Immunology, Philadelphia, PA 19102, USA

**Keywords:** iPSC, iPSC-Microglia, iPSC-Macrophages, Neuroinflammation, NF-κB, NeuroHIV, Macrophages, Antiretroviral therapy

## Abstract

Macrophages are innate immune cells that primarily function as professional phagocytes, playing essential roles throughout the body in host defense and maintaining homeostasis. The CNS is populated by two anatomically distinct macrophage populations; parenchymal microglia and border-resident CNS-associated macrophages (CAMs). These populations coordinate the neuroinflammatory response to pathogen-associated molecular patterns (PAMPs) through Toll-like receptor (TLR) signaling and are exposed to distinct microenvironmental cues that could shape their functions. CNS macrophages are also involved in infection with neurotropic viruses such as HIV, where the persistent infection contributes to chronic neuroinflammation and the constellation of other neurological symptoms collectively termed neuroHIV. Although antiretroviral therapy (cART) has transformed HIV care and lessened the severity of many neuroHIV symptoms, the persistence of CNS viral reservoirs and neurological deficits remains a major comorbidity and barrier to cure.

Understanding specific contributions of microglia and CAMs to inflammation and neuroHIV has been challenging due to the lack of genetically matched human model systems enabling direct mechanistic comparison between of microglia and CAMs. To address this, we optimized the generation of syngeneic, human-induced pluripotent stem cell (iPSC) derived microglia-like (iMg) and CAM-like (iMacs) cells. These are generated simultaneously from a single donor line and have the same genetic background, enabling side-by-side characterization and functional analyses of inflammation and HIV infection. Transcriptomic profiling by RNA sequencing revealed substantial baseline differences in inflammatory and immune signaling pathways between iMg and iMacs. These intrinsic differences were also observed in TLR-specific NF-κB responses. Stimulation of TLR4 drove heightened response in iMg relative to iMac, correlating closely with morphological changes, while stimulating TLR2 had a contrasting effect, eliciting a stronger and more robust early NF-κB response in iMacs. In HIV infection studies, iMacs were more permissive to viral replication at baseline, whereas iMg showed a reduced susceptibility to infection. However, iMacs responded more robustly to cART, while iMg exhibited relative resistance to cART-mediated suppression. These findings delineate fundamental inflammatory and virologic differences between human microglia and CAMs and provide a robust foundation on which to further examine HIV infection, cART efficacy, and immune function in these distinct CNS myeloid populations.

## Introduction

Myeloid cells are essential components of the immune system and are primarily derived from hematopoietic stem and progenitor cells in the bone marrow. Among the different types of myeloid cells, monocytes play critical roles in immune surveillance, tissue homeostasis, and repair. Monocytes also transmigrate from the bloodstream into tissues, differentiating into infiltrating macrophages in response to inflammation or damage. In some cases, monocytes can also differentiate to acquire tissue-specific phenotypes and functions, supplementing the existing population of tissue resident macrophages. Macrophages are key regulators of innate immunity, contributing to pathogen recognition, phagocytosis, antigen presentation, inflammatory signaling, and responses to diverse tissue insults^1,2^.

Within the central nervous system (CNS), macrophage populations can be broadly divided into two distinct groups: parenchymal microglia and non-parenchymal border-associated macrophages (BAMs), also referred to as CNS-associated macrophages (CAMs), which reside at CNS interfaces such as the perivascular, choroid plexus and meningeal regions^3–5^. Microglia originate from yolk sac progenitors and are maintained primarily through self-renewal, whereas BAMs are more readily replenished by circulating monocytes, particularly under inflammatory conditions^4,5^. Both cell types play a major role in the response to CNS insults, including neurotropic viruses, organizing the immune response to viral infection of the CNS, particularly through the promulgation of neuroinflammation, which is largely regulated by CNS myeloid cells^6,7^.Although both populations contribute to innate immune defense and tissue homeostasis, they differ substantially in their developmental origins, transcriptional programs, tissue localization, morphological alterations and responses to inflammatory cues ^8^.

Much of this neuroinflammation is regulated via the Nuclear factor kappa-light-chain-enhancer of activated B cells (NF-κB) pathway, a transcription factor that acts as a central mediator of immune responses in these cell types. Following detection of pathogen- or damage-associated molecular patterns (PAMPs and DAMPs), NF-κB rapidly activated by phosphorylation and subsequent nuclear translocation^9^, where it regulates the expression of numerous genes involved in inflammation, including those encoding various proinflammatory cytokines, chemokines, and other immunomodulatory molecules^10^. These mediators are crucial in innate immune responses, as well as involved in initiating responses from the adaptive immune system. Additionally, this signaling axis is specifically important in the context of HIV infection and how these cells (macrophages and microglia) contribute to both antiviral immune responses and persistent inflammation.

Human and rodent studies have found that primary macrophages and microglia have robust NF-κB activation in response to canonical stimuli such as LPS or viruses such as HIV ^11,12^, but direct comparisons between macrophages and microglia derived from the same genetic background are limited. This makes it difficult to determine how the inflammatory responses of these cell types differ within the CNS.

In HIV infection specifically, persistent infection of both types of brain macrophages, particularly microglia, is central to the formation of a CNS reservoir, where HIV can persist despite suppressive anti-retroviral therapy (ART)^13,14^. The presence of these reservoirs, and the sustained impact they have on the CNS inflammation are central to the development of HIV associated brain injury (HABI)^15^. Both microglia and CAMs express the HIV entry receptors CD4 and CCR5^16–18^ and support persistent infection and elaborate inflammatory mediators, with the specific differences in these processes between macrophage populations not well defined. Neurocognitive impairment resulting from HABI currently impacts 15 – 50% of people with HIV despite the use of ART, and current data show that persistent neuroinflammation, rather than infection alone, is a primary driver of neuroHIV ^19–21^. Thus, defining how responses in discrete cell types drive HABI is critical to developing effective therapies^22^. Significant progress has been made in this area using non-human primate models and post-mortem microglia^8,21,23^, but precisely defining these cell-type-specific mechanisms remains challenging, as most studies cannot directly compare primary microglia and macrophages from the same individual. This makes it difficult to parse the impact of human heterogeneity on the experimental readout, confounding attribution of specific changes to cell type rather than environment or genetic background.

The majority of human experimental models used to examine macrophage biology are insufficient to fully address these gaps. Immortalized cell lines lack HIV receptor expression and can display aberrant inflammatory profiles, while primary microglia and macrophages are limited by scarcity, donor variability, and tractability^24^. These constraints have hindered efforts to dissect how distinct CNS myeloid populations respond to innate immune stimulation (e.g., TLR-mediated stimulation) and viral pathogenesis (e.g., HIV). The development of induced pluripotent stem cell (iPSC) technology can help to address these challenges by enabling the generation of multiple CNS-relevant cell types from a single genetic background^25–28^. Using a syngeneic platform allows parallel derivation of microglia-like (iMg) and CAM-like (iMac) cells that retain key transcriptional and functional features of their primary cell counterparts. These systems provide a unique model to directly compare how brain myeloid cell populations respond to inflammatory stimuli, including TLR-derived NF-κB activation, and how these cells respond to HIV and cART response.

Using this model to compare syngeneic iMg and iMac, we show that these cell types exhibit fundamentally distinct inflammatory and virologic phenotypes. This includes differences in TLR-mediated NF-κB activation, cytokine secretion, and transcriptomic profiles by RNA sequencing. While both cell types displayed muted response NF-κB activation following Poly I:C (TLR3) and imiquimod (TLR7) exposure, LPS stimulation of TLR4 showed that iMg were more sensitive than iMacs and activated NF-κB in an ultrasensitive, switch-like manner. In contrast, stimulation of TLR2 with Pam3CSK4 showed that iMacs mounted a stronger and more sensitive NF-κB response at early time points, whereas prolonged stimulation led to convergence of responses between the two cell types. In response to infection with the M-tropic strain HIV_ADA_, iMacs were more permissive to viral replication than iMg, showing significantly higher numbers of infected cells and secretion of p24Gag. Yet iMacs also showed greater responsiveness to the antiretroviral drugs comprising Biktarvy, whereas iMg were less responsive to ART mediated viral suppression. These findings delineate key functional differences between human macrophages and microglia and suggest that current cART regimens may be more effective in macrophages than in microglia, highlighting the need for microglia-targeted therapeutic strategies to address persistent CNS reservoirs.

## Materials & Methods

### Reagents

RPMI-1640 (cat #11875119) medium, Advanced DMEM F12 (cat #12634010), penicillin/streptomycin (P/S) (cat #15140163), Glutamax (cat #35050061), and TrypLE Express (cat #12604013) from Invitrogen (ThermoFisher, Carlsbad, CA, USA). Bovine serum albumin (BSA) (cat #BP1600100), glycine (cat #G48500), and 4% Paraformaldehyde solution (cat #AAJ19943K2) from Fisher Scientific (Waltham, MA, USA). Fetal calf serum (FBS) from Corning (cat #MT35010CV). Hydroxyethyl piperazineethanesulfonic acid (HEPES) (cat #BP2991), Tween (cat #NC0689073), Lipopolysaccharide (LPS from E. Coli 055:B5; cat #L2880-10mg), Polyinosinic–polycytidylic acid sodium salt (Poly I: C, cat # P1530-25MG) and dimethyl sulfoxide (DMSO) (cat #472301) from Sigma-Aldrich (St. Louis, MO, USA). Pam3CSK4 (cat #4633) and Imiquimod (cat #3700) from R&D Systems (Minneapolis, MN). Macrophage colony-stimulating factor (M-CSF) (cat #300-25), IL-34 (cat #200-34), and TGF-β1 (cat #100-21) from Peprotech (Rocky Hill, NJ, USA). Bictegravir (cat #26532), Tenofovir Alafenamide (cat #26214), and Emtricitabine (cat #16879) from Cayman Chemical Company (Ann Arbor, MI, USA). All drugs were resuspended in their respective solvents, with Biktarvy (bictegravir, emtricitabine, and tenofovir alafenamide) in DMSO and others in dH_2_O, then aliquoted and stored in -20°C until use. TaqMan Fast Universal Master Mix, and PCR assay probes for TMEM119 (Hs01938722_u1), P2RY12 (Hs00224470_m1), IBA1/AIF1 (Hs00610419_g1), SALL1 (Hs01548765_m1), CD68 (Hs00154355_m1), CCR5 (Hs99999149_s1), MKI67 (Hs01032443_m1), and 18s (4319413E) genes were purchased from Applied Biosystems (ThermoFisher, Waltham, MA, USA).

### Differentiation and Culture of Human Induced Pluripotent Stem Cell Derived Microglia-like Cells (iMg)

Human microglia-like cells (iMg) were differentiated from induced pluripotent stem cells obtained from the Children’s Hospital of Philadelphia (CHOP) as common myeloid progenitors (CMPs). In this study, we included iMg derived from three (3) distinct iPSC lines (WT6, WT10, WT17). This process uses an established 11-day protocol^29–31^ generating iMg that express common microglia identity markers, including P2RY12, TMEM119, and IBA-1 (Figure 1). Briefly, CMPs are differentiated in RPMI-1640 supplemented with 10% FBS, 0.1% PenStrep, IL-34 (100ng/mL, PeproTech, 200-34-10UG), TGF-β (50ng/ml, PeproTech, 100-21-10UG), and M-CSF (25ng/ml, PeproTech, 300-25-10UG) at 37°C at 5% CO_2_. Half media changes were performed on days *in vitro* (DIV) 2 and 5, while full media changes were performed on DIV8 and 11, being careful to add media gently and avoid contact between pipette tip and plate. The growth factors IL-34, TGFβ, and M-CSF were added fresh with each media. At DIV11, cells were considered mature and ready for downstream assays. iMg were cultured in glass-like polymer plates (Cellvis, 96-well, P96-1.5P or 24-well, P24-1.5P, depending on the experiments) at a density of about 312.5 cells/mm^2^, 10,000 cells / well in 96-well plates, and 66,600-75,000 cells / well in 24-well plates. Optimal cell density and plating conditions are critical for successful differentiation and responsiveness to inflammatory stimuli.

**Figure 1:**
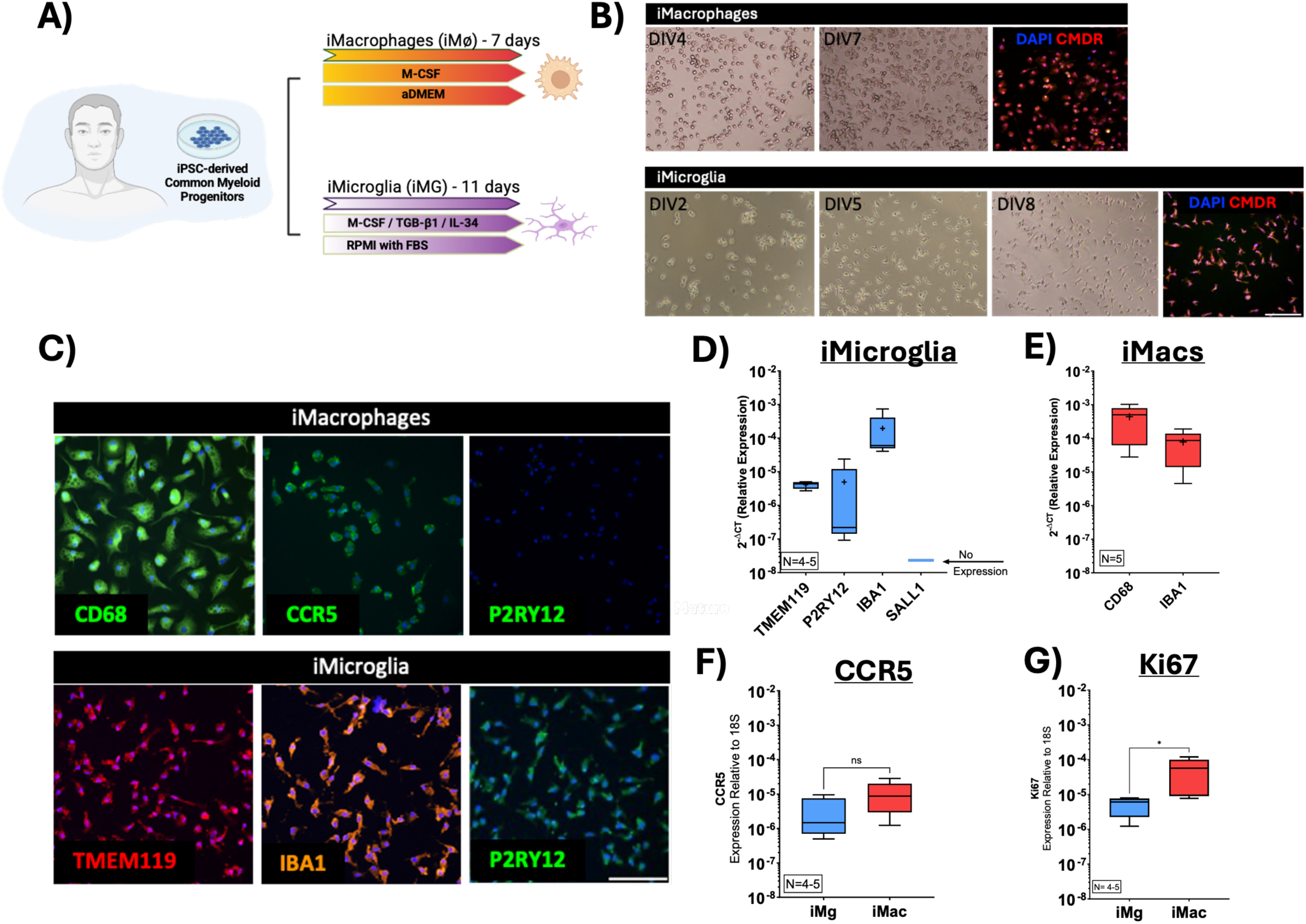
Generation of syngeneic iPSC -derived microglia and macrophages. (A) Schematic timeline and differentiation protocol for syngeneic iMg and iMac generation. iMacs were differentiated from common myeloid progenitors (CMPs) over 7 days in aDMEM supplemented with M-CSF (100 ng/mL). iMg were differentiated from CMPs using CSF1R-stimulating cytokines (IL-34 and M-CSF) and TGF-β1 for 11 days. (B) Representative brightfield images of iMacs at DIV4 and DIV7 (mature), and iMg at DIV2, DIV5, DIV8, and DIV11 (mature). Scale bar represents 20uM. . (C) Fully mature iMg and iMac after the differentiation period were stained with identity markers via immunofluerest imaging; CD68 (green), CCR5 (green) were used as markers for iMacs and TMEM119 (red), IBA1 (yellow), and P2RY12 (green) were used as identity markers for iMg. Scale bar represents 20uM. (D,E) expression of maturation markers was assessed using qPCR in iMg (D) as well as iMacs (E). (F,G) Gene expression of the CCR5 and proliferation marker MKI67 (Ki67) in iMg versus iMacs (Mann–Whitney test, *P < 0.05)

### Differentiation and Culture of Human Induced Pluripotent Stem Cell Macrophage-Like cells (iMacs)

Human, macrophage-like cells (iMacs) were also differentiated from induced pluripotent stem cells obtained from the Children’s Hospital of Philadelphia (CHOP) as common myeloid progenitors (CMPs). In this study, we included iMacs derived from 3 distinct iPSC lines (WT6, WT10, WT17), the same lines used to generate the iMg. The iMac differentiation process uses a 7-day protocol adapted from Vaughan Jackson et al. 2021^32^, generating iMac that express macrophage identity markers, including CD68, CCR5 but not P2RY12 (Figure 1). In brief, CMPs are differentiated in Advanced DMEM/F12 supplemented with 1% Glutamax, 1% PenStrep, 15mM HEPES, M-CSF (100ng/ml, PeproTech, 300-25-10UG) at 37°C at 5% CO_2_. A half media change is performed at DIV 4 with M-CSF added fresh with media change. At DIV 7, cells were considered mature and ready for downstream assays. iMacs were cultured in glass-like polymer plates (Cellvis, 96-well, P96-1.5P or 24-well, P24-1.5P, depending on the experiments) at a density of about 625 cells/mm^2^, 20,000 cells / well in 96-well plates, and 144,000 cells / well in 24-well plates.

### Immunofluorescence Staining

Cells were fixed and permeabilized using 4% PFA for 20 minutes at 4°C, then washed with PBS and incubated with blocking buffer (1% BSA, 0.1% Tween 20, and 22.52 mg/mL glycine in 1X PBS) for 1 hour at room temperature. Cells were then incubated with primary antibodies including anti-HIV-1 p24 monoclonal (AG3.0) primary antibody (4 µl/ml, which was obtained through the NIH HIV Reagent Program, Division of AIDS, NIAID, NIH: Monoclonal Anti-Human Immunodeficiency Virus Type 1 (HIV-1) p24 (AG3.0), ARP-4121, contributed by Dr. Marie-Claire Gauduin), anti-NF-κB (p65) primary antibody (2.5 µl/ml, Cell Signaling Technology, cat #8242), P2Y12 (2.5 µl/ml, Thermo Fisher Scientific, cat #702516), TMEM119 (2.5 µl/ml, Thermo Fisher Scientific, cat #PA5-119902), CD68 (2.5 µl/ml, Thermo Fisher Scientific, cat #MA513324), IBA-1 (2.5 µl/ml, Thermo Fisher Scientific, cat #PA5121836), and CCR5 (2.5 µl/ml, eBioscience, cat #12-1957-41) overnight at 4°C. The following day, cells were incubated with Alexa Fluor 488 secondary antibody (A-11001 or A-11008, Thermo Fisher) for 1 hour at room temperature and were counterstained with DAPI (0.2µg/mL, D1306, Thermo Fisher) and HCS Deep Red (6µg/mL, H32721, Thermo Fisher) for 10 minutes, then preserved in 1xPBS.

### High Content Imaging of NF-kB Nuclear Translocation

High-content imaging of the cells was performed using the Cellomics CellInsight CX7 HCS Platform, capturing 20 fields per well at 20x magnification as we have done previously^33^. To analyze NF-κB translocation into the nucleus from the cytoplasm, mature cells (iMg and iMac) plated in 96-wells were treated with various TLR agonists from 0.001 – 100ng/ml (TLR2, Pam3CSK4; TLR 3, polyinosinic:polycytidylic acid (Poly I:C); TLR 4, Lipopolysaccharide, (LPS); TLR 7, Imiquimod) for 30 or 60 minutes. Afterwards, the plates were fixed and stained. The ratio of the NF-κB translocation to the nucleus was assessed using high-content imaging and analysis. Following image acquisition, 3-channel colocalization was performed using the sample parameters **(Supp. Table 1**). In brief, binary masks were created to identify the cytoplasmic (HCS-CMDR) and the nuclear (DAPI) regions of interest (ROI); and then the average intensity of NF-κB in each ROI for every cell is quantified. The NF-κB average intensity in the nuclear ROI (DAPI) is divided by the NF-κB average intensity in the cytoplasmic ROI (HCS-CMDR), excluding the nuclear ROI, to generate a nuclear translocation ratio at cell levels. This quantifies the relative amount of NF-κB in the nucleus, while also controlling for cell size and differences in total NF-κB between cells. The colocalization ratio is averaged across all wells per condition, then used for statistical analysis.

### High Content Imaging of Morphological Parameters

Images acquired using the Cellomics CellInsight CX7 High-Content Screening (HCS) Platform for analysis of NF-κB nuclear translocation were used to quantify cellular morphological parameters **(Supp. Table 2)**, including the length-to-width ratio (LWR), perimeter-to-area ratio (P2A), and total process length (TPL). Cells were treated with the TLR4 agonist lipopolysaccharide (LPS) at concentrations ranging from 0.001 to 100 ng/mL and analyzed at two time points (30 and 60 min). Image analysis was performed using the HCS-CMDR channel set as the primary channel, with the threshold carefully optimized for this channel. The DAPI channel was additionally used to identify cell nuclei and improve segmentation accuracy.

### Correlative analysis of NF-κB activation and morphological remodeling

Well summaries exported from the CellInsight CX7 were analyzed in Python 3.14.4 with pandas 3.0.3. For each cell type, stimulation time, and LPS dose, the NF-κB translocation ratio, and the morphological measure were averaged across the four culture replicates with the arithmetic mean. The crosshair is ±1 standard error of that mean. The figure was specified in Altair 6.1.0 (Vega-Lite 6.4.1).

### Quantitative Polymerase Chain Reaction

Total RNA was collected cells from iMg and iMac cultured in 24-well plates (P24-1.5P, Cellvis) using the RLT buffer (74134, Qiagen). RNA extraction was performed using the RNeasy Plus kit (74134, Qiagen) according to the manufacturer’s protocol. RNA quantity and purity were determined by a NanoDropOne spectrophotometer (Nanodrop Technologies). RNA (200-250ng) was used to synthesize cDNA using the high-capacity cDNA reverse transcription kit (4368813, Thermo Fisher Scientific). Quantitative PCR (qPCR) was performed on a QuantStudio 7 (Thermo Fischer) using gene-specific, Taqman primers obtained from Thermo Fischer. The genes TMEM119, AIF1, P2RY12, SALL1, CCR5, CD68, and MKI67 were amplified from cDNA by quantitative PCR. The gene 18s was used as a housekeeping gene during all qPCR assays.

### Generation of HIV_ADA_ Viral Stocks

Viral stocks were generated by infecting CEM-SS cells with a blood-derived, M-tropic ^34^ strain of HIV (HIV_ADA_), as previously described^30,35^. Briefly, CEM-SS cells were cultured in T175 flasks in RPMI-1640 supplemented with 10% fetal bovine serum (FBS), 1% penicillin– streptomycin (PenStrep) HEPES buffer (10 mM). Each culture was inoculated with a single vial of HIV_ADA_ from an existing stock. Initial HIV_ADA_ stocks were obtained through the NIH HIV Reagent Program, Division of AIDS, NIAID, NIH: Human Immunodeficiency Virus-1 ADA, ARP-416, contributed by Dr. Howard Gendelman. Infected cultures were maintained until syncytium formation was observed, typically 15–18 days post-infection. Following the appearance of syncytia, supernatants were collected every 24 hours until viral replication outpaced cell growth and the culture was no longer viable. Cell-free supernatants were collected daily from days 18 to 41 post-infection. Following supernatant collection, supernatants were centrifuged at 1,200 × g to pellet cells and debris. The clarified supernatant was collected, aliquoted into 1 mL volumes, and stored at −80°C for use as viral stocks. Viral stock concentration was determined by quantifying p24 Gag antigen per milliliter using a high-sensitivity HIV-1 p24 AlphaLISA detection kit (Revvity, Waltham, MA), according to the manufacturer’s instructions.

### HIV Inoculation

Mature iMg and iMac cells were plated in 96-well plates and inoculated with HIV_ADA_ (0.1– 20 ng/mL) for 24 hours. Following inoculation, cultures were washed and replenished with complete fresh medium specific to each cell type. Beginning at 3 days post-infection (DPI3), half of the culture supernatant was collected and replaced with an equal volume of fresh medium every 48 hours through DPI9. Supernatants from all wells of the same inoculation condition were pooled, aliquoted, and stored at -80°C for analysis of p24Gag as a measure of viral replication. For studies that included combination antiretroviral therapy (cART), the antiretroviral drugs which comprise the first-line regimen Biktarvy (Bictegravir (BIC), Emtricitabine (FTC), Tenofovir Alafenamide (TAF)) were added to each well along with the fresh media starting after supernatant collection at DPI3. The concentrations used (BIC, 13.5 mM; TAF, 600 nM; FTC, 7 mM) are found at peak plasma levels (C_max_) in PLWH^36–39^. Supernatants were analyzed for p24Gag using AlphaLISAs performed according to the manufacturer’s protocol (Revvity). At DPI 9, plates were fixed with 4% PFA at 4°C for 20 minutes for high content imaging analysis for quantification of p24.

### High Content Image Analysis of HIV Infection

To analyze the percentage of p24+ cells, a 2-channel colocalization was performed using the sample parameters (**Supp. Table 3**). The CMDR stain (Channel 1) was used to identify cells in the well, and the anti-P24 stain (Channel 2) was used to identify p24+ cells. Both channels were gated by excluding border objects, object area, and object total/variant intensity (as needed). A region of interest (ROI) was formed based on the colocalization between CMDR and anti-p24 (Target1). Cell-level data was exported on ROI_A_ Target1_ObjectVarInten and used to calculate the %p24+ cells from the p24+ cell counts and total cell counts.

### Bulk RNA sequencing

#### Sample Preparation

RNA obtained from mature iMg and iMacs cultured in 24-well plates was evaluated for purity and integrity at the University of Nebraska Medical Center Genomics Core Facility. RNA purity was assessed using a NanoDrop spectrophotometer (Thermo Scientific, Wilmington, DE) by measuring absorbance ratios. RNA integrity and potential degradation were evaluated using a Fragment Analyzer (Advanced Analytical Technologies, Inc., Ames, IA). Samples with A260/280 ratios ≥ 1.8 and RNA Quality Number (RQN) scores > 8.0 were used for library preparation. Sequencing libraries were generated by University of Nebraska Medical Center NGS Core using 200 ng of total RNA per sample and the Universal Plus mRNA-Seq Library Kit (Tecan Genomics, Redwood City, CA), according to the manufacturer’s protocol. Library fragment size distribution was assessed by analyzing an aliquot of each library on a Bioanalyzer instrument (Agilent Technologies, Santa Clara, CA). Each library was assigned a unique index barcode to enable multiplexing. Indexed libraries were pooled and sequenced on a single SP-200 flow cell using the NovaSeq 6000 platform (Illumina) with a 2 × 100 bp paired-end configuration. Sequencing generated approximately 16–20 million paired-end reads per sample. The sequences will be deposited in NCBI GEO (accession number: in progress).

#### Analysis of RNA-Seq

Raw FASTQ files were processed using the nf-core RNA-seq pipeline (v3.10.1) to generate a gene-level count matrix. Briefly, sequencing reads were quality controlled, filtered, and aligned with the STAR aligner to a custom reference genome comprising hg38 together with the HIV-1 reference sequence (NC_001802.1)^40^. The resulting gene count matrix was imported into Partek® Flow® for downstream analyses. Differential gene expression was performed using DESeq2, and genes meeting a log fold change threshold of 1 and adjusted p value < 0.05 were considered significant^41^. Gene set enrichment analyses were conducted using Gene Ontology (GO) databases, including Hallmark, Biological Process, and Molecular Function, with full data sheets in **Supplementary Table 4 - 7**.

### Single-cell RNA sequencing

#### Sample preparation

Mature iMg cultured in 24-well plates (Cellvis, P24-1.5P) were inoculated with HIV_ADA_ (20 ng/ml) for 24 hours, after which virus was washed off cells. The cells remained in culture till 7 days post-infection, then detached at 37°C using TrypLE (12604013, Thermo Fisher). Detachment was halted by the addition of an equal volume of warm media, and cells were spun at 400xg for 5 minutes. iMg was cryopreserved in 90% FBS 10% DMSO at a minimum concentration of 1,000,000 cells/mL at -80°C > 1 day (Mr. Frosty; Thermo Fischer Scientific), then transferred to a liquid nitrogen tank.

#### Library preparation

Post-sorting, cell suspensions were concentrated to approximately 1,000 cells/µL, and viability and concentration were assessed by trypan blue exclusion. Cells were processed using the 10x Genomics (Pleasanton, CA, USA) Chromium 3’ v3 GEM kit, targeting a recovery of 8,000 cells. Based on the target cell number, the appropriate volume of cell suspension was loaded onto a 10x Genomics Chromium GEM Chip and processed on the Chromium Controller for GEM generation and cell capture. Briefly, cells are partitioned via microfluidics into nanoliter-scale droplets together with Single Cell 3’ Gel Beads carrying barcoded oligonucleotides containing a unique molecular identifier (UMI). Within each droplet, cells are lysed and RNA is reverse transcribed using barcoded primers containing a unique molecular identifier (UMI). Barcoded cDNA is subsequently amplified, fragmented (∼200 bp), and prepared into sequencing-ready libraries via adapter ligation and sample indexing, per the manufacturer’s version 3 chemistry protocol. Libraries were sequenced on Illumina (San Diego, CA, USA) NextSeq 550 and NovaSeq 6000 platforms. Sequencing data is deposited in NCBI GEO (accession number: in progress).

#### Bioinformatics

Sequenced samples were processed using the 10× Genomics Cell Ranger pipelines (v7.1.0). scRNA-seq reads were demultiplexed and aligned to a customized GRCh38 human reference containing an additional contig representing the HIV genome. To reduce ambiguous alignment between the highly similar 5′ and 3′ long terminal repeats (LTRs), the 3′ LTR was removed from the HIV reference sequence while the 5′ LTR was retained. Summary statistics from the Cell Ranger gene counting output are provided in **Supp Table 4**.

All downstream analyses were performed in Python (v3.12.7) using Scanpy (v1.10.3)^42^. Raw count matrices from microglia and macrophage samples were imported and merged. Features expressed in at least three cells were retained. Initial quality control (QC) involved filtering cells based on unique molecular identifiers (UMIs), gene counts, and mitochondrial transcript percentage. Following QC, data were normalized to a total count of 10,000 per cell using sc.pp.normalize_total and log-transformed using sc.pp.log1p. Highly variable genes were identified using sc.pp.highly_variable_genes and the dataset was restricted to highly variable genes. Total UMIs per cell and mitochondrial transcript percentage were regressed out using sc.pp.regress_out, after which expression values were scaled to unit variance using sc.pp.scale. Principal component analysis (PCA) was performed via sc.tl.pca, and a neighborhood graph was constructed using sc.pp.neighbors with 10 nearest neighbors and the first 15 PCs. UMAP was computed using sc.tl.umap for dimensionality reduction and visualization. Clustering was performed using the Leiden algorithm (sc.tl.leiden) with flavor=“igraph”, n_iterations=2, and resolution=0.5. Cluster-specific differentially expressed genes (DEGs) were identified using the Wilcoxon rank-sum test implemented in sc.tl.rank_genes_groups. DEGs were defined as adjusted p-value < 0.05 and absolute log fold change > 0.5.

To compare microglia and macrophages, differential expression analysis was performed using a t-test implemented in sc.tl.rank_genes_groups, with microglia designated as the reference group. Genes with a test statistic of zero were excluded, and the remaining genes were ranked by the test statistic and subjected to preranked gene set enrichment analysis (GSEA) using GSEApy (v1.1.3)^43^. Enrichment was performed using the MSigDB Hallmark and Gene Ontology (Biological Process) gene sets, using minimum and maximum gene set sizes of 5 and 500, respectively. Significantly enriched gene sets were defined by an FDR q-value < 0.05, and fold change of 1.

### Multiplex Cytokine Analysis

For cytokine analysis, supernatants were assessed using the MSD V-Plex Human Neuroinflammation panel (Cat #K15210D-1, Meso Scale Diagnostics, LLC, Rockville, Maryland, USA), assessing expression of 37 analytes. Samples were run in duplicate according to the manufacturer’s protocol. All plates were imaged on the MESO QuickPlex SQ 120MM and analyzed using MSD Discovery Workbench 4.0 Software (Meso Scale Diagnostics, LLC, Rockville, Maryland, USA).

### Statistical Analysis

Statistical analyses for all experiments except the sequencing studies were performed using GraphPad Prism. Data distributions were assessed for skewness, normality, and lognormality using the Shapiro-Wilk, Kolmogorov-Smirnov, and D’Agostino & Pearson tests. Outliers, presumed to be technical artifacts, were identified and excluded using the ROUT test (Q = 0.1%). p24 AlphaLISA data were log_10_-transformed prior to analysis. Gene expression data were analyzed as 2^−ΔCT^ prior to display as 2^−ΔΔCT^. Group comparisons were made using paired t-tests (for normally distributed data with equal variances) or Wilcoxon tests (for non-normal data). Multiple group comparisons were assessed using one-way ANOVA, two-way ANOVA, or mixed-effects analysis, followed by appropriate post hoc tests (Sidak’s, Dunnett’s, or Tukey’s) as necessary. Data are presented as mean ± SEM, with p < 0.05 considered significant. Sequencing analyses were performed as described in specific methods sections.

Nonlinear curve analysis was performed to quantify the differences in NF-kB response to distinct TLR agonists. In these analyses, the maximal response (E_max_), sensitivity (EC_50_), and cooperativity (HillSlope) was assessed. E_max_ represents the maximal NF-κB response to LPS, with a higher E_max_ reflecting a greater magnitude of NF-κB activation. EC_50_ is the concentration required to achieve 50% of the maximal response and measures cellular sensitivity to the stimulus, i.e. a lower EC_50_ indicates more sensitivity. Hillslope describes the slope of the dose-response curve and indicates how sharply each cell type responds to incremental changes in stimulus concentration.

## Results

### Generation of syngeneic iPSC-microglia and macrophages

To effectively model and compare syngeneic microglia and macrophages, we generated both iPSC-derived microglia (iMg) and CAMs (iMac) from the same induced pluripotent stem cell (iPSC) lines. Common myeloid progenitors (CMP) from the same iPSC line were patterned into microglia-like cells (iMg) over 11 days in the presence of IL-34, M-CSF, TGF-β^31^, or CAM-like cells (iMac) over 7 days in the presence of M-CSF **(Fig. 1A)**. At maturity these cells develop the phenotype associated with their respective cell types, with iMg becoming more ramified and iMac becoming larger over the course of the differentiation **(Fig. 1B**). Both cell types were positive for identity markers via immunofluorescence imaging. iMac expressed CD68 and CCR5, while lacking P2RY12, and iMg expressed P2RY12, TMEM119, and IBA1 **(Fig. 1C)**. We further characterized this at the genomic level using qPCR, confirming expression of TMEM119, P2RY12, and IBA1, but not SALL1 in iMg **(Fig 1D)**, while the iMacs express CD68 and IBA1 **(Fig 1E)**, but not P2RY12 **(Fig 1C)**. We then examined the expression of CCR5, as a marker of mature macrophages which is also a necessary coreceptor responsible for HIV entry in myeloid cells^44^. According to our qPCR data, both cell types express CCR5, with no difference in expression levels **(Fig 1F)**. We also noticed changes in cell proliferation between the cell types, with more rapid growth in iMac than iMg cultures. At maturity - DIV11 for iMg and DIV7 for iMac - we measured cell division marker MKI67 expression. As we anticipated from observations of culture growth, iMacs have a significantly greater expression of MKI67 compared to iMg **(Fig. 1G)**. This suggests that iMacs have a greater proliferative capacity, an important consideration to note for downstream assays.

To characterize baseline transcriptional differences between microglia and macrophages, bulk RNA sequencing was used. RNA from mature syngeneic iMg (DIV11) and iMacs (DIV7) was isolated from 2-3 independent lines and analyzed. iMg showed 2,111 genes differentially expressed compared with iMacs. Of these, 1,056 genes were upregulated and 1,055 genes downregulated, i.e. log_2_ FoldChange > 0.5 **(Fig. 2A)**. A full list of the differentially expressed genes is in **Supp. Table 5**. Pathway enrichment analysis was performed using the molecular signatures database (MsigDB) Hallmark and Gene Ontology (GO) databases. In the Hallmark dataset (**Supp. Table 6**), 15 pathways were differentially regulated in iMg compared to iMacs, including a number of inflammatory response pathways including TNF-α signaling via NF-κB, IL6-JAK-STAT3 signaling, interferon gamma response, and interferon alpha response **(Fig. 2B)**. GO Biological Processes (**Supp. Table 7**) analysis identified 139 differentially enriched pathways between the two cell types, including antigen presentation, chemokine signaling and response, inflammatory response, type II interferon signaling, chemotaxis, defense response to bacterium and lipopolysaccharide, and regulation of NF-κB activity **(Fig. 2C)**. In the GO Molecular Function category (**Supp. Table 8**), 19 pathways were differentially expressed, including immune receptor activity, peptide antigen binding, chemokine and cytokine receptor activity, and polysaccharide binding **(Fig. 2D)**. These data indicate substantial baseline differences in inflammatory and immune-related transcriptional programs between iMg and iMacs.

**Figure 2:**
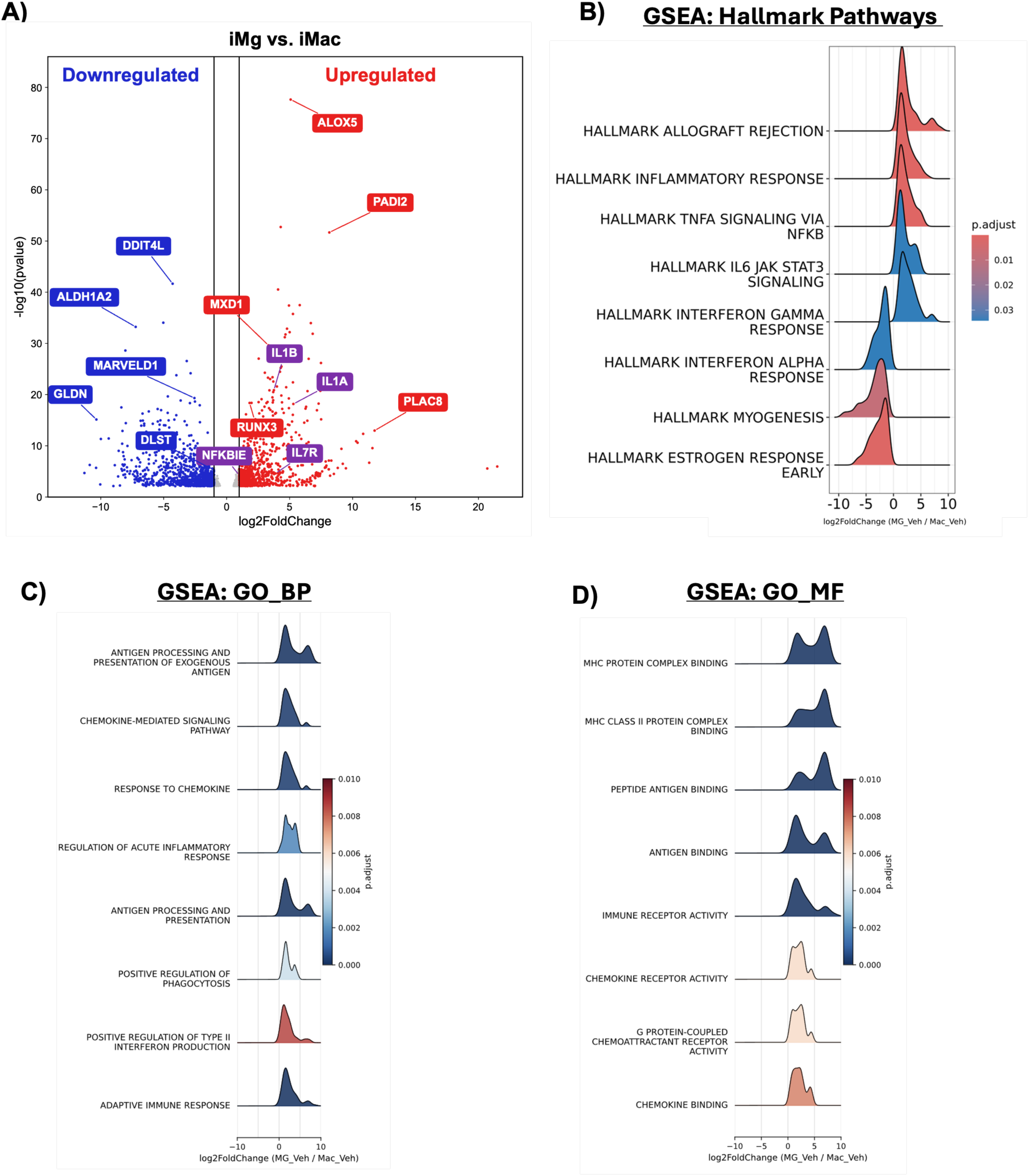
RNA sequencing identifies baseline differences in inflammatory response between iMg and iMac. (A) Volcano plot showing 2,111 differentially expressed genes in iMg compared to iMacs, including 1,056 upregulated and 1,055 downregulated genes (fold_change > 0.5, p_value < 0.05). Gene set enrichment analysis of differentially expressed genes using: (B) Molecular Signatures Database (MsigDB) Hallmark pathways, (C) Gene Ontology Biological Processes (GO_BP), and (D) Gene Ontology Molecular Functions (GO_MF).

### Syngeneic iMg and iMacs show differential NF-κB responses to TLR stimulation

Macrophages detect pathogen- and damage-associated molecular patterns (PAMPs and DAMPs) through a broad range of pathogen recognition receptors. Among these, Toll-like receptors (TLRs) detect diverse pathogen-associated signals across bacteria, viruses, fungi, and stressed cell^45^. Engagement of TLR activates downstream signaling pathways that converge on several central transcription factors, including NF-κB, a master regulator of inflammation^46^. To compare cell type specific differences in NF-κB-mediated TLR responses of microglia and macrophages to discrete inflammatory stimuli, gene expression of TLR 2, 3, 4, and 7 was first examined in both cell types. Both iMg and iMac expressed all the TLRs, with no significant differences in TLR expression between the two cell types. The relative expression levels were TLR2 > TLR7 = TLR4 > TLR3 in both iMac and iMg **(Fig. 3A**).

**Figure 3:**
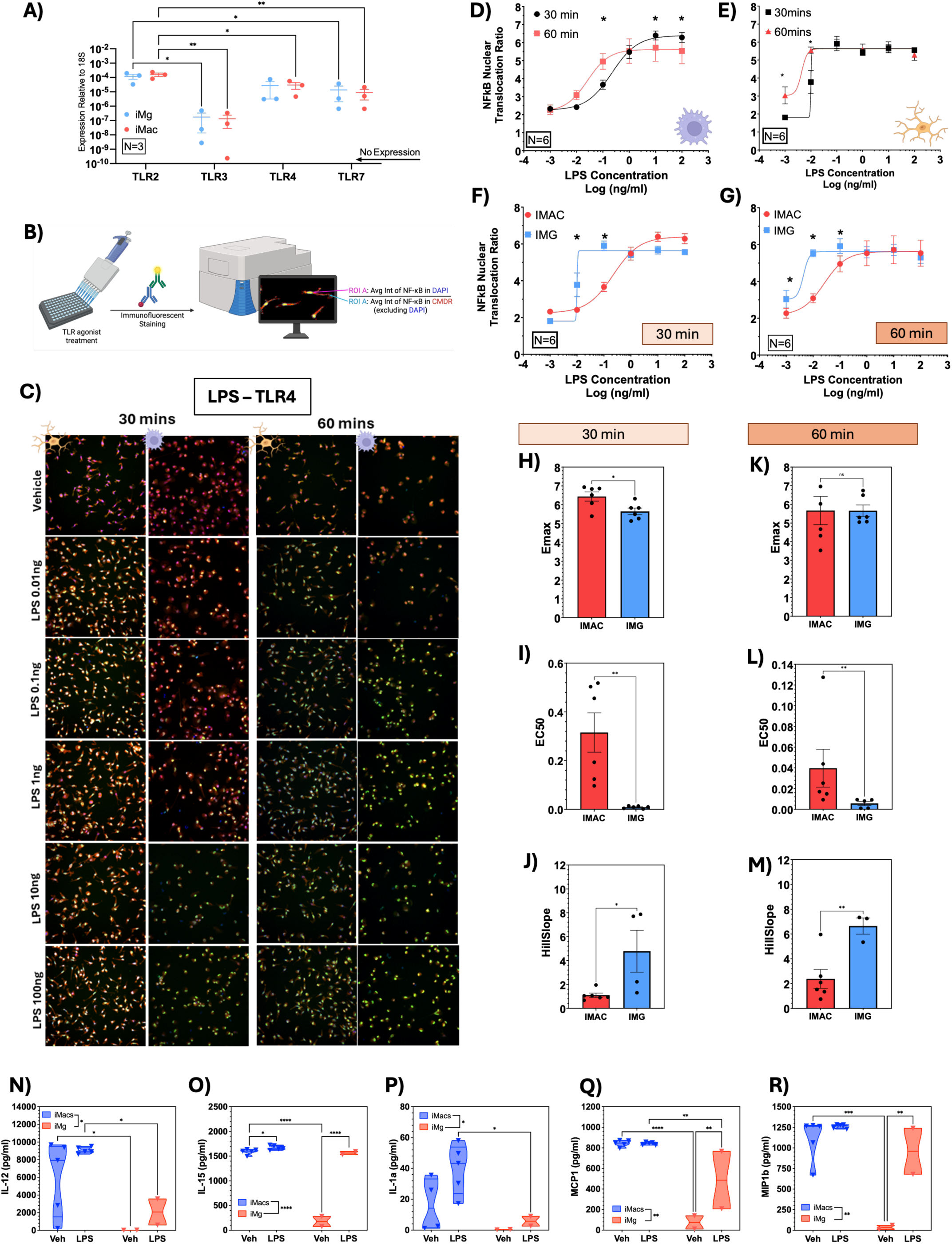
Differential NF-κB response of syngeneic iMg and iMacs to TLR-4 (LPS) mediated stimulation, linked to differential cytokine and chemokine secretion. (A) Gene expression of TLR2, TLR3, TLR4, and TLR7 in iMg and iMacs. Two-way ANOVA [F(3,16) = 12.66], Holm-Sidak post hoc test. *P < 0.05, **P < 0.005. (B) Schematic of the workflow for the NF-κB translocation assay in iMacs and iMg (created in Biorender.com). (C) Representative high-content images of iMg and iMacs treated with LPS (0–100 ng/mL) and stained for NF-κB (green), cell body (red), and nucleus (blue), scale bar represents 20uM. Nonlinear regression (sigmoidal curve fit) of NF-κB nuclear translocation in response to LPS at 30 and 60 minutes in: (D) iMacs, Two-way ANOVA [F(5,25) = 89.26], Holm-Sidak post hoc test; (E) iMg, Two-way ANOVA [F(5,25) = 85.69], Holm–Sidak post hoc test. *P < 0.05. Direct comparison of iMg and iMac responses at: (F) 30 minutes, Two-way ANOVA [F(5,25) = 139.6], Holm–Sidak post hoc test; (G) 60 minutes, Two-way mixed-effects ANOVA [F(2.245,11.23) = 40.56], Holm-Sidak post hoc test. *P < 0.05. (H-J) Quantification of Emax, EC50, and HillSlope parameters at 30 minutes. (K-M) Quantification of Emax, EC50, and HillSlope parameters at 60 minutes. T-test, *P < 0.05. Mature iMg and iMacs were treated with vehicle or LPS (10 ng/mL) for 24 hours. Supernatants were collected and analyzed using a multiplex cytokine panel. Differential baseline and LPS-induced secretion profiles were identified between iMg and iMacs for (N) IL-12, (O) IL-15, (P) IL-1α, (Q) MCP-1 and (R) MIP-1β.

To examine differences in TLR response, syngeneic iMac and iMg were treated with different TLR agonists, and NF-κB translocation was evaluated as a readout. The TLR agonists used were lipopolysaccharide (LPS, TLR4 agonist), Pam3CSK4 (Pam, TLR2 agonist), polyinosinic-polycytidylic acid (Poly I:C, TLR3 agonist), and imiquimod (Imi, TLR7 agonist). All TLR agonists were used at a range of concentrations from 0.001 - 100ng/ml (10^-3^ – 10^2^ng/mL), fixing cells 30 or 60 minutes post-treatment. Following fixation, cells were stained for cell membrane (Red, HCS Cell mask deep red), nuclei (Blue, DAPI) and NF-κB (Green, Alexafluor 488). Stained cells were imaged and evaluated for the ratio of NF-κB in the nucleus (amount of green NF-κB in blue nuclei) vs. NF-κB in the cytoplasm (amount of green NF-κB in red cytoplasm-excluding blue DAPI). The ratio of nuclear NF-κB: cytoplasmic NF-κB is referred to as the nuclear translocation ratio (**Fig. 3B**) and used as a measure of NF-κB activity^33^.

#### TLR4 Response

To assess changes the response to TLR4 activity, cells were treated with LPS, a commonly used pathogen-associated molecular pattern and endotoxin from the outer membrane of Gram-negative bacteria such as *E. coli*, *Salmonell*a, and *Pseudomonas*^47,48^. In iMacs, increasing concentrations of LPS elicited a graded sigmoidal increase in NF-κB translocation at both 30 and 60 mins **(Fig. 3C, 3D)**, consistent with our published work examining NF-κB in human monocyte-derived macrophages^33^. At higher LPS concentrations (10 – 100ng/ml), NF-κB translocation was greater at 30 mins than at 60 mins **(Fig. 3D)**. However, the 60-minute curve displayed a leftward shift, with significant activation detectable at 0.01ng/ml, indicating increased sensitivity to lower LPS concentrations over time, despite reduced maximal responses at later time points (**Fig. 3D**). In contrast, rather than a graded curve, NF-κB activation in iMg followed a switch-like pattern in which even the lowest LPS concentrations (0.01 ng/ml) produced near maximal NF-κB translocation (∼5.7 nuclear ratio versus ∼1.7 at baseline) **(Fig. 3C, 3E)**. The 60 min timepoint showed a diminished NF-κB response at lower concentrations (0.001ng/ml) compared to 30 mins but starting at 0.1ng/ml, there were no significant concentration dependent differences in iMg between the timepoints.

Comparing the responses between the iMac and iMg at each timepoint **(Fig. 3F, G)**, showed that iMg exhibited significantly greater NF-κB translocation than iMacs at lower concentrations (0.01 and 0.1 ng/ml) at 30 min. However, at higher concentrations, the responses converged **(Fig. 3F)**. This trend was maintained at 60 mins, with iMg showing a relatively greater sensitivity at low concentrations that became similar to iMac sensitivity at concentrations above 1 ng/mL **(Fig. 3G)**. Nonlinear curve analysis showed that at 30 mins, iMacs showed a higher E_max_ and EC_50_ than iMg, indicating the iMacs have a greater maximal capacity for TLR-4 mediated NF-κB activation, but reduced sensitivity to LPS **(Fig. 3H, I)**. iMg also displayed a significantly steeper Hillslope, consistent with a more switch-like activation response **(Fig. 3J)**. By 60 mins, assessment of the non-linear curve fit showed that E_max_ was comparable between the cell types, yet iMacs continued to have a greater EC_50_, reflecting their lower sensitivity to LPS, while iMg retained the higher Hillslope, indicating a more binary response pattern **(Fig. 3K-M)**. The EC_50_ values for both cell types were lower at 60 mins than at 30 mins, indicating a time-dependent increase in LPS responsiveness. Overall, these data suggest that microglia are more sensitive to LPS than macrophages and engage NF-κB in an ultrasensitive, switch-like manner, whereas macrophages mount a more graded response with higher maximal amplitude but reduced sensitivity.

To compare the broader responses to TLR4 agonist stimulation between these cell types, we assessed the production and secretion of cytokines and chemokines 24hrs post-treatment with LPS at 10ng/mL, a concentration and timepoint previously optimized in our laboratory^33^. This was done using the V-Plex Human Neuroinflammation panel from Mesoscale assessing the release of 37 different biomarkers associated with neuroinflammation. Our data suggested that iMacs and iMg mostly respond similarly in terms of biomarker production induced by LPS (data not shown). However, we observed differential release of IL-12, IL-15, MIP-1β, MCP1 and IL-1α when comparing iMacs and iMg. At baseline, iMacs exhibited a more pro-inflammatory secretory phenotype, with higher levels of pro-inflammatory cytokines (IL-12, IL-15) and chemokines (MIP-1β, MCP-1, IL-1α) compared to iMg **(Fig. 3N-R)**. These differences disappear following LPS stimulation, where iMg showed increased levels of all cytokines, with significant increases in IL-15, MIP-1 β and MCP-1. While there was some increase in cytokine release in iMac, the magnitude of the changes was small relative to the iMg and only reached significance for IL-15. Notably, IL-1α, IL-12, and IL-15 are involved in immune-cell activation, regulation, and coordination, whereas MIP-1β (CCL4) and MCP-1 (CCL2) regulate immune-cell trafficking and recruitment. In particular, MCP-1 promotes the recruitment of circulating monocytes to sites of inflammation, where they can differentiate into macrophages. These processes are relevant to HIV pathogenesis, as CNS macrophages drive to viral persistence in that compartment.

#### TLR4-Induced Morphological Changes in iMg and iMac

In addition to activating NF-κB, activation of TLR4 by LPS is often accompanied by rapid changes in cell morphology^49,50^. Although these morphological changes do not directly establish the magnitude of inflammatory signaling, they provide an additional phenotypic measure of cellular responses. Microglial morphology especially has been extensively investigated as an indicator of changes in cellular state and reactivity. Under homeostatic conditions, microglia typically exhibit a ramified morphology characterized by an elaborate network of processes, whereas activated microglia may exhibit process retraction and a more rounded or amoeboid morphology. These morphological features are more commonly characterized *in vivo* and in 3D models, but 2D microglial cultures also retain measurable morphological responses to inflammatory stimuli. In contrast, the morphological dynamics of macrophages have received comparatively less attention^51,52^. To characterize morphological responses in this system, high-content imaging to quantify morphological changes in both iMg and iMac following LPS stimulation for 30 or 60 minutes, broadly showing that in both cell types, increasing concentrations of LPS drove lower perimeter-to-area (P2A) and length-to-width (LWR) ratios **(Fig. 4A)**.

**Figure 4:**
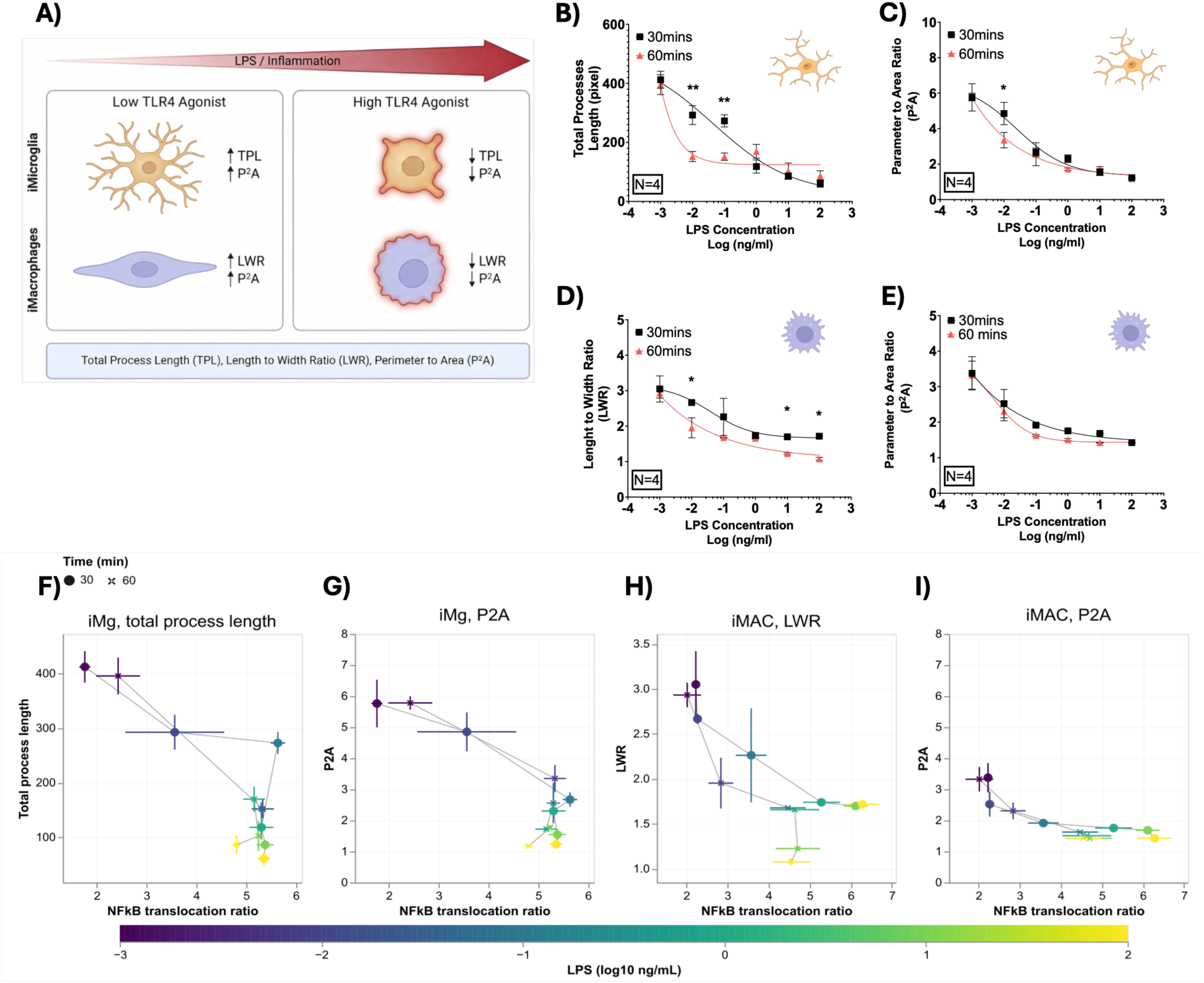
TLR4 modulation by LPS is correlated with morphology alterations of syngeneic iMg and iMac. (A) Schematic illustrating LPS-induced morphological remodeling in iMg and iMac, accompanied by increased NF-κB nuclear translocation. Increasing LPS concentrations are associated with reduced total process length (TPL) and perimeter-to-area ratio (P2A) in iMg, and reduced length-to-width ratio (LWR) and P2A in iMac (created with BioRender.com). (B,C) Quantification of TPL and P2A in iMg following exposure to increasing concentrations of LPS (Two-way ANOVA, Holm–Sidak post hoc test, *P < 0.05, **P < 0.01). (D,E) Quantification of LWR and P2A in iMac following exposure to increasing concentrations of LPS (Two-way ANOVA, Holm–Sidak post hoc test, *P < 0.05). (F,G) Correlation between NF-κB nuclear translocation and morphological parameters (TPL and P2A, respectively) in iMg across increasing LPS concentrations at 30 and 60 min. (H,I) Relationship between NF-κB nuclear translocation and morphological parameters (LWR and P2A, respectively) in iMac across increasing LPS concentrations. Circles represent 30-min measurements and crosses represent 60-min measurements. Data are presented as culture means ± SEM, with crosshairs indicating ±1 SEM. Color scale represents log10-transformed LPS concentration (ng/mL).

Specifically, in iMg, increasing LPS concentrations were associated with reductions in total process length (TPL) **(Fig. 4B)** and P2A **(Fig. 4C)**, consistent with process retraction and a shift toward a more compact cellular morphology. The concentration–response relationship for TPL differed between the two time points. At 30 min, TPL exhibited a more gradual, approximately linear decrease with increasing LPS concentrations, whereas at 60 min, the response was more sigmoidal, with concentrations above 0.01 ng/mL producing relatively similar TPL values **(Fig. 4B, C)**. These findings suggest that microglial morphological responses evolve over time, with greater separation across LPS concentrations at the earlier time point and a more pronounced plateau at the later time point. LPS stimulation also induced measurable morphological changes in iMac. Whereas unstimulated iMac retained a relatively elongated morphology, LPS exposure was associated with a rounder cellular appearance. At both 30 and 60 min, increasing LPS concentrations were associated with comparable reductions in length-to-width ratio (LWR) and P2A, suggesting that these morphological parameters exhibited broadly similar concentration-dependent response patterns across the two time points **(Fig. 4D, E)**.

Additionally, LPS-induced morphological changes were accompanied by increased NF-κB nuclear translocation in both cell types. To further characterize the relationship between NF-κB activation and morphological remodeling, we performed regression analyses across LPS concentrations at both time points (30 and 60 min). In iMg, total process length was strongly associated with NF-κB translocation at both time points. Notably, at LPS concentrations above 0.01 ng/mL, NF-κB translocation approached a plateau, whereas morphological remodeling continued with increasing LPS concentrations, as reflected by further reductions in total process length. This dissociation suggests that the extent of morphological remodeling is not necessarily proportional to the magnitude of NF-κB nuclear translocation. This pattern was generally consistent between the 30- and 60-min time points **(Fig. 4F, G)**. In iMacs, both P2A and LWR showed associations with NF-κB translocation. However, their temporal patterns differed slightly. LWR exhibited modest differences between the 30- and 60-min responses, paralleling the time-dependent variation observed in NF-κB translocation, whereas P2A displayed a more consistent relationship across the two time points **(Fig. 4H, I)**.

#### TLR2 Response

To assess changes in the NF-κB response to TLR2 activity, iMg and iMac were treated with Pam3CSK4 **(Fig. 5A)**, a synthetic tripalmitoylated lipopeptide that acts as a potent agonist of the TLR2 receptor^53,54^. Pam3CSK4 mimics the acylated amino terminus of bacterial lipoproteins found in both Gram-positive and Gram-negative bacteria, including pathogens such as *Leptospira* and *Mycobacterium*. In response to TLR2 activation, iMacs exhibited similar dose response curves at both 30 and 60 min after treatment, showing a modest but significantly response at 30 mins for 0.1 and 1 ng/ml **(Fig. 5B)**. In contrast, iMg displayed a distinctive response pattern, characterized by a sigmoidal graded curve at both 30 and 60 min (**Fig. 5C**), similar to the iMac response to LPS. In contrast to iMac, iMg exhibited greater NF-κB activation at 60 min compared to 30 min, particularly at 0.1 and 1ng/ml Pam3CSK4. The leftward shift in the curve at 60 min also indicates increased sensitivity of iMg to Pam3CSK4 with prolonged stimulation. Direct comparison of iMac and iMg responses at 30 min showed that iMacs exhibited a leftward shift and a higher NF-κB response **(Fig. 5D)**, with significant increases at 0.1 and 1ng/ml Pam3CSK4. By 60 mins, these differences were no longer observed, with both cell types displaying overlapping response curves and no significant differences at any concentration **(Fig. 5E)**. Non-linear analyses of iMacs showed a trend toward a higher E_max_ and a significantly lower EC_50_ **(Fig. 5F,G)** relative to iMg at 30 min, but by 60 mins differences in E_max_ and EC_50_ were no longer present **(Fig. 5I, J)**. Hillslope values were comparable between the two cell types at both time points, suggesting NF-κB similar activation dynamics of NF-κB **(Fig. 5H, K)**. These data indicate that iMacs mount a stronger and more sensitive NF-κB response to Pam3CSK4 than iMg after a short period of exposure, whereas longer stimulation results in convergence of responses between the two cell types.

**Figure 5:**
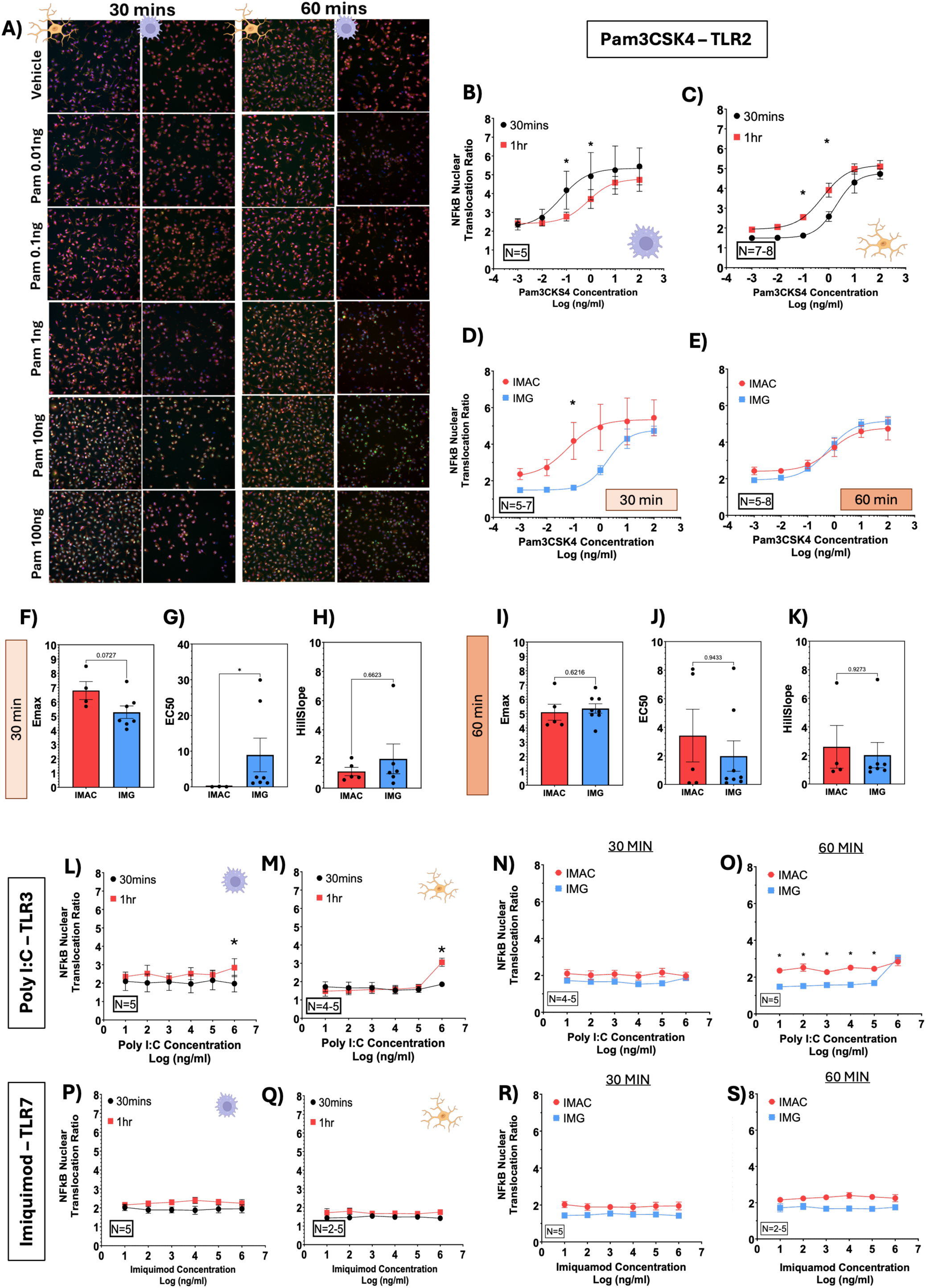
Differential NF-ΚB response of syngeneic iMg and iMacs to TLR-2 (Pam3CSK4), TLR-3 (Poly I:C) and TLR-7 (Imiquimod) mediated stimulation. (A) Representative high-content images showing NF-κB translocation in response to TLR2 agonist, Pam3CSK4 (0–100 ng/mL), 30 and 60 minutes post treatment. Scale bar shows 20uM. Nonlinear regression analysis of NF-κB translocation in response to Pam3CSK4 (0–100 ng/mL) at 30 and 60 minutes in: (B) iMacs, Two-way ANOVA [F(5,20) = 11.17], Holm-Sidak post hoc test; (C) iMg, Two-way mixed-effects ANOVA [F(5,35) = 71.03], Holm-Sidak post hoc test. *P < 0.05. Comparison between cell types at: (D) 30 minutes, Two-way mixed-effects ANOVA [F(5,30) = 33.89], Holm-Sidak post hoc test; (E) 60 minutes, Two-way mixed-effects ANOVA [F(5,55) = 45.21], Holm-Sidak post hoc test. *P < 0.05. (F-H) Quantification of Emax, EC50, and HillSlope at 30 minutes. (I-K) Quantification of Emax, EC50, and HillSlope at 60 minutes. T-test, *P < 0.05. NF-κB translocation in response to Poly I:C (0–100 ng/mL) at 30 and 60 minutes in: (L) iMacs, Two-way ANOVA [F(5,35) = 36.16], Holm-Sidak post hoc test; (M) iMg, Two-way mixed-effects ANOVA [F(2.448,19.10) = 1.424], Holm-Sidak post hoc test. *P < 0.05. Comparison between iMg and iMac responses to Poly I:C at: (N) 30 minutes, ANOVA, P = 0.1659; (O) 60 minutes, Two-way mixed-effects ANOVA [F(5,40) = 33.85], Holm-Sidak post hoc test. *P < 0.05. NF-κB translocation in response to Imiquimod (0–100 ng/mL) at 30 and 60 minutes in: (P) iMacs, Two-way ANOVA, P = 0.1939; (Q) iMg, Two-way mixed-effects ANOVA, P = 0.1683. Comparison between iMg and iMac responses to Imiquimod at: (R) 30 minutes, ANOVA, P = 0.1339; (S) 60 minutes, Two-way mixed-effects ANOVA [F(1,4) = 12.87], Holm-Sidak post hoc test. *P < 0.05.

#### TLR3 and TLR7 Responses

To assess changes in NF-κB in response to TLR3 activity, iMg and iMac were treated with poly I:C. This synthetic analog of double-stranded RNA (dsRNA) stimulates the TLR3 receptor, mimicking infection by viruses that contain dsRNA, including respiratory syncytial virus (RSV), SARS-CoV, and Chikungunya virus^48,53^. Both iMg and iMacs treated with poly I:C for 30 and 60 min showed an increase in NF-κB nuclear translocation only at the highest concentration of poly I:C (100 ng/ml) and only after a 60 minute stimulation **(Fig. 5L,M)**. Direct comparison between iMacs and iMg revealed no significant differences in NF-κB translocation at 30 mins, although iMac showed a significantly higher - although still quite low - NF-κB response at 60 minutes **(Fig. 5N,O)**. These data suggest that macrophages may be more responsive to dsRNA than microglia, but that both cell types exhibit limited responsiveness to TLR3 stimulation.

To assess changes in NF-κB in response to TLR7 activity, iMg and iMac were treated with the synthetic imidazoquinoline amine imiquimod, a specific TLR7 agonist^55^. TLR7 recognizes viral single-stranded RNA, including that from pathogens such as human papillomavirus (HPV)^48^. Treatment with imiquimod did not induce NF-κB nuclear translocation in either iMacs or iMg at 30 or 60 minutes **(Fig. 5P,Q)** and comparison between the two cell types showed no differences in NF-κB activation at either time point **(Fig. 5R,S)**. These findings suggest that, at least in iPSC-derived cells, TLR7 stimulation does not trigger detectable NF-κB translocation in either microglia or macrophages.

### Syngeneic iMg and iMacs exhibit distinct susceptibility to HIV replication and differential responses to antiretroviral drugs

Macrophages in the CNS, including both CAMs and microglia, are central to the development of neuroHIV^23,56^. These long-lived CNS macrophages sustain persistent viral infection, producing low levels of virus and driving chronic immune activation. These macrophages, specifically microglia, are also thought to comprise the majority of the HIV reservoir in the CNS^57^. Differences in the responses of these distinct myeloid populations to both HIV and antiretroviral therapy are not clear, therefore we examined differences in susceptibility to HIV infection and antiretroviral drug (ARV) effectiveness our syngeneic iMg and iMac populations. Both iMg and iMacs were inoculated with an M-tropic strain of HIV (HIV_ADA_) at concentrations ranging from 0.01ng– 20ng - p24/mL for 24 hours to establish the parameters of infection in these cells. Cultures were washed and given fresh media after a 24 hr inoculation, then supernatants were collected at 3, 5, 7, and 9 days post infection (dpi) and replaced with fresh media containing the antiretrovirals comprising the first line ART regimen Biktarvy. Supernatants were quantified for p24 to assess viral replication, and cultures were fixed at 9 dpi to assess the percentage of the culture that was infected with HIV **(Fig. 6A)**. Both iMg and iMac showed dose- and time-dependent increases in p24 secretion, with curve trends indicating an increasing capacity for viral replication over time. In iMac the p24 secretion may have plateaued at higher concentrations of HIV at 7 and 9 dpi **(Fig. 6B, C)**.

**Figure 6:**
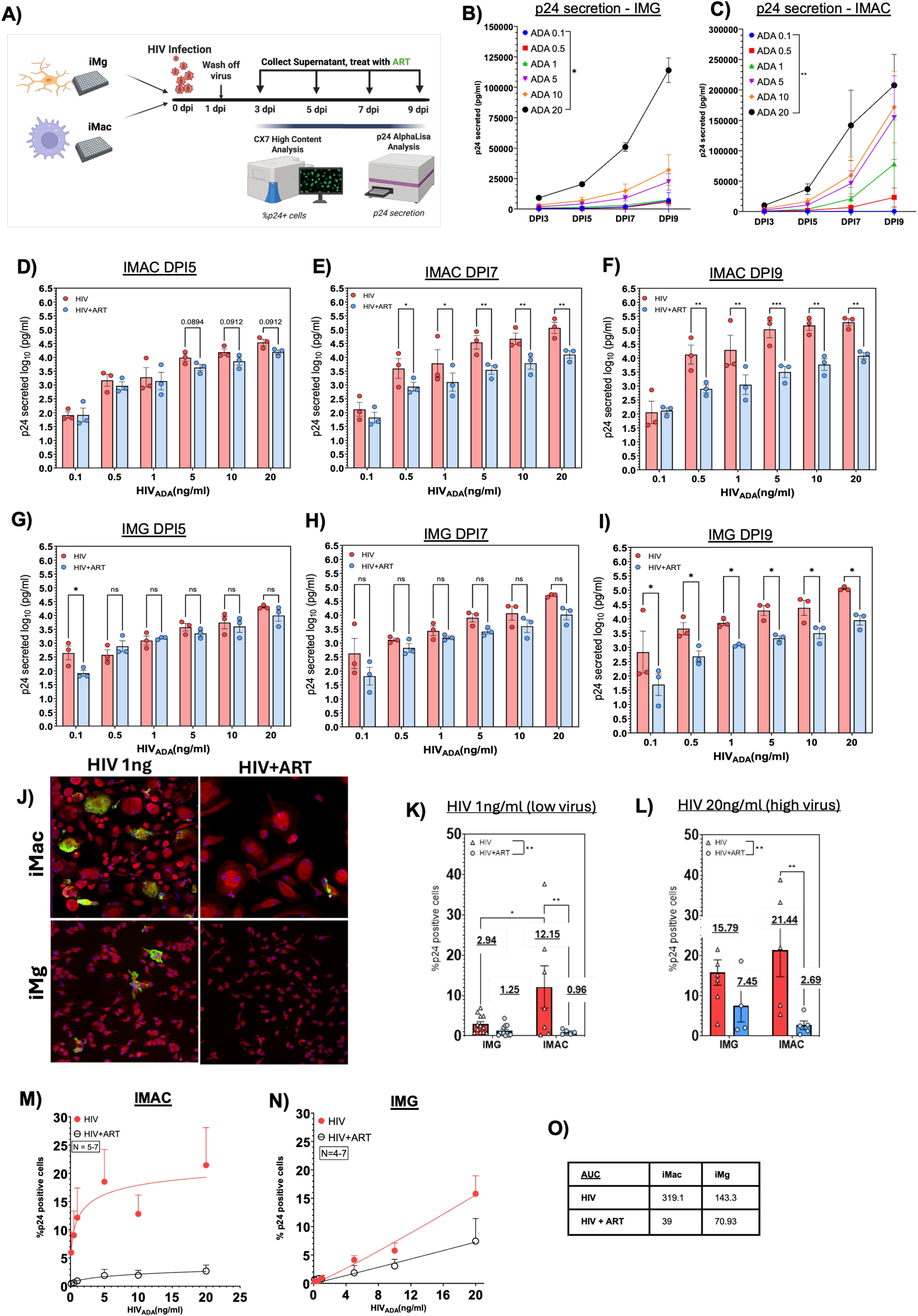
Microglia and macrophages exhibit distinct patterns of HIV replication and differential responses to cART. (A) Mature iMg and iMacs were inoculated with HIV_ADA_ (0–20 ng/mL p24) for 24 hours. Supernatants were collected every two days through day 9 post-infection (DPI9). cART treatment was initiated at DPI3. p24 secreted from DPI3-DPI9 in (B) iMg and (C) iMacs across HIV (ADA) 0.1 – 20ng/ml. iMg: Two-way ANOVA [F(1.063,2.127) = 17.26], iMacs: [F(15,30) = 4.457]. Secreted p24 levels measured by AlphaLISA in supernatants from iMacs at DPI5–DPI9 (D-F) and from iMg at DPI5–DPI9 (G–I). Two-way ANOVA with Holm-Sidak post hoc test. *P < 0.05, **P < 0.005, ***P < 0.0005. (J) Representative high-content images of iMg and iMacs inoculated with HIV(1 ng/mL) +/-cART and stained for p24 (green), cell body (red), and nucleus (blue). Quantification of % p24+ cells in iMg and iMac cultures at (K) 1ng-ml (low inoculum) and (L) 20ng/ml (high inoculum) at DPI9. Two-way mixed-effects ANOVA 1ng/ml: [F(1,22) = 8.444]; 20ng/ml: [F(1,7) = 14.06]; Fisher’s LSD post hoc test. *P < 0.05. Nonlinear regression analysis of % p24+ cells across increasing HIV concentrations +/-cART in: (M) iMacs and (N) iMg. (O) Table summarizing calculated AUC values for iMg and iMacs +/-cART.

Next, we examined the differences in the effectiveness of ARV on p24 secretion in iMg and iMacs 5, 7 and 9 dpi, excluding 3 dpi from this analysis because ARV treatment was initiated after supernatant collection that day. In iMacs, ARV treatment showed trends toward decreased p24 secretion at 5 dpi in the higher inoculum but robustly suppressed viral release at 7 and 9 dpi for all but the lowest concentration of HIV inoculum **(Fig. 6D–F)**. In contrast, iMg exhibited minimal ARV responsiveness at 5 and 7 dpi for all inocula, with significant suppression only observed primarily 9 dpi **(Fig. 6G–I)**. We then quantified the percentage of p24+ cells in HIV only and HIV + ARV treated cultures at 9 dpi to evaluate the effect of ARV on viral spread. Analyses were performed at 9 dpi because this was the time point at which both models showed measurable ARV effects and would show the fullest extent of viral spread. Analyses were performed at both low (1 ng/mL) and high (20 ng/mL) viral inoculum, to capture infection dynamics under conditions of limited versus almost high viral exposure.

At 1ng/ml HIV, significantly more iMacs (12.15%) than iMg (2.94%) were p24+. Treatment with ARV reduced the percentage of infected cells in both models, however, the magnitude of reduction was greater in iMacs, significantly reducing the number of infected cells from an average of 12.15% to 0.96%, whereas ARV treatment of iMg led to a non-significant decrease in the number of p24+ cells, from an average of 2.94% to 1.25% **(Fig. 6K, L)**. The trends were similar in response to inoculation with 20 ng/mL HIV. Both cell types showed higher infection levels, with less of an absolute difference between the number of p24+ cells in response to HIV inoculation, although iMacs again exhibited greater susceptibility (21.44%) than iMg (15.79%). Similarly, ARV treatment still reduced the percentage of p24+ cells in both models, but the reduction was much larger in iMacs. In iMacs, the number of p24+ cells dropped significantly from an average of 21.44% to 2.69%, while in iMg, there a smaller, non-significant reduction in the number of p24+ cells from an average of 15.79% to 7.45%. Analyzing the impact of ARV on the number of infected cells using area under the curve (AUC) across the full viral dose range further supported these findings. In the absence of ART, iMacs displayed a higher AUC (319.1) than iMg (143.3), consistent with the greater baseline permissiveness of macrophages to HIV infection **(Fig. 6M–O)**. Following ART treatment, the AUC decreased in iMacs to 39, but only to 70.93 in iMg, indicating that ART was more effective at reducing infection in macrophages than in microglia **(Fig. 6M–O)**. Overall, these results suggesting that human macrophages may be more susceptible to HIV infection than human microglia. They further suggest a relative bias in ART efficacy, with greater suppression of HIV infection in macrophages compared with microglia.

### iMg and iMacs exhibit distinct transcriptional profiles in response to HIV infection

To characterize cell-type specific transcriptional responses to HIV infection, scRNAseq was performed on syngeneic iMacs and iMg, generating RNA from 2 – 3 iPSC lines. Cultures of both iMac and iMg were infected with 20 ng/mL of HIV_ADA_ and collected at 9 dpi, maximizing the number of infected cells to provide the best sample for analysis. UMAP analysis of the combined data set identified distinct transcriptional clusters, with cluster segregation of iMacs and iMg and cell-type specific changes in cluster distribution (**Fig. 7A**). Quantification of cluster composition revealed a greater proportion of iMg in clusters 1 (iMacs: 5.47%; iMg: 38.32%), 4 (iMacs: 3.89%; iMg: 28.40%), and 7 (iMacs: 1.24%; iMg: 7.65%). In contrast, clusters 3 (iMacs: 41.66%; iMg: 0.65%) and 5 (iMacs: 25.92%; iMg: 3.58%) were predominantly composed of iMacs (**Fig. 7B**). Examining HIV transcript expression across the two populations showed that iMacs exhibited a higher proportion of HIV-positive cells than iMg (99.85% vs. 80.26%) at 9 dpi (**Fig. 7C, D)**, consistent with our protein-level measurements of p24. This difference was observed across nearly all clusters, although the magnitude of the difference varied by cluster (**Fig. 7E)**. Clusters 3 and 6 showed the largest differences in HIV expression, with approximately 20-fold and 15-fold higher expression in iMacs, respectively (FDR<0.05). In contrast, cluster 5 was the only population in which HIV expression was modestly higher in iMg, with a 1.08-fold difference. Looking beyond HIV transcripts, 10,882 differentially expressed genes were identified, including 4,062 upregulated and 6,820 genes downregulated in iMacs relative to iMg (**Fig. 7F**). GSEA using the Hallmark gene sets revealed differential enrichment of pathways related to apoptosis, complement, NF-κB signaling, and interferon responses, indicating distinct inflammatory and stress-response programs between iMacs and iMg during HIV infection (**Fig. 7G**). Gene Ontology biological process analysis identified differential regulation of pathways involved in antiviral responses, interferon signaling, and responses to viral genome replication (**Fig. 7H**). The increased amount of infection in iMac supports the supposition that iMacs are more permissive to HIV infection than iMg. Further, the differentially regulated genes indicate that iMacs and iMg not only differ in the extent of HIV infection but also exhibit distinct transcriptional responses to infection, suggesting that their intrinsic cellular programs may shape both HIV permissiveness and the inflammatory response to infection.

**Figure 7:**
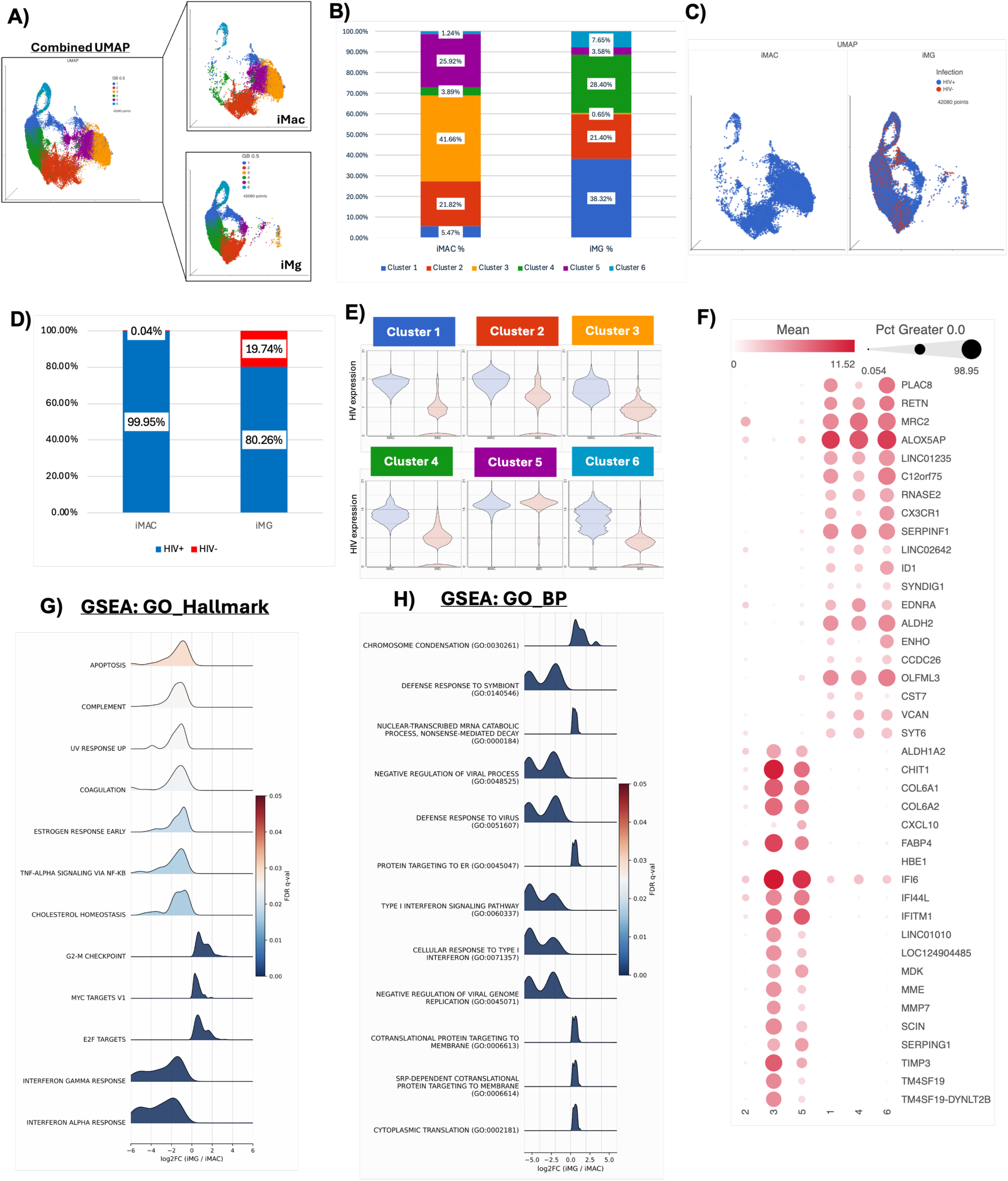
iMg and iMacs exhibit distinct transcriptional profiles in response to HIV infection. A) Combined and individual UMAP plots of HIV-infected iMg and iMacs. (B) Changes in the proportional representation of iMg and iMacs across identified clusters. (C) UMAP plots showing HIV-positive (blue) and HIV-negative (red) cells in iMg and iMacs. (D) Quantification of HIV-positive and HIV-negative cells in iMg and iMacs. (E) Cluster-level differences in HIV transcript expression between iMg and iMacs. (F) Top differentially expressed genes (DEGs) between iMg and iMacs within each cluster. Gene Ontology (GO) pathway analysis showing differences between iMg and iMacs for (G) Hallmark pathways and (H) biological process pathways.

## Discussion

A major limitation in studying human CNS myeloid cells is the limited access to these populations in human tissue. This challenge is particularly daunting when attempting to isolate more than one cell type from the same donor under controlled experimental conditions, making it difficult to evaluate discrete cell-type specific mechanisms that contribute to persistent inflammation, as well as to viral reservoir maintenance during HIV infection of the CNS. To address this limitation, we assessed differences in transcriptomic profiles, inflammatory response, susceptibility to HIV infection, and ART responsiveness in syngeneic microglia-like cells (iMg) and macrophage-like cells (iMac) from three independent iPSC donor lines. Both populations exhibited many of the morphological and molecular characteristics consistent with their primary counterparts. Morphologically, microglia typically display smaller cell bodies with highly ramified processes to support continuous surveillance of the brain parenchyma^58^, and similar morphology was observed in the iMg. In contrast, macrophages generally exhibit larger, amoeboid morphologies, as seen in the iMacs, associated with migratory and phagocytic activity^59^. Interestingly, the iMac and iMg also showed differential expression of the proliferation marker Ki67, with higher levels in iMacs compared to iMg. Macrophages have long been considered terminally differentiated, but our recent data show the presence of Ki-67 in primary monocyte derived macrophages^33^, and growing evidence indicates that macrophage populations may retain proliferative capacity^33,60–62^. Microglia are generally more quiescent under homeostatic conditions and proliferate primarily in response to inflammatory stimuli, potentially explaining the lower but detectable baseline Ki67 expression observed in iMg. This baseline expression is consistent with human studies estimating microglial renewal at ∼28% per year and ∼2% Ki67-positive cells in brain tissue at a given time^63,64^. This difference in proliferative status is an important experimental consideration, particularly in studies involving prolonged culture and infection paradigms.

Alongside their expected morphology, the iMg and iMac expressed microglial (P2RY12, TMEM119) and macrophage (CD68, CCR5, no P2RY12) identity markers respectively. The robust expression of TMEM119 is particularly important, as it is widely regarded as a microglia-specific marker that distinguishes resident parenchymal microglia from infiltrating macrophages and monocyte-derived cells^65,66^. Its presence suggests that the iMg recapitulate aspects of microglial identity and maturation. Notably, iMg did not express SALL1 a transcription factor critical for establishing and maintaining microglial identity *in vivo.* Although SALL1 is often considered a key transcriptional regulator for microglial identity, a loss of SALL1 expression in the iMg is not unusual. Other studies in iPSC microglia also find lower SALL1 expression^67^, and it is a recognized limitation of many iPSC-derived microglia-like models and likely reflects incomplete recapitulation of the CNS environment^68^. This is supported by the fact that *in vitro* culture of primary microglia show the same loss of SALL1^69^, and this should be considered when interpreting findings from *in vitro* systems. Despite this caveat, the capacity to generate these syngeneic models represents a key strength in the use of iPSCs. Many pathways relevant to inflammation and HIV infection, including innate sensing, interferon signaling, and inflammatory cytokine production, are highly donor dependent^70–73^. Deriving microglia and macrophages from the same donor minimizes confounds resulting from genetic variability and enables more precise interrogation of cell-type specific responses to infection or inflammatory stimuli. While this has been examined in murine models^74–76^, species differences in immune signaling, viral restriction factors, and HIV replication dynamics may limit the translational relevance of those studies.

Bulk RNA sequencing of both cell types demonstrated the expected expression of genes converging on immune and inflammatory signaling pathways, but there were still substantial differences between the discrete cell types in these processes. Hallmark pathway analysis highlighted differences in TNF-α signaling via NF-κB, IL6–JAK–STAT3 signaling, and both interferon-γ and interferon-α responses. Gene Ontology analyses further identified enrichment in antigen presentation, chemokine signaling, inflammatory responses, and regulation of NF-κB activity. The differential enrichment of TNF-α signaling, NF-κB regulatory pathways, interferon responses and antigen presentation indicates that these cell types may differ in how they activate or regulate of innate immune activation and cytokine production ^46,77^ or detect viral infection and coordinate immune signaling^78,79^. These differences may reflect their specialized roles *in vivo*; microglia operate within a tightly regulated CNS environment and must balance pathogen sensing with preservation of neuronal function, whereas macrophages are often adapted for more rapid and robust inflammatory responses in perivascular and peripheral tissues^80^.

Notably, these transcriptional differences are present at baseline, suggesting distinct pre-activation states that may shape downstream responses to infection or inflammatory stimuli, consistent with the functional differences observed in NF-κB activation. While we found no differences in baseline NF-κB activity, there were significant differences in the responses to stimulation of TLR2 and 4. Responses to TLR3 and TLR7 agonists were limited to very high concentrations of agonist in both cell types, consistent with prior observations in other macrophage models ^81–84^. These differences do not seem to be due to variation in TLR expression, and suggest that virus sensing TLR such as TLR3 and 7 may trigger distinct transcriptional programs compared to surface expressed bacterial/fungal sensors like TLR2 and TLR4, with myeloid cells deprioritizing interferon-mediated antiviral defenses relative to classical NF-κB-driven inflammatory responses.

Indeed, in response to TLR4 activation with LPS, iMg displayed a steep response consistent with switch-like activation of NF-κB, while iMacs exhibited a greater maximal NF-κB response but required higher LPS concentrations to reach comparable activation. This suggests that iMg have higher sensitivity to low levels of endotoxin like LPS, while the iMac response was relatively blunted, consistent environments with greater pathogen exposure. Considered alongside the baseline and LPS driven cytokine expression, these data suggest that iMacs exist in a more chemotactic inflammatory state, while iMg exhibit a more restrained baseline profile but retain the capacity for strong, stimulus-dependent cytokine induction following TLR4 engagement.

Similar distinctions in microglia and macrophage inflammatory responses have been reported *in vivo* in murine models, with macrophages showing rapid but more graded NF-κB activation dynamics over time while microglia are a bit more tightly regulated with some studies also observing a rapid response^81,85,86^. These differences may be attributed, at least in part, to differences in chromatin landscapes, with each cell type exhibiting distinct patterns of chromatin accessibility at inflammatory regulatory regions, including loci enriched for NF-κB and AP-1 motifs^87–89^. For example, a more restricted but plastic chromatin configuration at inflammatory loci in microglia^90,91^ may permit rapid activation while limiting excessive transcriptional amplification of pro-inflammatory genes, whereas a broader chromatin landscape in macrophages^87,88,92^ may support a wider dynamic range of inflammatory gene expression following TLR4 stimulation. This dichotomy supports functionally distinct roles for both myeloid cell types within the CNS, with perivascular/CAM-like macrophages poised for constitutive immune surveillance and leukocyte recruitment, and microglia acting as tightly regulated sensors that require pathogen signals to mount robust inflammatory programs.

While TLR2 stimulation also provoked differential responses, with iMacs mounting a stronger NF-κB response than iMg, although responses became more similar with prolonged stimulation. This difference may be due to the typical location of TLR2 responses, as TLR2 recognizes bacterial lipopeptides associated with Gram-positive organisms such as *Streptococcus pneumoniae* and *Staphylococcus aureus*, common causes of meningitis that access the CNS at meningeal or perivascular sites ^53,54^, while TLR4 is classically activated by Gram-negative bacterial LPS^47^. Although both microglia and CAMs are yolk sac-derived CNS macrophages, CAMs are positioned at CNS interfaces, while microglia are found in the parenchyma. Thus, CAMs are more likely to encounter circulating microbial products or systemic inflammatory mediators, and their heightened TLR2 sensitivity is consistent with the need to rapidly detecting Gram-positive or fungal ligands during bloodstream invasion or meningeal infection. Overall, these stimulus-specific differences suggest that microglia and macrophages are not hierarchically more or less inflammatory but are differentially tuned according to the pattern recognition pathways most relevant to their anatomical localization.

Similarly, both iMg and iMacs were permissive to infection with an M-tropic HIV^93–96^, but the degree of infection differed substantially between the two cell types. Both cell types showed a gradual, dose-dependent increase in infection, using p24 secretion as a surrogate for viral replication^97^. By 9 dpi, quantification of the number of infected (p24+) cells in each culture showed around 4-fold more iMac (12.15%) were p24+ than iMg (2.94%). Treatment with ARV significantly reduced the proportion of infected cells in both models, but the reduction in the number of p24+ cells was much greater in iMacs. This pattern was mirrored in the kinetics of p24 secretion between 5 and 9 dpi, with iMacs showing significant ARV-mediated reductions in p24 secretion during this period, while iMg exhibited significant reductions in secreted p24 only at 9 dpi. This is also shown by single cell RNA sequencing, where a greater proportion of iMacs are infected with HIV relative to the iMg. These findings suggest that CAMs are more susceptible to infection than microglia, correlating with several reports suggest that perivascular macrophages harbor a greater burden of productive infection during untreated disease^98–101^. These data also suggest that microglia may be comparatively less sensitive to viral suppression by ARVs. Single-cell RNA sequencing analyses of postmortem brain tissue from virally suppressed individuals^21,102^ or macaques^102,103^ on ART have detected low but persistent HIV transcripts and integrated viral DNA within brain macrophages, but the specific types of macrophages are not clear. However, a recent study from the Last Gift cohort found that approximately 0.5% of microglia contained detectable viral RNA and DNA, whereas infected perivascular macrophages were largely undetectable^21^.

Differences in viral infection and ART penetrance between microglia and CAMs may also reflect their distinct inflammatory signaling programs^104,105^. Stronger NF-κB signaling in macrophages may promote productive infection, whereas tighter regulation in microglia may limit overt inflammation while allowing low-level viral persistence. This distinction could have important implications for therapeutic strategies aimed at reservoir clearance, as many latency-reversing approaches rely on NF-κB activation to induce viral expression from latent proviruses^106,107^. If microglia regulate this pathway differently, the stimuli required to reactivate latent virus may differ from those effective in macrophages or peripheral CD4⁺ T cells, which most latency-reversal strategies target. Further, inflammatory signals present in the CNS could differentially modulate NF-κB activity in CAMs vs. microglia to influence reservoir stability or contribute to transient viral reactivation^108–110^, particularly in comorbid conditions that heighten neuroinflammation such as substance use or aging^111–113^. These data support the hypothesis that microglia, rather than CAMs, disproportionately contribute to residual viral activity in the CNS, but further studies are needed to dissect the relative contributions of transcriptionally active versus latent infection between CAMs versus microglia.

Overall, these data indicate substantial, cell-type specific differences in the inflammatory response and susceptibility to HIV infection and ART between macrophages and microglia. The distinct infection dynamics and inflammatory signaling profiles observed between macrophages and microglia suggest that these populations may differentially shape the neuroinflammatory response to brain insult, and more specifically HIV persistence, viral reactivation, and the progress of HIV associated brain injury (HABI). Understanding these cell-type specific mechanisms will therefore be important for designing therapeutic approaches capable of effectively targeting viral reservoirs within the CNS and may also provide broader insight into how myeloid-driven inflammation contribute to other neuroinflammatory and neurodegenerative diseases.

## Declarations

The authors declare no competing interests.

## Acknowledgements

We would like to thank all members of the Gaskill, Matt, and Fox Labs for their contributions, as well as for valuable feedback and discussions. We would also like to thank the Human Pluripotent Stem Cell Core, specifically Drs. Deborah French and Jean-Anne Maguire, for providing the common myeloid progenitors and assisting with the development of our iMicroglia and iMacrophage models.

