## Supplementary material for "Syngeneic human iPSC Macrophages and Microglia possess distinct neuroinflammatory profiles, responses to HIV infection, and ART treatment": Table S1-2-3

| Suplementary Table 1. High Content Imaging NF-kB Colocalization Analysis Parameters | |
|---|---|
| Condition | Value |
| **Ch1** | **DAPI** |
| Ch1: SmoothFactor | 1 |
| Ch1: Thresholding (Fixed) | 809 |
| Ch1: Segmentation (Intensity) | –999 |
| BackgroundCorrectionCh1 | 50 |
| ObjectAreaCh1 | 31.16 - 394.45 |
| Object.Average Intensity.Ch1 | 0 - 65535 |
| Object.Variable Intensity.Ch1 | 0 - 65535 |
| RejectBorderObjects | Yes |
| **Ch2** | **NF-kB** |
| Ch2: SmoothFactor | 4 |
| Ch2: Thresholding (Fixed) | 97 |
| Ch2: Segmentation (Intensity) | –2517 |
| BackgroundCorrectionCh2 | –255 |
| ObjectAreaCh2 | 53.52 - 5851.17 |
| Object.Average Intensity.Ch2 | 93.19 - 7748.41 |
| Object.Variable Intensity.Ch2 | 0 - 65535 |
| RejectBorderObjects | Yes |
| **Ch3** | **CMDR** |
| Ch3: SmoothFactor | 2 |
| Ch3: Thresholding (Fixed) | –3000 |
| Ch3: Segmentation (Intensity) | 10 |
| BackgroundCorrectionCh3 | –255 |
| ObjectAreaCh3 | 136.63 - 8935.92 |
| Object.Average Intensity.Ch3 | 0 - 65535 |
| Object.Variable Intensity.Ch3 | 0 - 65535 |
| RejectBorderObjects | Yes |
| **Colocalization** | |
| ROI A Channel | DAPI |
| ROI A Target | NF-kB |
| ROI B Channel | CMDR |
| ROI B Target | NF-kB |
| *CMDR: Cell Mask Deep Red, ROI: Region of Interest | |

| Suplementary Table 2. High Content Imaging Morphology Analysis Parameters | |
|---|---|
| Condition | Value |
| **Ch3** | **CMDR** |
| Ch3: SmoothFactor | 2 |
| Ch3: Thresholding (Fixed) | –3000 |
| Ch3: Segmentation (Intensity) | 10 |
| BackgroundCorrectionCh3 | –255 |
| ObjectAreaCh3 | 136.63 - 8935.92 |
| Object.Average Intensity.Ch3 | 0 - 65535 |
| Object.Variable Intensity.Ch3 | 0 - 65535 |
| RejectBorderObjects | Yes |
| **Ch1** | **DAPI** |
| Ch1: SmoothFactor | 1 |
| Ch1: Thresholding (Fixed) | 809 |
| Ch1: Segmentation (Intensity) | –999 |
| BackgroundCorrectionCh1 | 50 |
| ObjectAreaCh1 | 31.16 - 394.45 |
| Object.Average Intensity.Ch1 | 0 - 65535 |
| Object.Variable Intensity.Ch1 | 0 - 65535 |
| RejectBorderObjects | Yes |

| Suplementary Table 3 High Content Imaging p24 Colocalization Analysis Parameters | |
|---|---|
| Condition | Value |
| **Ch1** | **CMDR** |
| Ch1: SmoothFactor | 1 – 2 |
| Ch1: Thresholding (Isodata) | –0.62 – 0.857 |
| Ch1: Segmentation (Intensity) | –545 – - 1368 |
| BackgroundCorrectionCh1 | –255 – 0 |
| ObjectAreaCh1 | 0 - 372637 |
| Object.Average Intensity.Ch1 | 0 - 65535 |
| Object.Variable Intensity.Ch1 | 164.19 - 54033.18 |
| RejectBorderObjects | Yes |
| **Ch2** | **p24** |
| Ch2: SmoothFactor | 8 – 43 |
| Ch2: Thresholding (Isodata) | –0.99 - 0.145 |
| Ch2: Segmentation (Intensity) | 0 |
| BackgroundCorrectionCh2 | –255 – 0 |
| ObjectAreaCh2 | 0 - Max |
| Object.Average Intensity.Ch2 | 0 - 65535 |
| Object.Variable Intensity.Ch2 | 195.38 - 8819.57 |
| RejectBorderObjects | Yes |
| **Colocalization** | |
| ROI Channel | CMDR |
| ROI Target | P24 |
| *CMDR: Cell Mask Deep Red, ROI: Region of Interest | |
