## Supplementary material for "Syngeneic human iPSC Macrophages and Microglia possess distinct neuroinflammatory profiles, responses to HIV infection, and ART treatment": Table S4

**Supplementary Table 1: Summary Statstics for scRNAseq**

| | HIV | HIV+Cocaine |
|---|---|---|
| Estimated Number of Cells | 9,474 | 15,704 |
| Mean Reads per Cell | 43,512 | 31,749 |
| Median Genes per Cell | 4,293 | 3,430 |
| Number of Reads | 412,236,248 | 498,582,578 |
| Valid Barcodes | 95.30% | 95.30% |
| Sequencing Saturation | 22.70% | 22.50% |
| Q30 Bases in Barcode | 92.60% | 92.70% |
| Q30 Bases in RNA Read | 90.90% | 92.00% |
| Q30 Bases in UMI | 92.50% | 92.70% |
| Reads Mapped to Genome | 96.20% | 96.00% |
| Reads Mapped Confidently to Genome | 82.80% | 82.20% |
| Reads Mapped Confidently to Intergenic Regions | 1.70% | 1.80% |
| Reads Mapped Confidently to Intronic Regions | 20.90% | 21.30% |
| Reads Mapped Confidently to Exonic Regions | 60.20% | 59.10% |
| Reads Mapped Confidently to Transcriptome | 75.60% | 74.60% |
| Reads Mapped Antisense to Gene | 4.60% | 4.80% |
| Fraction Reads in Cells | 77.80% | 79.80% |
| Total Genes Detected | 31,040 | 31,571 |
| Median UMI Counts per Cell | 17,058 | 11,358 |
