## Supplementary material for "Syngeneic human iPSC Macrophages and Microglia possess distinct neuroinflammatory profiles, responses to HIV infection, and ART treatment": Table S5

| | ID | Description | setSize | enrichmentScore | NES | pvalue |
|---|---|---|---|---|---|---|
| **HALLMARK_INFLAMMATORY_RESPONSE** | HALLMARK_INFLAMMATORY_RESPONSE | HALLMARK_INFLAMMATORY_RESPONSE | 159 | 0.53080702 | 1.94922614 | 1.55E-16 |
| **HALLMARK_TNFA_SIGNALING_VIA_NFKB** | HALLMARK_TNFA_SIGNALING_VIA_NFKB | HALLMARK_TNFA_SIGNALING_VIA_NFKB | 190 | 0.50596928 | 1.87213174 | 3.28E-16 |
| **HALLMARK_KRAS_SIGNALING_UP** | HALLMARK_KRAS_SIGNALING_UP | HALLMARK_KRAS_SIGNALING_UP | 149 | 0.51834841 | 1.8949165 | 4.76E-14 |
| **HALLMARK_EPITHELIAL_MESENCHYMAL_TRANSITION** | HALLMARK_EPITHELIAL_MESENCHYMAL_TRANSITION | HALLMARK_EPITHELIAL_MESENCHYMAL_TRANSITION | 135 | 0.46346344 | 1.68414346 | 1.16E-08 |
| **HALLMARK_IL2_STAT5_SIGNALING** | HALLMARK_IL2_STAT5_SIGNALING | HALLMARK_IL2_STAT5_SIGNALING | 174 | 0.44262715 | 1.63004231 | 1.93E-08 |

| | | | | | | |
|---|---|---|---|---|---|---|
| **HALLMARK_ALLOGRAFT_REJECTION** | HALLMARK_ALLOGRAFT_REJECTION | HALLMARK_ALLOGRAFT_REJECTION | 146 | 0.44742281 | 1.6329284 | 9.60E-08 |
| **HALLMARK_TGF_BETA_SIGNALING** | HALLMARK_TGF_BETA_SIGNALING | HALLMARK_TGF_BETA_SIGNALING | 51 | 0.54734148 | 1.84629996 | 3.35E-06 |
| **HALLMARK_KRAS_SIGNALING_DN** | HALLMARK_KRAS_SIGNALING_DN | HALLMARK_KRAS_SIGNALING_DN | 75 | 0.48356092 | 1.68371911 | 7.16E-06 |
| **HALLMARK_ANGIOGENESIS** | HALLMARK_ANGIOGENESIS | HALLMARK_ANGIOGENESIS | 23 | 0.63891928 | 1.96278618 | 4.18E-05 |
| **HALLMARK_HYPOXIA** | HALLMARK_HYPOXIA | HALLMARK_HYPOXIA | 160 | 0.39569098 | 1.45368428 | 4.80E-05 |
| **HALLMARK_IL6_JAK_STAT3_SIGNALING** | HALLMARK_IL6_JAK_STAT3_SIGNALING | HALLMARK_IL6_JAK_STAT3_SIGNALING | 71 | 0.4584321 | 1.58813609 | 0.00015077 |
| **HALLMARK_UV_RESPONSE_DN** | HALLMARK_UV_RESPONSE_DN | HALLMARK_UV_RESPONSE_DN | 119 | 0.3969323 | 1.42713248 | 0.00048084 |
| **HALLMARK_INTERFERON_GAMMA_RESPONSE** | HALLMARK_INTERFERON_GAMMA_RESPONSE | HALLMARK_INTERFERON_GAMMA_RESPONSE | 183 | 0.36144026 | 1.33348208 | 0.00118733 |

| | | | | | | |
|---|---|---|---|---|---|---|
| **HALLMARK_APOPTOSIS** | HALLMARK_APOPTOSIS | HALLMARK_APOPTOSIS | 142 | 0.37232308 | 1.35629994 | 0.00279337 |
| **HALLMARK_ESTROGEN_RESPONSE_LATE** | HALLMARK_ESTROGEN_RESPONSE_LATE | HALLMARK_ESTROGEN_RESPONSE_LATE | 141 | 0.36188402 | 1.31815223 | 0.00312634 |

**Table 3: GO_Hallmark**

| p.adjust | qvalue | rank | leading_edge |
|---|---|---|---|
| 7.75E-15 | 4.73E-15 | 2506 | tags=48%, list=19%, signal=40% |
| 8.21E-15 | 5.01E-15 | 4126 | tags=57%, list=31%, signal=40% |
| 7.93E-13 | 4.84E-13 | 2969 | tags=46%, list=22%, signal=36% |
| 1.44E-07 | 8.82E-08 | 1422 | tags=28%, list=11%, signal=25% |
| 1.93E-07 | 1.18E-07 | 4023 | tags=47%, list=30%, signal=33% |

| | | | |
|---|---|---|---|
| 8.00E-07 | 4.88E-07 | 3217 | tags=45%, list=24%, signal=35% |
| 2.40E-05 | 1.46E-05 | 3636 | tags=63%, list=27%, signal=46% |
| 4.47E-05 | 2.73E-05 | 2131 | tags=33%, list=16%, signal=28% |
| 0.0002323 | 0.00014183 | 2903 | tags=61%, list=22%, signal=48% |
| 0.00023994 | 0.00014649 | 3441 | tags=40%, list=26%, signal=30% |
| 0.00068534 | 0.00041842 | 1916 | tags=39%, list=14%, signal=34% |
| 0.00200352 | 0.0012232 | 3945 | tags=40%, list=30%, signal=29% |
| 0.00456666 | 0.00278807 | 3703 | tags=37%, list=28%, signal=27% |

| | | | |
|---|---|---|---|
| 0.0099763 | 0.0060908 | 3622 | tags=40%, list=27%, signal=30% |
| 0.01042115 | 0.00636239 | 2409 | tags=27%, list=18%, signal=22% |

| core_enrichment |
| --- |
| 2015/3552/6372/10630/4210/6364/1441/3575/3553/1839/3576/7130/5142/4084/5008/2357/1240/10203/10507/133/23601/7097/64127/558/29933/5724/10135/6347/3586/929/114548/23533/7162/23308/2867/5329/9435/7098/7133/3587/4323/23166/1880/2769/3091/970/3554/490/7076/3606/1902/3386/7832/4067/2014/684/6004/719/728/6324/19/196/4792/7096/4973/3696/8743/1318/255488/7439/8578/3269/7855/5732/2643/391/5663 |
| 3552/6372/56937/57007/2919/6364/2921/5743/3575/3553/1839/7130/5142/4084/5055/9021/5328/7422/7124/2920/2114/3164/4929/7097/3914/7538/10769/9120/127544/1052/10135/6303/8942/50486/6347/8870/23308/7127/5329/10957/1846/51561/6351/388/2355/960/687/602/1960/1880/7262/374/4814/23135/490/3606/5271/7832/3398/10221/5187/4794/19/4170/1051/8061/25976/4792/10938/1326/9314/4973/2354/1647/7050/23258/2353/11080/2643/3433/10318/55332/694/51278/7128/941/1843/80149/4616/6648/10611/4783/8877/25816/8013/2683/5966/5970/9242/3726/7280/3460/10209/9945/6515/1958/4780/329/182 |
| 6590/56729/101/6364/6304/25960/5743/1438/3575/3553/1839/7852/5950/2591/2517/5328/51311/1240/1636/3075/8808/8829/50856/50486/56172/639/26499/23136/3394/5329/2119/5175/2556/25907/3486/2022/7133/3587/2207/55365/2745/57332/1794/23531/28959/1520/3398/10221/6004/719/6324/867/11179/9314/2118/4214/10912/10320/7805/9935/51129/2048/22903/5936/7128/55339/2115/23268 |
| 1462/6695/5999/4907/1004/1289/7474/6372/56937/7045/2919/2201/3576/5654/2517/2619/5352/7422/6678/5376/7057/3487/6303/6443/30008/5768/5329/10409/3486/4323/388/960/23705/374/7076/2014/3398/3685 |
| 8600/8530/1493/4907/399/3662/943/80760/1028/8728/4084/10875/942/27242/8829/3586/4050/3394/6095/84230/3563/8835/2027/7133/3587/5359/388/960/5339/8477/23075/51284/9144/81848/659/10924/3685/6004/196/5621/5325/1326/254013/28996/990/8784/8743/7850/2118/126669/64116/1154/23258/5451/200734/3459/5732/7494/56990/23710/79971/4953/9261/822/4616/7372/1690/4783/64332/23191/1522/11230/3566/4839/51651/9249/3664/8809/2184/975/4641/6515 |

3111/3122/729230/3117/2213/2833/3662/972/6775/1956/3553/3108/5552/7124/3109/2650/942/7903/7097/940/1234/6347/912/3586/114548/4050/3394/23308/9450/920/3603/7098/3059/6351/1230/23166/602/8477/3091/3133/7076/3606/3455/4067/3135/1520/7454/7096/9655/1445/27240/6688/3105/10333/567/2000/3459/5777/941/8661/3551/5788/6885/822/4689/6223

3397/56937/4091/1028/3399/4092/6498/7057/6497/7082/64750/7046/11031/2022/9612/90/659/3398/57154/28996/1499/3065/7050/2339/1025/51742/6885/5494/3726/387/7040/3460

50489/4210/5055/6498/54847/4929/3780/57118/3779/1952/7771/3425/9435/22849/51226/23031/5133/9194/7832/79917/5795/55890/2827/57089/56848

1462/3714/6372/7056/7422/27242/8829/7448/7076/3685/4973/7410/4043/6275

51316/1289/57007/7045/8406/1956/1028/3162/54541/9957/7852/7422/133/54206/7538/8870/7162/6095/5329/10957/9435/3486/2027/2355/2745/26136/26118/2908/8459/6478/23210/5578/25976/6513/3669/23327/2548/8509/1027/55818/230/7436/4214/2353/3309/3073/3423/51129/694/23036/10570/8819/7167/7128/1843/901/6275/1944/4783/7360/3340/7428/3098/55276

2919/2921/1441/1438/3553/3162/9021/7124/27242/7097/929/4050/23533/3676/3563/7133/1230/960/3554/952/1439/867/1326/7132/7850/2885/3597/23765

3397/5999/5577/4092/2650/26037/5376/8829/7082/6256/8821/6310/54468/2535/490/1902/2908/57178/8621/5578/8503/27109/5922/8455/9975/5728/1027/7436/1628/5002/11044/7410/7048/1843/51460/2064/10401/4255/10611/27032/11099/91746/1739/1859/3638/22884/6383/136

3123/3117/3662/2841/972/6775/5743/6398/3695/3108/7130/5142/9021/26053/2357/1240/10068/10875/942/7903/3075/4261/10135/834/6347/10437/3394/7127/55024/2209/10628/256586/3587/5359/3091/55016/952/3455/57162/1439/3135/684/84166/51735/4792/3669/3663/8743/3105/567/3106/2643/3433/841/694/5777/7128/10616/8737/6648/11054/7726/57169/8673/25939/10906/103/3566

3552/6304/3553/3162/4092/7124/834/6303/56172/
8870/929/10628/388/960/1960/2769/7076/3606/95
2/90427/7832/2878/5611/5058/4170/2876/4000/36
69/1499/3148/1027/8743/637/1647/2944/1806/300
5/3459/2643/841/3875/5663/1017/8567/5914/5117
6/2064/3301/4616/4255/6648/1387/5534/5970/210
7/150209/5538

7134/6280/5577/55806/5265/7022/7538/3487/4681
/1152/8645/1952/8140/5364/7162/10105/11031/96
12/6337/9060/960/1960/374/4665/3398/6478/2288/
9314/990/3669/3613/1154/7358/3716/2353/29107/
1345/7494
