## Supplementary material for "Syngeneic human iPSC Macrophages and Microglia possess distinct neuroinflammatory profiles, responses to HIV infection, and ART treatment": Table S6

| | baseMean | log2FoldChan | lfcSE | stat | pvalue | padj |
|---|---|---|---|---|---|---|
| ALOX5 | 1824.82669 | 5.08262183 | 0.27122476 | 18.7395198 | 2.357452644 | 3.310570749 |
| CSF3R | 953.497105 | 4.29216962 | 0.27885288 | 15.3922371 | 1.845401589 | 1.295748725 |
| PADI2 | 2767.7453 | 8.15368402 | 0.53524805 | 15.2334681 | 2.120390293 | 9.925546964 |
| DDIT4L | 1991.1102 | -4.2916593 | 0.3145304 | -13.644657 | 2.172335340 | 7.626526297 |
| MARCKSL1 | 1024.83012 | 4.08384381 | 0.30355397 | 13.4534355 | 2.938940444 | 8.254308133 |
| GAL3ST4 | 490.648689 | 5.80426519 | 0.44911694 | 12.9237279 | 3.307210283 | 7.740525669 |
| SEMA4C | 993.34397 | 4.94964679 | 0.38339566 | 12.910023 | 3.951827825 | 7.927931165 |
| ADAM8 | 291.95218 | 5.26532204 | 0.41740623 | 12.614383 | 1.759300660 | 3.088232396 |
| PPARG | 673.733613 | -5.0320434 | 0.40908163 | -12.30083 | 8.964861403 | 1.398817207 |
| FCGR3A | 3535.14522 | 6.68554208 | 0.5441312 | 12.2866361 | 1.068609575 | 1.500648426 |
| ALDH1A2 | 885.803279 | -7.2269876 | 0.59490908 | -12.148054 | 5.874721898 | 7.499883601 |
| GPR82 | 299.627981 | 4.79432915 | 0.39685552 | 12.0807924 | 1.334276700 | 1.561437308 |
| LINC01235 | 472.563388 | 5.0034119 | 0.41863253 | 11.9517992 | 6.353470216 | 6.863214019 |
| ALDH2 | 1791.53841 | 6.83342829 | 0.57462968 | 11.8918819 | 1.304341221 | 1.308347412 |
| LYZ | 4910.78022 | 4.67303847 | 0.39377918 | 11.8671546 | 1.753298254 | 1.641437825 |
| CXCL1 | 337.060748 | 4.6178653 | 0.39259637 | 11.762374 | 6.099480853 | 5.353438101 |
| MXD1 | 2056.1877 | 3.32309194 | 0.29024069 | 11.4494352 | 2.366826986 | 1.955138316 |
| FABP4 | 1081.3608 | -8.0436173 | 0.71524712 | -11.245928 | 2.425336798 | 1.892166925 |
| SEMA4A | 650.523742 | 4.83441286 | 0.43298862 | 11.1652192 | 6.034220208 | 4.459923915 |
| SLC8A1 | 1086.80724 | 2.54487273 | 0.23306429 | 10.9191877 | 9.332673178 | 6.552936472 |
| ENSG000002 | 356.417144 | 6.5128224 | 0.5968511 | 10.9119719 | 1.010404974 | 6.756722410 |
| EMP1 | 3606.61276 | -3.1694439 | 0.29290433 | -10.820748 | 2.745283315 | 1.752364254 |
| MRC2 | 4319.97464 | 7.49041793 | 0.697415 | 10.7402592 | 6.585920287 | 4.021133852 |
| SMAD6 | 200.221133 | 4.34994341 | 0.41067662 | 10.5921379 | 3.241038410 | 1.896412600 |
| PPP1R16B | 269.395491 | 4.29186257 | 0.4074703 | 10.5329457 | 6.089837904 | 3.420783747 |
| EDNRA | 1024.40479 | 4.23361412 | 0.40828272 | 10.36932 | 3.419495023 | 1.846921869 |
| DOCK4 | 2469.94977 | 3.03275765 | 0.29366771 | 10.3271742 | 5.310188984 | 2.761888292 |
| PLXNA1 | 487.991853 | -2.8639794 | 0.27791946 | -10.30507 | 6.684310273 | 3.352420327 |
| ATP13A2 | 205.22891 | -3.9936079 | 0.39080262 | -10.21899 | 1.630310883 | 7.894639908 |
| HMOX1 | 21577.2339 | 3.64246934 | 0.35950782 | 10.1318223 | 3.991406351 | 1.868377313 |
| ANKRD33B | 575.991934 | 3.28093583 | 0.32448815 | 10.1111112 | 4.932162583 | 2.234269650 |
| GPR84 | 1538.06098 | 3.45134971 | 0.34335072 | 10.051966 | 9.005266954 | 3.951905120 |
| HBEGF | 960.547161 | 3.70590121 | 0.37280894 | 9.94048384 | 2.774761025 | 1.180786942 |
| WNT5A | 1112.68737 | 5.26396049 | 0.53149858 | 9.9039972 | 3.999622242 | 1.651961622 |
| HLA-DRA | 7767.16799 | 7.41940386 | 0.75073166 | 9.88289721 | 4.938407036 | 1.981430000 |
| PTGS2 | 147.789227 | 3.99385112 | 0.41105929 | 9.71599774 | 2.577125459 | 1.005293689 |
| PDPN | 438.605679 | 4.72708832 | 0.4882437 | 9.68182145 | 3.602398406 | 1.367256238 |
| IL1B | 4411.03799 | 3.75099204 | 0.39289661 | 9.54702073 | 1.334795925 | 4.932773470 |
| NDRG2 | 2429.33294 | 3.39581404 | 0.35596782 | 9.53966587 | 1.432932448 | 5.159659070 |
| HLA-DPA1 | 3556.61535 | 7.44047797 | 0.78194599 | 9.51533485 | 1.811286172 | 6.358972930 |
| OLFML3 | 4899.92668 | 6.42158449 | 0.67584407 | 9.50157708 | 2.067352577 | 7.080934692 |
| AATK | 154.780489 | 3.69920178 | 0.39113748 | 9.45754871 | 3.152466191 | 1.054049588 |
| CD74 | 24252.7394 | 4.34214579 | 0.46725122 | 9.29295758 | 1.500594350 | 4.900661968 |
| KIAA1671 | 178.917234 | -4.6715719 | 0.50498381 | -9.250934 | 2.225394958 | 7.102550319 |
| SIGLEC5 | 452.570504 | 3.63354419 | 0.39476743 | 9.20426547 | 3.440159583 | 1.073559133 |
| MARVELD1 | 268.683755 | -2.5606483 | 0.27918318 | -9.171929 | 4.646289090 | 1.418431254 |
| UNC5B | 536.396232 | 4.94006114 | 0.53967426 | 9.15378318 | 5.497384386 | 1.642548275 |

| | | | | | | |
|---|---|---|---|---|---|---|
| CYP1B1 | 5459.67543 | 2.77170824 | 0.3040268 | 9.11665759 | 7.747690837 | 2.266683800 |
| GIMAP1 | 149.008815 | 3.69572598 | 0.40771558 | 9.0644709 | 1.252104519 | 3.588429339 |
| NREP | 160.286361 | -3.6527644 | 0.40350824 | -9.0525149 | 1.397128053 | 3.923973852 |
| SLC1A3 | 364.370863 | -2.5184589 | 0.27981137 | -9.0005597 | 2.245698587 | 6.183597109 |
| SMAD7 | 673.900082 | 3.24960501 | 0.36185942 | 8.98029684 | 2.700372413 | 7.292563423 |
| IRF2BP2 | 1428.98137 | 2.53674839 | 0.28272236 | 8.97257777 | 2.896546605 | 7.674755468 |
| C3 | 18335.5903 | 4.3951067 | 0.49044595 | 8.96144968 | 3.204374319 | 8.333153438 |
| TRPV4 | 226.790474 | 3.66149248 | 0.40888418 | 8.95484014 | 3.402282433 | 8.686954947 |
| FGD5 | 1871.98194 | 1.97628466 | 0.22121532 | 8.93376021 | 4.117692788 | 1.032584996 |
| RUNX3 | 3375.4144 | 1.82074559 | 0.20385613 | 8.93152241 | 4.201859297 | 1.035205440 |
| ABCC4 | 1644.05754 | 3.15964853 | 0.35399385 | 8.92571601 | 4.428253075 | 1.072171688 |
| VWA5A | 936.435737 | -2.9174689 | 0.32693575 | -8.9236766 | 4.510595264 | 1.073598123 |
| CST7 | 277.615187 | 7.30323392 | 0.82076341 | 8.89809883 | 5.681142194 | 1.329671330 |
| IL1A | 255.984423 | 5.29616644 | 0.59639956 | 8.880232 | 6.672105515 | 1.536006192 |
| ADGRE5 | 776.454966 | -2.1361846 | 0.24231563 | -8.8157111 | 1.189278238 | 2.693715209 |
| ICAM3 | 723.52358 | 3.00619909 | 0.34382512 | 8.7433958 | 2.262064961 | 5.042250516 |
| HLA-DPB1 | 1678.63807 | 6.30962084 | 0.72194799 | 8.73971665 | 2.336958092 | 5.127797264 |
| CSPG4 | 423.903388 | -8.3619574 | 0.96044781 | -8.7063111 | 3.139244934 | 6.782217941 |
| ENSG000002 | 228.65706 | 4.94521049 | 0.57042666 | 8.66931869 | 4.347136539 | 9.186935152 |
| HS3ST1 | 1054.71318 | 3.36237677 | 0.3878899 | 8.66837937 | 4.383142172 | 9.186935152 |
| S100A8 | 408.802797 | 6.2632599 | 0.72453399 | 8.64453561 | 5.402456627 | 1.115686741 |
| ENSG000002 | 104.650704 | 4.413183 | 0.51137706 | 8.62999794 | 6.135307676 | 1.248668488 |
| GALNT12 | 497.502513 | -5.4811719 | 0.63551993 | -8.624705 | 6.425809634 | 1.289109209 |
| SIRPB2 | 430.026299 | 3.47044948 | 0.40449299 | 8.5797518 | 9.507807167 | 1.880537127 |
| ALDH5A1 | 226.264287 | -5.2071112 | 0.60783723 | -8.5666211 | 1.065658632 | 2.078478358 |
| HLA-DRB1 | 4544.75618 | 7.17058392 | 0.83766267 | 8.56022859 | 1.126439539 | 2.166930199 |
| DNASE2 | 2795.2336 | 2.8081325 | 0.32830337 | 8.55346828 | 1.194441303 | 2.266694489 |
| MYL9 | 549.925704 | -6.1836742 | 0.72760169 | -8.4987079 | 1.917126908 | 3.589628423 |
| LRRK2 | 171.054577 | 3.64041592 | 0.42877278 | 8.49031482 | 2.060780857 | 3.807834945 |
| CYP19A1 | 105.895548 | -4.6832736 | 0.55596541 | -8.423678 | 3.648460341 | 6.644323114 |
| ANTXR1 | 3652.2727 | 4.53276599 | 0.53818393 | 8.42233621 | 3.690502050 | 6.644323114 |
| CSF2RA | 676.161054 | 3.91614213 | 0.46507751 | 8.42040742 | 3.751772985 | 6.669132661 |
| SLA | 3663.2334 | 2.58828723 | 0.30827656 | 8.39599099 | 4.619810849 | 8.109500469 |
| SCIN | 289.432135 | -6.6815908 | 0.7967405 | -8.3861568 | 5.022931392 | 8.708274758 |
| CD101 | 529.181415 | 3.85226986 | 0.45984568 | 8.37731006 | 5.415105496 | 9.273698351 |
| TXNIP | 2611.93367 | 1.5353819 | 0.18396212 | 8.34618487 | 7.050342069 | 1.192866911 |
| RHOH | 114.493678 | 5.25604477 | 0.63179455 | 8.31923095 | 8.853588905 | 1.480130345 |
| TBXA2R | 78.1511988 | -4.0004453 | 0.48200601 | -8.2995757 | 1.044839140 | 1.726197181 |
| PROCR | 351.910551 | 3.67713016 | 0.44409678 | 8.28001992 | 1.231546831 | 2.011001413 |
| SKIL | 1213.93272 | 2.90025378 | 0.35227649 | 8.23289062 | 1.827490489 | 2.949821717 |
| MGAT4A | 601.236548 | 2.28464735 | 0.28274733 | 8.08017305 | 6.467493254 | 1.032079633 |
| ASMTL | 161.720121 | 7.50326635 | 0.92908644 | 8.07596149 | 6.694700692 | 1.056333503 |
| GLDN | 100.112155 | -10.346771 | 1.28262221 | -8.0668893 | 7.211186042 | 1.125185395 |
| USP2 | 449.920783 | -4.1539492 | 0.51528398 | -8.0614755 | 7.537897353 | 1.163238379 |
| COL4A2 | 239.014104 | -5.260971 | 0.65585188 | -8.021584 | 1.043901218 | 1.593424436 |
| TSC22D1 | 1300.98951 | -4.6090068 | 0.57474095 | -8.0192767 | 1.063696163 | 1.606181207 |
| KCNJ5 | 99.4685092 | -6.6550772 | 0.83109342 | -8.0076162 | 1.169531905 | 1.747206016 |
| IGLON5 | 415.210381 | 6.40835497 | 0.80100762 | 8.00036706 | 1.240488552 | 1.833703236 |

| | | | | | | |
|---|---|---|---|---|---|---|
| TM4SF19 | 2224.28605 | -6.8348316 | 0.85472483 | -7.9965287 | 1.279759745 | 1.872048553 |
| NRP1 | 11788.1139 | 2.01968058 | 0.25275088 | 7.99079541 | 1.340708968 | 1.940987221 |
| VCL | 674.521911 | -3.3374447 | 0.41874205 | -7.9701685 | 1.584581823 | 2.270641076 |
| DHRS9 | 153.439226 | -5.4707149 | 0.68757424 | -7.9565443 | 1.769108421 | 2.509453491 |
| CSTA | 248.059241 | 5.19472151 | 0.65301721 | 7.95495347 | 1.791988391 | 2.516489298 |
| FFAR4 | 253.692395 | 3.76066602 | 0.47502857 | 7.91671545 | 2.438669759 | 3.368252796 |
| MT-CYB | 26730.2506 | -2.1223743 | 0.26810124 | -7.9163168 | 2.446498506 | 3.368252796 |
| ADGRE1 | 291.519917 | 6.16096358 | 0.78076115 | 7.89097096 | 2.998444263 | 4.088073086 |
| CRABP2 | 851.566868 | -8.0103045 | 1.01849167 | -7.8648699 | 3.694811594 | 4.989061463 |
| ITGA3 | 870.555485 | -6.5482797 | 0.83556743 | -7.8369255 | 4.617118145 | 6.175065724 |
| GOLGA7B | 332.751787 | -5.8680147 | 0.74948552 | -7.8293903 | 4.902415220 | 6.494775183 |
| RNF19B | 832.432542 | 2.2675316 | 0.28972223 | 7.82657089 | 5.013566479 | 6.579954585 |
| SH2D2A | 122.142852 | 4.86936696 | 0.62567932 | 7.78252824 | 7.108932937 | 9.243587522 |
| ATP6V1C2 | 87.407037 | -5.3261599 | 0.68487807 | -7.7768001 | 7.438200188 | 9.582994976 |
| SH3PXD2A | 924.327479 | 1.93422873 | 0.24892559 | 7.77030884 | 7.829498774 | 9.995422844 |
| ACKR3 | 77.331978 | 4.98131371 | 0.64230145 | 7.75541411 | 8.805565793 | 1.106058438 |
| NEDD9 | 224.902261 | -3.1658929 | 0.40822911 | -7.7551865 | 8.821373292 | 1.106058438 |
| ENG | 13774.4435 | 1.48673194 | 0.19178902 | 7.75191385 | 9.051779766 | 1.124903922 |
| ATF3 | 1208.04468 | -4.9784017 | 0.64262307 | -7.7470012 | 9.408803769 | 1.159016064 |
| SHANK3 | 109.378909 | -4.8661121 | 0.62871678 | -7.7397522 | 9.961074052 | 1.216377068 |
| INPP5D | 1351.9727 | 1.56533092 | 0.20230637 | 7.73742783 | 1.014482370 | 1.228135855 |
| FBP1 | 657.862228 | -6.5539594 | 0.84798496 | -7.7288628 | 1.085115591 | 1.302416944 |
| COL4A1 | 351.663151 | -6.7830515 | 0.87854396 | -7.7207878 | 1.156128665 | 1.375891089 |
| CXCL8 | 2098.4275 | 3.53555712 | 0.46016275 | 7.68327538 | 1.550717501 | 1.829976964 |
| NFAM1 | 1348.40285 | 1.83425905 | 0.23991648 | 7.64540665 | 2.082860153 | 2.437467095 |
| NT5DC2 | 3229.5594 | 2.7581111 | 0.36207725 | 7.61746593 | 2.587041588 | 3.002464877 |
| EMILIN1 | 72.4866875 | -4.5615359 | 0.59915649 | -7.6132629 | 2.672614055 | 3.076354031 |
| GPR132 | 207.798135 | 2.13291687 | 0.28135717 | 7.5808159 | 3.433885760 | 3.920492498 |
| SLC16A10 | 1674.89306 | 2.784995 | 0.36900361 | 7.54733816 | 4.442440647 | 5.031064033 |
| RASAL1 | 345.101778 | 4.64740327 | 0.61690528 | 7.53341469 | 4.943031232 | 5.553199008 |
| PCSK5 | 150.002622 | 4.43806996 | 0.59166374 | 7.50100038 | 6.333260704 | 7.058569847 |
| ARHGAP23 | 224.883306 | -5.3468957 | 0.713653 | -7.4922907 | 6.768182366 | 7.483904328 |
| PDE4B | 276.327395 | 3.34803439 | 0.44757836 | 7.4803313 | 7.413547710 | 8.133472694 |
| CABLES1 | 405.916343 | 2.46774874 | 0.33044915 | 7.46786226 | 8.150809426 | 8.873009052 |
| PLAC8 | 378.527741 | 11.7497926 | 1.58052993 | 7.43408424 | 1.052948563 | 1.137427436 |
| C1orf162 | 674.928803 | 3.32898303 | 0.44878865 | 7.41770766 | 1.191647606 | 1.277428041 |
| RNF128 | 254.954432 | -8.2750249 | 1.1162007 | -7.4135636 | 1.229500072 | 1.308020418 |
| COL6A1 | 7167.15355 | -6.3829558 | 0.86624311 | -7.3685502 | 1.724936081 | 1.821299052 |
| SNAI1 | 111.796809 | 5.49723513 | 0.75010074 | 7.32866249 | 2.324609920 | 2.436156500 |
| SLC5A3 | 1829.60796 | -3.9627943 | 0.54109288 | -7.3236859 | 2.412509666 | 2.509546166 |
| RNF213 | 2052.61624 | -1.5055512 | 0.20599145 | -7.3088042 | 2.695301919 | 2.783097415 |
| OLFM2 | 266.560796 | 4.76009703 | 0.65198639 | 7.30091472 | 2.858178647 | 2.929737426 |
| GPC4 | 4317.11876 | -5.8655689 | 0.80362434 | -7.2988941 | 2.901426528 | 2.952516864 |
| PTPRS | 185.117634 | -6.4445213 | 0.88382514 | -7.2916248 | 3.062390058 | 3.093895222 |
| SIGLEC10 | 798.317281 | 4.01604666 | 0.55183519 | 7.27761976 | 3.397613515 | 3.408049043 |
| CLEC12A | 128.446986 | 4.25892133 | 0.58692151 | 7.25637295 | 3.976078523 | 3.960005014 |
| VCAN | 546.69792 | 10.8409558 | 1.49789769 | 7.23744746 | 4.572077845 | 4.521527407 |
| WIPF3 | 102.30832 | -5.6919203 | 0.78757196 | -7.2271748 | 4.931451875 | 4.842823684 |

| | | | | | | |
|---|---|---|---|---|---|---|
| MRPS6 | 1011.03185 | -3.9538218 | 0.54882821 | -7.2041155 | 5.842183732 | 5.697346260 |
| SYT6 | 221.386238 | 5.77036996 | 0.80600275 | 7.15924348 | 8.112349324 | 7.856670452 |
| GPR160 | 423.606218 | 3.62759189 | 0.50691827 | 7.15616725 | 8.296391748 | 7.979878721 |
| VAT1 | 5486.48514 | -2.8123549 | 0.39340444 | -7.1487626 | 8.756362410 | 8.365006620 |
| GYPC | 950.2681 | -2.9713197 | 0.4170093 | -7.125308 | 1.038482111 | 9.853651553 |
| SATB1 | 1431.77836 | 4.23909933 | 0.59545019 | 7.11915019 | 1.085945232 | 1.023485161 |
| HCLS1 | 4316.41745 | 1.41376743 | 0.19878367 | 7.11209043 | 1.142983283 | 1.070060949 |
| MALT1 | 1253.17769 | 2.56624332 | 0.36176593 | 7.09365675 | 1.306138610 | 1.214708907 |
| NES | 284.004644 | -6.2075349 | 0.88075555 | -7.0479658 | 1.815522936 | 1.677328197 |
| ADORA3 | 2003.88486 | 5.13966924 | 0.73028955 | 7.03785129 | 1.952268772 | 1.791876495 |
| ITGB7 | 3411.47628 | 3.62126714 | 0.51650645 | 7.01107823 | 2.364884522 | 2.156498269 |
| SPOCD1 | 631.658506 | -5.6176609 | 0.80192551 | -7.0052153 | 2.466063036 | 2.234253110 |
| YWHAH | 2635.15931 | 1.84292581 | 0.26369278 | 6.98891267 | 2.770247828 | 2.493755785 |
| ARRDC4 | 392.647861 | 1.91279727 | 0.27389206 | 6.98376306 | 2.873765697 | 2.570464438 |
| CDR2L | 153.465424 | 2.34895955 | 0.33690159 | 6.97224243 | 3.119284044 | 2.772411762 |
| SH3KBP1 | 1484.92764 | 2.01673923 | 0.28962509 | 6.96327535 | 3.324511313 | 2.936233482 |
| RASGRP1 | 86.0450933 | -4.2623764 | 0.61225984 | -6.9617116 | 3.361631332 | 2.950461800 |
| HERPUD1 | 2427.09974 | 2.2823682 | 0.32827609 | 6.95258743 | 3.586464130 | 3.128243216 |
| VNN1 | 50.779633 | -9.3676941 | 1.34792469 | -6.9497163 | 3.660216554 | 3.172865498 |
| CDK18 | 271.021933 | -4.854374 | 0.69973971 | -6.9373997 | 3.993827712 | 3.440817335 |
| RNASE2 | 437.068416 | 5.92524669 | 0.85611658 | 6.92107459 | 4.482303756 | 3.838109246 |
| GEM | 137.819077 | -5.1839763 | 0.74914626 | -6.9198453 | 4.521370041 | 3.848096938 |
| F8A3 | 78.2309138 | -9.9912199 | 1.44403129 | -6.9189775 | 4.549152585 | 3.848418659 |
| HS3ST3B1 | 193.166852 | 2.68287956 | 0.38791674 | 6.9161221 | 4.641743655 | 3.903233901 |
| ATP9A | 472.797506 | -4.4855184 | 0.65056036 | -6.8948534 | 5.392037203 | 4.507165383 |
| GLIPR2 | 538.71185 | -3.7151744 | 0.53899316 | -6.8928043 | 5.470309388 | 4.545535783 |
| TNFSF13 | 1209.56146 | 2.92097272 | 0.42383055 | 6.89184086 | 5.507496914 | 4.549516421 |
| APMAP | 2970.16983 | 1.68621929 | 0.24477315 | 6.88890635 | 5.622293439 | 4.617185191 |
| SLC36A1 | 2830.98547 | 1.72984911 | 0.25211839 | 6.86125729 | 6.825709957 | 5.572874705 |
| GUCY1A2 | 460.44636 | -5.9266338 | 0.86524222 | -6.8496818 | 7.401445871 | 6.008006032 |
| JAG2 | 244.267205 | 5.17704556 | 0.75606562 | 6.8473495 | 7.523076723 | 6.071641748 |
| MAN1C1 | 138.122643 | -2.6378429 | 0.38740297 | -6.8090416 | 9.825104244 | 7.884225080 |
| EPB41L1 | 456.65503 | -5.292288 | 0.7797203 | -6.7874185 | 1.141577412 | 9.108620229 |
| ENSG000002 | 137.099627 | 10.2836072 | 1.51820005 | 6.77355212 | 1.256582804 | 9.969600185 |
| ST3GAL1 | 1230.35862 | 1.39473572 | 0.20606182 | 6.76853059 | 1.300968893 | 1.026376751 |
| RXRA | 3487.86117 | 1.9109737 | 0.28248849 | 6.76478434 | 1.335079161 | 1.047403165 |
| BMP2 | 55.9550908 | -4.4182588 | 0.65667452 | -6.7282323 | 1.717365428 | 1.339831261 |
| KL | 132.783707 | 6.50418744 | 0.96855369 | 6.71536079 | 1.876014569 | 1.455517822 |
| GDF15 | 328.642075 | -5.6830044 | 0.84960426 | -6.6890018 | 2.246980057 | 1.733692659 |
| GIMAP8 | 162.44005 | 2.94813995 | 0.44080719 | 6.68804868 | 2.261659166 | 1.733692659 |
| VSIR | 1634.84755 | 2.19589045 | 0.328362 | 6.68740728 | 2.271590467 | 1.733692659 |
| PTGFRN | 868.188368 | 2.41572811 | 0.36156525 | 6.68130614 | 2.368219123 | 1.797670332 |
| SPOCK1 | 148.354589 | 10.3980754 | 1.55756777 | 6.67584143 | 2.458176481 | 1.855923243 |
| CDKN2B | 183.684608 | -2.3152246 | 0.34887589 | -6.6362413 | 3.217829166 | 2.416469250 |
| STX6 | 607.120195 | 1.67203931 | 0.25232669 | 6.62648619 | 3.437716267 | 2.567864337 |
| GFOD1 | 121.603516 | -3.4218266 | 0.51645423 | -6.6256144 | 3.458068185 | 2.569399551 |
| TLE1 | 93.1195106 | -4.4089652 | 0.66698833 | -6.6102584 | 3.836497269 | 2.825812086 |
| CTLA4 | 118.685417 | 6.33942804 | 0.95914673 | 6.60944549 | 3.857622264 | 2.825812086 |

| | | | | | | |
|---|---|---|---|---|---|---|
| TIE1 | 175.700669 | -8.2982839 | 1.25556199 | -6.6092188 | 3.863532867 | 2.825812086 |
| OTOAP1 | 56.4003178 | -4.1563922 | 0.62921267 | -6.6057033 | 3.956345248 | 2.878702400 |
| CCL20 | 79.798265 | 4.55055473 | 0.68916783 | 6.60297029 | 4.030003221 | 2.913685303 |
| ENSG000002 | 126.046122 | 3.5690786 | 0.54057405 | 6.60238612 | 4.045920631 | 2.913685303 |
| USP53 | 543.829031 | 2.94198139 | 0.44622021 | 6.59311551 | 4.306912257 | 3.085814736 |
| FGFR1 | 364.533605 | -4.3271235 | 0.65750558 | -6.5811206 | 4.669156814 | 3.328374068 |
| LMO7 | 89.3365921 | 4.89036255 | 0.74331091 | 6.57916153 | 4.731085639 | 3.355486648 |
| ARL4C | 2417.32577 | 3.15095429 | 0.47948364 | 6.57155752 | 4.979164580 | 3.513688854 |
| TNFRSF8 | 55.1530028 | 4.92123784 | 0.74930648 | 6.56772362 | 5.109025285 | 3.587302104 |
| COL23A1 | 319.958727 | 3.73896709 | 0.56944304 | 6.56600715 | 5.168233463 | 3.610821021 |
| ITPKB | 706.79241 | -2.9089407 | 0.44339565 | -6.5605982 | 5.359236271 | 3.725730443 |
| GPC6 | 168.532461 | 2.20753111 | 0.33840688 | 6.52330439 | 6.877509099 | 4.757677846 |
| NCF1C | 531.140582 | 3.653634 | 0.56113929 | 6.51109992 | 7.460249597 | 5.135504171 |
| CD300LF | 241.543339 | 4.48365561 | 0.68936457 | 6.50404127 | 7.819043040 | 5.356235191 |
| LRRC39 | 74.716809 | -6.744332 | 1.03900074 | -6.4911715 | 8.517147167 | 5.806130955 |
| KLF13 | 1951.4675 | 1.7559649 | 0.27059089 | 6.48937176 | 8.619503713 | 5.847521287 |
| TRIM47 | 194.808642 | -3.1039306 | 0.47924723 | -6.4766792 | 9.376326888 | 6.330373004 |
| PEAK3 | 106.209194 | 4.78618091 | 0.74045601 | 6.46382882 | 1.020863518 | 6.851377635 |
| OAS3 | 465.916833 | -2.4613817 | 0.38085613 | -6.4627599 | 1.028103751 | 6.851377635 |
| PLEC | 7646.79334 | 1.34267576 | 0.2077621 | 6.46256359 | 1.029438639 | 6.851377635 |
| TPM2 | 130.586459 | -7.8798657 | 1.21984775 | -6.4597125 | 1.049020986 | 6.948774390 |
| NID1 | 389.127618 | -6.5471486 | 1.01582265 | -6.4451689 | 1.154717212 | 7.613001793 |
| SGPL1 | 3819.19737 | 1.69863311 | 0.26372638 | 6.44089186 | 1.187734817 | 7.794093479 |
| SLC40A1 | 692.165156 | 3.63970131 | 0.56774471 | 6.41080621 | 1.447521017 | 9.454668671 |
| TRIM2 | 123.854806 | -5.4908979 | 0.85845444 | -6.3962601 | 1.592286967 | 1.035207679 |
| MYO1G | 693.239531 | 5.78381111 | 0.90478251 | 6.39248778 | 1.632082717 | 1.056190672 |
| C1RL | 194.778093 | 2.62484865 | 0.41095426 | 6.38720385 | 1.689463673 | 1.088309099 |
| ADGRG3 | 128.97685 | 5.90907166 | 0.92525108 | 6.38645208 | 1.697786058 | 1.088676238 |
| DLL1 | 57.6892732 | 3.96856073 | 0.62474167 | 6.35232277 | 2.120877176 | 1.353794463 |
| SMAD3 | 226.920876 | -4.1540962 | 0.6551688 | -6.3404976 | 2.290242251 | 1.455288323 |
| PPIF | 2112.93698 | 1.53696426 | 0.2428137 | 6.32980868 | 2.454653592 | 1.552734251 |
| DIP2C | 76.0709961 | -5.5973754 | 0.88601891 | -6.3174447 | 2.659235988 | 1.674603183 |
| BCL7A | 83.9666276 | -2.5402743 | 0.40241021 | -6.3126488 | 2.742995881 | 1.719637998 |
| FCER1A | 1826.44151 | 11.576873 | 1.83487655 | 6.30934707 | 2.802150950 | 1.748915813 |
| NPIPB5 | 174.607297 | -2.3218331 | 0.3683924 | -6.3026085 | 2.926774707 | 1.818614921 |
| MMP7 | 828.458003 | -6.2687201 | 0.99547399 | -6.2972213 | 3.030284657 | 1.874638213 |
| HEG1 | 76.1495531 | -2.9047245 | 0.46170858 | -6.2912509 | 3.149178988 | 1.931425688 |
| FBN2 | 141.9782 | 4.16469039 | 0.66198341 | 6.29123078 | 3.149586858 | 1.931425688 |
| CYP2S1 | 1056.58024 | 2.29166876 | 0.36474783 | 6.28288521 | 3.323464691 | 2.029191941 |
| PARM1 | 1290.60992 | 3.95023628 | 0.62983511 | 6.27185784 | 3.567652557 | 2.168854756 |
| ID1 | 165.893709 | 9.11490688 | 1.45614882 | 6.25959846 | 3.859699801 | 2.336282944 |
| BCAT1 | 3561.05339 | -3.402166 | 0.54507139 | -6.2416889 | 4.328712594 | 2.608931800 |
| PIPOX | 248.446478 | 3.96587815 | 0.6355763 | 6.23981444 | 4.380902608 | 2.629103219 |
| RGCC | 1507.00333 | -3.3361645 | 0.53516337 | -6.2339178 | 4.549117949 | 2.718436739 |
| PTPN14 | 131.726014 | 4.79681582 | 0.77302508 | 6.20525254 | 5.460913648 | 3.249475015 |
| BCAR3 | 212.900696 | -2.7724154 | 0.44776404 | -6.1916884 | 5.952311812 | 3.526933113 |
| CAPN3 | 108.600561 | -4.9758448 | 0.80385351 | -6.1899896 | 6.016818414 | 3.550175672 |
| HTRA1 | 4305.86959 | 3.45084287 | 0.55927248 | 6.17023539 | 6.818840230 | 4.006567922 |

| | | | | | | |
|---|---|---|---|---|---|---|
| SORL1 | 1385.12992 | 2.61261955 | 0.42409632 | 6.16043911 | 7.254350677 | 4.244701940 |
| CITED2 | 1055.91207 | -2.2580548 | 0.36678577 | -6.1563316 | 7.444925698 | 4.338136580 |
| ZFP36 | 1233.30251 | 2.31808851 | 0.37668911 | 6.15385058 | 7.562393905 | 4.388375934 |
| NR1H3 | 949.870224 | -4.3456491 | 0.70784797 | -6.1392407 | 8.291682548 | 4.791773581 |
| CHI3L1 | 8401.93328 | -5.3882868 | 0.87866323 | -6.1323686 | 8.658022251 | 4.968271974 |
| GZF1 | 423.528461 | 2.01867211 | 0.32919279 | 6.1321881 | 8.667853263 | 4.968271974 |
| MYOZ1 | 132.901247 | -4.0883437 | 0.66786482 | -6.1215138 | 9.269051268 | 5.291271827 |
| ELOVL6 | 57.3401807 | -8.5749482 | 1.40159054 | -6.1180124 | 9.474971471 | 5.386924063 |
| CX3CR1 | 838.565999 | 5.08999271 | 0.83216434 | 6.11657152 | 9.560998567 | 5.413915438 |
| RCBTB2 | 509.487488 | 1.38006726 | 0.22574814 | 6.11330526 | 9.758844171 | 5.503752959 |
| ADAM19 | 496.906445 | 3.59573741 | 0.58962055 | 6.09839225 | 1.071405765 | 6.018300468 |
| MAN1A1 | 935.621924 | -2.0221101 | 0.33185086 | -6.0934305 | 1.105163203 | 6.183189988 |
| FCGR2A | 5740.93771 | 2.93509829 | 0.48222541 | 6.0865691 | 1.153558853 | 6.428344036 |
| S100A9 | 4482.90577 | 3.85653763 | 0.63390559 | 6.08377286 | 1.173868753 | 6.515667553 |
| KCNJ5-AS1 | 412.751854 | -4.1756012 | 0.68721832 | -6.0760912 | 1.231473803 | 6.808498669 |
| PXDC1 | 1321.77243 | 1.75857271 | 0.29032643 | 6.05722573 | 1.384891921 | 7.626681273 |
| COTL1 | 14600.1052 | 1.14558832 | 0.18959034 | 6.04244048 | 1.518003564 | 8.327079710 |
| ISM1 | 68.1627582 | -5.5643531 | 0.92307499 | -6.0280618 | 1.659375718 | 9.067164672 |
| VCAM1 | 45.1732773 | -8.2296183 | 1.36692178 | -6.0205481 | 1.738274136 | 9.461466548 |
| SPTBN1 | 248.57618 | -3.7829276 | 0.62867932 | -6.0172612 | 1.773927741 | 9.618249912 |
| LINC01094 | 680.093303 | 2.37119745 | 0.3948379 | 6.00549607 | 1.907477807 | 1.030258109 |
| HMGB3 | 869.777236 | 2.00378702 | 0.33370588 | 6.00465004 | 1.917450276 | 1.031676407 |
| FZD1 | 320.885356 | 1.64551516 | 0.27419966 | 6.00115681 | 1.959166740 | 1.050098417 |
| CELSR1 | 203.928217 | -3.4484604 | 0.57479137 | -5.9994993 | 1.979269359 | 1.056839528 |
| TPD52L1 | 70.1642016 | -6.3829061 | 1.06448493 | -5.9962391 | 2.019396170 | 1.074181076 |
| CCNG1 | 747.353379 | -1.5491679 | 0.2583857 | -5.9955636 | 2.027808974 | 1.074585714 |
| NAGLU | 713.207189 | -1.6393691 | 0.27353018 | -5.9933755 | 2.055294809 | 1.078030464 |
| AFAP1L1 | 311.46674 | -5.7006296 | 0.95116577 | -5.9933082 | 2.056146319 | 1.078030464 |
| CAMK2B | 59.6854716 | -4.2240912 | 0.70481236 | -5.9932139 | 2.057339347 | 1.078030464 |
| ASPH | 1537.42956 | -1.1771764 | 0.19669043 | -5.9849196 | 2.164968384 | 1.130210075 |
| RRAS2 | 45.0576911 | -4.31252 | 0.72112136 | -5.9802972 | 2.227308956 | 1.158448136 |
| PLOD2 | 129.832778 | 3.04685331 | 0.51127735 | 5.95929653 | 2.533260631 | 1.312715093 |
| ADAMTSL4 | 307.597151 | 2.84406827 | 0.47857484 | 5.94278681 | 2.802167682 | 1.445968778 |
| SLC28A3 | 96.038532 | -6.6295853 | 1.11566629 | -5.9422655 | 2.811095941 | 1.445968778 |
| PMEPA1 | 173.723761 | 5.01163387 | 0.84347204 | 5.94167159 | 2.821302037 | 1.445968778 |
| GPRIN3 | 3207.86257 | 1.62607513 | 0.27407138 | 5.93303522 | 2.973850166 | 1.518610104 |
| MEF2A | 1977.33566 | 1.52564834 | 0.25800538 | 5.91324244 | 3.354378713 | 1.706722473 |
| FUCA1 | 10407.2276 | 3.23248025 | 0.54723194 | 5.90696561 | 3.484664585 | 1.766611724 |
| ZSWIM6 | 711.737147 | 2.00801578 | 0.34038363 | 5.89927256 | 3.651076888 | 1.844319163 |
| ZNF462 | 138.16878 | -2.6175016 | 0.4438227 | -5.897629 | 3.687619792 | 1.849971593 |
| ALAD | 364.370251 | 1.64519981 | 0.27896163 | 5.8975845 | 3.688613872 | 1.849971593 |
| ST6GAL1 | 2509.29637 | -2.8470634 | 0.48313823 | -5.8928546 | 3.795803265 | 1.896956058 |
| NAPSB | 98.596739 | 4.93351676 | 0.84236124 | 5.85677083 | 4.719536518 | 2.350228770 |
| PRDM1 | 706.076467 | 1.89372807 | 0.32345163 | 5.85474883 | 4.777314456 | 2.370594590 |
| NCF1 | 885.177688 | 2.71781412 | 0.46535101 | 5.84035287 | 5.209037015 | 2.575722070 |
| DAPK1 | 1410.64769 | 1.20637099 | 0.20666347 | 5.83736915 | 5.303151283 | 2.613058016 |
| OAF | 137.224147 | -3.8966184 | 0.6681484 | -5.8319655 | 5.477824434 | 2.683873180 |
| SLC9A7P1 | 193.983133 | 3.4867248 | 0.59788712 | 5.8317443 | 5.485092949 | 2.683873180 |

| | | | | | | |
|---|---|---|---|---|---|---|
| CIPC | 296.629066 | -2.4529466 | 0.42075363 | -5.8298881 | 5.546454136 | 2.704474147 |
| PLEKHA1 | 346.628064 | -1.4872252 | 0.25583639 | -5.8131887 | 6.129391592 | 2.973618801 |
| CFD | 124.02905 | 3.45582018 | 0.59451104 | 5.8128781 | 6.140777984 | 2.973618801 |
| RAB6B | 87.8378674 | -6.1129781 | 1.05175717 | -5.8121573 | 6.167284818 | 2.976191776 |
| SMIM15 | 597.492017 | 1.26173822 | 0.21725675 | 5.80759031 | 6.337836927 | 3.048022053 |
| CUX1 | 1349.12064 | 1.90549548 | 0.32823533 | 5.80527236 | 6.426145783 | 3.079944206 |
| AQP9 | 1440.76665 | -5.6233589 | 0.96926708 | -5.8016609 | 6.566125401 | 3.136329898 |
| ENHO | 89.7375186 | 5.45146103 | 0.94083202 | 5.79429792 | 6.860757476 | 3.265953127 |
| KDM7A | 437.560517 | 2.40251265 | 0.41517105 | 5.78680197 | 7.173906646 | 3.403485507 |
| DBI | 4091.39022 | -2.5907991 | 0.44796477 | -5.7834884 | 7.316723051 | 3.459553596 |
| IFI30 | 15506.1452 | 1.83002966 | 0.31652856 | 5.78156262 | 7.400992351 | 3.479502150 |
| CLIC4 | 1913.99023 | -1.5994929 | 0.27666221 | -5.7813928 | 7.408467870 | 3.479502150 |
| OGT | 791.262682 | -2.0479017 | 0.3545413 | -5.7762008 | 7.640619690 | 3.576574077 |
| OSM | 77.1770603 | 2.93008604 | 0.50734329 | 5.77535186 | 7.679243721 | 3.582711613 |
| SNTB2 | 43.8959766 | -3.3654067 | 0.58404487 | -5.7622401 | 8.300481568 | 3.859723929 |
| MT-CO3 | 39371.7125 | -1.3520539 | 0.23469391 | -5.7609244 | 8.365452523 | 3.877097352 |
| MIOS | 427.553438 | 1.32238666 | 0.23027107 | 5.74273901 | 9.315725482 | 4.303313583 |
| SLC16A3 | 9629.54456 | 1.47425951 | 0.2570347 | 5.73564391 | 9.714256945 | 4.472698697 |
| DNASE2B | 43.8541595 | -8.1857066 | 1.42925833 | -5.7272408 | 1.020772423 | 4.684544817 |
| ASTN2 | 110.269661 | -4.3363642 | 0.75828439 | -5.7186515 | 1.073727708 | 4.911517333 |
| PODXL | 288.805897 | -5.0306088 | 0.88095261 | -5.7104193 | 1.126981400 | 5.138376559 |
| CD109 | 1001.1592 | -3.5662856 | 0.62473934 | -5.7084376 | 1.140179630 | 5.181728979 |
| GNA11 | 174.122527 | -2.8657042 | 0.50285203 | -5.6989015 | 1.205818558 | 5.462358068 |
| RAI14 | 426.804877 | -5.0908532 | 0.89395776 | -5.6947357 | 1.235631704 | 5.579413514 |
| CXCL5 | 47.3859503 | 5.27065016 | 0.92704336 | 5.6854408 | 1.304756775 | 5.872660065 |
| USP18 | 43.9719385 | -3.7642572 | 0.66374551 | -5.6712357 | 1.417711789 | 6.360679444 |
| NCOR2 | 3021.19119 | 1.4295253 | 0.2522888 | 5.66622571 | 1.459772072 | 6.528528411 |
| P2RY11 | 574.005109 | 1.4648894 | 0.25958591 | 5.64317755 | 1.669400258 | 7.421277160 |
| AHNAK | 2319.97831 | -2.0567234 | 0.36446565 | -5.64312 | 1.669959113 | 7.421277160 |
| ETS2 | 3493.02699 | 2.80760523 | 0.49775953 | 5.6404851 | 1.695717393 | 7.511974559 |
| SUSD6 | 2562.09761 | 1.44256126 | 0.25644107 | 5.62531292 | 1.851719970 | 8.177265267 |
| TMEM43 | 684.360991 | -1.5396806 | 0.27406601 | -5.6179187 | 1.932713447 | 8.504253018 |
| ENDOD1 | 197.302183 | -4.0817526 | 0.72661921 | -5.6174576 | 1.937877209 | 8.504253018 |
| IFNAR2 | 730.606443 | 1.18195762 | 0.21045171 | 5.61628909 | 1.951021344 | 8.535262536 |
| TRIM8 | 1481.37055 | 1.26244972 | 0.22545075 | 5.59966967 | 2.147606382 | 9.366098268 |
| CDC42EP3 | 216.072985 | -3.4507054 | 0.61652919 | -5.5969862 | 2.181100904 | 9.482724460 |
| S1PR2 | 756.644702 | 1.8939612 | 0.3386926 | 5.59197699 | 2.244985825 | 9.730350600 |
| DOCK3 | 57.5102551 | -7.0886719 | 1.26806308 | -5.5901572 | 2.268641586 | 9.802625782 |
| VWA1 | 75.5173647 | 2.49666222 | 0.44752843 | 5.5787791 | 2.422125982 | 1.043371631 |
| ACP5 | 7674.95494 | -3.9869201 | 0.71493201 | -5.5766423 | 2.452053189 | 1.053033117 |
| CYP27A1 | 1525.88994 | -5.0757408 | 0.91032174 | -5.5757658 | 2.464432547 | 1.055122752 |
| SERTAD2 | 474.652859 | -2.8489004 | 0.51132581 | -5.5715952 | 2.524174222 | 1.076331806 |
| IGF2BP2 | 258.84559 | -2.978678 | 0.53465243 | -5.5712419 | 2.529299268 | 1.076331806 |
| LY6E | 658.317134 | -3.5642844 | 0.64141898 | -5.5568738 | 2.746491353 | 1.165225923 |
| PALM2AKAP2 | 362.513312 | -3.5580429 | 0.6403951 | -5.556012 | 2.760079516 | 1.167463754 |
| GPX1 | 24795.1029 | 0.97214637 | 0.17504304 | 5.55375611 | 2.795955134 | 1.179087025 |
| MT-ND5 | 34472.9903 | -1.7720379 | 0.31920275 | -5.5514495 | 2.833106091 | 1.191176911 |
| TWSG1 | 669.669574 | -1.3683891 | 0.24651862 | -5.5508547 | 2.842762119 | 1.191668908 |

| | | | | | | |
|---|---|---|---|---|---|---|
| TGFBI | 48765.2266 | 4.70583461 | 0.84808221 | 5.54879532 | 2.876446818 | 1.202200674 |
| SUFU | 644.678963 | 1.30394395 | 0.23512382 | 5.54577572 | 2.926537639 | 1.219506470 |
| RNF152 | 40.8568951 | 5.51058309 | 0.99388668 | 5.54447827 | 2.948319389 | 1.224948200 |
| BIRC3 | 148.161116 | -2.6087344 | 0.4710195 | -5.538485 | 3.050993358 | 1.263867248 |
| MBD4 | 739.924999 | 1.96419332 | 0.35519801 | 5.52985457 | 3.204963755 | 1.323744294 |
| QKI | 3791.72385 | 1.66617372 | 0.30213071 | 5.51474468 | 3.492866555 | 1.438425954 |
| SOCS3 | 196.711605 | 3.17784752 | 0.57737456 | 5.50396176 | 3.713501211 | 1.524815716 |
| TATDN2 | 957.811595 | 1.25870799 | 0.22872012 | 5.50326743 | 3.728162397 | 1.526372727 |
| PNRC1 | 798.599531 | 1.53269006 | 0.27878831 | 5.49768415 | 3.848113738 | 1.570902942 |
| CXCL3 | 58.1254579 | 4.53977675 | 0.82650946 | 5.49270994 | 3.958125904 | 1.611129335 |
| TOX2 | 124.99884 | 3.26437859 | 0.59455609 | 5.49044678 | 4.009182864 | 1.627195230 |
| HDAC9 | 222.700524 | -3.2164768 | 0.58652867 | -5.4839208 | 4.160009730 | 1.683545148 |
| BMP7 | 41.2981127 | -6.1905576 | 1.12936944 | -5.4814283 | 4.219057583 | 1.702535219 |
| FBXL7 | 133.870201 | -4.7596693 | 0.86894002 | -5.4775579 | 4.312359209 | 1.735199437 |
| DOK5 | 21.1380237 | -8.1030149 | 1.48007462 | -5.4747341 | 4.381691215 | 1.758059706 |
| LRP5 | 238.96939 | 2.75609234 | 0.50364022 | 5.47234364 | 4.441225685 | 1.776869866 |
| SLC35F5 | 910.587096 | -1.2320962 | 0.22523345 | -5.4703074 | 4.492557677 | 1.792300780 |
| EHD3 | 86.2143431 | -6.9792292 | 1.27744452 | -5.4634303 | 4.670205810 | 1.857895189 |
| ETV5 | 1514.984 | 1.56940606 | 0.287534 | 5.45815821 | 4.810988860 | 1.908494818 |
| NPDC1 | 88.3322144 | 3.22764682 | 0.59173515 | 5.45454634 | 4.909803086 | 1.942207457 |
| ADARB1 | 372.582223 | -1.7346622 | 0.3181044 | -5.4531221 | 4.949305213 | 1.952334076 |
| KANK1 | 343.87079 | -1.6024312 | 0.29470436 | -5.4374194 | 5.405777943 | 2.126424080 |
| MPDZ | 158.36043 | -3.2276569 | 0.59389355 | -5.4347398 | 5.487644985 | 2.152597724 |
| ARMC9 | 348.459618 | -2.8238892 | 0.51967978 | -5.4339024 | 5.513474291 | 2.156705277 |
| SWAP70 | 1215.84423 | 1.25320039 | 0.23074605 | 5.43108065 | 5.601381195 | 2.179623002 |
| SLC11A1 | 2835.43965 | 2.69270856 | 0.4958011 | 5.43102577 | 5.603104066 | 2.179623002 |
| SIGLEC17P | 88.8111161 | -4.3809338 | 0.80919552 | -5.4139373 | 6.165377210 | 2.391723540 |
| THBS2 | 28.3137885 | -7.5522762 | 1.3954921 | -5.411909 | 6.235639432 | 2.412316378 |
| FGL2 | 751.904907 | 2.76005367 | 0.51066211 | 5.40485307 | 6.486154022 | 2.491345776 |
| NT5E | 82.4114104 | 5.73881642 | 1.06182541 | 5.40467047 | 6.492764938 | 2.491345776 |
| PHYH | 163.502105 | -1.6090974 | 0.29772409 | -5.4046599 | 6.493146437 | 2.491345776 |
| RAB38 | 93.8772462 | -4.2917821 | 0.79447894 | -5.4020086 | 6.589879745 | 2.521571696 |
| TEAD1 | 153.961918 | 3.17186019 | 0.58723549 | 5.40134276 | 6.614390108 | 2.524072834 |
| WBP1L | 1661.143 | 1.43607339 | 0.26644585 | 5.38973831 | 7.056035520 | 2.685309127 |
| HS3ST2 | 208.184609 | -4.3901465 | 0.8147171 | -5.3885533 | 7.102710807 | 2.689397696 |
| NRGN | 111.339671 | -2.2109293 | 0.41030566 | -5.3884933 | 7.105081146 | 2.689397696 |
| RAB40B | 138.883295 | -2.5952407 | 0.48241556 | -5.3796787 | 7.461889236 | 2.816863186 |
| GLT1D1 | 83.1269597 | 8.11581272 | 1.50958328 | 5.37619411 | 7.607677337 | 2.864198735 |
| XACT | 34.9587516 | 8.31242016 | 1.54650695 | 5.37496464 | 7.659771400 | 2.876100796 |
| GPD1 | 346.976498 | -5.6221702 | 1.04950636 | -5.3569663 | 8.463089956 | 3.169257926 |
| DAG1 | 207.892386 | -1.7980634 | 0.33598547 | -5.3516107 | 8.717477057 | 3.255838572 |
| CYTIP | 649.779754 | 2.55600475 | 0.4777718 | 5.34984434 | 8.802991374 | 3.279055911 |
| HAPLN4 | 677.685451 | -3.357682 | 0.62925784 | -5.3359399 | 9.505088136 | 3.531215679 |
| GIPC3 | 205.232198 | -3.3652122 | 0.63104288 | -5.3327789 | 9.672103645 | 3.583782361 |
| IFITM2 | 160.494012 | -2.9395307 | 0.55197746 | -5.3254543 | 1.007010172 | 3.721432594 |
| CST3 | 15567.6578 | 2.64227873 | 0.4963458 | 5.32346344 | 1.018099212 | 3.752537331 |
| MT-ND4L | 5093.72951 | -1.6906661 | 0.31787164 | -5.318707 | 1.045072914 | 3.841874068 |
| ARG2 | 98.643039 | -2.3052756 | 0.43363908 | -5.3161158 | 1.060056806 | 3.886782698 |

| | | | | | | |
|---|---|---|---|---|---|---|
| TBC1D12 | 261.322009 | 1.36771652 | 0.25751142 | 5.31128501 | 1.088549330 | 3.980858917 |
| LGALS9 | 3337.46194 | -2.2416528 | 0.42234766 | -5.3076009 | 1.110775418 | 4.051589401 |
| PRXL2A | 467.506855 | -2.680328 | 0.50537732 | -5.3036174 | 1.135301031 | 4.123887738 |
| KCNJ6-AS1 | 69.568358 | 7.86347413 | 1.48271488 | 5.3034297 | 1.136469810 | 4.123887738 |
| SERPINB9 | 721.383957 | 2.82790917 | 0.53539638 | 5.28189821 | 1.278522305 | 4.617089869 |
| IL18BP | 539.746852 | 2.78814593 | 0.52787448 | 5.28183506 | 1.278963155 | 4.617089869 |
| CBX4 | 154.499016 | 1.72128923 | 0.32598191 | 5.2803213 | 1.289575434 | 4.643463545 |
| RAB31 | 6729.44954 | 1.50239409 | 0.28489842 | 5.27343785 | 1.338916099 | 4.808797643 |
| CMKLR1 | 5713.08535 | 2.83022417 | 0.53676568 | 5.27273688 | 1.344041919 | 4.814893028 |
| CACNB3 | 36.8312735 | -3.2501118 | 0.6173962 | -5.2642239 | 1.407827877 | 5.030566636 |
| IKZF2 | 29.4902298 | -4.3096368 | 0.8203322 | -5.2535263 | 1.492143331 | 5.310082476 |
| FHL1 | 108.352958 | -5.0165524 | 0.95492538 | -5.2533449 | 1.493614311 | 5.310082476 |
| REPS2 | 183.016406 | 2.98545379 | 0.56851893 | 5.25128303 | 1.510433830 | 5.356318757 |
| NLRP3 | 822.475226 | 1.81558982 | 0.34594001 | 5.24827943 | 1.535263068 | 5.430654726 |
| DST | 97.1074411 | -2.157605 | 0.41181132 | -5.2393048 | 1.611825904 | 5.687153562 |
| SLC38A6 | 552.773311 | -1.8493672 | 0.35321549 | -5.2358045 | 1.642677981 | 5.781485436 |
| TESC | 841.009522 | 2.18420924 | 0.41733911 | 5.2336558 | 1.661898029 | 5.834508506 |
| WWTR1 | 63.5240854 | -6.8673014 | 1.3126001 | -5.2318306 | 1.678395233 | 5.877731737 |
| FAS | 62.6204807 | -2.6319815 | 0.50421167 | -5.2199932 | 1.789297340 | 6.250523023 |
| OXR1 | 347.593394 | -1.4228747 | 0.27291217 | -5.2136725 | 1.851381568 | 6.451352698 |
| SRSF9 | 1023.27278 | 1.02328698 | 0.1966111 | 5.20462472 | 1.943890394 | 6.756943764 |
| PPP1R1A | 124.01413 | -4.809563 | 0.92442617 | -5.2027552 | 1.963554829 | 6.808444561 |
| CES1 | 281.281699 | 11.321264 | 2.17768009 | 5.19877279 | 2.006085550 | 6.938783101 |
| PALD1 | 1052.37686 | 2.87558414 | 0.55347417 | 5.19551639 | 2.041523493 | 7.044008458 |
| ST8SIA4 | 1409.68567 | 2.44512637 | 0.47118524 | 5.18931015 | 2.110745759 | 7.265000660 |
| CTDSPL | 62.5589376 | -3.698992 | 0.71423639 | -5.1789465 | 2.231424798 | 7.661588862 |
| CAMK1D | 327.765899 | 1.94846662 | 0.37630461 | 5.17789727 | 2.244007322 | 7.685998738 |
| MERTK | 3236.51147 | 1.95623705 | 0.37838642 | 5.16994514 | 2.341627315 | 8.000844863 |
| PCSK6 | 355.270593 | -4.5867724 | 0.88786707 | -5.1660576 | 2.390831897 | 8.149138916 |
| CD99L2 | 1532.96605 | 1.27893343 | 0.24778639 | 5.16143529 | 2.450636027 | 8.332755868 |
| UBE2J1 | 1979.18008 | -1.0988737 | 0.21304744 | -5.1578828 | 2.497579188 | 8.471861000 |
| EPS8 | 1301.12883 | -1.2138935 | 0.23542388 | -5.1562039 | 2.520064466 | 8.527533809 |
| HTR2B | 79.9393242 | -5.7771601 | 1.12072474 | -5.154843 | 2.538434716 | 8.569047769 |
| SLC23A2 | 2304.99328 | 1.0534619 | 0.20439868 | 5.15395636 | 2.550473193 | 8.589039580 |
| SECTM1 | 258.738475 | 3.93227265 | 0.76387275 | 5.14781117 | 2.635435973 | 8.853929995 |
| TLR8 | 392.256485 | 2.92347675 | 0.57036701 | 5.12560634 | 2.965817051 | 9.940088032 |
| CDKN1C | 102.994119 | 4.08648299 | 0.79929141 | 5.11263216 | 3.177005617 | 1.062254520 |
| HLA-DMA | 1301.58149 | 3.58173926 | 0.70067186 | 5.11186398 | 3.189954887 | 1.064050747 |
| TNFSF11 | 65.6822967 | 9.22228634 | 1.80494208 | 5.10946387 | 3.230742692 | 1.074859073 |
| ACTA2 | 61.7158605 | -6.1745941 | 1.20855789 | -5.1090594 | 3.237665656 | 1.074859073 |
| SLCO4C1 | 95.4188528 | 2.28158632 | 0.44678179 | 5.10671284 | 3.278113264 | 1.085720390 |
| TNKS1BP1 | 305.240233 | -1.6422367 | 0.32174018 | -5.1042326 | 3.321395799 | 1.097467322 |
| IL1RAP | 1395.25693 | 3.33732876 | 0.65437721 | 5.10000762 | 3.396397994 | 1.119615423 |
| TMEM150B | 48.6520892 | 3.27958566 | 0.64399984 | 5.0925256 | 3.533249852 | 1.162000648 |
| MPG | 290.951392 | -1.8716051 | 0.36784234 | -5.0880633 | 3.617387079 | 1.186891746 |
| GRAMD1B | 47.463083 | 3.30097226 | 0.64896084 | 5.08655076 | 3.646341632 | 1.193603159 |
| ITGB1BP1 | 774.835125 | -2.6494652 | 0.52099997 | -5.0853461 | 3.669563126 | 1.198411046 |
| YWHAQ | 2032.38507 | -1.259449 | 0.24794739 | -5.0795012 | 3.784272068 | 1.233005398 |

| | | | | | | |
|---|---|---|---|---|---|---|
| SPNS3 | 53.4796474 | 3.24802018 | 0.64026632 | 5.07292054 | 3.917561185 | 1.273479438 |
| TTC28 | 50.1502706 | -3.6170819 | 0.71374553 | -5.0677472 | 4.025514966 | 1.305549807 |
| SLCO2B1 | 8082.9575 | 1.31641246 | 0.25993281 | 5.06443353 | 4.096165244 | 1.325402039 |
| FCGR2B | 67.953335 | 6.79100686 | 1.34143045 | 5.06251133 | 4.137694695 | 1.335761990 |
| SMURF2 | 699.144754 | 1.89953106 | 0.37557309 | 5.05768683 | 4.243725866 | 1.364941478 |
| STUM | 29.6542684 | -4.6970393 | 0.9287245 | -5.0575163 | 4.247521371 | 1.364941478 |
| B3GNT7 | 319.412011 | 2.65879642 | 0.5260493 | 5.05427235 | 4.320347423 | 1.385174403 |
| ATP2B2 | 46.3267517 | -8.2661339 | 1.63718249 | -5.0489997 | 4.441294677 | 1.420708454 |
| OMA1 | 77.8821446 | -2.2066881 | 0.43749279 | -5.0439417 | 4.560382847 | 1.455487643 |
| LANCL3 | 55.4386523 | -4.0848746 | 0.81015556 | -5.0420867 | 4.604825821 | 1.466339433 |
| FAM81A | 21.141817 | -8.1016758 | 1.60752996 | -5.0398288 | 4.659484254 | 1.480387723 |
| FYCO1 | 118.868603 | -3.3951472 | 0.67457014 | -5.0330529 | 4.827297219 | 1.527091562 |
| CALM3 | 5936.72338 | -1.3589221 | 0.27000156 | -5.0330158 | 4.828232242 | 1.527091562 |
| RBP1 | 381.504232 | 4.08125283 | 0.81210234 | 5.02554002 | 5.020178606 | 1.584232992 |
| LONRF1 | 135.3548 | -1.7930729 | 0.35690468 | -5.0239547 | 5.061819209 | 1.593792088 |
| OSBP2 | 22.0437591 | -8.1619294 | 1.62746449 | -5.0151198 | 5.300043151 | 1.665067247 |
| ADRA2B | 281.552009 | -10.875065 | 2.17012953 | -5.0112514 | 5.407721476 | 1.693241458 |
| ITGAX | 3195.76182 | -2.5230673 | 0.50350234 | -5.0110338 | 5.413839027 | 1.693241458 |
| CLDN1 | 510.572943 | 4.0842356 | 0.81514369 | 5.01044868 | 5.430327998 | 1.694624357 |
| GRIN2D | 322.568376 | 2.59772236 | 0.51953996 | 5.00004342 | 5.731740472 | 1.784719100 |
| PCNX1 | 1188.34346 | 1.22374183 | 0.24483465 | 4.998238 | 5.785654795 | 1.797521024 |
| ZMIZ1 | 6570.48514 | 1.09266611 | 0.21894243 | 4.99065502 | 6.017488463 | 1.863140191 |
| CXCR3 | 36.8465703 | 5.93209038 | 1.18869187 | 4.99043572 | 6.024324519 | 1.863140191 |
| XKRX | 36.382362 | -8.8852007 | 1.7805869 | -4.9900405 | 6.036664439 | 1.863140191 |
| TGFBR1 | 2581.41874 | 1.66730022 | 0.33416596 | 4.9894377 | 6.055530040 | 1.864864218 |
| PMP22 | 908.503389 | 2.27028569 | 0.45506441 | 4.98893265 | 6.071380819 | 1.865654285 |
| ADGRE3 | 39.3052187 | 6.04542243 | 1.21226681 | 4.9868745 | 6.136389834 | 1.881513590 |
| THBD | 2050.5899 | 3.45563247 | 0.69395836 | 4.97959628 | 6.371704928 | 1.949408547 |
| FUT8 | 596.517059 | 1.2585421 | 0.25293273 | 4.97579771 | 6.497948063 | 1.983710535 |
| TNFSF15 | 258.854786 | -1.6605158 | 0.33384437 | -4.9739218 | 6.561179085 | 1.998668934 |
| GPR34 | 2879.54587 | 1.49195629 | 0.30014351 | 4.97080985 | 6.667381546 | 2.026624222 |
| ENSG000002 | 61.4150786 | -2.1553118 | 0.43368349 | -4.9697806 | 6.702870470 | 2.033011015 |
| UBE2L6 | 557.608768 | -1.4538038 | 0.29263128 | -4.9680398 | 6.763308151 | 2.046921042 |
| TNNI2 | 87.2540607 | -4.2478404 | 0.85524813 | -4.966793 | 6.806917564 | 2.055689104 |
| TTC7B | 276.242607 | -1.7267238 | 0.34779465 | -4.9647795 | 6.877915185 | 2.072673024 |
| SLITRK4 | 74.5042147 | -2.6491781 | 0.53402107 | -4.960812 | 7.019910406 | 2.110933658 |
| TPCN2 | 259.165782 | -1.7503436 | 0.35304674 | -4.957824 | 7.128710981 | 2.139070263 |
| SLC4A8 | 167.026172 | -3.2786807 | 0.66147331 | -4.9566333 | 7.172516547 | 2.147625796 |
| ENSG000002 | 26.6122457 | 7.92152953 | 1.59861476 | 4.9552461 | 7.223879874 | 2.158403086 |
| DPYSL3 | 148.999643 | -4.7728966 | 0.96461395 | -4.9479863 | 7.498519117 | 2.235704967 |
| ME3 | 44.3941091 | -5.2288777 | 1.05740126 | -4.945027 | 7.613332749 | 2.265127792 |
| PDGFRA | 43.8957536 | -8.1866423 | 1.65861866 | -4.9358195 | 7.981491153 | 2.369642288 |
| GJA1 | 163.580561 | -4.6514653 | 0.94252707 | -4.9351 | 8.010971056 | 2.373376931 |
| CDIP1 | 497.883737 | 1.17546358 | 0.2382046 | 4.93468043 | 8.028210102 | 2.373476936 |
| PCED1B | 49.3433514 | 3.28419055 | 0.66593126 | 4.93172606 | 8.150616816 | 2.404603192 |
| PDE4A | 847.067596 | 1.83423722 | 0.37207557 | 4.92974372 | 8.233755836 | 2.424038431 |
| INAVA | 58.8926768 | 5.6491861 | 1.14619502 | 4.92864304 | 8.280270045 | 2.432632473 |
| CD55 | 285.984212 | -2.1028743 | 0.42759595 | -4.9179004 | 8.747741885 | 2.564604160 |

| | | | | | | |
|---|---|---|---|---|---|---|
| MYOF | 945.959964 | -3.15068 | 0.64122623 | -4.9135232 | 8.945416095 | 2.612639654 |
| NISCH | 1466.13616 | 1.59967541 | 0.32557077 | 4.91344909 | 8.948797789 | 2.612639654 |
| CBX1 | 568.222778 | 1.24722763 | 0.25418345 | 4.90680104 | 9.257385625 | 2.697125857 |
| RAB11FIP4 | 324.975213 | -2.953235 | 0.60339134 | -4.8943941 | 9.860905717 | 2.867012401 |
| MFHAS1 | 903.25349 | -2.5677335 | 0.52538508 | -4.8873362 | 1.022094581 | 2.961297294 |
| ALOX5AP | 8107.02329 | 4.2406514 | 0.8677035 | 4.88721253 | 1.022736728 | 2.961297294 |
| LINGO1 | 60.5538945 | -7.1906674 | 1.47367522 | -4.8794112 | 1.064030149 | 3.074521683 |
| ITGBL1 | 50.0181625 | -6.9268294 | 1.42013299 | -4.877592 | 1.073887417 | 3.096632649 |
| 5_8S_rRNA | 48.9930724 | 21.4890104 | 4.40839584 | 4.87456462 | 1.090486864 | 3.138054720 |
| TFRC | 4076.46105 | -3.5819084 | 0.7351174 | -4.8725665 | 1.101577791 | 3.163488124 |
| JDP2 | 450.1416 | 2.00382307 | 0.41165502 | 4.8677241 | 1.128907838 | 3.235357708 |
| GALNT3 | 77.9805136 | 3.27924502 | 0.67403423 | 4.86510159 | 1.143980304 | 3.271876868 |
| NEXMIF | 31.5186745 | 3.47332621 | 0.71446858 | 4.86141209 | 1.165513377 | 3.326687877 |
| ITGA9 | 336.678607 | 1.85945847 | 0.38326295 | 4.85165207 | 1.224372779 | 3.487599786 |
| PLS3 | 123.882068 | -4.976113 | 1.02642198 | -4.8480187 | 1.247006184 | 3.544880131 |
| SIGLEC15 | 228.698637 | -3.8379573 | 0.79191758 | -4.84641 | 1.257155870 | 3.566513108 |
| PTPRG | 405.851416 | -2.9014449 | 0.59876803 | -4.8456911 | 1.261717218 | 3.572236874 |
| TPR | 238.445525 | -1.6948864 | 0.34991115 | -4.8437621 | 1.274034368 | 3.599852038 |
| CAMKK2 | 235.021778 | -1.390338 | 0.28763103 | -4.8337553 | 1.339812648 | 3.778110245 |
| LARGE1 | 52.1042164 | -2.7490862 | 0.56903579 | -4.8311305 | 1.357599692 | 3.820595688 |
| ADCY8 | 59.2249517 | -5.3860701 | 1.11643325 | -4.8243547 | 1.404572438 | 3.944882150 |
| QPCT | 86.7591226 | -2.1624149 | 0.44830815 | -4.8235012 | 1.410599512 | 3.953901987 |
| TSHZ3 | 283.214657 | 3.35434776 | 0.69594636 | 4.81983667 | 1.436758006 | 4.018175311 |
| CEACAM4 | 28.8631717 | 4.97259625 | 1.03176802 | 4.81949059 | 1.439252425 | 4.018175311 |
| MCEMP1 | 70.4651644 | 7.88522287 | 1.6372248 | 4.81621269 | 1.463085738 | 4.076609728 |
| ENSG000002 | 25.1504531 | 7.8356152 | 1.62787186 | 4.8134103 | 1.483762139 | 4.126034004 |
| ACSL5 | 317.057425 | -1.4337831 | 0.29798139 | -4.8116531 | 1.496869753 | 4.153130386 |
| NCR3LG1 | 267.440572 | 2.20597965 | 0.45849849 | 4.81131283 | 1.499421139 | 4.153130386 |
| FILIP1L | 115.018672 | 1.62358725 | 0.33759574 | 4.8092646 | 1.514865712 | 4.187649447 |
| DIRAS2 | 36.4391843 | -7.9170339 | 1.64775036 | -4.8047532 | 1.549424706 | 4.274768399 |
| GBP2 | 151.241513 | -1.5355244 | 0.31976002 | -4.8021152 | 1.569983428 | 4.322995546 |
| AGPAT4 | 127.814866 | -1.9611473 | 0.40844485 | -4.8014984 | 1.574828022 | 4.327849300 |
| LSS | 666.882841 | -1.7667364 | 0.36805334 | -4.8002184 | 1.584926999 | 4.347095674 |
| MT-ND4 | 56935.6679 | -1.5090319 | 0.31473789 | -4.794567 | 1.630265431 | 4.462732446 |
| MREG | 119.102945 | -4.7955584 | 1.00039485 | -4.7936656 | 1.637611418 | 4.474120069 |
| LRRC75A | 73.6843946 | -2.800837 | 0.58434829 | -4.7930952 | 1.642276124 | 4.478152157 |
| PXK | 299.729009 | 2.24758728 | 0.46944751 | 4.78772859 | 1.686795533 | 4.590633657 |
| MBOAT1 | 118.661396 | -1.9468631 | 0.40704762 | -4.7828877 | 1.727946953 | 4.693531733 |
| TYMP | 1820.36374 | 1.48544231 | 0.31096099 | 4.77694105 | 1.779819241 | 4.825096834 |
| RNASET2 | 2490.04571 | 1.47729316 | 0.30986682 | 4.76750992 | 1.865168161 | 5.046735353 |
| STON2 | 32.4882665 | -3.6043394 | 0.75622136 | -4.7662492 | 1.876871353 | 5.068635465 |
| KAZN | 93.1500546 | -4.4004434 | 0.92374606 | -4.7636938 | 1.900808891 | 5.123427882 |
| SLA2 | 66.4187825 | 7.80158967 | 1.63800804 | 4.76285189 | 1.908759400 | 5.135001581 |
| CR1 | 27.4401271 | -4.580172 | 0.96195098 | -4.7613362 | 1.923153445 | 5.163832474 |
| COL6A2 | 3489.57682 | -6.3354185 | 1.33142497 | -4.7583744 | 1.951582924 | 5.230167749 |
| SHISA2 | 198.619368 | -10.371283 | 2.18100156 | -4.7552846 | 1.981670265 | 5.300684864 |
| FAT1 | 288.484307 | -4.8475332 | 1.01949544 | -4.7548356 | 1.986079906 | 5.302380252 |
| DPP4 | 365.333672 | -4.0692333 | 0.85601075 | -4.7537175 | 1.997100305 | 5.320400514 |

| | | | | | | |
|---|---|---|---|---|---|---|
| NOTCH1 | 1359.26037 | 1.68472173 | 0.35442582 | 4.75338319 | 2.000406944 | 5.320400514 |
| OCA2 | 49.7912841 | 8.8230359 | 1.85862905 | 4.7470666 | 2.063880092 | 5.478840856 |
| TNFRSF21 | 2576.87569 | 2.5770146 | 0.54381263 | 4.73879137 | 2.149967215 | 5.696601813 |
| CSF1 | 1756.86428 | -4.3082901 | 0.90968621 | -4.7360178 | 2.179584512 | 5.764200623 |
| ATP2B4 | 1527.73332 | -0.999922 | 0.2113934 | -4.7301477 | 2.243565086 | 5.922252727 |
| AGPAT5 | 203.133313 | -1.4286327 | 0.30210238 | -4.7289687 | 2.256631167 | 5.945566883 |
| HSD3B7 | 851.065014 | -3.3724456 | 0.71334273 | -4.7276652 | 2.271162506 | 5.972647017 |
| CTIF | 352.722863 | 2.13436581 | 0.45225853 | 4.71934892 | 2.366006891 | 6.210436406 |
| TBX15 | 78.0396336 | 9.47089817 | 2.009232 | 4.7136907 | 2.432698449 | 6.365581077 |
| ZMIZ1-AS1 | 89.690702 | -4.7171477 | 1.00075969 | -4.7135668 | 2.434178621 | 6.365581077 |
| TRAP1 | 372.978568 | -1.6239736 | 0.34480489 | -4.7098335 | 2.479192923 | 6.471246509 |
| FCMR | 2815.93089 | -2.4052254 | 0.51073168 | -4.7093719 | 2.484812791 | 6.473882379 |
| SKI | 1303.67218 | 2.09161695 | 0.44509461 | 4.69926376 | 2.611010537 | 6.790077960 |
| MT-CO1 | 99814.369 | -1.1383196 | 0.2424574 | -4.6949262 | 2.667029504 | 6.911372477 |
| ENSG000002 | 27.9960414 | 20.6989017 | 4.40915412 | 4.69452895 | 2.672217653 | 6.911372477 |
| ADAMDEC1 | 697.740876 | -1.6282437 | 0.3468397 | -4.6945137 | 2.672417044 | 6.911372477 |
| PLAU | 748.922973 | 3.00673744 | 0.6433449 | 4.67360109 | 2.959639177 | 7.640112678 |
| MYO1E | 1893.48841 | -1.8452271 | 0.39487666 | -4.67292 | 2.969474167 | 7.651435915 |
| FGD3 | 89.2723897 | 1.79378616 | 0.38399312 | 4.67140182 | 2.991510809 | 7.694100053 |
| AK1 | 288.06309 | 1.67632321 | 0.35888393 | 4.67093421 | 2.998329714 | 7.697540069 |
| LHFPL2 | 4456.33312 | 2.15109658 | 0.46083057 | 4.66786863 | 3.043403819 | 7.798999969 |
| TUBA8 | 25.1432788 | -8.3518692 | 1.79414099 | -4.6550796 | 3.238555137 | 8.283976282 |
| SEC14L1 | 1823.8873 | 1.00288825 | 0.21571805 | 4.64906974 | 3.334354648 | 8.513516787 |
| P3H2 | 296.560413 | 1.90974398 | 0.41211101 | 4.63405225 | 3.585764241 | 9.138818011 |
| SCD | 11051.5678 | -1.6149386 | 0.34857371 | -4.6329904 | 3.604212019 | 9.147026005 |
| HSPB1 | 913.226317 | -2.3137715 | 0.49942877 | -4.6328358 | 3.606906347 | 9.147026005 |
| PKD1 | 199.356464 | -1.9152829 | 0.4134231 | -4.6327429 | 3.608525533 | 9.147026005 |
| CTTNBP2 | 83.1209797 | 2.46618826 | 0.53319793 | 4.62527723 | 3.740976923 | 9.462182400 |
| FLT1 | 124.041122 | -3.6843457 | 0.79661859 | -4.6249808 | 3.746331563 | 9.462182400 |
| OPTN | 363.429793 | -3.6330392 | 0.78561388 | -4.624459 | 3.755773448 | 9.468999377 |
| F3 | 68.2597002 | -5.139091 | 1.11166086 | -4.6228946 | 3.784220065 | 9.508771281 |
| DOCK10 | 2109.12249 | 1.17585555 | 0.25435745 | 4.6228469 | 3.785090896 | 9.508771281 |
| RGS4 | 30.517879 | 8.11479376 | 1.75625678 | 4.6205053 | 3.828065616 | 9.596844028 |
| HLA-DQB1 | 2341.06982 | 8.10633711 | 1.75454481 | 4.62019382 | 3.833817204 | 9.596844028 |
| TOP6BL | 27.3977135 | -3.2430386 | 0.70209177 | -4.6191092 | 3.853909468 | 9.623891153 |
| IRF4 | 106.962279 | 5.0666267 | 1.09694041 | 4.61887142 | 3.858328505 | 9.623891153 |
| COL18A1 | 175.480742 | -2.029635 | 0.43956284 | -4.6173945 | 3.885882086 | 9.675433003 |
| CCNJL | 31.0312215 | 5.11804289 | 1.11081022 | 4.60748633 | 4.075659935 | 0.0001013 |
| SLC16A6 | 338.256482 | 2.27793227 | 0.49455416 | 4.60603195 | 4.104253672 | 0.00010181 |
| TMEM51 | 977.546704 | 1.94692256 | 0.42271937 | 4.6057094 | 4.110621019 | 0.00010181 |
| DLST | 1086.42244 | -1.0602057 | 0.23024426 | -4.6046998 | 4.130612673 | 0.00010212 |
| SEPTIN3 | 71.2171497 | 2.99547879 | 0.65153413 | 4.59757773 | 4.274311772 | 0.00010549 |
| CD244 | 48.4735933 | 4.11056428 | 0.89459004 | 4.59491399 | 4.329278060 | 0.00010666 |
| ABLIM3 | 173.563254 | -2.6394952 | 0.57468909 | -4.5929099 | 4.371078241 | 0.0001075 |
| ENSG000002 | 118.253795 | 1.93058259 | 0.42128313 | 4.58262496 | 4.591751530 | 0.00011273 |
| NCF1B | 345.338691 | 3.57845329 | 0.78097978 | 4.58200502 | 4.605388404 | 0.00011287 |
| FAM234A | 406.739049 | -1.0189052 | 0.22260794 | -4.5771286 | 4.714016755 | 0.00011533 |
| CYSTM1 | 677.888301 | -3.0000783 | 0.65586899 | -4.5742036 | 4.780349138 | 0.00011675 |

| | | | | | | |
|---|---|---|---|---|---|---|
| ZFAND3 | 1145.52685 | 1.09405971 | 0.23932604 | 4.57141951 | 4.844313547 | 0.00011811 |
| HMGA2 | 353.78907 | -3.134163 | 0.68576289 | -4.5703304 | 4.869557768 | 0.00011852 |
| SH3RF2 | 38.4916251 | -6.4910626 | 1.42060693 | -4.5692179 | 4.895475543 | 0.00011894 |
| ABCB4 | 296.934778 | 1.23530971 | 0.27043035 | 4.56794029 | 4.925402884 | 0.00011946 |
| ENSG000002 | 25.444989 | -6.5083834 | 1.42534426 | -4.5661835 | 4.966840512 | 0.00012026 |
| RADX | 21.5041111 | -8.1261757 | 1.7799833 | -4.5653101 | 4.987565430 | 0.00012055 |
| VDR | 474.084759 | 1.38104377 | 0.30272265 | 4.5620761 | 5.065028135 | 0.00012221 |
| LINC00504 | 25.9120065 | -6.4928015 | 1.42510153 | -4.5560273 | 5.213016441 | 0.00012557 |
| KLHL4 | 31.4993467 | -6.2218353 | 1.36680832 | -4.5520906 | 5.311545785 | 0.00012772 |
| IQCK | 20.2949092 | -3.8683215 | 0.85008752 | -4.5504979 | 5.351912617 | 0.00012847 |
| BST1 | 306.161482 | 1.90953767 | 0.42059513 | 4.54008508 | 5.623153199 | 0.00013475 |
| FMN1 | 522.004403 | 1.38536248 | 0.30519225 | 4.53931084 | 5.643838444 | 0.00013502 |
| ZBTB7C | 176.538337 | -3.2639476 | 0.72040491 | -4.5307126 | 5.878506945 | 0.00014039 |
| PDE6G | 28.4785233 | -5.2882523 | 1.16734598 | -4.5301499 | 5.894183829 | 0.00014053 |
| AMDHD2 | 414.41297 | -1.5342866 | 0.33875417 | -4.5292036 | 5.920642408 | 0.00014092 |
| OS9 | 4335.07541 | 0.94936663 | 0.20994021 | 4.52208106 | 6.123459989 | 0.0001455 |
| PIK3C2B | 800.651494 | 1.76102897 | 0.38953327 | 4.52086924 | 6.158622276 | 0.00014609 |
| KYNU | 528.96039 | 1.99925976 | 0.44234667 | 4.51966724 | 6.193690390 | 0.00014667 |
| SSR1 | 2982.00437 | 0.92650308 | 0.20516469 | 4.51589933 | 6.304861321 | 0.00014906 |
| NCKAP1 | 87.5750884 | -4.8615597 | 1.07665979 | -4.5154094 | 6.319457073 | 0.00014915 |
| BST2 | 963.436696 | 1.03595983 | 0.2297592 | 4.50889386 | 6.516649988 | 0.00015355 |
| MOV10 | 295.243689 | -1.4733895 | 0.3268781 | -4.5074585 | 6.560874639 | 0.00015433 |
| SH3PXD2B | 1628.87927 | 1.53321927 | 0.34022811 | 4.50644495 | 6.592277885 | 0.00015481 |
| CD4 | 5887.29931 | 1.59675938 | 0.35456128 | 4.5034793 | 6.684988143 | 0.00015672 |
| RERE | 320.203658 | -1.9601462 | 0.43578968 | -4.4979178 | 6.862221547 | 0.00016061 |
| ZNF516 | 643.153815 | 1.69465647 | 0.37688414 | 4.49649183 | 6.908383166 | 0.00016142 |
| ISY1 | 195.697515 | 1.26030979 | 0.28046301 | 4.49367562 | 7.000423674 | 0.0001633 |
| HSPA12A | 40.4729875 | -7.1926252 | 1.60125292 | -4.4918732 | 7.059943837 | 0.00016417 |
| MT-ND6 | 80.2751745 | -1.9000424 | 0.42299931 | -4.4918333 | 7.061268886 | 0.00016417 |
| SEPTIN11 | 1077.35393 | -1.6870032 | 0.37592241 | -4.4876366 | 7.201762923 | 0.00016716 |
| RAP1GAP | 24.158181 | -8.2942259 | 1.84972448 | -4.4840331 | 7.324527890 | 0.00016971 |
| KLHL15 | 183.581154 | 1.34461074 | 0.29988801 | 4.48370953 | 7.335650381 | 0.00016971 |
| KCNK6 | 399.109839 | 1.24146055 | 0.27695335 | 4.48256189 | 7.375224410 | 0.00017035 |
| S100A10 | 1529.90923 | 2.827203 | 0.63092455 | 4.4810477 | 7.427750710 | 0.00017128 |
| UQCC5 | 334.340244 | 1.08136239 | 0.24135029 | 4.4804686 | 7.447933833 | 0.00017146 |
| MAST4 | 55.165057 | -6.6444964 | 1.48484547 | -4.474874 | 7.645637397 | 0.00017572 |
| CDYL2 | 50.4329579 | -3.6356953 | 0.81326146 | -4.4705123 | 7.803246714 | 0.00017905 |
| ATG7 | 884.482179 | -1.0091026 | 0.22587231 | -4.4675797 | 7.910952894 | 0.00018123 |
| CHD9 | 558.872816 | 1.26868181 | 0.28430261 | 4.46243464 | 8.103365037 | 0.00018533 |
| DBNDD2 | 456.432859 | 1.4725019 | 0.33000425 | 4.46206956 | 8.117186492 | 0.00018535 |
| DNAJB6 | 1819.36576 | 1.13928102 | 0.25571288 | 4.45531329 | 8.377077288 | 0.00019097 |
| MAP2 | 15.42342 | -7.6465585 | 1.71662859 | -4.4544047 | 8.412628100 | 0.00019147 |
| VAV3 | 605.478573 | 1.99451402 | 0.44803591 | 4.45168344 | 8.519970332 | 0.0001936 |
| GPBAR1 | 141.833656 | 3.55391638 | 0.79906312 | 4.44760408 | 8.683338782 | 0.000197 |
| MICALL2 | 58.4509026 | -2.4422246 | 0.54928853 | -4.4461598 | 8.741892626 | 0.000198 |
| SHTN1 | 606.211621 | -2.0942128 | 0.47127767 | -4.4436919 | 8.842819868 | 0.00019997 |
| TAL1 | 927.510169 | 2.49127968 | 0.56087577 | 4.44176733 | 8.922300653 | 0.00020144 |
| ACOT1 | 128.375683 | -3.0600001 | 0.68948779 | -4.4380773 | 9.076601275 | 0.0002046 |

| | | | | | | |
|---|---|---|---|---|---|---|
| SRPX | 27.2025493 | 4.46851424 | 1.00699277 | 4.43748393 | 9.101651435 | 0.00020483 |
| HYAL2 | 451.646787 | -1.2310822 | 0.27768739 | -4.4333386 | 9.278497613 | 0.00020848 |
| MALAT1 | 565.309634 | -2.0517179 | 0.46325905 | -4.4288783 | 9.472445486 | 0.00021249 |
| SARDH | 35.8491774 | -7.8944747 | 1.78264436 | -4.4285192 | 9.488229454 | 0.00021251 |
| TBC1D2B | 384.977384 | -1.0154544 | 0.229326 | -4.4279951 | 9.511305432 | 0.00021269 |
| LIMD2 | 916.453095 | -1.5778692 | 0.35679048 | -4.422397 | 9.761188250 | 0.00021793 |
| NAPSB.1 | 21.0626117 | 7.5829921 | 1.71557299 | 4.42009297 | 9.865844544 | 0.00021991 |
| CYP4X1 | 51.92772 | -4.484663 | 1.01555926 | -4.415954 | 1.005654522 | 0.00022368 |
| DDIT4 | 537.126137 | 3.37334189 | 0.76393572 | 4.4157405 | 1.006647891 | 0.00022368 |
| RGS19 | 1575.38949 | 1.09967058 | 0.24926948 | 4.41157333 | 1.026221865 | 0.00022767 |
| TPST2 | 920.425744 | 1.03519719 | 0.23493403 | 4.40633146 | 1.051360421 | 0.00023287 |
| CRACD | 12.4532911 | -7.3387501 | 1.66721088 | -4.4018127 | 1.073502002 | 0.0002374 |
| UBA7 | 243.267126 | -2.0082311 | 0.45662292 | -4.3980077 | 1.092491082 | 0.00024122 |
| TFAP2C | 72.0625232 | 3.03923273 | 0.69115598 | 4.39731811 | 1.095966602 | 0.00024132 |
| ATOH8 | 125.258322 | 4.47430196 | 1.01752459 | 4.39724209 | 1.096350409 | 0.00024132 |
| FCHSD2 | 555.613954 | 1.36881759 | 0.31155536 | 4.39349711 | 1.115416731 | 0.00024513 |
| NAIPP2 | 362.563775 | 3.12906812 | 0.71254674 | 4.39138645 | 1.126301507 | 0.00024714 |
| NEO1 | 149.049227 | -2.0145887 | 0.45892331 | -4.3898156 | 1.134468316 | 0.00024854 |
| QSOX1 | 3154.84035 | 1.69657644 | 0.38771613 | 4.37582111 | 1.209762036 | 0.00026462 |
| EIF4A2 | 545.944742 | -1.0580006 | 0.24180236 | -4.3754766 | 1.211674204 | 0.00026463 |
| ELL | 389.222801 | 1.5842152 | 0.36220513 | 4.37380666 | 1.220985238 | 0.00026625 |
| RALGAPA1 | 152.708987 | -1.5319121 | 0.350361 | -4.3723819 | 1.228982841 | 0.00026755 |
| B4GALT2 | 118.549827 | -1.672723 | 0.38259329 | -4.3720657 | 1.230764754 | 0.00026755 |
| TNFRSF10D | 431.298015 | 1.38653022 | 0.31727955 | 4.37005852 | 1.242132454 | 0.0002696 |
| RB1 | 2517.668 | 1.9965923 | 0.45702212 | 4.36869944 | 1.249886422 | 0.00027087 |
| UBXN7 | 380.195445 | -1.0469516 | 0.23968725 | -4.3679903 | 1.253950736 | 0.00027133 |
| MATK | 1507.77213 | -2.4965895 | 0.5716229 | -4.3675462 | 1.256502045 | 0.00027146 |
| ACTR1B | 357.48388 | -1.4930337 | 0.34190009 | -4.3668714 | 1.260388939 | 0.00027148 |
| TIMP3 | 3669.74505 | -3.4757462 | 0.79593632 | -4.3668646 | 1.260428018 | 0.00027148 |
| PSD3 | 177.913053 | -2.4719716 | 0.56663622 | -4.3625372 | 1.285627144 | 0.00027648 |
| TMCC2 | 20.9491723 | -7.1116765 | 1.63143976 | -4.3591413 | 1.305737630 | 0.00028021 |
| TLNRD1 | 1221.12093 | 1.26476916 | 0.29015533 | 4.3589383 | 1.306949504 | 0.00028021 |
| CLEC7A | 1157.79588 | 1.45866725 | 0.33468969 | 4.35826767 | 1.310959906 | 0.00028064 |
| HDGFL3 | 68.7309638 | -2.8473942 | 0.65364403 | -4.3561848 | 1.323490305 | 0.00028289 |
| CENPV | 29.7600247 | -3.4644493 | 0.79570841 | -4.3539182 | 1.337256164 | 0.0002854 |
| RBP4 | 28.6533876 | 3.32483367 | 0.76373341 | 4.35339558 | 1.340449580 | 0.00028564 |
| ABCA3 | 47.0725446 | -3.5160798 | 0.80810719 | -4.3510067 | 1.355139132 | 0.00028834 |
| SEZ6 | 28.3088344 | 4.23612605 | 0.97485553 | 4.34538852 | 1.390293634 | 0.00029518 |
| MT-ND1 | 22846.2268 | -1.027615 | 0.23649452 | -4.3451959 | 1.391513943 | 0.00029518 |
| SLC43A2 | 3968.07828 | 2.76287061 | 0.63617039 | 4.34297263 | 1.405676498 | 0.00029774 |
| LAT | 114.042102 | -4.2547897 | 0.98011753 | -4.3411015 | 1.417701800 | 0.00029983 |
| GIMAP5 | 137.632631 | 1.86228378 | 0.429357 | 4.33737838 | 1.441922762 | 0.0003045 |
| CAVIN3 | 33.8440593 | -5.5174636 | 1.27347779 | -4.3325951 | 1.473619684 | 0.00031072 |
| ENSG000002 | 257.019143 | -1.5351386 | 0.35441497 | -4.3314722 | 1.481156289 | 0.00031184 |
| ID3 | 929.927355 | 3.73350663 | 0.86343374 | 4.32402214 | 1.532099072 | 0.00032208 |
| CCDC26 | 15.4524177 | 7.13488823 | 1.65061516 | 4.32256312 | 1.542269420 | 0.00032374 |
| TDRD3 | 403.272999 | -3.7728526 | 0.87344731 | -4.3194965 | 1.563855593 | 0.00032778 |
| PTGES3L | 30.0899445 | -4.6529647 | 1.07877739 | -4.3131834 | 1.609205206 | 0.00033678 |

| | | | | | | |
|---|---|---|---|---|---|---|
| ITM2B | 16574.6174 | 1.75054177 | 0.40638689 | 4.30757439 | 1.650545931 | 0.00034481 |
| B4GALT5 | 937.443709 | -1.2545991 | 0.29127155 | -4.3073177 | 1.652461625 | 0.00034481 |
| ZNF780A | 164.527389 | 1.35261713 | 0.31413796 | 4.305806 | 1.663788186 | 0.00034666 |
| COLGALT2 | 152.546447 | 3.39075704 | 0.78782941 | 4.30392289 | 1.678000840 | 0.0003491 |
| FLI1 | 848.452557 | 0.8503568 | 0.19770389 | 4.30116374 | 1.699034533 | 0.00035295 |
| CRACDL | 32.3701544 | 4.44134514 | 1.03304481 | 4.29927637 | 1.713566832 | 0.00035544 |
| ADTRP | 42.1431379 | 2.44077955 | 0.56781551 | 4.29854332 | 1.719243027 | 0.0003561 |
| RPS6KA2 | 157.900053 | -2.8100164 | 0.65400198 | -4.2966482 | 1.734000643 | 0.00035862 |
| GRN | 32161.9315 | 0.9341676 | 0.21746499 | 4.29571496 | 1.741312110 | 0.00035961 |
| NLRX1 | 175.827721 | -1.7955176 | 0.41827934 | -4.292628 | 1.765707024 | 0.00036411 |
| LSR | 46.2540976 | -5.0156512 | 1.16940968 | -4.2890454 | 1.794426638 | 0.00036949 |
| MRO | 59.0012265 | 2.49160986 | 0.58098966 | 4.28856144 | 1.798340564 | 0.00036975 |
| IGSF3 | 54.3905254 | 3.25377992 | 0.75896937 | 4.28710308 | 1.810183109 | 0.00037164 |
| ABLIM1 | 64.4390027 | -3.6509274 | 0.85210418 | -4.2846021 | 1.830665361 | 0.0003753 |
| ENSG000002 | 47.9201453 | 2.46128283 | 0.57468712 | 4.2828223 | 1.845375487 | 0.00037776 |
| TCF12 | 827.573027 | 0.85841383 | 0.20053645 | 4.28058746 | 1.864006097 | 0.00038067 |
| S100B | 2958.2254 | 2.92270884 | 0.68281011 | 4.28041238 | 1.865473231 | 0.00038067 |
| MGLL | 1309.19919 | -1.8865361 | 0.44076447 | -4.2801457 | 1.867710271 | 0.00038067 |
| RBFOX2 | 93.5499066 | -1.5189424 | 0.35501704 | -4.2785057 | 1.881521993 | 0.00038293 |
| CCR5 | 561.693139 | 1.9397625 | 0.45363767 | 4.27601727 | 1.902664676 | 0.00038667 |
| TAP1 | 251.855314 | -2.0058578 | 0.46923163 | -4.274771 | 1.913338677 | 0.00038828 |
| PRKAR2B | 32.1962694 | 3.6512954 | 0.85466884 | 4.27217562 | 1.935750285 | 0.00039226 |
| SLC4A7 | 1151.71267 | 1.89235251 | 0.44306976 | 4.27100357 | 1.945952933 | 0.00039376 |
| UBE2Q2 | 656.027097 | 1.01904769 | 0.2388207 | 4.26699896 | 1.981200807 | 0.00040032 |
| TNNC1 | 14.9654972 | 7.08944856 | 1.66284565 | 4.26344355 | 2.013003681 | 0.00040616 |
| SPON2 | 193.562878 | -11.297415 | 2.65050918 | -4.2623566 | 2.022823175 | 0.00040755 |
| CYRIB | 2745.37195 | 1.1101323 | 0.26050196 | 4.26151224 | 2.030482314 | 0.00040839 |
| PFKP | 322.052226 | -1.9988559 | 0.46907648 | -4.261258 | 2.032793676 | 0.00040839 |
| PRKG1 | 30.3285227 | -5.7594347 | 1.35188407 | -4.2603022 | 2.041507032 | 0.00040956 |
| TENT5A | 293.36814 | 1.2680924 | 0.29776243 | 4.25873878 | 2.055835365 | 0.00041184 |
| STAT4 | 199.789702 | 4.25049937 | 0.99939552 | 4.25307026 | 2.108593816 | 0.00042181 |
| NUPR1 | 17.0631968 | 7.28096927 | 1.71242983 | 4.25183511 | 2.120259480 | 0.00042354 |
| MYBPH | 96.9081185 | -3.774073 | 0.88799324 | -4.2501146 | 2.136611540 | 0.0004262 |
| CTTN | 76.0917872 | -4.1336349 | 0.9734165 | -4.2465224 | 2.171141301 | 0.00043247 |
| DOP1B | 946.186072 | -2.6424611 | 0.62256359 | -4.2444838 | 2.190971699 | 0.0004358 |
| MTCO1P12 | 486.254321 | -1.1944692 | 0.28147463 | -4.2436122 | 2.199502929 | 0.00043688 |
| MOB3A | 2341.10103 | 1.23658153 | 0.291572 | 4.24108459 | 2.224422733 | 0.00044116 |
| PIM1 | 360.979356 | -2.3863314 | 0.56270863 | -4.240794 | 2.227304996 | 0.00044116 |
| OTOA | 52.4191794 | -2.6375868 | 0.6221278 | -4.2396221 | 2.238963643 | 0.00044284 |
| GAPT | 46.1318976 | 4.07889536 | 0.96302579 | 4.23549962 | 2.280440910 | 0.00045041 |
| AKNA | 1076.33462 | 1.46784969 | 0.3470901 | 4.22901625 | 2.347153927 | 0.00046294 |
| IFI27 | 17.4280795 | -4.8799722 | 1.15512357 | -4.2246322 | 2.393313786 | 0.00047138 |
| HYI | 83.3671636 | -1.685577 | 0.39908708 | -4.2235819 | 2.404499013 | 0.00047292 |
| CHORDC1 | 381.395381 | 1.10175162 | 0.26093687 | 4.22229182 | 2.418307307 | 0.00047497 |
| IL7R | 5871.23494 | 3.8868515 | 0.92100586 | 4.22022451 | 2.440591376 | 0.00047868 |
| CASTOR1 | 138.313216 | 2.44251754 | 0.57884152 | 4.21966539 | 2.446651749 | 0.0004792 |
| LACC1 | 1953.93734 | 2.3830409 | 0.56482461 | 4.21908118 | 2.452999428 | 0.00047977 |
| ENSG000002 | 22.0206605 | 3.81004237 | 0.90332946 | 4.21777717 | 2.467224514 | 0.0004813 |

| | | | | | | |
|---|---|---|---|---|---|---|
| USP2-AS1 | 20.6031111 | -7.0891329 | 1.68079148 | -4.2177349 | 2.467686911 | 0.0004813 |
| DEF6 | 401.140391 | 1.45570717 | 0.34535493 | 4.21510464 | 2.496624061 | 0.00048576 |
| NINL | 35.0742246 | -3.4298699 | 0.81372418 | -4.2150276 | 2.497476697 | 0.00048576 |
| TENM4 | 26.6256125 | 3.58818461 | 0.85142604 | 4.21432333 | 2.505281731 | 0.00048661 |
| RASGRP2 | 23.3364324 | 5.26731662 | 1.25056368 | 4.21195395 | 2.531711615 | 0.00049106 |
| MAN2B1 | 11986.1528 | 1.15970121 | 0.27539513 | 4.21104471 | 2.541924283 | 0.00049236 |
| SH3RF3 | 109.774635 | -2.5214881 | 0.59894567 | -4.2098779 | 2.555087394 | 0.00049423 |
| PHC2 | 2453.29207 | 1.23062965 | 0.2923648 | 4.20922651 | 2.562464014 | 0.00049497 |
| TIMP1 | 2546.50983 | 1.20974019 | 0.28759183 | 4.20644842 | 2.594152588 | 0.00050041 |
| PHF12 | 315.2555 | 1.06630668 | 0.25359483 | 4.20476507 | 2.613534773 | 0.00050345 |
| CAD | 357.904454 | -1.5501223 | 0.36872098 | -4.2040522 | 2.621783741 | 0.00050435 |
| MGAT4B | 1111.94919 | -1.8948843 | 0.4508491 | -4.2029236 | 2.634895542 | 0.00050618 |
| SYNDIG1 | 19.0034671 | 7.43286492 | 1.76906179 | 4.20158582 | 2.650517290 | 0.00050849 |
| ZDHHC9 | 216.823005 | -1.3914626 | 0.33134529 | -4.1994337 | 2.675833710 | 0.00051264 |
| TMED8 | 342.828079 | -1.1301418 | 0.2691968 | -4.1981992 | 2.690459663 | 0.00051419 |
| PREX1 | 1847.25086 | 1.67459612 | 0.39889039 | 4.19813606 | 2.691209267 | 0.00051419 |
| CBLB | 435.004378 | -1.7626185 | 0.42010634 | -4.1956485 | 2.720919800 | 0.00051916 |
| ACOT4 | 39.013895 | -4.1034469 | 0.97855226 | -4.1933856 | 2.748217349 | 0.00052365 |
| NRG1 | 15.34042 | 7.1252154 | 1.69977491 | 4.1918582 | 2.766789169 | 0.00052648 |
| GPER1 | 31.7178077 | -7.7148833 | 1.84076583 | -4.191127 | 2.775722488 | 0.00052676 |
| TJP1 | 171.445637 | 1.97304214 | 0.47076697 | 4.19112274 | 2.775774361 | 0.00052676 |
| SLC39A13-AS | 399.722213 | 1.39077832 | 0.33203942 | 4.18859397 | 2.806880816 | 0.00053194 |
| TMEM106B | 205.109531 | -1.4351637 | 0.34278288 | -4.1868011 | 2.829135839 | 0.00053544 |
| PSTPIP2 | 679.400293 | 1.9612047 | 0.46856761 | 4.18553194 | 2.844990604 | 0.00053771 |
| DLEU1 | 67.7578994 | 2.10738012 | 0.50386598 | 4.18242196 | 2.884200366 | 0.00054439 |
| SPSB1 | 592.876426 | -2.8013163 | 0.67010067 | -4.180441 | 2.909443158 | 0.00054842 |
| SMG9 | 465.843032 | 1.02548031 | 0.24534365 | 4.17977121 | 2.918025178 | 0.0005493 |
| JAM3 | 18.9715114 | -4.1329181 | 0.99003808 | -4.1745041 | 2.986357826 | 0.00056141 |
| MTURN | 279.325407 | 1.48898436 | 0.35675647 | 4.17367157 | 2.997297515 | 0.00056271 |
| SERPINA1 | 1070.52111 | 3.20531405 | 0.76861076 | 4.17026954 | 3.042395810 | 0.00057042 |
| HIVEP1 | 543.130731 | 1.76744928 | 0.42399771 | 4.16853493 | 3.065637896 | 0.00057343 |
| HLA-DOA | 40.2185886 | 8.51531109 | 2.04282067 | 4.16840853 | 3.067338050 | 0.00057343 |
| TUSC3 | 20.2343853 | -5.5663062 | 1.33543486 | -4.16816 | 3.070683030 | 0.00057343 |
| GNA15 | 1292.17038 | 1.33475087 | 0.32040661 | 4.16580315 | 3.102583393 | 0.00057861 |
| TRANK1 | 91.3571339 | -1.8665105 | 0.4486075 | -4.1606761 | 3.173068201 | 0.00059097 |
| FADS2 | 2589.48114 | -1.2329476 | 0.29679327 | -4.1542304 | 3.263842258 | 0.00060707 |
| LTC4S | 262.393178 | 3.7660005 | 0.90695817 | 4.15234201 | 3.290899485 | 0.0006113 |
| IL2RG | 277.429211 | -2.6611259 | 0.64092622 | -4.1520003 | 3.295818257 | 0.0006114 |
| ELMO1 | 964.1158 | 1.39298978 | 0.33568596 | 4.14968143 | 3.329383047 | 0.00061681 |
| PSEN2 | 685.804167 | -2.8747361 | 0.69295713 | -4.1485049 | 3.346536297 | 0.00061918 |
| FBXO38 | 658.004134 | -1.6632886 | 0.40100871 | -4.1477617 | 3.357415644 | 0.00062037 |
| C14orf132 | 48.6267892 | -4.8430212 | 1.1689926 | -4.1429015 | 3.429392722 | 0.00063234 |
| PPL | 25.8462946 | -6.4618524 | 1.55989664 | -4.1424875 | 3.435590706 | 0.00063234 |
| GBP1 | 17.3876391 | -6.8396776 | 1.6511067 | -4.1424807 | 3.435692675 | 0.00063234 |
| CD300E | 41.5205744 | 2.84293914 | 0.68680373 | 4.13937639 | 3.482511845 | 0.00064012 |
| SLC45A3 | 165.220798 | -1.6477677 | 0.39823159 | -4.1377122 | 3.507860167 | 0.00064393 |
| CLEC11A | 1356.7728 | 1.70077009 | 0.41116741 | 4.13644184 | 3.527326977 | 0.00064666 |
| C3orf62 | 69.0793546 | 2.55891781 | 0.61893811 | 4.13436785 | 3.559329944 | 0.00065168 |

| | | | | | | |
|---|---|---|---|---|---|---|
| LINC01010 | 259.271379 | -7.4387628 | 1.8000339 | -4.1325682 | 3.587323184 | 0.00065595 |
| DMRTA1 | 12.9352063 | -6.4084637 | 1.55140532 | -4.1307475 | 3.615855270 | 0.00065975 |
| ALDH9A1 | 538.581343 | -1.0345972 | 0.25046878 | -4.1306434 | 3.617494263 | 0.00065975 |
| MIPEP | 124.59249 | -2.0888988 | 0.50604561 | -4.1278863 | 3.661130248 | 0.00066684 |
| SOX13 | 26.4901251 | -3.959411 | 0.95939612 | -4.1269825 | 3.675543267 | 0.00066798 |
| TMEM255A | 110.330239 | -1.9887501 | 0.48189954 | -4.1268976 | 3.676900336 | 0.00066798 |
| CDH23 | 197.984276 | -8.0722853 | 1.9582869 | -4.1221158 | 3.754084568 | 0.00068112 |
| EGR2 | 412.496259 | -3.3132926 | 0.80415665 | -4.1202079 | 3.785306179 | 0.00068544 |
| ARF4 | 2221.99311 | 0.95510448 | 0.23181789 | 4.12006376 | 3.787675767 | 0.00068544 |
| LDAH | 228.656429 | -1.1751343 | 0.28541026 | -4.1173512 | 3.832518284 | 0.00069266 |
| GATB | 186.567174 | -1.5281512 | 0.37144926 | -4.1140242 | 3.888207265 | 0.00070096 |
| KLK4 | 81.5839885 | -4.1882043 | 1.01809207 | -4.1137775 | 3.892365857 | 0.00070096 |
| KCNC3 | 183.129943 | 1.38219797 | 0.33599734 | 4.113717 | 3.893387078 | 0.00070096 |
| G0S2 | 223.794812 | 1.95554199 | 0.47561865 | 4.11157549 | 3.929683397 | 0.00070659 |
| NFATC3 | 1223.65709 | 1.2855896 | 0.31272468 | 4.110931 | 3.940669510 | 0.00070766 |
| PILRA | 1892.09074 | 2.01425543 | 0.49013201 | 4.10961817 | 3.963138737 | 0.00071078 |
| LIPN | 18.6456983 | 7.40419331 | 1.80239252 | 4.10798049 | 3.991338126 | 0.00071493 |
| OLFML2B | 456.291332 | 2.35784355 | 0.57418088 | 4.10644732 | 4.017910442 | 0.00071822 |
| CASP1 | 652.664159 | 2.0052548 | 0.48833247 | 4.10633106 | 4.019932310 | 0.00071822 |
| SEMA6B | 1803.12724 | 2.47391638 | 0.60284548 | 4.10373213 | 4.065381444 | 0.00072541 |
| ENOX1 | 26.9171836 | 4.16212928 | 1.01453188 | 4.10251205 | 4.086885619 | 0.00072807 |
| ADAP1 | 732.958264 | 1.64615929 | 0.40127708 | 4.10230081 | 4.090619644 | 0.00072807 |
| MAP1LC3A | 125.293318 | 1.57820446 | 0.38477589 | 4.10161994 | 4.102677439 | 0.00072929 |
| ARHGAP30 | 1795.50965 | 0.97971434 | 0.23902424 | 4.09880747 | 4.152842928 | 0.00073727 |
| ARMT1 | 524.656948 | 1.02844859 | 0.2509589 | 4.09807585 | 4.165987833 | 0.00073833 |
| PLEKHO1 | 1606.48195 | -1.3327356 | 0.32522469 | -4.097892 | 4.169298056 | 0.00073833 |
| ADORA1 | 32.2704936 | 3.31805001 | 0.81027027 | 4.09499166 | 4.221835917 | 0.00074669 |
| GRAMD4 | 494.388027 | 1.450829 | 0.35432777 | 4.0945958 | 4.229055167 | 0.00074703 |
| ISYNA1 | 320.230987 | -2.1340832 | 0.52129717 | -4.0937939 | 4.243716245 | 0.00074801 |
| A4GALT | 137.810329 | 3.1738348 | 0.77534567 | 4.09344492 | 4.250110755 | 0.00074801 |
| TMCC3 | 317.999903 | 2.86990386 | 0.70110183 | 4.09341942 | 4.250578426 | 0.00074801 |
| CH25H | 13.492298 | -6.4696733 | 1.5807942 | -4.0926727 | 4.264293462 | 0.00074948 |
| RTN4RL2 | 93.7364945 | 1.8894575 | 0.4623344 | 4.0867768 | 4.374075448 | 0.00076781 |
| WAPL | 985.813007 | 0.97352849 | 0.23836032 | 4.0842725 | 4.421512012 | 0.00077517 |
| TLR2 | 2568.13661 | 2.36826663 | 0.58030819 | 4.08104982 | 4.483274237 | 0.00078502 |
| GSTM3 | 113.96672 | 3.0906126 | 0.75763648 | 4.07928167 | 4.517507394 | 0.0007899 |
| CCNA1 | 60.0428593 | -4.7923296 | 1.17487804 | -4.0790018 | 4.522949630 | 0.0007899 |
| AGAP2 | 26.4209479 | 5.47718217 | 1.34286086 | 4.07874139 | 4.528017170 | 0.0007899 |
| MT-ATP8 | 6877.47746 | -1.0018331 | 0.24577356 | -4.0762443 | 4.576893368 | 0.00079744 |
| NT5C2 | 1114.66451 | 1.02598369 | 0.25173406 | 4.07566492 | 4.588304918 | 0.00079759 |
| GIMAP6 | 483.657304 | 1.79552642 | 0.44055267 | 4.07562256 | 4.589140440 | 0.00079759 |
| DNAJC5 | 2913.04981 | 0.74215827 | 0.18212066 | 4.07509098 | 4.599636150 | 0.00079843 |
| FGFRL1 | 827.192394 | 1.68824382 | 0.41457799 | 4.07219838 | 4.657148793 | 0.00080741 |
| AMT | 40.0922089 | 2.315383 | 0.56871715 | 4.07123824 | 4.676389177 | 0.00080975 |
| EGFL7 | 575.514383 | -2.7312869 | 0.67106716 | -4.0700649 | 4.700004490 | 0.00081283 |
| THRB | 53.1551361 | -5.6302179 | 1.3834803 | -4.0696047 | 4.709298145 | 0.00081315 |
| DAGLB | 466.088055 | 1.08749432 | 0.26723695 | 4.0694011 | 4.713414289 | 0.00081315 |
| SCUBE1 | 22.6130784 | -8.1989768 | 2.01724478 | -4.0644432 | 4.814725825 | 0.00082961 |

| | | | | | | |
|---|---|---|---|---|---|---|
| PCDH1 | 58.0006104 | -3.7410156 | 0.92088922 | -4.0623948 | 4.857183952 | 0.0008359 |
| RUFY3 | 129.41708 | -1.8822828 | 0.46346758 | -4.061304 | 4.879937152 | 0.00083879 |
| MMS19 | 452.475078 | -1.1620035 | 0.28618837 | -4.0602751 | 4.901493849 | 0.00084146 |
| CBR4 | 81.2555345 | -1.6575362 | 0.40829693 | -4.0596341 | 4.914967365 | 0.00084275 |
| CALU | 3063.72382 | -1.591203 | 0.39199552 | -4.0592378 | 4.923316196 | 0.00084315 |
| TMEM140 | 48.0875885 | -3.6782071 | 0.90634008 | -4.0583079 | 4.942956073 | 0.00084548 |
| ANO5 | 64.965296 | -4.7854604 | 1.17933877 | -4.0577487 | 4.954803103 | 0.00084648 |
| ENSG000002 | 14.6790616 | 7.06358308 | 1.74093838 | 4.05734237 | 4.963429437 | 0.00084692 |
| SLC6A7 | 22.0655449 | -5.7229709 | 1.41347364 | -4.0488699 | 5.146553690 | 0.00087661 |
| ZNF652 | 421.210584 | 1.92807671 | 0.476257 | 4.04839553 | 5.156994269 | 0.00087661 |
| CKB | 112.023225 | 1.82028587 | 0.44965447 | 4.04818812 | 5.161565440 | 0.00087661 |
| NR4A1 | 132.311691 | 2.59974494 | 0.64220579 | 4.04814933 | 5.162420601 | 0.00087661 |
| MAFF | 644.843517 | -1.7720985 | 0.43780484 | -4.0476904 | 5.172549442 | 0.00087727 |
| AK3 | 415.644008 | -1.5860399 | 0.39196091 | -4.0464237 | 5.200606382 | 0.00088097 |
| CD226 | 115.551598 | 3.12881435 | 0.77345273 | 4.04525607 | 5.226596279 | 0.0008843 |
| CNR2 | 16.2032257 | 7.20717027 | 1.78260286 | 4.04305996 | 5.275812091 | 0.00089097 |
| SMARCA1 | 86.8776223 | -2.2337239 | 0.55250104 | -4.0429315 | 5.278703722 | 0.00089097 |
| PNRC2 | 790.485178 | 1.10858513 | 0.27456841 | 4.03755522 | 5.401113485 | 0.00091054 |
| MAF | 2775.72656 | 1.36757987 | 0.33909583 | 4.03301882 | 5.506486813 | 0.00092719 |
| SEL1L3 | 111.002317 | 2.88062975 | 0.71494844 | 4.02914336 | 5.598047352 | 0.00094148 |
| MYCL | 69.8196025 | 2.07750452 | 0.51622248 | 4.02443634 | 5.711193937 | 0.00095936 |
| NFKBIE | 1315.89624 | 1.00361621 | 0.24963564 | 4.02032429 | 5.811808426 | 0.0009746 |
| LY86 | 872.185204 | 1.63726634 | 0.4072637 | 4.0201627 | 5.815796294 | 0.0009746 |
| RCSD1 | 704.55357 | 1.50736619 | 0.37522153 | 4.01727001 | 5.887624984 | 0.00098546 |
| CAMSAP2 | 545.231154 | 1.26155692 | 0.3143783 | 4.01286255 | 5.998684612 | 0.00100285 |
| STX2 | 207.444623 | -2.4740315 | 0.61730612 | -4.0077871 | 6.129032112 | 0.00102328 |
| SRGN | 11548.6289 | 3.29106154 | 0.82121739 | 4.00754001 | 6.135447023 | 0.00102328 |
| CYB5A | 97.4724251 | -2.7664809 | 0.69107005 | -4.0031845 | 6.249553711 | 0.00104107 |
| FCGR2C | 43.5167809 | 7.16469357 | 1.79145203 | 3.99937786 | 6.350921322 | 0.00105671 |
| BBS2 | 84.9264274 | -1.4744427 | 0.36914581 | -3.9942014 | 6.491266322 | 0.00107878 |
| TSPAN15 | 1249.58101 | 1.6110781 | 0.40354617 | 3.99230179 | 6.543502272 | 0.00108617 |
| TTYH2 | 213.121585 | -2.6737293 | 0.66994228 | -3.9909846 | 6.579956022 | 0.00109094 |
| ARHGAP5-AS1 | 45.9433205 | -3.75935 | 0.94343122 | -3.9847632 | 6.754746269 | 0.0011186 |
| ENSG000002 | 32.6942631 | 8.21584302 | 2.0621111 | 3.98419029 | 6.771059913 | 0.00111998 |
| TANC1 | 56.6237963 | 2.00390606 | 0.50307039 | 3.98335121 | 6.795022009 | 0.00112262 |
| HLA-DQA1 | 287.385952 | 6.81994215 | 1.71257919 | 3.98226381 | 6.826194774 | 0.00112644 |
| EPHB3 | 290.413666 | 3.18093437 | 0.800069 | 3.97582506 | 7.013567057 | 0.001156 |
| TSHZ1 | 78.2635455 | -2.1408998 | 0.53858467 | -3.9750479 | 7.036509972 | 0.00115843 |
| TRPV2 | 1897.2117 | -2.8122532 | 0.70758084 | -3.974462 | 7.053853323 | 0.00115992 |
| MYO19 | 125.769736 | -1.5391938 | 0.38742644 | -3.9728672 | 7.101263067 | 0.00116635 |
| RAB43 | 366.165367 | 1.13864254 | 0.28669306 | 3.97164323 | 7.137856437 | 0.00117099 |
| FBXO17 | 13.6943191 | -6.4963951 | 1.63617639 | -3.9704736 | 7.172991304 | 0.00117538 |
| SH3BP4 | 22.8172005 | 5.26046957 | 1.32514164 | 3.96974134 | 7.195069576 | 0.00117763 |
| TRIM36 | 238.082357 | 1.40329782 | 0.35378452 | 3.96653253 | 7.292582517 | 0.0011922 |
| BMPR2 | 1220.90092 | 1.06618341 | 0.26903908 | 3.96293137 | 7.403507050 | 0.00120892 |
| ENSG000002 | 10.3333479 | -7.0711488 | 1.78535495 | -3.9606403 | 7.474905989 | 0.00121916 |
| SLPI | 18.5434954 | 7.39851837 | 1.87092827 | 3.95446394 | 7.670643816 | 0.00124964 |
| CD6 | 73.0913901 | -2.5481074 | 0.64445601 | -3.9538888 | 7.689116510 | 0.0012512 |

| | | | | | | |
|---|---|---|---|---|---|---|
| PRDX3 | 1533.59507 | -0.7512746 | 0.19005892 | -3.9528509 | 7.722556376 | 0.00125518 |
| ACAT2 | 841.345821 | -2.1966561 | 0.55578397 | -3.952356 | 7.738550653 | 0.00125633 |
| DNAJC3 | 1992.68419 | 1.05325521 | 0.26654335 | 3.95153438 | 7.765171318 | 0.0012592 |
| DTX1 | 84.5561307 | -1.9337454 | 0.48956098 | -3.9499582 | 7.816484749 | 0.00126517 |
| SPCS3 | 2512.72904 | 0.95784422 | 0.24250144 | 3.94984955 | 7.820033123 | 0.00126517 |
| IL16 | 211.612016 | 1.56224936 | 0.3956589 | 3.94847521 | 7.865055883 | 0.00127099 |
| IGSF21 | 20.7956492 | 4.06410074 | 1.02938787 | 3.94807523 | 7.878205144 | 0.00127165 |
| NF1 | 692.697656 | 0.9315914 | 0.23602279 | 3.94704003 | 7.912333284 | 0.00127498 |
| ENSG000002 | 15.910888 | -4.7678044 | 1.20798749 | -3.9468988 | 7.917000134 | 0.00127498 |
| TCTN2 | 69.804381 | -2.9172872 | 0.73934666 | -3.9457636 | 7.954605739 | 0.00127957 |
| GPC3 | 51.403836 | -5.8903818 | 1.49298013 | -3.9453853 | 7.967177011 | 0.00128013 |
| SULT1B1 | 16.8886902 | 7.26362471 | 1.84321115 | 3.94074478 | 8.122901313 | 0.00130366 |
| EML2 | 211.349821 | 1.22839252 | 0.31201984 | 3.93690519 | 8.253919868 | 0.00132247 |
| BAALC | 12.203824 | -7.3086033 | 1.85650451 | -3.9367549 | 8.259088012 | 0.00132247 |
| ABCB10 | 180.058322 | -1.6441078 | 0.41765894 | -3.9364842 | 8.268405291 | 0.00132247 |
| CLUH | 817.597477 | -1.2498493 | 0.31765994 | -3.9345512 | 8.335230959 | 0.00133165 |
| KCNN4 | 200.156064 | -3.1616718 | 0.80405014 | -3.9321824 | 8.417814657 | 0.00134331 |
| B3GALNT1 | 233.077173 | 1.23063829 | 0.31299694 | 3.93179014 | 8.431565690 | 0.00134398 |
| ANKS6 | 71.5145948 | -3.2135568 | 0.81767153 | -3.9301317 | 8.489936330 | 0.00135175 |
| PTAFR | 3793.30131 | 2.06557354 | 0.52568217 | 3.92932011 | 8.518637515 | 0.00135478 |
| CCR2 | 17.1252623 | 7.28359056 | 1.85486934 | 3.92674051 | 8.610478984 | 0.00136784 |
| SPARC | 4266.99467 | 2.81102349 | 0.71605873 | 3.92568846 | 8.648202721 | 0.00137228 |
| VASN | 14.6165878 | -5.0632804 | 1.29096346 | -3.9220943 | 8.778263004 | 0.00139134 |
| DCSTAMP | 1447.82822 | -3.7363185 | 0.95323792 | -3.9196075 | 8.869327934 | 0.00140379 |
| TSPAN14 | 1611.92722 | 1.1144238 | 0.28433496 | 3.91940472 | 8.876794586 | 0.00140379 |
| ASCL2 | 25.5591005 | 3.56740596 | 0.91056533 | 3.91779244 | 8.936359877 | 0.00141162 |
| ANKRD28 | 343.898167 | -1.5223018 | 0.38869863 | -3.9164064 | 8.987867141 | 0.00141816 |
| HSPA6 | 30.1468547 | 3.10217649 | 0.7922724 | 3.91554279 | 9.020104550 | 0.00142165 |
| ARHGAP35 | 941.315841 | 1.2216704 | 0.3121487 | 3.91374495 | 9.087562245 | 0.00143068 |
| PDGFC | 178.133694 | -2.2564601 | 0.57661073 | -3.9133161 | 9.103722568 | 0.00143162 |
| TRUB2 | 199.928358 | -1.3517602 | 0.34598441 | -3.9069975 | 9.345012805 | 0.00146792 |
| G2E3 | 143.58054 | 1.7054661 | 0.4367297 | 3.90508388 | 9.419270829 | 0.00147793 |
| NMRK2 | 229.620144 | -8.619186 | 2.20855876 | -3.9026292 | 9.515340148 | 0.00149134 |
| JAK3 | 106.584247 | -2.7425993 | 0.70363334 | -3.8977677 | 9.708346085 | 0.00151981 |
| CYP20A1 | 235.322877 | 1.18359381 | 0.30367936 | 3.89751156 | 9.718618398 | 0.00151981 |
| CBFB | 1472.02245 | 1.33202021 | 0.34184701 | 3.89653906 | 9.757707627 | 0.00152422 |
| UBA6 | 771.286141 | 1.41785237 | 0.3640248 | 3.89493344 | 9.822569716 | 0.00153265 |
| GPR35 | 1323.23548 | 1.97178122 | 0.50667471 | 3.8916117 | 9.958052671 | 0.00155206 |
| SNX18 | 948.130886 | 1.21415999 | 0.31206994 | 3.89066622 | 9.996937012 | 0.0015564 |
| SELENOM | 50.1881718 | 2.93378605 | 0.75460638 | 3.88783626 | 0.00010114 | 0.00157278 |
| PLXNB2 | 2322.80538 | -0.981805 | 0.25254869 | -3.8875869 | 0.00010125 | 0.00157278 |
| PPM1E | 15.1531408 | -6.6384772 | 1.70842539 | -3.8857285 | 0.00010202 | 0.00158311 |
| TJP2 | 169.263498 | 2.17431282 | 0.55966798 | 3.88500486 | 0.00010233 | 0.00158399 |
| RCAN1 | 590.465301 | -1.1197342 | 0.28822624 | -3.8849142 | 0.00010237 | 0.00158399 |
| CPAMD8 | 19.3438202 | -3.057075 | 0.78693435 | -3.8847904 | 0.00010242 | 0.00158399 |
| HS6ST1 | 232.009921 | -1.878147 | 0.48359499 | -3.8837189 | 0.00010287 | 0.00158924 |
| MMP19 | 1193.45807 | -2.3085822 | 0.59449797 | -3.8832465 | 0.00010307 | 0.00159058 |
| SDC1 | 216.555872 | -3.7172743 | 0.95751538 | -3.8822084 | 0.00010351 | 0.00159563 |

| | | | | | | |
|---|---|---|---|---|---|---|
| ADM | 136.797002 | 2.58133622 | 0.66502758 | 3.88154763 | 0.00010379 | 0.00159822 |
| ATXN1 | 748.078738 | 1.37744811 | 0.35494313 | 3.88075718 | 0.00010413 | 0.00160167 |
| TLE4 | 522.817027 | 1.87472088 | 0.48340248 | 3.87817802 | 0.00010524 | 0.00161591 |
| NXPE3 | 233.577022 | 1.00476901 | 0.25908989 | 3.87807104 | 0.00010529 | 0.00161591 |
| TNF | 256.479187 | 2.89421221 | 0.746421 | 3.8774528 | 0.00010556 | 0.00161825 |
| IGF2BP3 | 915.643841 | -1.0548228 | 0.27222824 | -3.8747736 | 0.00010672 | 0.00163438 |
| FNIP2 | 1392.75227 | -2.3347069 | 0.60270885 | -3.8736894 | 0.0001072 | 0.00163988 |
| SCARB1 | 1305.87229 | -1.08144 | 0.27923514 | -3.8728649 | 0.00010756 | 0.00164365 |
| KLHDC2 | 83.6094005 | -1.4970683 | 0.38694147 | -3.8689787 | 0.00010929 | 0.00166825 |
| COL1A2 | 163.695524 | -3.2833237 | 0.84902957 | -3.8671488 | 0.00011012 | 0.00167899 |
| ARHGEF9 | 144.353428 | -1.1568502 | 0.29917222 | -3.8668369 | 0.00011026 | 0.00167931 |
| KCNQ1 | 312.539111 | 2.16283265 | 0.55944448 | 3.86603627 | 0.00011062 | 0.00168301 |
| CKLF | 661.886424 | 1.13820214 | 0.29457245 | 3.86391232 | 0.00011159 | 0.00169543 |
| RNF166 | 440.27672 | 1.11822576 | 0.28941742 | 3.86371272 | 0.00011168 | 0.00169543 |
| CD1D | 92.2078416 | 1.89087046 | 0.49021779 | 3.85720486 | 0.00011469 | 0.00173787 |
| TYROBP | 8248.59369 | 0.80771059 | 0.20940633 | 3.85714495 | 0.00011472 | 0.00173787 |
| TBXAS1 | 1811.83672 | 1.66924667 | 0.43293523 | 3.85564985 | 0.00011542 | 0.00174664 |
| SERPINF1 | 2900.06242 | 5.08872764 | 1.32025403 | 3.85435493 | 0.00011604 | 0.00175402 |
| IPO13 | 412.900413 | -0.9691311 | 0.25149981 | -3.8534071 | 0.00011649 | 0.00175893 |
| ASAP2 | 35.8380504 | -3.4658306 | 0.89992617 | -3.8512388 | 0.00011752 | 0.00177267 |
| TRPM2 | 897.371836 | 1.70994408 | 0.44425596 | 3.84900654 | 0.0001186 | 0.00178698 |
| ENSG000002 | 57.3643189 | 4.99139502 | 1.29791664 | 3.84569769 | 0.00012021 | 0.00180933 |
| BTAF1 | 343.341815 | 0.96158766 | 0.25036633 | 3.84072282 | 0.00012267 | 0.00184442 |
| CPEB4 | 887.503902 | 0.76131778 | 0.19824803 | 3.84022878 | 0.00012292 | 0.00184616 |
| ACVR2A | 198.512456 | -1.9507113 | 0.50809708 | -3.8392492 | 0.00012341 | 0.00185156 |
| LINC00649 | 52.9999262 | -2.0554354 | 0.53621314 | -3.8332433 | 0.00012646 | 0.00189392 |
| MAEA | 732.189655 | 0.93304445 | 0.24341347 | 3.83316682 | 0.0001265 | 0.00189392 |
| C1orf226 | 27.5435159 | -4.9908083 | 1.30257252 | -3.8315013 | 0.00012736 | 0.00190476 |
| TACC2 | 237.749792 | -1.9078436 | 0.49801713 | -3.8308795 | 0.00012769 | 0.00190755 |
| SMAGP | 92.2208505 | 1.67997733 | 0.43857848 | 3.83050559 | 0.00012788 | 0.00190842 |
| CSF2RB | 680.323774 | 1.1741577 | 0.30683606 | 3.82666131 | 0.00012989 | 0.00193446 |
| FBXW8 | 107.075922 | -1.916519 | 0.50083511 | -3.8266467 | 0.0001299 | 0.00193446 |
| ZNF687 | 408.730715 | 1.12101314 | 0.2931784 | 3.82365531 | 0.00013149 | 0.00195602 |
| FBLIM1 | 31.9571209 | -3.2631531 | 0.85373522 | -3.8222075 | 0.00013226 | 0.00196546 |
| ZCCHC24 | 228.401253 | 1.2343852 | 0.32307318 | 3.82076034 | 0.00013304 | 0.00197494 |
| RETN | 719.670252 | 6.99318311 | 1.83053835 | 3.82028767 | 0.0001333 | 0.00197576 |
| FUT4 | 210.853657 | -1.2447415 | 0.32583683 | -3.8201376 | 0.00013338 | 0.00197576 |
| TMEM134 | 159.014598 | -1.3369979 | 0.35005803 | -3.8193609 | 0.0001338 | 0.0019799 |
| SERINC5 | 1100.52712 | 1.51694713 | 0.39747033 | 3.81650408 | 0.00013536 | 0.00200084 |
| HR | 40.3721691 | 3.55330244 | 0.932436 | 3.81077354 | 0.00013853 | 0.00204135 |
| OAS1 | 248.882369 | -1.7929381 | 0.47051977 | -3.8105479 | 0.00013866 | 0.00204135 |
| TFCP2 | 414.660234 | 1.07137107 | 0.28116055 | 3.8105312 | 0.00013867 | 0.00204135 |
| TMEM243 | 145.322145 | 1.23009762 | 0.32281663 | 3.81051507 | 0.00013868 | 0.00204135 |
| LAG3 | 64.4828299 | 4.3044816 | 1.1303476 | 3.80810434 | 0.00014004 | 0.00205919 |
| DIXDC1 | 55.0266674 | -2.4277857 | 0.63795987 | -3.8055461 | 0.00014149 | 0.00207705 |
| PRKACB | 145.809897 | -3.2098686 | 0.84349239 | -3.8054506 | 0.00014155 | 0.00207705 |
| CNIH3 | 183.659391 | -2.7099467 | 0.71220912 | -3.8049874 | 0.00014181 | 0.00207877 |
| DSTN | 871.774977 | -1.0671378 | 0.28057203 | -3.8034361 | 0.0001427 | 0.00208965 |

| | | | | | | |
|---|---|---|---|---|---|---|
| ELAVL4 | 160.82699 | 1.6696749 | 0.43930422 | 3.80072583 | 0.00014427 | 0.00211044 |
| KCTD5 | 627.958281 | 0.91083936 | 0.23989422 | 3.79683752 | 0.00014655 | 0.0021395 |
| MT-ND3 | 23004.607 | 0.87685944 | 0.23094574 | 3.79682007 | 0.00014656 | 0.0021395 |
| RBMS2 | 204.903554 | 1.43699101 | 0.3787506 | 3.79402963 | 0.00014822 | 0.00216146 |
| TRAM2 | 314.479512 | -1.7759689 | 0.46820014 | -3.7931832 | 0.00014873 | 0.00216659 |
| TMEM123 | 3848.44738 | 1.04792074 | 0.27631564 | 3.79247718 | 0.00014915 | 0.00217051 |
| ARID3B | 194.206604 | 1.66324156 | 0.43872095 | 3.79111497 | 0.00014997 | 0.00218019 |
| PFDN1 | 917.09447 | 1.1612421 | 0.3063712 | 3.79031086 | 0.00015046 | 0.002185 |
| PLOD3 | 755.433024 | -1.2434893 | 0.32809455 | -3.7900335 | 0.00015063 | 0.00218518 |
| CD99 | 1320.33157 | -1.1256365 | 0.29713319 | -3.788323 | 0.00015167 | 0.00219801 |
| TRIM54 | 20.3268239 | -8.0452165 | 2.12450919 | -3.7868589 | 0.00015256 | 0.00220871 |
| DNAJB9 | 568.705348 | 1.23322365 | 0.32579467 | 3.78527877 | 0.00015354 | 0.00221816 |
| EHD2 | 22.7018796 | -2.7177974 | 0.71802169 | -3.7851189 | 0.00015364 | 0.00221816 |
| FRAT2 | 188.398849 | 1.32320594 | 0.34958929 | 3.78502995 | 0.00015369 | 0.00221816 |
| BLVRA | 225.467421 | -1.4229173 | 0.37611138 | -3.7832339 | 0.0001548 | 0.00223194 |
| OCSTAMP | 213.309261 | -7.1694598 | 1.89641174 | -3.7805397 | 0.00015649 | 0.00225392 |
| FOXJ2 | 279.858135 | 1.02943438 | 0.27233439 | 3.78003807 | 0.0001568 | 0.00225615 |
| PRPF38B | 248.249683 | 1.0301743 | 0.27256454 | 3.77956101 | 0.00015711 | 0.00225814 |
| EPN2 | 197.822912 | -1.725994 | 0.4566989 | -3.7792822 | 0.00015728 | 0.00225814 |
| FCGR1BP | 246.15119 | 1.82320218 | 0.4824493 | 3.77905446 | 0.00015742 | 0.00225814 |
| METRNL | 2317.28881 | 1.25035616 | 0.33090285 | 3.77862009 | 0.0001577 | 0.00225977 |
| TMT1B | 50.1139611 | -4.47167 | 1.18359605 | -3.7780373 | 0.00015807 | 0.00226276 |
| SLC7A5 | 3374.00732 | 1.75384689 | 0.4644582 | 3.77611351 | 0.00015929 | 0.00227798 |
| IRAK3 | 211.828352 | 1.5514349 | 0.41104327 | 3.77438337 | 0.0001604 | 0.00229151 |
| MSX2 | 19.1674547 | 3.6110771 | 0.95716818 | 3.77266733 | 0.00016151 | 0.00230281 |
| SLCO3A1 | 335.644151 | 1.53336209 | 0.40644168 | 3.77264975 | 0.00016152 | 0.00230281 |
| S100A12 | 37.2255146 | 3.96325393 | 1.05111947 | 3.77050758 | 0.00016292 | 0.00232031 |
| TNFSF12-TNF | 10.9675798 | 6.64166848 | 1.76380653 | 3.76553118 | 0.0001662 | 0.00236353 |
| GP1BA | 15.9639547 | -3.9468458 | 1.04818946 | -3.7653935 | 0.00016629 | 0.00236353 |
| PPP3CA | 767.535355 | 1.07608787 | 0.2859673 | 3.76297529 | 0.0001679 | 0.0023841 |
| MCOLN3 | 34.9234243 | -4.5799329 | 1.21760693 | -3.7614215 | 0.00016895 | 0.00239654 |
| MIDN | 1013.44235 | 1.1148656 | 0.29657974 | 3.75907543 | 0.00017054 | 0.00241668 |
| TRAPPC2L | 383.616178 | -1.4437484 | 0.38440733 | -3.7557774 | 0.0001728 | 0.00244626 |
| TNFRSF1B | 5903.66504 | 1.45183772 | 0.38668872 | 3.75453856 | 0.00017366 | 0.00245591 |
| SLC17A9 | 278.276677 | -2.7079551 | 0.72171443 | -3.7521143 | 0.00017535 | 0.0024773 |
| GRB2 | 4708.60998 | 0.83315324 | 0.22213295 | 3.75069635 | 0.00017634 | 0.00248851 |
| COG3 | 162.466775 | 1.0947284 | 0.29189034 | 3.75047835 | 0.0001765 | 0.00248851 |
| SLC17A5 | 916.120673 | -1.6458492 | 0.43887915 | -3.7501193 | 0.00017675 | 0.00248958 |
| IARS1 | 1264.09446 | -2.499036 | 0.66681873 | -3.7476992 | 0.00017846 | 0.00250985 |
| BCAR1 | 55.0253892 | -3.5692861 | 0.95242363 | -3.7475825 | 0.00017855 | 0.00250985 |
| EPB41L3 | 1772.79293 | 1.76328778 | 0.470793 | 3.74535686 | 0.00018014 | 0.00252967 |
| PTGER4 | 218.510551 | -2.2984637 | 0.61411367 | -3.7427333 | 0.00018203 | 0.00255369 |
| CYB5R2 | 26.5946064 | 2.6344276 | 0.70573014 | 3.73291071 | 0.00018928 | 0.00265275 |
| FIGNL2 | 8.84000157 | -6.8455918 | 1.83583107 | -3.7288789 | 0.00019233 | 0.00268917 |
| SUGCT-AS1 | 64.7702882 | -4.1038109 | 1.10059197 | -3.7287305 | 0.00019245 | 0.00268917 |
| FECH | 457.620713 | -1.3044543 | 0.34983945 | -3.7287226 | 0.00019245 | 0.00268917 |
| TIMP4 | 9.12466125 | -6.8896013 | 1.84807659 | -3.7279847 | 0.00019302 | 0.00269437 |
| RP2 | 1184.75803 | 0.83413673 | 0.22378928 | 3.72733109 | 0.00019352 | 0.00269868 |

| | | | | | | |
|---|---|---|---|---|---|---|
| FCER1G | 7860.74218 | 1.37359698 | 0.36862326 | 3.72628945 | 0.00019432 | 0.00270717 |
| SHMT1 | 217.005001 | -1.3721824 | 0.36858507 | -3.7228377 | 0.000197 | 0.00274174 |
| SIPA1L1 | 1186.79906 | 2.48378696 | 0.66729981 | 3.72214548 | 0.00019754 | 0.00274655 |
| PRNP | 2120.81779 | 0.94853805 | 0.25494026 | 3.72062866 | 0.00019873 | 0.00276037 |
| PPP2R3A | 447.664745 | -1.6237494 | 0.43653381 | -3.7196418 | 0.00019951 | 0.00276844 |
| MBNL3 | 414.186712 | 1.4700527 | 0.39531374 | 3.71869875 | 0.00020025 | 0.00277605 |
| ZBTB18 | 76.9624978 | 1.52242372 | 0.40974814 | 3.71551105 | 0.00020279 | 0.00280851 |
| STK17B | 927.430331 | 2.14852012 | 0.57835815 | 3.71486098 | 0.00020332 | 0.00281296 |
| HLA-DRB5 | 1965.55869 | 7.03715589 | 1.89471294 | 3.71410134 | 0.00020393 | 0.00281865 |
| VASH1 | 1829.26695 | 1.37546679 | 0.37046496 | 3.71281213 | 0.00020497 | 0.00283027 |
| MRTFA | 898.035312 | 1.1618917 | 0.31300109 | 3.71210111 | 0.00020555 | 0.00283544 |
| SRSF1 | 1964.94694 | 0.9080245 | 0.24480728 | 3.70914004 | 0.00020796 | 0.00286599 |
| GRK2 | 1380.78308 | 0.84374226 | 0.22752824 | 3.70829683 | 0.00020866 | 0.00287273 |
| LST1 | 1453.43731 | 1.50610488 | 0.40643542 | 3.70564378 | 0.00021085 | 0.00290013 |
| MAST3 | 703.433107 | 1.27602046 | 0.34465065 | 3.70235909 | 0.0002136 | 0.00293507 |
| INO80C | 103.53055 | -1.371619 | 0.37054874 | -3.7015886 | 0.00021425 | 0.00294112 |
| IRF8 | 1627.27068 | 1.70352392 | 0.46034385 | 3.7005467 | 0.00021514 | 0.00295034 |
| RNASEH2B | 126.269387 | -1.3053155 | 0.35287086 | -3.6991309 | 0.00021634 | 0.00296395 |
| ENTPD5 | 142.789127 | -1.4007194 | 0.37870954 | -3.6986641 | 0.00021674 | 0.00296651 |
| RBBP9 | 152.322276 | -1.1051548 | 0.29894596 | -3.696838 | 0.0002183 | 0.00298501 |
| TNS3 | 2957.41984 | -1.8601153 | 0.50343091 | -3.694877 | 0.00021999 | 0.00300522 |
| C2 | 486.37438 | -2.4804112 | 0.67167304 | -3.6928848 | 0.00022172 | 0.00302593 |
| SPIRE2 | 21.8322179 | -4.418324 | 1.19732334 | -3.6901678 | 0.00022411 | 0.00305546 |
| ARMCX1 | 107.428403 | -2.1178801 | 0.57401771 | -3.6895727 | 0.00022463 | 0.00305964 |
| ALAS1 | 1919.25543 | -1.2936537 | 0.35119138 | -3.6836146 | 0.00022995 | 0.00312906 |
| CIZ1 | 446.30498 | -1.0660205 | 0.28961701 | -3.680794 | 0.00023251 | 0.00315848 |
| CTNND1 | 1964.57684 | 1.01144706 | 0.27479479 | 3.68073591 | 0.00023256 | 0.00315848 |
| CUL7 | 191.180313 | -1.5003311 | 0.4076544 | -3.6803997 | 0.00023287 | 0.00315959 |
| ZNF264 | 318.989429 | 1.51617639 | 0.41207296 | 3.67938816 | 0.00023379 | 0.00316908 |
| GUCY1A1 | 28.9659854 | -4.235623 | 1.15214011 | -3.676309 | 0.00023663 | 0.00320447 |
| BTG2 | 2677.34775 | 1.12622035 | 0.30638626 | 3.67581871 | 0.00023709 | 0.00320754 |
| ACTG1 | 48906.3965 | -0.8149802 | 0.22176751 | -3.6749306 | 0.00023791 | 0.00321562 |
| SYT4 | 74.4998157 | 4.04273864 | 1.10016037 | 3.6746812 | 0.00023815 | 0.00321567 |
| MTHFD1 | 736.614105 | -2.3272008 | 0.63380112 | -3.6718156 | 0.00024083 | 0.00324882 |
| CTC1 | 131.924511 | -1.8220166 | 0.49671283 | -3.668149 | 0.00024431 | 0.00329089 |
| AGO4 | 293.467156 | -1.097874 | 0.29930838 | -3.6680362 | 0.00024442 | 0.00329089 |
| NRAP | 13.5059229 | -6.4725025 | 1.76555899 | -3.6659792 | 0.00024639 | 0.00331428 |
| KLF9 | 256.070744 | 1.36249971 | 0.37175267 | 3.66507043 | 0.00024727 | 0.00331972 |
| RAB42 | 409.221384 | -1.0523955 | 0.28716308 | -3.6648009 | 0.00024753 | 0.00331972 |
| TES | 233.303254 | 1.15221927 | 0.31441842 | 3.66460489 | 0.00024772 | 0.00331972 |
| ZNF33A | 223.748485 | 1.70087069 | 0.46413779 | 3.66458132 | 0.00024774 | 0.00331972 |
| NEU1 | 2559.39545 | -0.7001191 | 0.19106681 | -3.6642631 | 0.00024805 | 0.00332068 |
| PRSS22 | 25.2469745 | -4.590979 | 1.2535511 | -3.6623788 | 0.00024988 | 0.00333824 |
| UTS2R | 53.4338432 | 2.91162127 | 0.79502763 | 3.6622894 | 0.00024997 | 0.00333824 |
| SLC27A3 | 262.027545 | -1.8823856 | 0.5140067 | -3.6621811 | 0.00025008 | 0.00333824 |
| RRAGD | 716.778145 | -1.2904431 | 0.35244772 | -3.6613746 | 0.00025087 | 0.00334278 |
| STK38 | 222.234286 | -1.3079924 | 0.35724712 | -3.66131 | 0.00025093 | 0.00334278 |
| SLC28A1 | 17.2202044 | -5.943587 | 1.62344132 | -3.6611037 | 0.00025113 | 0.00334278 |

| | | | | | | |
|---|---|---|---|---|---|---|
| TMEM18 | 167.05801 | -1.1872029 | 0.32433599 | -3.6604105 | 0.00025181 | 0.00334866 |
| SH2D3C | 102.175294 | 1.63186596 | 0.44597282 | 3.65911523 | 0.00025309 | 0.00336245 |
| HS2ST1 | 668.566847 | -1.0716438 | 0.29307551 | -3.6565451 | 0.00025564 | 0.00339312 |
| RNF122 | 161.927775 | -2.1187008 | 0.57972325 | -3.6546763 | 0.00025751 | 0.0034147 |
| TMEM38B | 224.617245 | -1.7481434 | 0.47855694 | -3.6529477 | 0.00025925 | 0.00343453 |
| ENSG000002 | 18.9748528 | -6.9693932 | 1.90886052 | -3.6510751 | 0.00026114 | 0.00345642 |
| C5orf15 | 548.232946 | -0.8817811 | 0.24154234 | -3.6506273 | 0.0002616 | 0.00345919 |
| KCTD15 | 46.9731408 | 3.64500163 | 0.99853759 | 3.65033992 | 0.00026189 | 0.0034598 |
| NUDT17 | 26.0208314 | -3.3377598 | 0.91472537 | -3.6489201 | 0.00026334 | 0.00347344 |
| FOXN3 | 1671.76897 | 1.32152733 | 0.36217676 | 3.64884629 | 0.00026342 | 0.00347344 |
| ZBTB4 | 944.874932 | 1.21148114 | 0.33205388 | 3.64844743 | 0.00026383 | 0.00347557 |
| RIC1 | 350.086207 | 0.95604733 | 0.26208806 | 3.64780954 | 0.00026449 | 0.00348095 |
| ANOS1 | 1341.86323 | 2.60936734 | 0.7155225 | 3.64679984 | 0.00026553 | 0.00349138 |
| CXCL2 | 31.3030416 | 2.88067918 | 0.79018885 | 3.64555787 | 0.00026681 | 0.00350244 |
| DAZAP2 | 6449.89315 | 1.02432736 | 0.28098365 | 3.64550521 | 0.00026687 | 0.00350244 |
| MTF1 | 635.897626 | 1.11343038 | 0.30567119 | 3.64257545 | 0.00026992 | 0.00353925 |
| MYRF | 23.4440008 | -4.9920042 | 1.37143167 | -3.6399949 | 0.00027264 | 0.00357158 |
| CCDC47 | 1146.02738 | 0.83191408 | 0.22864272 | 3.63848921 | 0.00027424 | 0.00358917 |
| MOB3B | 539.659676 | 2.0600335 | 0.56624166 | 3.63808184 | 0.00027468 | 0.00359151 |
| ARHGAP45 | 948.653266 | 1.04656455 | 0.28777776 | 3.63671103 | 0.00027614 | 0.00360731 |
| MIDEAS | 805.086387 | 1.44097226 | 0.39628123 | 3.6362365 | 0.00027665 | 0.0036106 |
| NCK1-DT | 33.7722144 | 2.57820501 | 0.70929343 | 3.63489199 | 0.0002781 | 0.00362611 |
| MPC1 | 352.195147 | -1.2142566 | 0.33409447 | -3.6344708 | 0.00027855 | 0.00362867 |
| LPIN2 | 756.563196 | 0.9495679 | 0.26133849 | 3.63347891 | 0.00027963 | 0.00363928 |
| LRRC8C | 661.856279 | 1.63316345 | 0.44961374 | 3.63236997 | 0.00028083 | 0.00365157 |
| GALC | 1552.24883 | 1.12120614 | 0.30881452 | 3.63067818 | 0.00028268 | 0.00367008 |
| TREM1 | 710.886879 | 3.63537602 | 1.00131875 | 3.63058817 | 0.00028278 | 0.00367008 |
| RIGI | 64.1306413 | -2.3768093 | 0.65474946 | -3.6301049 | 0.00028331 | 0.00367356 |
| LPAR5 | 457.244708 | 1.49578612 | 0.41220828 | 3.62871436 | 0.00028484 | 0.00368999 |
| DUSP13B | 11.0888803 | -6.1803947 | 1.70395907 | -3.6270793 | 0.00028665 | 0.00371001 |
| CMTM4 | 109.501955 | 1.62161252 | 0.4473666 | 3.62479565 | 0.00028919 | 0.0037395 |
| CIBAR1 | 36.3929516 | -2.713655 | 0.74888182 | -3.6236092 | 0.00029052 | 0.00375325 |
| RWDD2B | 88.2197148 | -1.5401998 | 0.42513955 | -3.62281 | 0.00029142 | 0.00376141 |
| NPTXR | 36.6853567 | -3.1247019 | 0.86290037 | -3.6211618 | 0.00029328 | 0.00378086 |
| CALCRL | 78.3753992 | 2.81396008 | 0.77712219 | 3.62100083 | 0.00029347 | 0.00378086 |
| PTGR3 | 328.752673 | -1.6188794 | 0.44711777 | -3.6207002 | 0.00029381 | 0.00378179 |
| DTX4 | 188.490265 | -2.1374109 | 0.5909943 | -3.6166354 | 0.00029846 | 0.00383813 |
| MDH1 | 1736.74736 | -1.1245671 | 0.31119044 | -3.6137585 | 0.00030179 | 0.00387744 |
| ZNRF2 | 329.218647 | 1.30744629 | 0.36183601 | 3.61336701 | 0.00030225 | 0.00387975 |
| B3GNT9 | 38.2477894 | 2.03249227 | 0.56274746 | 3.61173069 | 0.00030416 | 0.00390075 |
| ZDHHC8 | 220.28779 | -1.3793186 | 0.38203438 | -3.6104567 | 0.00030566 | 0.00391639 |
| CCRL2 | 514.276491 | -2.3068849 | 0.63907849 | -3.6097051 | 0.00030655 | 0.00392417 |
| GIMAP4 | 150.659555 | 2.13243371 | 0.59080388 | 3.60937664 | 0.00030693 | 0.00392556 |
| ASS1 | 64.0764088 | -2.3194637 | 0.64267826 | -3.6090588 | 0.00030731 | 0.0039268 |
| ERP44 | 1621.99738 | 0.74784796 | 0.20723333 | 3.60872437 | 0.00030771 | 0.00392829 |
| GGT7 | 120.584618 | 1.41162975 | 0.39143092 | 3.60633175 | 0.00031056 | 0.00396107 |
| AKAP1 | 211.351485 | -1.0535718 | 0.29222009 | -3.605405 | 0.00031167 | 0.00397163 |
| NAMPT | 2193.87628 | 2.05014232 | 0.56876469 | 3.60455276 | 0.00031269 | 0.00398107 |

| | | | | | | |
|---|---|---|---|---|---|---|
| PLEKHO2 | 6921.6631 | 0.86491286 | 0.24003005 | 3.60335244 | 0.00031414 | 0.00399588 |
| SHISAL2A | 10.8754251 | 6.62623854 | 1.8398199 | 3.60156913 | 0.0003163 | 0.00401976 |
| PPM1D | 183.49852 | 0.98936255 | 0.2748243 | 3.59998211 | 0.00031824 | 0.00404072 |
| LAMB3 | 108.785467 | 2.32390781 | 0.64615318 | 3.59652767 | 0.00032249 | 0.00409014 |
| PSAP | 70030.4946 | 0.99782884 | 0.2774592 | 3.5963083 | 0.00032277 | 0.00409014 |
| STK4 | 1292.29893 | 0.88303667 | 0.24555296 | 3.59611494 | 0.00032301 | 0.00409014 |
| PSAT1 | 97.2860673 | -2.5843888 | 0.7189341 | -3.5947506 | 0.0003247 | 0.00410792 |
| KLHL36 | 644.229609 | 0.76174291 | 0.21191812 | 3.59451519 | 0.000325 | 0.00410794 |
| SCRN1 | 747.205649 | -1.6560054 | 0.46082017 | -3.5936044 | 0.00032613 | 0.00411863 |
| CCN3 | 17.6995742 | -6.8679005 | 1.91163601 | -3.5926821 | 0.00032729 | 0.00412952 |
| SRD5A1 | 171.203172 | -1.9157763 | 0.53372679 | -3.5894325 | 0.0003314 | 0.00417427 |
| ACE | 158.806922 | 2.57593063 | 0.7176483 | 3.58940533 | 0.00033143 | 0.00417427 |
| REEP4 | 343.893017 | 1.40427641 | 0.39131981 | 3.5885646 | 0.0003325 | 0.004184 |
| NAB2 | 864.534332 | 1.11989535 | 0.31247396 | 3.58396374 | 0.00033842 | 0.00425463 |
| MINDY1 | 147.900559 | -2.6111708 | 0.72902524 | -3.5817289 | 0.00034133 | 0.00428736 |
| CLN8 | 977.786138 | 1.2777911 | 0.35684428 | 3.58080872 | 0.00034253 | 0.0042965 |
| CD44 | 8021.06893 | 1.3729135 | 0.38341983 | 3.58070548 | 0.00034267 | 0.0042965 |
| LPCAT2 | 2745.39887 | 1.22398529 | 0.34199343 | 3.57897317 | 0.00034495 | 0.00432122 |
| ACOT11 | 311.184729 | 1.04124667 | 0.29103755 | 3.57770562 | 0.00034662 | 0.00433835 |
| CD99.1 | 1278.52396 | -0.8137747 | 0.22750324 | -3.5769806 | 0.00034759 | 0.00434653 |
| ENSG000002 | 12.2306442 | -6.3272063 | 1.76967166 | -3.575356 | 0.00034975 | 0.00436972 |
| PIK3R5 | 4490.37104 | 1.72388916 | 0.48230764 | 3.57425224 | 0.00035123 | 0.00438429 |
| MMP14 | 12721.6874 | 1.39169541 | 0.38943625 | 3.5736155 | 0.00035209 | 0.00439107 |
| TLN1 | 6584.06039 | -0.901236 | 0.25237358 | -3.5710394 | 0.00035557 | 0.00443055 |
| SULT1C2 | 28.0878898 | -3.5354638 | 0.99015466 | -3.5706177 | 0.00035614 | 0.00443376 |
| SEC61G | 182.30947 | 1.18675195 | 0.33240988 | 3.57014642 | 0.00035678 | 0.00443781 |
| HLA-DMB | 1636.75115 | 2.80215782 | 0.78523885 | 3.56854201 | 0.00035897 | 0.00446112 |
| GRIK4 | 11.5089072 | -7.2239602 | 2.02500676 | -3.5673759 | 0.00036057 | 0.00447705 |
| DNAH17 | 9.19493948 | 6.38579548 | 1.79060236 | 3.5662834 | 0.00036208 | 0.0044881 |
| CD163L1 | 1486.69422 | 3.19799873 | 0.89673584 | 3.56626621 | 0.0003621 | 0.0044881 |
| SLC5A4-AS1 | 10.8453897 | -7.1382467 | 2.00219118 | -3.5652173 | 0.00036355 | 0.00450212 |
| ADGRA2 | 35.9779315 | 4.09357293 | 1.14866099 | 3.56377814 | 0.00036555 | 0.0045229 |
| CD93 | 1505.24687 | 2.86625929 | 0.80477078 | 3.56158468 | 0.00036862 | 0.00455684 |
| TMSB10 | 6986.53767 | -0.8845606 | 0.24840436 | -3.5609703 | 0.00036949 | 0.0045635 |
| DYRK4 | 181.101741 | -0.9746033 | 0.27373566 | -3.5603814 | 0.00037032 | 0.00456973 |
| CD207 | 16.3335285 | 5.76405219 | 1.61947224 | 3.55921641 | 0.00037196 | 0.00458602 |
| KCNF1 | 12.1724247 | -7.3048896 | 2.05263489 | -3.5587866 | 0.00037257 | 0.00458842 |
| AGRN | 503.122305 | -2.1212218 | 0.59608024 | -3.5586179 | 0.00037281 | 0.00458842 |
| MCL1 | 2671.27792 | 0.98172389 | 0.27589301 | 3.55834996 | 0.00037319 | 0.00458908 |
| SPI1 | 6259.18263 | 0.83143039 | 0.233745 | 3.5569976 | 0.00037512 | 0.00460873 |
| ANXA6 | 2377.19282 | 1.5916773 | 0.44756198 | 3.55632819 | 0.00037607 | 0.00461145 |
| LPAR1 | 499.232739 | 1.19087823 | 0.33486244 | 3.55632068 | 0.00037608 | 0.00461145 |
| WDR13 | 329.219274 | -1.341546 | 0.37724635 | -3.5561537 | 0.00037632 | 0.00461145 |
| ARHGAP5 | 806.101633 | -1.4976917 | 0.42147692 | -3.5534371 | 0.00038023 | 0.00465527 |
| TNFAIP2 | 1061.81192 | 1.65208709 | 0.46550318 | 3.54903503 | 0.00038665 | 0.00472967 |
| GTF3A | 962.053369 | -1.4587608 | 0.41107612 | -3.5486392 | 0.00038723 | 0.00473259 |
| ATE1 | 268.514012 | -1.0024968 | 0.28251968 | -3.5484141 | 0.00038756 | 0.00473259 |
| CSF1R | 22529.7407 | 1.76507153 | 0.49751563 | 3.54777099 | 0.00038851 | 0.00474004 |

| | | | | | | |
|---|---|---|---|---|---|---|
| JPT1 | 785.394092 | 1.01173824 | 0.28523379 | 3.54704899 | 0.00038957 | 0.00474892 |
| NRAS | 1846.17688 | -0.8228729 | 0.23215389 | -3.5445146 | 0.00039334 | 0.00479066 |
| C6orf47 | 268.747537 | 1.09552557 | 0.30920152 | 3.54307949 | 0.00039548 | 0.00481263 |
| NAMPTP1 | 72.9208783 | 2.33044648 | 0.65788655 | 3.54232273 | 0.00039662 | 0.00482228 |
| ENSG000002 | 125.352368 | -2.9559593 | 0.83478605 | -3.5409784 | 0.00039865 | 0.00484182 |
| ANKH | 9894.56759 | 1.93269734 | 0.54586489 | 3.54061489 | 0.0003992 | 0.00484182 |
| INKA1 | 56.2630585 | -2.7872305 | 0.78722611 | -3.5405717 | 0.00039926 | 0.00484182 |
| SOX4 | 158.0282 | -1.5921711 | 0.44988005 | -3.5391013 | 0.00040149 | 0.00486296 |
| HLA-E | 8366.0154 | 1.27333535 | 0.35980432 | 3.53896627 | 0.0004017 | 0.00486296 |
| CUBN | 10.7737633 | -5.2380931 | 1.48095209 | -3.5369768 | 0.00040474 | 0.00489552 |
| ID2 | 2360.54574 | 1.03971576 | 0.29397729 | 3.53672137 | 0.00040513 | 0.00489604 |
| SUCNR1 | 870.950916 | -2.3632553 | 0.66869482 | -3.5341313 | 0.00040912 | 0.00494002 |
| CARS2 | 337.556521 | -1.972669 | 0.55836883 | -3.5329138 | 0.00041101 | 0.00495856 |
| MCCC1 | 176.796568 | -1.0266513 | 0.29070135 | -3.5316358 | 0.000413 | 0.00497625 |
| PLEK2 | 239.802084 | 1.8281918 | 0.51767872 | 3.53151818 | 0.00041318 | 0.00497625 |
| TMEM39B | 168.756604 | 1.91703808 | 0.54350547 | 3.52717348 | 0.00042002 | 0.00505429 |
| MFSD14CP | 77.7181281 | -1.4759744 | 0.41880018 | -3.5242927 | 0.00042461 | 0.00510519 |
| FGR | 1734.69217 | -2.4138866 | 0.68508679 | -3.5234756 | 0.00042593 | 0.00511658 |
| VWA8 | 245.949217 | -1.7869085 | 0.50717742 | -3.5232414 | 0.0004263 | 0.00511673 |
| POMT2 | 112.692485 | -1.8683362 | 0.53038228 | -3.5226219 | 0.0004273 | 0.00512432 |
| VWF | 196.014024 | -2.9295003 | 0.8322113 | -3.52014 | 0.00043132 | 0.00516709 |
| LYRM9 | 60.7441763 | -1.667413 | 0.47370149 | -3.5199658 | 0.0004316 | 0.00516709 |
| SGK1 | 2092.54477 | -0.9322975 | 0.26490193 | -3.5194061 | 0.00043251 | 0.00517359 |
| TMEM131L | 65.4400567 | 2.17066944 | 0.61729454 | 3.5164242 | 0.0004374 | 0.0052276 |
| TMEM53 | 185.739021 | -1.3113703 | 0.37309465 | -3.5148463 | 0.00044001 | 0.00525429 |
| TNFAIP6 | 29.8745947 | 3.36262695 | 0.95676685 | 3.51457303 | 0.00044046 | 0.00525523 |
| DOCK2 | 2871.58655 | 1.14296864 | 0.32559814 | 3.51036595 | 0.00044749 | 0.00533456 |
| PCK2 | 440.254478 | -1.2687389 | 0.36150031 | -3.5096481 | 0.0004487 | 0.00534394 |
| TBX3 | 19.2995214 | 3.33439447 | 0.95011945 | 3.50944764 | 0.00044904 | 0.00534394 |
| ENPP4 | 44.0074751 | -2.7633274 | 0.78775131 | -3.5078677 | 0.00045171 | 0.00537122 |
| SPCS2 | 895.635207 | 0.88435676 | 0.2521332 | 3.50749828 | 0.00045234 | 0.00537414 |
| COL27A1 | 14.9971735 | -5.105945 | 1.45794185 | -3.5021596 | 0.0004615 | 0.00547835 |
| B3GNT5 | 702.475897 | 1.03167414 | 0.29464401 | 3.50142577 | 0.00046278 | 0.00548882 |
| MT-RNR2 | 170133.051 | 0.88334426 | 0.25236726 | 3.50023323 | 0.00046485 | 0.00550878 |
| MID1IP1 | 412.593453 | -1.5635006 | 0.44694949 | -3.4981595 | 0.00046848 | 0.00554711 |
| ACOT2 | 405.626844 | -1.6684494 | 0.47716996 | -3.4965515 | 0.00047131 | 0.00557237 |
| FIGN | 17.1042863 | -3.2657837 | 0.93401556 | -3.4964981 | 0.00047141 | 0.00557237 |
| FDPS | 962.432375 | -0.8695735 | 0.24885415 | -3.49431 | 0.00047529 | 0.00561352 |
| SERPINB8 | 873.4459 | 1.16558542 | 0.33366924 | 3.49323604 | 0.0004772 | 0.00563141 |
| NR1H2 | 744.950137 | 0.70893531 | 0.20304083 | 3.49158987 | 0.00048015 | 0.00566147 |
| MAP4K3 | 209.975997 | -1.3118511 | 0.37576254 | -3.4911705 | 0.00048091 | 0.00566561 |
| IPO9 | 645.760977 | -0.8838198 | 0.25327287 | -3.4895953 | 0.00048375 | 0.00569433 |
| GYG2 | 17.4540209 | -3.4656785 | 0.99361719 | -3.4879414 | 0.00048675 | 0.00572487 |
| VWCE | 31.9674294 | -3.072108 | 0.88139049 | -3.4855243 | 0.00049117 | 0.00577201 |
| DOT1L | 1426.94425 | -1.2273587 | 0.35217135 | -3.485118 | 0.00049192 | 0.00577305 |
| CBL | 1750.70219 | 0.9430461 | 0.27059925 | 3.48502849 | 0.00049208 | 0.00577305 |
| RGS20 | 32.5022781 | -4.7059447 | 1.35110807 | -3.4830261 | 0.00049578 | 0.00581155 |
| TM9SF3 | 2315.33875 | 0.89928811 | 0.25823685 | 3.48241587 | 0.00049691 | 0.00581995 |

| | | | | | | |
|---|---|---|---|---|---|---|
| EGFR | 15.6616827 | 4.10810404 | 1.18008098 | 3.4812052 | 0.00049916 | 0.00584145 |
| AZU1 | 16.7506335 | 4.80650213 | 1.38163338 | 3.47885496 | 0.00050356 | 0.00588802 |
| SLC41A2 | 1126.19646 | 1.66461786 | 0.47901803 | 3.4750631 | 0.00051073 | 0.00596691 |
| SSH1 | 1680.41149 | 1.02176564 | 0.2940637 | 3.47464043 | 0.00051154 | 0.00596834 |
| ECE1 | 30.4206541 | -3.8850148 | 1.11813375 | -3.4745528 | 0.00051171 | 0.00596834 |
| TP53I3 | 141.090796 | -1.6430091 | 0.47302482 | -3.4734099 | 0.00051389 | 0.00598884 |
| CD36 | 13001.1732 | -1.3268594 | 0.38212753 | -3.4722947 | 0.00051603 | 0.00600407 |
| TEX2 | 1504.93225 | -2.1136533 | 0.60872147 | -3.4722832 | 0.00051605 | 0.00600407 |
| SLC7A11 | 147.562946 | 2.96956767 | 0.85588452 | 3.46958917 | 0.00052126 | 0.00605793 |
| RHBDF1 | 35.0569073 | -2.6542848 | 0.76509492 | -3.4692229 | 0.00052197 | 0.00605793 |
| ENSG000002 | 16.687201 | -6.779834 | 1.95428295 | -3.4692182 | 0.00052198 | 0.00605793 |
| CTSA | 7221.51359 | 0.67433432 | 0.19449186 | 3.46715963 | 0.00052599 | 0.00609948 |
| FUBP1 | 631.829135 | 0.77375407 | 0.22324094 | 3.46600438 | 0.00052825 | 0.00612069 |
| TMEM220 | 39.001646 | -1.8410532 | 0.53125636 | -3.4654705 | 0.0005293 | 0.00612758 |
| CD86 | 5479.44259 | 2.61326806 | 0.75413358 | 3.46525884 | 0.00052972 | 0.00612758 |
| CLEC17A | 14.1845768 | 3.93652348 | 1.13608669 | 3.46498511 | 0.00053026 | 0.00612877 |
| ADAMTS10 | 19.3975968 | 6.01297203 | 1.73613414 | 3.46342595 | 0.00053334 | 0.00615933 |
| MYO10 | 299.839076 | -1.5696548 | 0.45359337 | -3.4604889 | 0.0005392 | 0.00622179 |
| HIF1A | 5572.92305 | 1.33059314 | 0.38466252 | 3.45911826 | 0.00054195 | 0.00624841 |
| CXCR4 | 85.1871665 | 3.34866198 | 0.9688242 | 3.4564186 | 0.0005474 | 0.00630606 |
| RTN4R | 68.9551685 | -4.1286952 | 1.19457586 | -3.4562017 | 0.00054785 | 0.00630606 |
| UCHL1 | 547.885272 | -5.8486269 | 1.69277687 | -3.4550489 | 0.00055019 | 0.00631953 |
| TNFSF12 | 245.081472 | 1.30904245 | 0.37889134 | 3.4549284 | 0.00055044 | 0.00631953 |
| PTPRJ | 3316.48133 | 0.88561992 | 0.25634735 | 3.4547653 | 0.00055077 | 0.00631953 |
| BMAL2 | 83.3779352 | -2.2160115 | 0.64144135 | -3.4547376 | 0.00055083 | 0.00631953 |
| PLEKHA6 | 18.5577357 | -3.7014664 | 1.07148387 | -3.4545237 | 0.00055127 | 0.00631953 |
| PON2 | 336.69888 | 1.14012606 | 0.33012134 | 3.45365754 | 0.00055304 | 0.00633469 |
| HNRNPH2 | 1041.27527 | 0.90079906 | 0.26088172 | 3.45290212 | 0.00055459 | 0.00634728 |
| IL1RN | 3855.29333 | -2.5764932 | 0.74648036 | -3.4515218 | 0.00055743 | 0.0063719 |
| PAK1 | 800.136686 | 1.01138215 | 0.29303379 | 3.4514182 | 0.00055765 | 0.0063719 |
| NEMP1 | 813.335667 | 1.62346028 | 0.47058194 | 3.44989926 | 0.0005608 | 0.00639883 |
| EFR3B | 40.4707859 | -4.3548227 | 1.26232574 | -3.4498407 | 0.00056092 | 0.00639883 |
| MX1 | 325.076294 | -1.7129073 | 0.49658996 | -3.4493393 | 0.00056196 | 0.00640552 |
| NUP210 | 462.429328 | -1.7997529 | 0.52221074 | -3.4464112 | 0.00056809 | 0.00647009 |
| ENSG000002 | 15.8540687 | 3.30747479 | 0.96029229 | 3.44423757 | 0.00057267 | 0.00651674 |
| SSC5D | 9.6592748 | -5.0745436 | 1.47343128 | -3.4440314 | 0.00057311 | 0.00651674 |
| SIGLEC9 | 1180.67748 | 1.39093343 | 0.40398544 | 3.4430286 | 0.00057524 | 0.00653566 |
| RUSC2 | 154.075379 | -1.6462495 | 0.478409 | -3.4410922 | 0.00057937 | 0.00657729 |
| FAM135A | 239.1718 | 1.21414294 | 0.35295164 | 3.43996965 | 0.00058178 | 0.0065993 |
| CSTB | 6606.12178 | -2.1237112 | 0.6178263 | -3.437392 | 0.00058734 | 0.00665705 |
| FLNA | 10509.112 | -2.2182979 | 0.64546146 | -3.4367627 | 0.00058871 | 0.00666715 |
| IDUA | 98.6356709 | 1.61601628 | 0.47065428 | 3.43355272 | 0.00059573 | 0.00674116 |
| CDK4 | 577.834549 | -1.0317918 | 0.3005717 | -3.4327643 | 0.00059746 | 0.00675535 |
| PTGES3L-AAR | 9.57011107 | -6.9577147 | 2.02714717 | -3.4322691 | 0.00059855 | 0.00676226 |
| GPR85 | 58.5252033 | -2.5790695 | 0.75205791 | -3.4293497 | 0.00060503 | 0.00682993 |
| SLC5A6 | 467.428441 | -1.3873401 | 0.40466197 | -3.4283927 | 0.00060717 | 0.00684855 |
| SLC12A5 | 294.0802 | 1.30080152 | 0.3794555 | 3.42807397 | 0.00060788 | 0.00684891 |
| ITGA4 | 1929.18242 | 1.67623873 | 0.48899265 | 3.42794258 | 0.00060817 | 0.00684891 |

| | | | | | | |
|---|---|---|---|---|---|---|
| ENSG000002 | 18.0066591 | 4.32450193 | 1.26197125 | 3.42678325 | 0.00061078 | 0.0068727 |
| SUB1 | 1410.25899 | 0.97009346 | 0.28342165 | 3.4227924 | 0.00061981 | 0.00696881 |
| CXCL6 | 19.4892448 | 5.03842874 | 1.47237146 | 3.42198208 | 0.00062166 | 0.00698402 |
| CA2 | 371.971999 | -2.92926 | 0.85646616 | -3.4201701 | 0.00062582 | 0.00702509 |
| PTEN | 905.206843 | 0.88276929 | 0.2581442 | 3.41967505 | 0.00062696 | 0.00703227 |
| LPL | 9585.83899 | -1.6384918 | 0.4793255 | -3.4183281 | 0.00063007 | 0.00706152 |
| PLSCR1 | 1093.237 | 1.43299765 | 0.41944378 | 3.41642368 | 0.00063449 | 0.00710543 |
| SLITRK5 | 7.42428972 | -6.5925996 | 1.93031337 | -3.4153002 | 0.00063712 | 0.00712912 |
| BDH1 | 89.7539016 | -2.0906792 | 0.61227394 | -3.4146141 | 0.00063872 | 0.00714141 |
| SNRK | 415.561126 | 0.96975264 | 0.28403648 | 3.41418342 | 0.00063973 | 0.00714701 |
| FHL3 | 476.820343 | 1.21084875 | 0.35468955 | 3.41382697 | 0.00064057 | 0.00715068 |
| DHRS11 | 92.062625 | -2.5480262 | 0.74667449 | -3.4124993 | 0.0006437 | 0.0071799 |
| CCR5AS | 128.277054 | 1.65585615 | 0.48536415 | 3.4115749 | 0.00064589 | 0.00719857 |
| AIDA | 529.887136 | -0.8467651 | 0.24822589 | -3.4112684 | 0.00064661 | 0.00719862 |
| ARHGAP18 | 1874.05135 | 1.17407094 | 0.34418721 | 3.41114054 | 0.00064692 | 0.00719862 |
| NTRK3 | 58.1401381 | 3.39583159 | 0.99753141 | 3.40423526 | 0.0006635 | 0.00737725 |
| TSPAN31 | 439.624589 | 1.06395466 | 0.31261055 | 3.40345084 | 0.0006654 | 0.00739262 |
| N4BP1 | 654.695646 | 0.90911521 | 0.26716458 | 3.40282839 | 0.00066692 | 0.00740362 |
| PTPRB | 9.2960386 | -5.9242716 | 1.74159969 | -3.4016265 | 0.00066986 | 0.00743038 |
| NBL1 | 1210.59219 | 1.89374355 | 0.55685017 | 3.40081343 | 0.00067186 | 0.00744663 |
| BRSK2 | 20.1583594 | -2.5023349 | 0.7360116 | -3.399858 | 0.00067421 | 0.00746681 |
| INPP5A | 72.2368528 | -1.7536561 | 0.51594557 | -3.3989168 | 0.00067653 | 0.00748664 |
| METTL1 | 211.963266 | -0.9222763 | 0.27140388 | -3.3981692 | 0.00067838 | 0.00750122 |
| THBS3 | 83.597466 | -2.0088463 | 0.59128469 | -3.3974264 | 0.00068023 | 0.0075157 |
| NR3C1 | 943.347214 | 1.11232999 | 0.32747046 | 3.39673378 | 0.00068195 | 0.00752882 |
| LINC02712 | 11.1093001 | 6.65997451 | 1.96144954 | 3.39543505 | 0.0006852 | 0.00755869 |
| HHAT | 74.781246 | -2.9054843 | 0.85591226 | -3.3946053 | 0.00068728 | 0.00757568 |
| PDE7B | 15.2201256 | 4.60138784 | 1.35621813 | 3.39280808 | 0.0006918 | 0.00761958 |
| ENSG000002 | 11.6087205 | 6.7219632 | 1.98197992 | 3.39153951 | 0.00069501 | 0.00764894 |
| EIF2D | 337.55027 | -0.8584403 | 0.25318045 | -3.3906266 | 0.00069733 | 0.00766845 |
| HCFC1R1 | 406.285772 | -1.4994711 | 0.44242166 | -3.3892352 | 0.00070088 | 0.00770144 |
| TMT1A | 447.464298 | -1.5821833 | 0.46696903 | -3.3881974 | 0.00070354 | 0.00772459 |
| TBC1D1 | 1315.40356 | -1.362947 | 0.4023912 | -3.3871192 | 0.00070631 | 0.00774896 |
| H2AC6 | 85.7910449 | -1.7040255 | 0.50324147 | -3.3860991 | 0.00070894 | 0.00777175 |
| GASK1B | 3413.25426 | 1.01251125 | 0.29909571 | 3.38524166 | 0.00071116 | 0.00778999 |
| XPNPEP2 | 27.101518 | -4.2716545 | 1.26272823 | -3.3828772 | 0.00071731 | 0.00785124 |
| FCER2 | 10.3961466 | -7.0804811 | 2.09510195 | -3.3795401 | 0.00072607 | 0.00794099 |
| PPARD | 1758.561 | 1.10853861 | 0.32805391 | 3.37913552 | 0.00072714 | 0.0079465 |
| NDFIP2 | 104.231158 | -1.4701667 | 0.4351104 | -3.3788361 | 0.00072793 | 0.00794897 |
| MRRF | 235.92053 | -0.9341957 | 0.2766479 | -3.37684 | 0.00073324 | 0.00800066 |
| HPS5 | 641.170572 | 0.69534229 | 0.20601084 | 3.3752704 | 0.00073743 | 0.00804018 |
| FAM86B1 | 13.1114704 | -3.5621491 | 1.05559725 | -3.3745342 | 0.00073941 | 0.00805547 |
| AP1B1 | 12462.5551 | 1.29673319 | 0.38459705 | 3.37166702 | 0.00074715 | 0.00813348 |
| VOPP1 | 1268.2091 | -0.9530864 | 0.2827542 | -3.3707243 | 0.00074971 | 0.00815504 |
| NLRC4 | 202.723048 | 1.22332697 | 0.36306876 | 3.36940848 | 0.0007533 | 0.00818588 |
| SYNGR1 | 51.6394644 | -3.2666551 | 0.96954749 | -3.3692575 | 0.00075371 | 0.00818588 |
| ETV6 | 1279.04523 | 1.06971094 | 0.31761174 | 3.36798296 | 0.0007572 | 0.00821542 |
| FZD6 | 46.2895489 | -2.3833966 | 0.70771915 | -3.3677153 | 0.00075794 | 0.00821542 |

| | | | | | | |
|---|---|---|---|---|---|---|
| MDK | 12.9318076 | -4.3955187 | 1.30522777 | -3.3676258 | 0.00075818 | 0.00821542 |
| WDR90 | 29.8707338 | -2.3271309 | 0.69116149 | -3.3669858 | 0.00075995 | 0.00822816 |
| GPX1P1 | 210.33303 | 0.93762906 | 0.27858356 | 3.36570137 | 0.00076349 | 0.0082602 |
| SRRM3 | 35.3020034 | 4.08724413 | 1.2145459 | 3.36524469 | 0.00076476 | 0.00826751 |
| NAT8L | 144.872205 | 1.20983535 | 0.35959318 | 3.36445575 | 0.00076695 | 0.00828481 |
| C7orf50 | 362.762282 | -1.5609385 | 0.46411936 | -3.3632265 | 0.00077037 | 0.00831539 |
| FHIP1A | 15.413476 | 4.68361461 | 1.3933144 | 3.36149156 | 0.00077523 | 0.00835936 |
| PTPRA | 576.319162 | -1.561998 | 0.46469415 | -3.3613463 | 0.00077564 | 0.00835936 |
| CCR1 | 5233.67718 | 1.38824156 | 0.41307354 | 3.36076129 | 0.00077728 | 0.00837066 |
| LRRC8D | 452.287029 | 1.12343752 | 0.33443264 | 3.35923407 | 0.00078159 | 0.00841061 |
| TP53I11 | 1110.48293 | -0.9271959 | 0.27625075 | -3.3563562 | 0.00078977 | 0.00849212 |
| ANAPC1P2 | 44.2370534 | -1.8919017 | 0.5637403 | -3.3559809 | 0.00079084 | 0.00849715 |
| TBC1D2 | 432.565943 | -2.2128475 | 0.65981447 | -3.353742 | 0.00079727 | 0.00855964 |
| FAIM | 48.9020757 | -3.4314622 | 1.02340379 | -3.3529895 | 0.00079944 | 0.00857639 |
| MB21D2 | 616.463727 | 2.04298844 | 0.60958243 | 3.35145556 | 0.00080388 | 0.00861746 |
| GALNT11 | 306.10643 | -1.1174188 | 0.3335692 | -3.3498859 | 0.00080845 | 0.00865983 |
| ARHGEF3 | 525.578168 | -1.5788141 | 0.47137931 | -3.3493496 | 0.00081002 | 0.00867 |
| RPN2 | 5462.39096 | 0.82304204 | 0.24579011 | 3.34855634 | 0.00081234 | 0.00868824 |
| DNAJC7 | 652.195032 | 0.73978749 | 0.22108508 | 3.3461665 | 0.00081937 | 0.0087568 |
| MLXIPL | 42.4418257 | -3.0470115 | 0.91076012 | -3.3455697 | 0.00082114 | 0.008769 |
| ROR2 | 124.13735 | 1.80363796 | 0.53982732 | 3.34113874 | 0.00083436 | 0.00889966 |
| RNASE4 | 49.9675621 | 2.31825613 | 0.69387181 | 3.34104383 | 0.00083464 | 0.00889966 |
| INAFM2 | 489.268377 | 0.78498142 | 0.23500012 | 3.34034481 | 0.00083674 | 0.00891533 |
| TTC8 | 49.9476906 | -1.922177 | 0.57551954 | -3.3398987 | 0.00083809 | 0.00892289 |
| FAM86DP | 71.2504181 | -2.4541489 | 0.73485436 | -3.3396398 | 0.00083887 | 0.00892444 |
| UGCG | 589.690088 | -1.764297 | 0.52858702 | -3.3377607 | 0.00084456 | 0.00897821 |
| SERPINE2 | 16.9917057 | -5.3750494 | 1.61064115 | -3.337211 | 0.00084624 | 0.00898192 |
| PTPRO | 1106.72707 | 0.76923312 | 0.23050947 | 3.33709987 | 0.00084658 | 0.00898192 |
| PDP1 | 386.038686 | 1.14970364 | 0.3445305 | 3.33701552 | 0.00084683 | 0.00898192 |
| HMG20B | 952.138593 | -1.5045406 | 0.4512011 | -3.3345233 | 0.00085446 | 0.00905312 |
| DCP2 | 805.493228 | 0.96902385 | 0.2906141 | 3.33440061 | 0.00085483 | 0.00905312 |
| SLC12A9 | 466.282436 | 1.23087676 | 0.36917965 | 3.33408614 | 0.0008558 | 0.00905653 |
| TLR1 | 992.003323 | 0.93367014 | 0.2801283 | 3.33300893 | 0.00085912 | 0.00907476 |
| PSMB5 | 652.000718 | -0.8628705 | 0.25889374 | -3.3329138 | 0.00085942 | 0.00907476 |
| APBB2 | 153.497261 | -2.3946858 | 0.71849944 | -3.3328986 | 0.00085946 | 0.00907476 |
| ARHGAP22 | 191.562374 | 1.05175172 | 0.31564543 | 3.33206697 | 0.00086204 | 0.00909509 |
| ENSG000002 | 37.2225465 | -1.9422274 | 0.58296865 | -3.3316156 | 0.00086343 | 0.0091025 |
| BCAM | 30.6339086 | -2.6650798 | 0.79998255 | -3.3314224 | 0.00086403 | 0.0091025 |
| HPGD | 158.524818 | -3.9230369 | 1.17821902 | -3.329633 | 0.00086961 | 0.00915432 |
| ZNF620 | 23.3852344 | -3.3629279 | 1.01027726 | -3.3287178 | 0.00087247 | 0.00917757 |
| WARS1 | 2031.06143 | -2.0445239 | 0.61441713 | -3.3275828 | 0.00087603 | 0.00919613 |
| CDK13 | 587.291378 | 1.08460272 | 0.32596992 | 3.32730919 | 0.00087689 | 0.00919613 |
| NBEAL2 | 375.814865 | -3.383872 | 1.01710095 | -3.3269775 | 0.00087793 | 0.00919613 |
| CBFA2T3 | 57.0035171 | -2.3273123 | 0.69955337 | -3.3268545 | 0.00087832 | 0.00919613 |
| ST6GALNAC6 | 1337.15878 | -1.6646244 | 0.50040563 | -3.3265502 | 0.00087928 | 0.00919613 |
| ACAA2 | 720.341553 | -1.2811889 | 0.38514795 | -3.326485 | 0.00087949 | 0.00919613 |
| PAPLN | 47.8824815 | -3.0542792 | 0.91818292 | -3.3264387 | 0.00087963 | 0.00919613 |
| MT-ATP6 | 28326.7128 | -0.8393421 | 0.25233014 | -3.3263648 | 0.00087987 | 0.00919613 |

| | | | | | | |
|---|---|---|---|---|---|---|
| OPN3 | 373.777896 | -1.574473 | 0.47334309 | -3.3262828 | 0.00088013 | 0.00919613 |
| SALL4 | 23.3445771 | -4.0110875 | 1.20630093 | -3.3251135 | 0.00088383 | 0.00922793 |
| SIGLEC12 | 8.04846401 | 6.19217974 | 1.86260095 | 3.32448008 | 0.00088584 | 0.00924204 |
| MGAT5 | 1740.42891 | 0.92499019 | 0.27825614 | 3.32424 | 0.0008866 | 0.00924314 |
| FAM53B | 798.82945 | 0.80020902 | 0.24077171 | 3.32351759 | 0.0008889 | 0.00926024 |
| INKA2 | 241.949087 | 1.54639676 | 0.46542968 | 3.3225143 | 0.0008921 | 0.00928671 |
| CAST | 612.282241 | -1.3673118 | 0.41162173 | -3.3217677 | 0.00089449 | 0.0093047 |
| ENSG000002 | 63.8582651 | 2.58154189 | 0.77735942 | 3.32091155 | 0.00089724 | 0.00931222 |
| TMEM63B | 626.775662 | -1.1275504 | 0.33953045 | -3.3209109 | 0.00089724 | 0.00931222 |
| GASK1A | 33.2989128 | 2.65601695 | 0.7998076 | 3.32081985 | 0.00089753 | 0.00931222 |
| IER3 | 859.565819 | 1.86206456 | 0.56077386 | 3.32052666 | 0.00089848 | 0.00931222 |
| RAC1 | 4631.78459 | 0.75131786 | 0.22626576 | 3.32051056 | 0.00089853 | 0.00931222 |
| ENSG000002 | 9.62186737 | -6.9654894 | 2.0984265 | -3.3193869 | 0.00090215 | 0.00934288 |
| IQCD | 14.0837706 | 5.5507064 | 1.6723226 | 3.31916008 | 0.00090289 | 0.00934358 |
| GALM | 717.066443 | -1.6891375 | 0.50911423 | -3.3177966 | 0.00090731 | 0.00937916 |
| SSR4 | 1195.04971 | 0.8783708 | 0.26476061 | 3.31760374 | 0.00090793 | 0.00937916 |
| WNK1 | 2906.46778 | 0.72666726 | 0.21904183 | 3.31748169 | 0.00090833 | 0.00937916 |
| FDFT1 | 1487.36382 | -0.7011926 | 0.21139769 | -3.3169361 | 0.0009101 | 0.00939059 |
| ANGPT4 | 25.759295 | -4.9629672 | 1.4967603 | -3.3158063 | 0.00091379 | 0.00942171 |
| APOBEC3G | 50.5503191 | -1.7738857 | 0.5352262 | -3.314273 | 0.00091882 | 0.00946658 |
| SH3BP1 | 1546.83857 | 0.91909723 | 0.27736908 | 3.31362538 | 0.00092095 | 0.00948158 |
| NCAPH | 287.333253 | -2.5776232 | 0.77797957 | -3.3132273 | 0.00092226 | 0.00948813 |
| CHD3 | 391.782476 | 1.83612957 | 0.55425758 | 3.31277302 | 0.00092376 | 0.0094966 |
| RN7SL2 | 219.460923 | 2.30052212 | 0.69455487 | 3.31222515 | 0.00092557 | 0.00950825 |
| SORBS1 | 33.4332227 | -3.2432007 | 0.97966931 | -3.3105055 | 0.00093128 | 0.00955988 |
| ADRB1 | 9.1783974 | 6.38117472 | 1.92800895 | 3.30972256 | 0.00093388 | 0.00957965 |
| SYNGR2 | 5458.29035 | 1.2020809 | 0.36365505 | 3.3055526 | 0.00094789 | 0.00971625 |
| EPB41L2 | 923.749626 | -1.7194831 | 0.52053853 | -3.3032772 | 0.00095562 | 0.0097883 |
| GAS1 | 41.6756363 | 3.21486585 | 0.97335158 | 3.30288246 | 0.00095696 | 0.00979494 |
| SLC44A2 | 997.68287 | -1.4609009 | 0.44233953 | -3.3026686 | 0.0009577 | 0.00979527 |
| ANXA4 | 906.245223 | -1.0114821 | 0.30643013 | -3.3008573 | 0.0009639 | 0.00985155 |
| RBPMS2 | 63.8716425 | -4.4379444 | 1.34519984 | -3.2990968 | 0.00096996 | 0.00990634 |
| ZHX3 | 99.7867151 | -1.1256777 | 0.34154986 | -3.295793 | 0.00098144 | 0.01001628 |
| ESD | 436.296814 | -0.7540954 | 0.22890278 | -3.2943916 | 0.00098635 | 0.01005905 |
| RMRP | 78.8362954 | 2.66634248 | 0.80944397 | 3.29404207 | 0.00098758 | 0.01006366 |
| MAPRE2 | 910.647243 | -1.4120473 | 0.42869143 | -3.2938547 | 0.00098824 | 0.01006366 |
| HEATR5B | 222.002132 | -0.9656177 | 0.29318261 | -3.2935707 | 0.00098923 | 0.01006654 |
| LYSMD3 | 244.492168 | 1.17502713 | 0.35687201 | 3.29257297 | 0.00099275 | 0.01009501 |
| PLAUR | 4718.91241 | 1.57343892 | 0.47814192 | 3.29073616 | 0.00099926 | 0.0101538 |
| BCOR | 548.506204 | 1.48002667 | 0.45004389 | 3.2886274 | 0.00100677 | 0.01022278 |
| SLC29A2 | 29.3636078 | -2.1895232 | 0.66590405 | -3.2880461 | 0.00100885 | 0.01023651 |
| ZFP90 | 109.644602 | -1.3748696 | 0.41834039 | -3.2864854 | 0.00101446 | 0.01028073 |
| CRISPLD1 | 21.4463171 | -4.1484551 | 1.26233521 | -3.286334 | 0.00101501 | 0.01028073 |
| LMLN | 75.0826077 | -1.3992532 | 0.42579384 | -3.2862224 | 0.00101541 | 0.01028073 |
| RPLP0P2 | 16.4916431 | -5.8115835 | 1.76940529 | -3.2844841 | 0.00102169 | 0.01033691 |
| SFMBT2 | 729.898487 | 1.34022611 | 0.40811189 | 3.28396732 | 0.00102357 | 0.01034843 |
| SYTL4 | 15.4229411 | -3.2967064 | 1.00409039 | -3.2832765 | 0.00102608 | 0.01036636 |
| PCED1B-AS1 | 219.913727 | 2.05680754 | 0.62657121 | 3.2826397 | 0.0010284 | 0.01038233 |

| | | | | | | |
|---|---|---|---|---|---|---|
| GNGT2 | 177.00874 | 1.65290425 | 0.50356213 | 3.28242363 | 0.00102919 | 0.01038283 |
| CAMK1 | 1875.75933 | 1.06482261 | 0.32452511 | 3.28117176 | 0.00103377 | 0.01042154 |
| TTLL4 | 582.066931 | -1.5468112 | 0.4719913 | -3.2772027 | 0.00104841 | 0.01054526 |
| MAPK8IP1 | 17.078013 | -4.8587966 | 1.48260697 | -3.277198 | 0.00104843 | 0.01054526 |
| TMEM41B | 446.799751 | 1.25424227 | 0.38271927 | 3.27718612 | 0.00104847 | 0.01054526 |
| SOCS5 | 208.365307 | 0.88527989 | 0.27015725 | 3.27690589 | 0.00104951 | 0.01054526 |
| CASP7 | 205.265553 | -1.332487 | 0.40663902 | -3.2768302 | 0.00104979 | 0.01054526 |
| RN7SL1 | 262.222905 | 2.05410995 | 0.62690737 | 3.2765765 | 0.00105074 | 0.01054719 |
| COL6A3 | 50.6295341 | -2.8735388 | 0.87706761 | -3.2763025 | 0.00105176 | 0.01054831 |
| PLEK | 6275.103 | -1.1934176 | 0.36427518 | -3.276143 | 0.00105235 | 0.01054831 |
| SMAD5 | 436.604324 | 0.93352103 | 0.2850855 | 3.27453001 | 0.00105838 | 0.01060115 |
| TREML2 | 13.253107 | 4.46008179 | 1.36299519 | 3.27226523 | 0.00106689 | 0.01067883 |
| VARS2 | 164.310897 | -1.3832983 | 0.42276862 | -3.2719985 | 0.0010679 | 0.0106813 |
| ZCCHC4 | 92.1807698 | -1.1951858 | 0.36535114 | -3.2713345 | 0.00107041 | 0.01069879 |
| RASL11B | 24.0211128 | -4.7083899 | 1.44021488 | -3.2692274 | 0.00107842 | 0.01077112 |
| SAT1 | 6380.88136 | 2.00177178 | 0.61238331 | 3.26882157 | 0.00107996 | 0.01077891 |
| ENSG000002 | 29.1437748 | -4.6614819 | 1.42659893 | -3.267549 | 0.00108483 | 0.0108198 |
| SINHCAF | 394.396952 | 0.91532401 | 0.28020872 | 3.26657933 | 0.00108855 | 0.01084922 |
| FAM200B | 203.220114 | 1.10935946 | 0.33969701 | 3.26573216 | 0.00109181 | 0.01087401 |
| CLIC6 | 12.1733667 | -6.3204154 | 1.93603483 | -3.2646186 | 0.00109612 | 0.01090911 |
| IQGAP2 | 1092.31268 | -1.0372103 | 0.31803772 | -3.2612808 | 0.0011091 | 0.01103053 |
| AMDHD1 | 39.0285258 | -3.0026114 | 0.92119143 | -3.2594869 | 0.00111614 | 0.01109267 |
| DLC1 | 65.637263 | -2.1888635 | 0.67162079 | -3.2590764 | 0.00111776 | 0.01110088 |
| ENSG000002 | 10.5961361 | -3.7766741 | 1.15981929 | -3.2562608 | 0.0011289 | 0.01120364 |
| MAPKBP1 | 158.841596 | -1.2605065 | 0.3872511 | -3.2550108 | 0.00113388 | 0.01124512 |
| ABAT | 79.1225763 | 1.40351512 | 0.43125577 | 3.25448429 | 0.00113598 | 0.01125803 |
| P2RY6 | 980.080334 | 2.6005632 | 0.79957345 | 3.25243816 | 0.00114419 | 0.0113314 |
| CSK | 1943.32849 | 0.85992046 | 0.26441963 | 3.25210517 | 0.00114554 | 0.01133669 |
| SLC29A3 | 860.943761 | -2.2197542 | 0.68370348 | -3.2466621 | 0.00116767 | 0.01154759 |
| ICOSLG | 831.727327 | 1.67116791 | 0.51496708 | 3.24519369 | 0.00117371 | 0.011591 |
| NECTIN2 | 2188.80651 | 0.93740396 | 0.28885922 | 3.24519319 | 0.00117371 | 0.011591 |
| SELPLG | 383.842205 | 1.26089158 | 0.38862606 | 3.24448539 | 0.00117663 | 0.01159838 |
| PLPPR3 | 13.1169551 | 4.38101359 | 1.35030299 | 3.24446706 | 0.00117671 | 0.01159838 |
| MTHFR | 488.878294 | -1.0603997 | 0.32683884 | -3.2444117 | 0.00117694 | 0.01159838 |
| AHNAK2 | 52.9818124 | -3.9644999 | 1.22339877 | -3.2405623 | 0.00119294 | 0.01174789 |
| SYK | 2050.42313 | 0.80667953 | 0.24913458 | 3.23792681 | 0.00120402 | 0.01184864 |
| ENSG000002 | 110.809373 | -4.2025046 | 1.29801841 | -3.237631 | 0.00120527 | 0.01185263 |
| PRPS1 | 684.314648 | -1.6257985 | 0.50222171 | -3.2372128 | 0.00120703 | 0.01186171 |
| C4orf19 | 7.61604489 | 6.11593615 | 1.89029406 | 3.23544166 | 0.00121455 | 0.01192719 |
| GNB4 | 1612.40455 | 0.68116386 | 0.21056708 | 3.23490205 | 0.00121684 | 0.0119414 |
| CPLX2 | 10.1927263 | 6.53948999 | 2.02170657 | 3.23463854 | 0.00121797 | 0.01194408 |
| MAPK10 | 11.9016336 | -4.8333109 | 1.49469174 | -3.2336506 | 0.00122219 | 0.01197711 |
| FABP3 | 756.011495 | -1.3153276 | 0.40682031 | -3.2331905 | 0.00122416 | 0.01198216 |
| ABCD4 | 309.126718 | -1.518766 | 0.46975075 | -3.2331317 | 0.00122441 | 0.01198216 |
| TDRD9 | 92.4993465 | -3.9099508 | 1.20942089 | -3.2329116 | 0.00122535 | 0.01198305 |
| NDUFV1 | 922.254443 | -0.7683379 | 0.23782 | -3.230754 | 0.00123464 | 0.01206546 |
| LINC02207 | 14.6625283 | 4.54507354 | 1.40857105 | 3.2267265 | 0.00125215 | 0.01221697 |
| TET2 | 265.289777 | 1.24186956 | 0.38487059 | 3.22671983 | 0.00125218 | 0.01221697 |

| | | | | | | |
|---|---|---|---|---|---|---|
| PPARA | 287.451408 | -1.2389068 | 0.38396804 | -3.2265884 | 0.00125276 | 0.01221697 |
| ZNF768 | 356.392469 | 0.77439751 | 0.24004935 | 3.225993 | 0.00125536 | 0.01223392 |
| ANGPTL2 | 10.2506171 | -6.0618528 | 1.88051291 | -3.2235103 | 0.0012663 | 0.01233191 |
| B2M | 27789.5519 | 0.69428831 | 0.21542759 | 3.22283843 | 0.00126927 | 0.01235231 |
| GAL | 76.1090762 | -3.7789269 | 1.17272407 | -3.2223496 | 0.00127144 | 0.01236483 |
| MAPK8IP3 | 41.9877108 | -1.5923715 | 0.49422456 | -3.2219596 | 0.00127317 | 0.01237311 |
| VARS1 | 1115.97818 | -1.4228575 | 0.44188902 | -3.2199431 | 0.00128216 | 0.01245185 |
| RPL23AP7 | 197.424881 | -0.9230062 | 0.28680724 | -3.218211 | 0.00128993 | 0.01251864 |
| RRBP1 | 925.823492 | -1.1682661 | 0.36317681 | -3.2167972 | 0.0012963 | 0.0125718 |
| HADH | 501.834043 | -1.2957367 | 0.40285806 | -3.2163604 | 0.00129828 | 0.01258226 |
| NINJ2 | 180.800628 | 1.16499462 | 0.36225152 | 3.21598265 | 0.00129999 | 0.0125885 |
| LDLRAD3 | 157.698677 | 1.617464 | 0.50297056 | 3.21582238 | 0.00130071 | 0.0125885 |
| ENSG000002 | 7.43667122 | 6.07766987 | 1.89009248 | 3.215541 | 0.00130199 | 0.01259217 |
| MSI2 | 165.56183 | 1.51809366 | 0.47221974 | 3.2148035 | 0.00130534 | 0.01260574 |
| RTKN | 35.4121634 | -2.1975493 | 0.68359745 | -3.2146834 | 0.00130588 | 0.01260574 |
| MTATP6P1 | 4617.55181 | -0.8839967 | 0.27499094 | -3.2146394 | 0.00130609 | 0.01260574 |
| SNAPC3 | 165.911647 | -1.5148665 | 0.47127122 | -3.214426 | 0.00130706 | 0.01260645 |
| MLLT11 | 90.2599808 | -2.7979632 | 0.87211931 | -3.2082345 | 0.00133553 | 0.01287219 |
| GALNT18 | 21.6487462 | 2.54871798 | 0.79463199 | 3.20741929 | 0.00133932 | 0.01288106 |
| MAP3K2 | 982.687581 | 0.86592497 | 0.26997633 | 3.20741073 | 0.00133936 | 0.01288106 |
| SDC4 | 787.571561 | -1.1414938 | 0.35590733 | -3.2072781 | 0.00133997 | 0.01288106 |
| ALG2 | 679.021716 | 1.06685007 | 0.33263723 | 3.20724799 | 0.00134011 | 0.01288106 |
| SLC16A1-AS1 | 51.7072378 | 2.13144277 | 0.66496887 | 3.20532717 | 0.00134909 | 0.01295847 |
| MAP3K5-AS2 | 551.525632 | -1.1705612 | 0.36529327 | -3.2044422 | 0.00135324 | 0.01298948 |
| CEBPD | 452.132479 | 2.26598455 | 0.70737038 | 3.2033919 | 0.00135819 | 0.01302805 |
| SOAT1 | 1475.08525 | -2.0025535 | 0.62527698 | -3.2026663 | 0.00136162 | 0.013052 |
| LIMD1 | 804.150512 | 1.3164773 | 0.41108555 | 3.20244121 | 0.00136268 | 0.0130533 |
| NHS | 22.1293857 | 3.54921575 | 1.10846884 | 3.20190846 | 0.0013652 | 0.01306855 |
| NAGK | 1083.58846 | -0.9058513 | 0.28304394 | -3.200391 | 0.00137241 | 0.0131286 |
| CENPE | 15.7660271 | -3.463845 | 1.08264915 | -3.199416 | 0.00137706 | 0.01316412 |
| HIPK2 | 1597.37678 | 0.90828708 | 0.28410528 | 3.19700875 | 0.00138861 | 0.01326545 |
| ARHGAP9 | 601.169767 | 1.18944143 | 0.37223281 | 3.19542339 | 0.00139626 | 0.01332947 |
| FOXP4 | 329.970576 | 1.97682876 | 0.61881388 | 3.19454497 | 0.00140051 | 0.01336102 |
| RAPGEF3 | 79.3478991 | -3.2645297 | 1.0219881 | -3.1942932 | 0.00140174 | 0.0133636 |
| RNASE3 | 7.99156346 | 6.18598329 | 1.93716963 | 3.19331007 | 0.00140652 | 0.01340009 |
| SLC25A44 | 637.571139 | 0.76277245 | 0.2389535 | 3.19213758 | 0.00141224 | 0.01344549 |
| MYO5A | 978.28713 | 0.95689203 | 0.29980292 | 3.19173682 | 0.0014142 | 0.01345503 |
| FRRS1 | 197.999757 | -2.4264715 | 0.76055829 | -3.1903821 | 0.00142085 | 0.01350912 |
| GCNT1 | 1668.11532 | 2.70751128 | 0.84966577 | 3.18656037 | 0.00143975 | 0.01367961 |
| MRGPRF | 12.0633908 | -7.292125 | 2.29006257 | -3.1842471 | 0.00145131 | 0.01378009 |
| DENND11 | 452.674013 | 1.51361052 | 0.47568154 | 3.18198286 | 0.0014627 | 0.01387036 |
| MIR124-1HG | 6.88671478 | 5.96913843 | 1.87592811 | 3.18196545 | 0.00146279 | 0.01387036 |
| ASPHD1 | 112.385816 | -3.0027024 | 0.94397025 | -3.1809291 | 0.00146804 | 0.01391068 |
| APOBEC3C | 720.801759 | -1.1330952 | 0.35668109 | -3.1767741 | 0.00148923 | 0.01410199 |
| LCP1 | 19237.336 | -1.8393854 | 0.5794522 | -3.1743523 | 0.00150171 | 0.01421062 |
| ACTR2 | 9170.20917 | 0.62277157 | 0.19634662 | 3.17179677 | 0.00151499 | 0.0143266 |
| ARHGEF17 | 29.1961954 | -2.3010966 | 0.72558665 | -3.1713602 | 0.00151727 | 0.0143385 |
| DARS1 | 468.93728 | -1.003404 | 0.31644919 | -3.1708219 | 0.00152008 | 0.01435543 |

| | | | | | | |
|---|---|---|---|---|---|---|
| PEMT | 72.8761905 | -1.5131904 | 0.47740852 | -3.1695924 | 0.00152653 | 0.01440662 |
| MRAS | 1277.02338 | -1.5052829 | 0.47501694 | -3.1689035 | 0.00153015 | 0.01442069 |
| VEGFA | 213.459825 | 2.99373779 | 0.94473883 | 3.16885226 | 0.00153042 | 0.01442069 |
| PEPD | 2203.93544 | -0.8404536 | 0.26523415 | -3.1687232 | 0.0015311 | 0.01442069 |
| ZNF320 | 61.7903943 | -1.4891967 | 0.4700509 | -3.1681605 | 0.00153407 | 0.01443016 |
| SIDT2 | 455.273004 | 1.5480047 | 0.48862564 | 3.16807912 | 0.0015345 | 0.01443016 |
| YTHDC2 | 337.793816 | 1.12608824 | 0.35546295 | 3.16794824 | 0.00153519 | 0.01443016 |
| HDAC1 | 662.978197 | 0.75979358 | 0.23989571 | 3.16718288 | 0.00153923 | 0.01445602 |
| TPCN1 | 2463.39402 | 1.14928206 | 0.36288856 | 3.16703859 | 0.00154 | 0.01445602 |
| ZMYM2 | 432.997974 | 1.14249718 | 0.36100633 | 3.1647566 | 0.00155213 | 0.01456014 |
| GALE | 431.090003 | -0.8935722 | 0.28239718 | -3.1642392 | 0.00155489 | 0.01457631 |
| DUSP3 | 1786.3618 | -0.746484 | 0.23598516 | -3.1632669 | 0.00156009 | 0.01461533 |
| TADA2B | 439.569974 | 0.76453611 | 0.24181293 | 3.16168411 | 0.0015686 | 0.0146852 |
| OAS2 | 105.070072 | -2.1354631 | 0.67655288 | -3.1563876 | 0.00159736 | 0.01494456 |
| MEFV | 14.1284206 | 4.56126486 | 1.44573684 | 3.15497589 | 0.00160511 | 0.01500707 |
| KCNK13 | 989.976305 | 0.90992151 | 0.28859463 | 3.15293987 | 0.00161635 | 0.01508288 |
| LINC01857 | 18.4570809 | -2.882055 | 0.9141169 | -3.1528298 | 0.00161696 | 0.01508288 |
| GRIN3A | 29.1735388 | 3.14397226 | 0.99719654 | 3.15281103 | 0.00161706 | 0.01508288 |
| IFRD2 | 265.914766 | -1.107557 | 0.35130105 | -3.1527291 | 0.00161752 | 0.01508288 |
| APOL2 | 271.471251 | -2.2809317 | 0.7237635 | -3.1514875 | 0.00162441 | 0.01513709 |
| PDLIM1 | 24.3065634 | 3.19964591 | 1.01549929 | 3.15081059 | 0.00162818 | 0.01516216 |
| SLC25A4 | 82.2149426 | -1.6444884 | 0.52206989 | -3.1499392 | 0.00163304 | 0.01519738 |
| PLEKHF1 | 106.422396 | -1.5967397 | 0.50694758 | -3.1497137 | 0.00163431 | 0.01519904 |
| MAGEH1 | 74.4873361 | -1.2375079 | 0.39313282 | -3.1478111 | 0.00164498 | 0.01528818 |
| GRB10 | 116.755684 | 1.44351903 | 0.45896346 | 3.145172 | 0.00165989 | 0.01541658 |
| NCEH1 | 3278.0984 | -1.901851 | 0.60486253 | -3.1442698 | 0.00166502 | 0.01544811 |
| SLC49A3 | 226.716117 | -2.09843 | 0.66739979 | -3.1441874 | 0.00166549 | 0.01544811 |
| MDN1 | 204.537102 | -1.1476661 | 0.36520454 | -3.1425297 | 0.00167495 | 0.01552211 |
| LGALS3 | 2803.1638 | -2.0467898 | 0.65134557 | -3.1424023 | 0.00167568 | 0.01552211 |
| P2RX4 | 1973.88414 | -0.7919541 | 0.25207054 | -3.1417956 | 0.00167915 | 0.01554406 |
| PCCA | 131.548101 | -1.6071501 | 0.51173881 | -3.1405672 | 0.00168621 | 0.01559911 |
| SMOX | 2012.81352 | 2.49363707 | 0.79417823 | 3.13989602 | 0.00169008 | 0.0156246 |
| DALRD3 | 98.2708993 | -1.5122975 | 0.48166942 | -3.1397 | 0.00169121 | 0.01562477 |
| UBE2H | 833.614731 | 0.86826519 | 0.27658239 | 3.13926418 | 0.00169373 | 0.01563774 |
| RDH10 | 107.739808 | -1.6186057 | 0.51565261 | -3.138946 | 0.00169557 | 0.01564444 |
| LYST | 208.699247 | -1.970902 | 0.62818726 | -3.1374434 | 0.00170428 | 0.01571453 |
| XPR1 | 930.718674 | -1.3723114 | 0.43773253 | -3.1350455 | 0.00171827 | 0.01583316 |
| C1QTNF5 | 15.855341 | -3.2981703 | 1.05257091 | -3.1334424 | 0.00172769 | 0.01590946 |
| AKAP8L | 267.015734 | 1.00778872 | 0.32166463 | 3.13304178 | 0.00173005 | 0.01592075 |
| LMTK2 | 113.946141 | -1.1738327 | 0.37471512 | -3.1326002 | 0.00173265 | 0.01593428 |
| HGSNAT | 2079.35913 | 0.74007832 | 0.23638152 | 3.13086376 | 0.00174293 | 0.01600818 |
| ZNF124 | 124.861871 | 1.1230522 | 0.35870437 | 3.1308573 | 0.00174297 | 0.01600818 |
| ENSG000002 | 6.60550333 | -6.4257603 | 2.05349597 | -3.1291808 | 0.00175294 | 0.01608928 |
| KCNQ3 | 126.719561 | 2.98752071 | 0.95499161 | 3.12832143 | 0.00175808 | 0.01612586 |
| DTNB | 187.280749 | -1.2824108 | 0.40997404 | -3.1280292 | 0.00175983 | 0.01613136 |
| DGAT2 | 62.4193678 | -3.5910389 | 1.14898794 | -3.1253931 | 0.00177568 | 0.01625849 |
| DOCK6 | 11.8036989 | -5.3414774 | 1.70908813 | -3.1253376 | 0.00177601 | 0.01625849 |
| CCL22 | 661.66209 | -3.1901323 | 1.02109104 | -3.1242389 | 0.00178266 | 0.01629942 |

| | | | | | | |
|---|---|---|---|---|---|---|
| SASH3 | 1854.88645 | 1.05000725 | 0.33608678 | 3.12421468 | 0.0017828 | 0.01629942 |
| CTSS | 5745.22967 | 1.07512199 | 0.34427677 | 3.12284212 | 0.00179114 | 0.01636496 |
| ANXA11 | 3478.79724 | -1.4081014 | 0.45099939 | -3.1221804 | 0.00179517 | 0.01639113 |
| ENSG000002 | 19.5469641 | 2.70140809 | 0.86532936 | 3.12182644 | 0.00179733 | 0.01640019 |
| CREM | 865.164285 | 1.59048742 | 0.5095618 | 3.12128462 | 0.00180064 | 0.01641972 |
| ITGB3 | 520.162352 | -1.7935041 | 0.5747926 | -3.120263 | 0.0018069 | 0.01646609 |
| CYBA | 5853.22734 | 0.69103753 | 0.22157335 | 3.11877539 | 0.00181604 | 0.0165289 |
| CCDC74A | 13.1900469 | -3.366143 | 1.07932124 | -3.1187592 | 0.00181614 | 0.0165289 |
| ATP6V1H | 996.854652 | -1.8539685 | 0.59498303 | -3.1160024 | 0.00183321 | 0.0166734 |
| CARD19 | 800.033911 | 0.70090402 | 0.22497223 | 3.11551352 | 0.00183625 | 0.01668578 |
| SRC | 2093.25476 | 1.10970614 | 0.35620001 | 3.11540179 | 0.00183695 | 0.01668578 |
| MEF2D | 1301.45472 | 0.88961652 | 0.28566395 | 3.11420642 | 0.0018444 | 0.01674271 |
| KYAT3 | 184.948372 | -1.0007803 | 0.3214598 | -3.1132363 | 0.00185048 | 0.01678657 |
| ENSG000002 | 7.26013122 | -5.554114 | 1.78413722 | -3.1130531 | 0.00185163 | 0.01678657 |
| TMPO | 1109.78363 | 1.06187941 | 0.34115086 | 3.11263884 | 0.00185423 | 0.0167993 |
| RSPO4 | 10.2849296 | -6.0693361 | 1.95084285 | -3.1111353 | 0.0018637 | 0.01687419 |
| SYP | 28.2203677 | -2.7833038 | 0.89497974 | -3.1099071 | 0.00187146 | 0.0169336 |
| ZC3H18 | 458.862513 | 0.93389559 | 0.300359 | 3.10926455 | 0.00187554 | 0.01695954 |
| FPR3 | 18.6086408 | -6.01826 | 1.93653465 | -3.1077471 | 0.00188519 | 0.01703589 |
| MRPS27 | 475.567341 | -1.0585961 | 0.34069374 | -3.1071781 | 0.00188883 | 0.01705773 |
| VAMP3 | 1056.64619 | -1.1246824 | 0.36202583 | -3.1066357 | 0.00189229 | 0.01707808 |
| TGFB1I1 | 17.3222657 | -3.0364279 | 0.97771411 | -3.1056398 | 0.00189868 | 0.01712469 |
| FCGR3B | 19.4206463 | 3.10271588 | 0.99914766 | 3.1053627 | 0.00190046 | 0.01712974 |
| ANKRD35 | 19.291362 | 2.37308179 | 0.76427825 | 3.10499715 | 0.00190281 | 0.01713287 |
| PEX10 | 190.368139 | -1.0259862 | 0.33044012 | -3.1049082 | 0.00190338 | 0.01713287 |
| SMIM29 | 263.989032 | -0.8694952 | 0.28005414 | -3.1047396 | 0.00190447 | 0.01713287 |
| PYGO1 | 36.2602672 | -3.6630112 | 1.18093933 | -3.1017776 | 0.00192362 | 0.01729414 |
| IVNS1ABP | 1506.84184 | -0.8877461 | 0.28622737 | -3.1015417 | 0.00192516 | 0.01729686 |
| CCNC | 378.246009 | 0.71344306 | 0.23010374 | 3.1005279 | 0.00193176 | 0.01734508 |
| DYSF | 599.548576 | -3.0291395 | 0.97730863 | -3.0994708 | 0.00193867 | 0.01739598 |
| CDH6 | 45.3315302 | 5.61926739 | 1.81315953 | 3.09915775 | 0.00194072 | 0.01740325 |
| KCNN1 | 28.7811988 | 2.36261852 | 0.76262004 | 3.09802839 | 0.00194813 | 0.01745856 |
| RMDN1 | 196.848571 | -1.0748101 | 0.3469606 | -3.0977872 | 0.00194971 | 0.01746163 |
| ZNF296 | 44.3677037 | 1.69860082 | 0.54864805 | 3.0959753 | 0.00196167 | 0.01755748 |
| APOBEC3D | 10.4535803 | -5.2379636 | 1.69239721 | -3.0949966 | 0.00196815 | 0.0176043 |
| HAAO | 44.2888432 | -2.5180216 | 0.81399616 | -3.0934072 | 0.00197872 | 0.0176876 |
| PITPNA | 2028.69518 | 0.82006509 | 0.26517413 | 3.09255317 | 0.00198443 | 0.01772729 |
| BNIP2 | 841.09258 | 0.87969463 | 0.28448414 | 3.0922449 | 0.00198649 | 0.0177319 |
| BIRC7 | 6.14901107 | -6.3213509 | 2.04435618 | -3.0920986 | 0.00198747 | 0.0177319 |
| ANKRD44 | 406.248399 | 1.17050769 | 0.37863115 | 3.09141945 | 0.00199202 | 0.01776123 |
| DAAM1 | 111.158198 | -1.7940118 | 0.58068079 | -3.0894974 | 0.00200495 | 0.01786521 |
| FCN1 | 35.6220999 | 3.50641873 | 1.13502447 | 3.08928912 | 0.00200636 | 0.01786641 |
| MYO1A | 32.5463515 | 2.89621491 | 0.93808787 | 3.08735993 | 0.00201943 | 0.01797138 |
| CBR3 | 11.4990202 | -3.6302966 | 1.1759399 | -3.0871447 | 0.00202089 | 0.01797301 |
| POLR2A | 1355.47092 | -0.6357031 | 0.20617621 | -3.0833001 | 0.00204719 | 0.01819534 |
| SLC44A5 | 6.89662628 | -5.4857911 | 1.77977215 | -3.0822997 | 0.00205408 | 0.01823607 |
| SLC25A1 | 695.143293 | -0.9912484 | 0.32159812 | -3.0822581 | 0.00205437 | 0.01823607 |
| JMJD6 | 202.807175 | 0.97313978 | 0.31588105 | 3.08071592 | 0.00206504 | 0.0183192 |

| | | | | | | |
|---|---|---|---|---|---|---|
| MFGE8 | 362.032484 | -3.0007012 | 0.97437763 | -3.0796081 | 0.00207273 | 0.01837587 |
| LYN | 4417.41172 | 1.12457573 | 0.36521852 | 3.07918597 | 0.00207567 | 0.01839031 |
| PILRB | 12.4133338 | 3.70817184 | 1.20564211 | 3.07568208 | 0.00210022 | 0.01859605 |
| COQ8A | 142.979625 | -1.2053998 | 0.39200279 | -3.0749775 | 0.00210518 | 0.0186283 |
| IFI6 | 755.580421 | -2.1697526 | 0.70570039 | -3.0746088 | 0.00210779 | 0.01863959 |
| MAP1S | 1045.91468 | 0.68045114 | 0.22140645 | 3.07331213 | 0.00211697 | 0.018709 |
| TFE3 | 1865.37398 | 0.67453868 | 0.21950364 | 3.07301822 | 0.00211906 | 0.01871566 |
| ENSG000002 | 14.8078066 | 3.60258854 | 1.17247753 | 3.07262907 | 0.00212182 | 0.0187283 |
| NAGS | 17.85523 | 2.40766224 | 0.78376345 | 3.07192463 | 0.00212683 | 0.01875268 |
| KICS2 | 59.8207163 | -1.4159459 | 0.46094003 | -3.0718657 | 0.00212725 | 0.01875268 |
| DDAH1 | 10.2082372 | -5.1622866 | 1.68096256 | -3.0710301 | 0.00213322 | 0.01879345 |
| TMEM9 | 201.566864 | 0.88209139 | 0.28727504 | 3.07054658 | 0.00213667 | 0.0188121 |
| CDKN2A | 154.790503 | -1.6324567 | 0.53175846 | -3.0699214 | 0.00214115 | 0.01883972 |
| PLK2 | 23.4287981 | 2.29006642 | 0.74650866 | 3.06770242 | 0.00215711 | 0.01896667 |
| ARHGAP27 | 531.24044 | 1.17228866 | 0.38215914 | 3.06754059 | 0.00215828 | 0.01896667 |
| RAN | 2994.10822 | -0.6405397 | 0.20883856 | -3.0671525 | 0.00216109 | 0.01897944 |
| ZFHX2 | 57.9543801 | -1.2973017 | 0.42311451 | -3.066077 | 0.00216887 | 0.01903594 |
| HEXA | 4431.07494 | 0.67718753 | 0.22088979 | 3.06572586 | 0.00217142 | 0.0190464 |
| PTPN9 | 1527.78346 | 0.73865113 | 0.24108308 | 3.06388619 | 0.00218482 | 0.01914657 |
| TNRC18 | 891.871141 | 0.68920891 | 0.22495831 | 3.06371838 | 0.00218605 | 0.01914657 |
| ADCY3 | 1789.59999 | -2.9078046 | 0.9491651 | -3.0635393 | 0.00218735 | 0.01914657 |
| ANK2 | 38.7786581 | -2.531439 | 0.82634665 | -3.0634105 | 0.0021883 | 0.01914657 |
| MTARC1 | 13.4218568 | -4.9548572 | 1.6177038 | -3.0628952 | 0.00219207 | 0.01916763 |
| TPRG1 | 25.1243006 | -4.0147669 | 1.31107584 | -3.0621928 | 0.00219722 | 0.01920071 |
| GAPVD1 | 782.327318 | 0.66179313 | 0.21615596 | 3.06164637 | 0.00220123 | 0.01922383 |
| SEMA4D | 117.853414 | 2.67208377 | 0.87342387 | 3.05932076 | 0.00221839 | 0.01936167 |
| FOXN2 | 343.066601 | 1.19643849 | 0.39111235 | 3.05906602 | 0.00222028 | 0.0193661 |
| DNAAF5 | 275.916711 | -1.7412949 | 0.56981831 | -3.0558775 | 0.00224403 | 0.01956108 |
| ENSG000002 | 13.3124351 | 3.86819074 | 1.26593192 | 3.05560724 | 0.00224605 | 0.01956657 |
| JUND | 1545.09995 | 0.80562368 | 0.26369543 | 3.05512947 | 0.00224963 | 0.01958562 |
| NRP2 | 2044.52864 | -1.1432223 | 0.37453246 | -3.0523985 | 0.0022702 | 0.01975247 |
| CPEB3 | 73.8377711 | 1.34914175 | 0.44216459 | 3.05122067 | 0.00227913 | 0.01981785 |
| SNAI3 | 236.544014 | 1.83884745 | 0.60270515 | 3.0509901 | 0.00228088 | 0.0198208 |
| POPDC3 | 26.3356809 | -3.2639623 | 1.07013643 | -3.0500431 | 0.00228808 | 0.0198711 |
| B9D1 | 10.0284874 | -5.1760922 | 1.69719391 | -3.0497943 | 0.00228998 | 0.0198746 |
| MAX | 1531.0598 | 0.92548089 | 0.30347426 | 3.04961903 | 0.00229132 | 0.0198746 |
| TNFSF14 | 114.818427 | -3.4379694 | 1.12745051 | -3.0493306 | 0.00229352 | 0.01987777 |
| INPP4B | 165.429341 | 1.56224361 | 0.51234536 | 3.04920026 | 0.00229451 | 0.01987777 |
| ATP11A | 1132.23933 | 0.77885756 | 0.25554607 | 3.04781663 | 0.00230511 | 0.01995721 |
| RARRES1 | 12.6906563 | -4.4135578 | 1.44939422 | -3.0451051 | 0.00232599 | 0.02012562 |
| ENTPD1 | 1858.9836 | 0.80964367 | 0.26590894 | 3.0448155 | 0.00232823 | 0.0201326 |
| PAG1 | 680.560345 | 1.35524723 | 0.44522728 | 3.04394468 | 0.00233498 | 0.02016199 |
| MOCS1 | 36.3091167 | -1.8815734 | 0.61815999 | -3.0438291 | 0.00233588 | 0.02016199 |
| SP110 | 166.080194 | -1.3376947 | 0.43949236 | -3.043727 | 0.00233667 | 0.02016199 |
| ENSG000002 | 15.9941649 | -3.2891795 | 1.08067416 | -3.0436366 | 0.00233737 | 0.02016199 |
| ENSG000002 | 13.7686998 | 3.8976029 | 1.28148781 | 3.04146701 | 0.00235428 | 0.0202954 |
| CD52 | 2035.1959 | -4.7220676 | 1.55299755 | -3.040615 | 0.00236096 | 0.02034043 |
| PDCD4 | 221.236716 | -1.0787569 | 0.35480521 | -3.04042 | 0.00236248 | 0.02034112 |

| | | | | | | |
|---|---|---|---|---|---|---|
| TMEM167A | 1158.62171 | 0.69929437 | 0.23014778 | 3.03845798 | 0.00237792 | 0.0204615 |
| DHDDS | 448.678912 | -0.7017792 | 0.23101156 | -3.0378533 | 0.0023827 | 0.02048319 |
| CA11 | 47.0032772 | 1.82495137 | 0.60075372 | 3.0377696 | 0.00238336 | 0.02048319 |
| ARID3A | 1096.28965 | -1.5256944 | 0.50232411 | -3.0372709 | 0.00238731 | 0.02050456 |
| ZNF217 | 861.056861 | 1.43819048 | 0.47371057 | 3.03601094 | 0.00239731 | 0.02057786 |
| ERRFI1 | 166.414291 | 2.32558943 | 0.76618268 | 3.03529366 | 0.00240302 | 0.02061427 |
| OGDH | 2023.89522 | -0.7434317 | 0.24497409 | -3.0347358 | 0.00240746 | 0.02063982 |
| SDF2L1 | 325.550845 | 1.1514441 | 0.37953479 | 3.03383022 | 0.0024147 | 0.02068924 |
| SNHG9 | 12.3300311 | 4.35380808 | 1.43548709 | 3.03298308 | 0.00242149 | 0.02072519 |
| NMI | 144.083665 | -1.3091934 | 0.43165843 | -3.0329384 | 0.00242185 | 0.02072519 |
| MRPS35 | 390.655763 | -0.8992452 | 0.29656128 | -3.0322409 | 0.00242745 | 0.0207605 |
| CADPS2 | 9.43474387 | -5.9420065 | 1.96008281 | -3.0315079 | 0.00243336 | 0.0207983 |
| GCDH | 102.979989 | -1.3123304 | 0.43308599 | -3.0301844 | 0.00244404 | 0.02087571 |
| CPSF3 | 516.951398 | -0.7547738 | 0.24909872 | -3.0300188 | 0.00244538 | 0.02087571 |
| RAMP1 | 141.98102 | -2.009176 | 0.66383806 | -3.0266056 | 0.00247316 | 0.02110003 |
| TOR3A | 260.611792 | -1.0108087 | 0.33407335 | -3.0257089 | 0.00248051 | 0.02114985 |
| STOML1 | 53.8469412 | -1.3001158 | 0.4297416 | -3.0253431 | 0.00248351 | 0.02115677 |
| TMEM135 | 166.450761 | -1.4510595 | 0.47965056 | -3.025243 | 0.00248433 | 0.02115677 |
| PHKA1 | 51.8390928 | -2.3991325 | 0.79321214 | -3.0245786 | 0.0024898 | 0.02119044 |
| FUS | 2259.63579 | 0.79553069 | 0.2630506 | 3.02424965 | 0.00249251 | 0.02119343 |
| RMND5B | 142.284874 | -1.0359089 | 0.34254328 | -3.0241695 | 0.00249317 | 0.02119343 |
| DYRK3 | 67.136592 | -2.0203323 | 0.66825923 | -3.0232764 | 0.00250054 | 0.02122736 |
| UACA | 16.6959435 | -3.2586762 | 1.07790138 | -3.0231673 | 0.00250144 | 0.02122736 |
| SMPD1 | 495.578564 | -1.7523754 | 0.57965474 | -3.0231365 | 0.00250169 | 0.02122736 |
| SBF2-AS1 | 45.4084509 | -2.4140166 | 0.79888869 | -3.0217184 | 0.00251344 | 0.02131418 |
| PHB1 | 865.56674 | -1.0800626 | 0.35750665 | -3.0210979 | 0.0025186 | 0.02134501 |
| ILDR2 | 17.8737476 | 3.23585794 | 1.07127024 | 3.02058043 | 0.00252291 | 0.02136863 |
| SESN3 | 326.158087 | 2.07266979 | 0.68653304 | 3.01903865 | 0.00253578 | 0.02146473 |
| LGALS12 | 55.5423538 | 3.09359823 | 1.02497213 | 3.0182267 | 0.00254259 | 0.02150936 |
| SIRT5 | 45.5811743 | -1.459492 | 0.48361011 | -3.0179105 | 0.00254524 | 0.02151882 |
| CFAP36 | 94.501484 | -1.8258255 | 0.60503304 | -3.0177286 | 0.00254677 | 0.02151882 |
| NOTCH3 | 246.036171 | 2.11811918 | 0.70196353 | 3.01742053 | 0.00254936 | 0.02152775 |
| FKBP4 | 589.112516 | 0.91650727 | 0.30377103 | 3.01709905 | 0.00255206 | 0.02153764 |
| SLC16A13 | 83.9368356 | -1.391736 | 0.46136831 | -3.01654 | 0.00255677 | 0.02156444 |
| TET3 | 597.758253 | 0.96322486 | 0.31947746 | 3.015001 | 0.00256978 | 0.02166115 |
| ENSG000002 | 24.6135364 | 3.3380036 | 1.10754646 | 3.01387231 | 0.00257936 | 0.02172885 |
| EEPD1 | 1010.70171 | 1.0415726 | 0.34561615 | 3.01366874 | 0.00258109 | 0.0217304 |
| HUWE1 | 1967.00699 | -0.6225786 | 0.20668395 | -3.0122254 | 0.0025934 | 0.02182092 |
| ABCB9 | 106.614396 | -1.7792517 | 0.5907254 | -3.0119776 | 0.00259552 | 0.02182566 |
| LRRC7 | 45.6399712 | 1.72701695 | 0.57353419 | 3.01118396 | 0.00260231 | 0.02186969 |
| CPNE2 | 436.430963 | -0.7816634 | 0.25963248 | -3.0106534 | 0.00260686 | 0.02189484 |
| ASGR2 | 259.116898 | 2.70394662 | 0.8985927 | 3.00909033 | 0.00262031 | 0.02199464 |
| PNKP | 204.980457 | -0.898653 | 0.2986714 | -3.0088352 | 0.00262251 | 0.02199997 |
| CADM1 | 1430.2883 | 1.30506009 | 0.43384733 | 3.00810909 | 0.00262879 | 0.02203944 |
| SERPINE1 | 62.7735137 | -2.2666774 | 0.75362763 | -3.0076888 | 0.00263243 | 0.02205586 |
| CYGB | 66.1030437 | -3.8277412 | 1.27272341 | -3.0075201 | 0.00263389 | 0.02205586 |
| OGFRL1 | 5168.58542 | 1.6036782 | 0.53361883 | 3.00528787 | 0.0026533 | 0.02220515 |
| MEGF6 | 106.638924 | -3.6457989 | 1.21432498 | -3.0023255 | 0.00267926 | 0.02240904 |

| | | | | | | |
|---|---|---|---|---|---|---|
| RN7SK | 141.900247 | 1.92848629 | 0.64259299 | 3.00110071 | 0.00269006 | 0.02248598 |
| MAP3K7CL | 10.6552977 | -3.2664326 | 1.0885577 | -3.0006977 | 0.00269362 | 0.02250237 |
| SEC62 | 1288.67923 | 0.59500628 | 0.19840696 | 2.99891838 | 0.0027094 | 0.02262074 |
| NDN | 154.295612 | -1.1505411 | 0.38370807 | -2.9984803 | 0.0027133 | 0.02263983 |
| PTP4A1 | 973.862371 | -0.5860608 | 0.19546489 | -2.998292 | 0.00271497 | 0.02264037 |
| SLC22A16 | 9.07199451 | -4.9472755 | 1.65083136 | -2.9968388 | 0.00272795 | 0.02273507 |
| TXNDC12 | 1271.04193 | 0.73228075 | 0.24453749 | 2.99455415 | 0.00274846 | 0.02289244 |
| MED29 | 622.808879 | 0.80610673 | 0.26929948 | 2.99334682 | 0.00275936 | 0.02296959 |
| ATP8B3 | 28.9461692 | 2.15872852 | 0.72126021 | 2.99299545 | 0.00276254 | 0.02298242 |
| ADCY10 | 13.1053425 | -4.4721073 | 1.49464128 | -2.9920941 | 0.00277071 | 0.02303675 |
| SATB1-AS1 | 29.9921101 | 3.14094282 | 1.05039686 | 2.9902439 | 0.00278755 | 0.02316304 |
| PDZD7 | 17.5188223 | -3.1746175 | 1.06175574 | -2.9899697 | 0.00279005 | 0.02317013 |
| PALLD | 228.446911 | -2.595492 | 0.86819727 | -2.9895188 | 0.00279417 | 0.02319064 |
| FDX2 | 122.06081 | -1.0114142 | 0.33834941 | -2.9892596 | 0.00279654 | 0.02319661 |
| IL17RA | 2783.24795 | 0.79008191 | 0.26438695 | 2.98835438 | 0.00280484 | 0.0232517 |
| SAMD9L | 17.3114098 | -3.0623726 | 1.02573327 | -2.9855447 | 0.00283074 | 0.02345026 |
| S100A1 | 20.9593787 | -3.8736698 | 1.29754052 | -2.985394 | 0.00283213 | 0.02345026 |
| ZC3H12C | 637.334213 | -0.7938468 | 0.26595487 | -2.9848931 | 0.00283677 | 0.02347485 |
| ST7 | 240.895143 | -1.1089626 | 0.37170353 | -2.9834599 | 0.00285009 | 0.02357118 |
| RNF114 | 829.240917 | 0.81238643 | 0.27233029 | 2.98309239 | 0.00285352 | 0.02358561 |
| SBNO1 | 638.616466 | 0.68767404 | 0.23072065 | 2.98054826 | 0.00287733 | 0.02376843 |
| AUTS2 | 23.3900678 | 2.89629213 | 0.97182022 | 2.98027564 | 0.00287989 | 0.02377561 |
| RAB13 | 938.371321 | -0.8419673 | 0.2825816 | -2.9795548 | 0.00288668 | 0.02381762 |
| SLC6A17 | 11.2632912 | 6.68138584 | 2.24361525 | 2.97795526 | 0.00290178 | 0.02392821 |
| ATL3 | 654.457272 | -0.8557602 | 0.28738743 | -2.977723 | 0.00290398 | 0.0239323 |
| FBXL6 | 56.5616307 | -1.3792922 | 0.46324227 | -2.9774747 | 0.00290634 | 0.02393764 |
| GPR155 | 78.2539909 | 1.31051389 | 0.44017191 | 2.97727741 | 0.00290821 | 0.02393901 |
| CUL4B | 435.951019 | -0.6628393 | 0.22265544 | -2.9769733 | 0.00291109 | 0.02394873 |
| LRCH1 | 486.845162 | -1.7678554 | 0.59391759 | -2.9766006 | 0.00291463 | 0.02396383 |
| CCS | 203.560136 | 1.27380469 | 0.4281027 | 2.97546524 | 0.00292545 | 0.02403863 |
| RHOB | 904.929845 | 1.37967904 | 0.4637826 | 2.97484002 | 0.00293141 | 0.02407359 |
| BOK | 87.2470998 | -1.1415429 | 0.38376522 | -2.9745867 | 0.00293384 | 0.0240794 |
| SLC43A1 | 53.347243 | -1.425394 | 0.47946096 | -2.9729095 | 0.00294991 | 0.02419722 |
| CLTCL1 | 44.7505304 | 2.13765684 | 0.71948676 | 2.97108571 | 0.00296749 | 0.02432718 |
| CD164 | 7198.95846 | -1.3821731 | 0.46523974 | -2.9708835 | 0.00296944 | 0.024329 |
| TCEAL3 | 316.375192 | 1.19011591 | 0.40076046 | 2.96964405 | 0.00298145 | 0.02441312 |
| ATP6V1B2 | 8976.95226 | -0.772923 | 0.26030489 | -2.9692989 | 0.0029848 | 0.02442631 |
| VEZF1 | 415.560668 | 1.11565181 | 0.37577239 | 2.96895627 | 0.00298813 | 0.02443933 |
| DENND5A | 686.635247 | 0.72920011 | 0.24569737 | 2.96787918 | 0.00299862 | 0.02451085 |
| CD99P1 | 37.7253739 | -4.284035 | 1.44387003 | -2.9670503 | 0.00300672 | 0.02456273 |
| ADAMTSL5 | 12.6705303 | 2.95658994 | 0.99663786 | 2.96656396 | 0.00301148 | 0.02458731 |
| CCL2 | 1851.64898 | 1.90209167 | 0.64126731 | 2.96614475 | 0.00301559 | 0.02460655 |
| LPP | 482.16009 | -1.3536814 | 0.45656563 | -2.9649218 | 0.0030276 | 0.02469023 |
| BID | 1469.15382 | 0.81472565 | 0.27481311 | 2.96465347 | 0.00303024 | 0.02469743 |
| RGS2 | 269.679284 | 1.47164427 | 0.49655388 | 2.96371518 | 0.00303949 | 0.02475848 |
| OXA1L | 1702.62986 | -0.7746706 | 0.26145627 | -2.9629069 | 0.00304749 | 0.02480919 |
| ORC2 | 80.2363989 | -1.4126947 | 0.47684394 | -2.9625933 | 0.00305059 | 0.02482009 |
| LYVE1 | 810.506654 | -5.3620487 | 1.81018444 | -2.962156 | 0.00305493 | 0.02484098 |

| | | | | | | |
|---|---|---|---|---|---|---|
| LYSMD2 | 117.354742 | 1.45573755 | 0.49199486 | 2.95884707 | 0.00308792 | 0.02508492 |
| CATSPER1 | 51.1571429 | 1.51991501 | 0.5136949 | 2.95878939 | 0.0030885 | 0.02508492 |
| FHL2 | 14.2425858 | -2.6757114 | 0.90535365 | -2.9554323 | 0.00312231 | 0.02534487 |
| SAXO4 | 7.76070571 | -6.6552419 | 2.25286651 | -2.9541217 | 0.0031356 | 0.02543806 |
| FCGR1A | 1630.5651 | 1.58303083 | 0.53598137 | 2.95351841 | 0.00314174 | 0.02547312 |
| NIBAN1 | 351.182983 | -1.34521 | 0.45592529 | -2.9505054 | 0.00317255 | 0.02570805 |
| DNM1 | 23.8522347 | -2.092891 | 0.70975144 | -2.9487662 | 0.00319045 | 0.02583825 |
| ACSL1 | 2242.03226 | -0.8938564 | 0.30330047 | -2.9470987 | 0.00320771 | 0.02595202 |
| TBC1D9 | 290.013527 | -1.1488544 | 0.38983177 | -2.9470516 | 0.0032082 | 0.02595202 |
| RAC2 | 3870.52217 | -1.4357272 | 0.48725332 | -2.9465725 | 0.00321317 | 0.02597729 |
| TECPR1 | 251.92714 | -0.7961374 | 0.27027028 | -2.9457082 | 0.00322216 | 0.026035 |
| PRKCA | 1735.12741 | 0.96654468 | 0.32826527 | 2.94440132 | 0.0032358 | 0.02613016 |
| INA | 668.321063 | 4.34401567 | 1.47663036 | 2.94184367 | 0.00326265 | 0.02632967 |
| PLEKHA7 | 37.7904205 | 1.69805459 | 0.57723876 | 2.94168496 | 0.00326432 | 0.02632967 |
| PLCB2 | 2078.22739 | -0.8141512 | 0.27677976 | -2.941513 | 0.00326613 | 0.02632967 |
| RHOC | 2200.99904 | -1.727477 | 0.58737274 | -2.9410235 | 0.0032713 | 0.02635619 |
| IRS1 | 18.5400798 | -4.5912315 | 1.56185602 | -2.9395997 | 0.00328637 | 0.0264624 |
| SLAMF9 | 13.6424152 | -3.4381758 | 1.16977144 | -2.939186 | 0.00329076 | 0.02648044 |
| ABHD3 | 316.347985 | 0.99289461 | 0.33783034 | 2.93903326 | 0.00329238 | 0.02648044 |
| CLPB | 142.980474 | -1.3530406 | 0.46058175 | -2.9376774 | 0.00330681 | 0.02658129 |
| PM20D2 | 54.6954442 | -2.1158444 | 0.72037411 | -2.9371466 | 0.00331247 | 0.0266116 |
| COQ2 | 189.451303 | 1.08713055 | 0.37041389 | 2.9349076 | 0.00333647 | 0.02678905 |
| SLC9A7P1.1 | 54.2498296 | 2.17969792 | 0.74273421 | 2.93469441 | 0.00333876 | 0.02679214 |
| TMEM63A | 610.503545 | -1.7444902 | 0.59506296 | -2.9316061 | 0.00337214 | 0.02704453 |
| NPIPA5 | 5.99374215 | -6.2857624 | 2.14504173 | -2.9303684 | 0.0033856 | 0.02712422 |
| LSM10 | 404.328724 | 0.80335204 | 0.27415 | 2.9303375 | 0.00338594 | 0.02712422 |
| ABCB8 | 386.133998 | -1.2457854 | 0.42521056 | -2.9298082 | 0.00339171 | 0.02715497 |
| SLC30A3 | 30.9146883 | -3.8998076 | 1.3311931 | -2.9295582 | 0.00339444 | 0.02716134 |
| SPON1 | 5.83295202 | -6.2462899 | 2.13267884 | -2.9288469 | 0.00340222 | 0.02720807 |
| BPHL | 125.982695 | -1.3275554 | 0.45332716 | -2.928471 | 0.00340634 | 0.02722549 |
| ACSS1 | 83.9700601 | -1.0644427 | 0.36358615 | -2.9276216 | 0.00341565 | 0.02727503 |
| STK10 | 1854.10184 | -0.8017421 | 0.27390086 | -2.9271253 | 0.00342111 | 0.02727503 |
| SRM | 535.988527 | -1.1638109 | 0.39759562 | -2.9271221 | 0.00342115 | 0.02727503 |
| FAM66D | 15.640373 | -3.3178577 | 1.13352363 | -2.9270301 | 0.00342216 | 0.02727503 |
| RCN1 | 218.577187 | -1.5055162 | 0.5143622 | -2.9269573 | 0.00342296 | 0.02727503 |
| TNKS2 | 1048.16558 | 0.61149219 | 0.20892533 | 2.92684554 | 0.00342419 | 0.02727503 |
| ZNF483 | 23.9950727 | 3.27762358 | 1.11995631 | 2.92656377 | 0.00342729 | 0.02728428 |
| S100A16 | 80.9653584 | -10.039906 | 3.43323054 | -2.9243321 | 0.00345196 | 0.02746511 |
| ACAD11 | 43.1531499 | -1.9228656 | 0.65760865 | -2.9240273 | 0.00345534 | 0.02747644 |
| TTC13 | 83.6684664 | -1.6111906 | 0.55106856 | -2.9237571 | 0.00345834 | 0.02748474 |
| LINC02009 | 13.1270651 | -6.4313766 | 2.20002407 | -2.923321 | 0.00346319 | 0.02750273 |
| MAMDC2 | 96.1534493 | -3.911609 | 1.33812512 | -2.9232012 | 0.00346453 | 0.02750273 |
| ZNF319 | 156.898244 | 1.31057923 | 0.44839011 | 2.92285488 | 0.00346838 | 0.02751778 |
| TMEM62 | 134.828276 | -1.7711229 | 0.60602794 | -2.9225103 | 0.00347222 | 0.0275327 |
| ENSG000002 | 32.1200609 | -1.7440125 | 0.5969187 | -2.9216918 | 0.00348136 | 0.02758957 |
| PYCARD | 683.175199 | 0.71318765 | 0.24413137 | 2.92132739 | 0.00348543 | 0.02759661 |
| ABCA7 | 383.676122 | 2.20461109 | 0.75467791 | 2.92126092 | 0.00348618 | 0.02759661 |
| MEX3B | 48.7198845 | 1.52450544 | 0.52191096 | 2.92100676 | 0.00348902 | 0.02759709 |

| | | | | | | |
|---|---|---|---|---|---|---|
| AKAP7 | 14.2852391 | -3.2960332 | 1.12842898 | -2.9209044 | 0.00349017 | 0.02759709 |
| IGF1R | 102.535055 | -1.4621035 | 0.50061628 | -2.9206071 | 0.0034935 | 0.0276079 |
| LLGL1 | 220.37639 | -1.2208865 | 0.41807206 | -2.9202776 | 0.0034972 | 0.02762156 |
| ACVR1 | 428.754516 | 1.1083533 | 0.37969807 | 2.91903856 | 0.00351113 | 0.027716 |
| PCYT2 | 247.535479 | -1.5003376 | 0.51418742 | -2.9178808 | 0.00352419 | 0.02780349 |
| RAB24 | 144.178329 | 0.90930942 | 0.3117145 | 2.9171226 | 0.00353277 | 0.02785552 |
| RAPH1 | 1337.16369 | 1.1908097 | 0.40824717 | 2.91688414 | 0.00353547 | 0.02786118 |
| GPSM3 | 1929.41716 | 0.65057771 | 0.2230552 | 2.91666691 | 0.00353793 | 0.02786495 |
| FUT7 | 16.8126674 | 3.41175483 | 1.17004978 | 2.9159057 | 0.00354658 | 0.02791736 |
| L3MBTL4-AS1 | 9.43980987 | -5.005574 | 1.71721512 | -2.9149371 | 0.0035576 | 0.02798847 |
| ZNF93 | 22.2271617 | -2.5602798 | 0.87840695 | -2.9146852 | 0.00356047 | 0.02799538 |
| LRRC37A6P | 5.70505387 | 5.69566119 | 1.95431757 | 2.914399 | 0.00356374 | 0.02800538 |
| CRELD2 | 210.136655 | 0.85879152 | 0.29471492 | 2.91397371 | 0.0035686 | 0.02802788 |
| ZNF746 | 366.633023 | 0.84123586 | 0.2888178 | 2.91268705 | 0.00358334 | 0.02812788 |
| PDE8A | 351.352263 | -1.3103413 | 0.45006426 | -2.9114538 | 0.00359751 | 0.02822338 |
| PLPP3 | 25.1156477 | -5.5573056 | 1.90895173 | -2.9111819 | 0.00360064 | 0.02823218 |
| HMGCL | 242.871982 | -0.8519281 | 0.29266291 | -2.9109534 | 0.00360328 | 0.02823707 |
| FOSL2 | 663.127287 | 1.37840286 | 0.47366645 | 2.91007069 | 0.00361347 | 0.02827622 |
| ARL6 | 20.3715576 | -2.7020653 | 0.92856109 | -2.9099489 | 0.00361488 | 0.02827622 |
| LINC03040 | 8.90200193 | 6.33538084 | 2.17738362 | 2.90963006 | 0.00361857 | 0.02827622 |
| OSCAR | 370.035983 | 1.4645079 | 0.50334255 | 2.90956508 | 0.00361932 | 0.02827622 |
| SAMD4A | 232.930426 | -2.1246736 | 0.73025219 | -2.9095067 | 0.00362 | 0.02827622 |
| MYL6 | 7622.97526 | -0.7194516 | 0.24728236 | -2.9094336 | 0.00362084 | 0.02827622 |
| TLR6 | 546.650951 | 0.70249316 | 0.24146451 | 2.90930196 | 0.00362237 | 0.02827622 |
| ITGAV | 2270.23327 | 1.03017308 | 0.35414088 | 2.90893576 | 0.00362661 | 0.02829363 |
| SETD3 | 1440.51395 | 0.70246948 | 0.24154854 | 2.90819182 | 0.00363525 | 0.02834264 |
| CHIT1 | 14683.2734 | -5.1325699 | 1.76495399 | -2.9080474 | 0.00363693 | 0.02834264 |
| EIF2AK2 | 214.431018 | -0.7592748 | 0.26113184 | -2.9076302 | 0.00364179 | 0.02835345 |
| MICAL1 | 1217.68689 | 0.66890528 | 0.23006614 | 2.9074477 | 0.00364391 | 0.02835345 |
| FMR1 | 135.212498 | 0.94344213 | 0.32449596 | 2.90740793 | 0.00364438 | 0.02835345 |
| ENSG000002 | 11.7430455 | 3.6821314 | 1.26656254 | 2.90718482 | 0.00364698 | 0.02835797 |
| SLC35G2 | 17.4890271 | -3.4311107 | 1.18112001 | -2.9049637 | 0.00367296 | 0.02853475 |
| POGZ | 418.203712 | 0.69039033 | 0.23766455 | 2.90489401 | 0.00367377 | 0.02853475 |
| BLVRB | 1360.19347 | -1.6403258 | 0.5647703 | -2.9044123 | 0.00367943 | 0.02856289 |
| HMG20A | 445.107014 | 0.91911097 | 0.31648305 | 2.9041396 | 0.00368264 | 0.02857199 |
| BCL2A1 | 578.281356 | -2.2183983 | 0.76427214 | -2.9026288 | 0.00370045 | 0.02869432 |
| ZBTB7B | 1102.4699 | 0.85066296 | 0.2931965 | 2.90134075 | 0.0037157 | 0.02879664 |
| TTC39B | 102.210822 | -1.9189891 | 0.6614925 | -2.900999 | 0.00371975 | 0.02881217 |
| LGALSL | 32.1587889 | -2.6222942 | 0.90402924 | -2.9006741 | 0.00372361 | 0.02881873 |
| FYTTD1 | 1584.28871 | 0.57249647 | 0.19737296 | 2.90058207 | 0.0037247 | 0.02881873 |
| DYNC2I2 | 80.2576953 | -1.4993499 | 0.5171149 | -2.8994521 | 0.00373815 | 0.02889076 |
| FANCA | 93.5577147 | -2.6127786 | 0.90115607 | -2.8993631 | 0.00373922 | 0.02889076 |
| LDLRAP1 | 127.524267 | -1.8066678 | 0.6231432 | -2.8992819 | 0.00374018 | 0.02889076 |
| MCTP1 | 372.684993 | 0.82888371 | 0.28618347 | 2.89633681 | 0.00377547 | 0.02914729 |
| INTS8 | 230.113295 | 0.91173016 | 0.31481188 | 2.89611106 | 0.00377819 | 0.02915224 |
| BHLHE41 | 685.62215 | -1.0729153 | 0.37069605 | -2.8943262 | 0.00379973 | 0.02930238 |
| PNPLA3 | 50.5328464 | -2.469065 | 0.85313921 | -2.8940939 | 0.00380254 | 0.02930798 |
| ENSG000002 | 20.1579296 | 5.04466918 | 1.74341494 | 2.89355625 | 0.00380906 | 0.02934209 |

| | | | | | | |
|---|---|---|---|---|---|---|
| TMEM44 | 38.6214397 | -1.7399476 | 0.60162317 | -2.8920888 | 0.0038269 | 0.02946334 |
| RNASE6 | 1771.02474 | 1.35813888 | 0.47014982 | 2.88873633 | 0.00386793 | 0.02976295 |
| NPIPB13 | 43.1898129 | -1.8902268 | 0.65455325 | -2.8878122 | 0.00387931 | 0.02983418 |
| MFSD12 | 3903.48923 | -2.0551067 | 0.71181621 | -2.8871311 | 0.00388772 | 0.02988247 |
| MACROH2A1 | 3479.44148 | -0.6021239 | 0.20856866 | -2.8869339 | 0.00389016 | 0.02988485 |
| ELF1 | 1046.09367 | 0.72777645 | 0.25224545 | 2.8851916 | 0.00391176 | 0.03003432 |
| CTBS | 352.23667 | 0.86508929 | 0.299958 | 2.88403471 | 0.00392615 | 0.03012841 |
| F11R | 1111.80232 | -1.2147634 | 0.42143976 | -2.8824128 | 0.00394642 | 0.03025429 |
| MFNG | 427.674775 | 0.92498651 | 0.32091097 | 2.88237736 | 0.00394687 | 0.03025429 |
| NDUFA4 | 1650.86557 | 0.63560057 | 0.22054526 | 2.88195072 | 0.00395222 | 0.03027876 |
| DUBR | 6.2793439 | -5.344296 | 1.85705361 | -2.8778361 | 0.00400413 | 0.03065977 |
| MLPH | 57.4599439 | -2.8397148 | 0.98685056 | -2.8775531 | 0.00400773 | 0.03067056 |
| GTF2IP1 | 339.749606 | 0.95049726 | 0.33035628 | 2.87718843 | 0.00401236 | 0.0306893 |
| CLEC2B | 19.698327 | -2.614612 | 0.90881799 | -2.8769369 | 0.00401556 | 0.03069394 |
| SAMSN1 | 1437.50199 | 1.77321722 | 0.6163859 | 2.87679719 | 0.00401734 | 0.03069394 |
| RAB4B | 337.324757 | 0.74326532 | 0.25853684 | 2.87489129 | 0.00404167 | 0.03086303 |
| CEP68 | 141.404309 | -0.8621874 | 0.29993474 | -2.8745833 | 0.00404561 | 0.03087636 |
| ST20 | 78.6348774 | -1.8709125 | 0.6509312 | -2.8742092 | 0.00405041 | 0.03089617 |
| SMIM10 | 12.9710056 | -4.4002658 | 1.53132593 | -2.8735004 | 0.00405951 | 0.03094877 |
| ENSG000002 | 15.1860599 | 3.97624922 | 1.38418742 | 2.87262344 | 0.00407079 | 0.03099812 |
| RUNDC3A | 13.7265774 | 3.92563372 | 1.36662405 | 2.87250449 | 0.00407232 | 0.03099812 |
| CNR1 | 7.78658386 | -6.6600829 | 2.31858053 | -2.8724829 | 0.0040726 | 0.03099812 |
| ZNF626 | 7.55174101 | -3.6128874 | 1.25822797 | -2.8714092 | 0.00408646 | 0.03108677 |
| ZNF148 | 519.414351 | 0.8969497 | 0.31256519 | 2.86964043 | 0.00410939 | 0.03124425 |
| MANSC1 | 5.4854936 | -6.157307 | 2.14695625 | -2.8679239 | 0.00413175 | 0.03139727 |
| MAD2L2 | 221.995111 | -1.0139124 | 0.35377296 | -2.8659973 | 0.00415698 | 0.0315719 |
| SMIM10L2A | 17.0762854 | 3.17074526 | 1.10692294 | 2.86446792 | 0.0041771 | 0.03170761 |
| NINJ1 | 3632.13621 | 1.24687912 | 0.43532728 | 2.86423383 | 0.00418019 | 0.03171391 |
| SETX | 1283.77028 | 0.71295373 | 0.24906481 | 2.86252293 | 0.00420283 | 0.03186842 |
| MIR9-3HG | 7.0171267 | 5.99974182 | 2.0977591 | 2.86007188 | 0.00423545 | 0.03209193 |
| CIMAP1B | 66.236683 | 1.21016722 | 0.42314054 | 2.85996517 | 0.00423688 | 0.03209193 |
| ENSG000002 | 9.58222167 | 4.95700779 | 1.73372917 | 2.85915924 | 0.00424765 | 0.03215624 |
| B4GAT1 | 84.8511557 | -1.7719737 | 0.6198781 | -2.8585841 | 0.00425536 | 0.03219723 |
| USF2 | 2157.77782 | -0.9666365 | 0.33830001 | -2.8573351 | 0.00427215 | 0.03230681 |
| ARHGAP31 | 3417.59451 | 1.32103921 | 0.46245231 | 2.85659553 | 0.00428211 | 0.03236474 |
| SYAP1 | 466.459969 | 0.95373874 | 0.33399024 | 2.85558864 | 0.00429571 | 0.03245008 |
| LIMA1 | 364.257527 | -1.7085475 | 0.5984197 | -2.855099 | 0.00430234 | 0.03248268 |
| CELSR2 | 71.6835256 | 1.75978703 | 0.61645135 | 2.85470547 | 0.00430768 | 0.03250548 |
| PTGR1 | 290.749458 | -0.8837042 | 0.30965183 | -2.8538639 | 0.0043191 | 0.0325742 |
| KIAA1549 | 18.2733877 | -2.7759145 | 0.97283041 | -2.8534414 | 0.00432485 | 0.03260004 |
| AFG1L | 53.540689 | -1.7306856 | 0.60660057 | -2.8530894 | 0.00432964 | 0.03261867 |
| TM7SF2 | 193.007439 | -1.2249896 | 0.42947566 | -2.852291 | 0.00434053 | 0.03268318 |
| MYO1D | 1385.09725 | -2.0021646 | 0.70221636 | -2.8512076 | 0.00435535 | 0.03277332 |
| ZKSCAN4 | 69.4014357 | 1.1601017 | 0.40690106 | 2.85106582 | 0.00435729 | 0.03277332 |
| MT1X | 19.8458469 | 2.17702234 | 0.76362515 | 2.85090443 | 0.00435951 | 0.03277332 |
| LIPA | 36913.7849 | -1.2758166 | 0.44768692 | -2.8497965 | 0.00437472 | 0.0328701 |
| NOD2 | 19.6234181 | 2.2644351 | 0.79534136 | 2.84712353 | 0.00441162 | 0.03312964 |
| KRT79 | 15.2017153 | -5.8157067 | 2.04306163 | -2.8465645 | 0.00441938 | 0.03317013 |

| | | | | | | |
|---|---|---|---|---|---|---|
| PRKX | 385.790097 | -1.4586541 | 0.51348388 | -2.8407009 | 0.00450145 | 0.03376809 |
| HABP4 | 84.0916832 | -1.4134146 | 0.49762226 | -2.8403364 | 0.0045066 | 0.03378866 |
| NOTCH2NLA | 143.125242 | -1.7113789 | 0.60265913 | -2.8397129 | 0.00451542 | 0.03383671 |
| EPDR1 | 38.3698635 | -2.8525713 | 1.00506031 | -2.8382091 | 0.00453675 | 0.03397841 |
| LRCH4 | 482.674484 | 0.7915493 | 0.27892658 | 2.83784108 | 0.00454198 | 0.03399947 |
| OVOL1 | 15.4617708 | -3.6489308 | 1.28614034 | -2.8371171 | 0.00455229 | 0.03405851 |
| PIP4K2C | 281.297956 | -1.0588157 | 0.3732303 | -2.8368964 | 0.00455544 | 0.03406392 |
| ABCB5 | 29.5611686 | -4.2453242 | 1.49682745 | -2.8362148 | 0.00456517 | 0.03411833 |
| TEC | 69.0322124 | 1.66804904 | 0.58815991 | 2.83604681 | 0.00456758 | 0.03411833 |
| ALCAM | 1791.81592 | -1.1275668 | 0.39762036 | -2.8357874 | 0.00457129 | 0.03412789 |
| TRMT5 | 179.197056 | -1.6430045 | 0.57946457 | -2.8353838 | 0.00457707 | 0.03415289 |
| ARHGAP19 | 167.000556 | 0.99504558 | 0.3510057 | 2.83484166 | 0.00458484 | 0.03419273 |
| NABP1 | 769.769149 | -1.2485177 | 0.4405071 | -2.8342738 | 0.004593 | 0.03422064 |
| ZBTB8OS | 134.112485 | 0.90434653 | 0.31908125 | 2.83422021 | 0.00459377 | 0.03422064 |
| SEC61B | 606.711791 | 0.75040529 | 0.26477987 | 2.8340723 | 0.00459589 | 0.03422064 |
| TUT4 | 171.291626 | 1.00283413 | 0.35393572 | 2.8333793 | 0.00460587 | 0.03427675 |
| SORT1 | 2536.05614 | -1.3775294 | 0.48643497 | -2.8318882 | 0.0046274 | 0.0344034 |
| HSPB8 | 17.9140048 | -5.4714648 | 1.93210887 | -2.8318616 | 0.00462779 | 0.0344034 |
| SNHG4 | 73.3451394 | -1.6808421 | 0.59374812 | -2.8309009 | 0.00464171 | 0.03448864 |
| ACO1 | 655.350714 | -1.4253008 | 0.50351 | -2.8307299 | 0.00464419 | 0.03448884 |
| CD274 | 44.3099876 | -1.9422846 | 0.68625976 | -2.8302469 | 0.00465121 | 0.03452269 |
| ILK | 1348.01314 | -0.5976202 | 0.21121471 | -2.829444 | 0.0046629 | 0.03458733 |
| ATP6V1F | 4861.3327 | -0.733161 | 0.25913064 | -2.8293104 | 0.00466484 | 0.03458733 |
| CDK19 | 539.638619 | 0.66214702 | 0.23409568 | 2.82853158 | 0.00467621 | 0.0346533 |
| TBL1XR1 | 926.894525 | 0.96697066 | 0.34194817 | 2.82782816 | 0.0046865 | 0.03471121 |
| UBE2D4 | 291.97787 | -1.1849544 | 0.41913757 | -2.8271252 | 0.0046968 | 0.03476916 |
| MRPL20-AS1 | 90.0683754 | 1.14599524 | 0.40544187 | 2.82653405 | 0.00470547 | 0.03481505 |
| PLD4 | 2917.695 | 1.69292391 | 0.59903332 | 2.82609304 | 0.00471196 | 0.03484467 |
| CD22 | 33.2908094 | -2.7843196 | 0.98544224 | -2.8254519 | 0.0047214 | 0.0348961 |
| THEM6 | 167.647876 | -1.6509114 | 0.58466494 | -2.8236881 | 0.00474746 | 0.03507024 |
| TRIM38 | 381.316924 | 0.81379891 | 0.28824634 | 2.82327583 | 0.00475357 | 0.03509691 |
| PPT2 | 29.35578 | -2.2337165 | 0.79147052 | -2.8222359 | 0.00476901 | 0.0351791 |
| DPYD | 780.749637 | 0.76751541 | 0.2719575 | 2.82218884 | 0.00476971 | 0.0351791 |
| ARIH2 | 708.230395 | 0.65436349 | 0.23190084 | 2.82173833 | 0.00477641 | 0.03521006 |
| PAM | 134.753217 | -1.7278477 | 0.61307422 | -2.8183336 | 0.00482736 | 0.03556698 |
| SERHL2 | 9.97748261 | 4.04334369 | 1.43482126 | 2.81801212 | 0.0048322 | 0.03558394 |
| SLC6A12 | 93.6624791 | -2.7418856 | 0.97344759 | -2.8166751 | 0.00485236 | 0.0357021 |
| FARS2 | 71.293948 | -1.2735886 | 0.45217061 | -2.8166108 | 0.00485333 | 0.0357021 |
| VSTM2L | 7.36149098 | -6.5790442 | 2.33643198 | -2.8158509 | 0.00486482 | 0.03576791 |
| IGFBP4 | 2351.43794 | 2.0801751 | 0.73913422 | 2.81434014 | 0.00488775 | 0.03591766 |
| ASCC1 | 181.514022 | -0.8479206 | 0.30139117 | -2.8133557 | 0.00490274 | 0.03600897 |
| PPP1R16A | 89.9954702 | -1.1458897 | 0.40733369 | -2.8131474 | 0.00490592 | 0.03601348 |
| TMC4 | 12.7974753 | 4.42422568 | 1.57320772 | 2.81223237 | 0.00491989 | 0.03609722 |
| GDPD5 | 100.212121 | -2.5507198 | 0.90745051 | -2.8108638 | 0.00494087 | 0.03623219 |
| DDX60L | 372.026894 | -1.5002554 | 0.5338113 | -2.8104601 | 0.00494707 | 0.0362545 |
| FAM168B | 1775.94242 | 0.57096261 | 0.2031657 | 2.81032974 | 0.00494908 | 0.0362545 |
| ELK1 | 769.259356 | 0.6092114 | 0.2167963 | 2.81006359 | 0.00495317 | 0.03626558 |
| SMURF1 | 447.824912 | 0.98248556 | 0.34973474 | 2.80923067 | 0.00496601 | 0.0363406 |

| | | | | | | |
|---|---|---|---|---|---|---|
| CD2BP2-DT | 45.9980373 | -2.4264828 | 0.8640197 | -2.8083651 | 0.00497937 | 0.03640153 |
| GOLGA3 | 265.025295 | -0.8234549 | 0.29321599 | -2.808356 | 0.00497952 | 0.03640153 |
| DNAJA4 | 264.619703 | -1.6460239 | 0.58618459 | -2.80803 | 0.00498456 | 0.03641943 |
| DLAT | 588.166217 | -1.0777849 | 0.38387897 | -2.8076164 | 0.00499096 | 0.03644727 |
| SEC22B | 1155.73354 | 0.97513206 | 0.34739188 | 2.80700879 | 0.00500039 | 0.03649711 |
| RAC1P2 | 68.5042753 | 1.36854134 | 0.48768116 | 2.80622147 | 0.00501262 | 0.03656741 |
| ZFP36L1 | 4889.66258 | 0.86680007 | 0.30891237 | 2.80597396 | 0.00501647 | 0.03657651 |
| CYP4V2 | 415.971435 | -1.0644908 | 0.37945724 | -2.8052984 | 0.005027 | 0.03663424 |
| FAM20A | 252.642721 | -1.659664 | 0.59174283 | -2.8047049 | 0.00503627 | 0.03668272 |
| GPR150 | 8.63631441 | -5.8113543 | 2.07298388 | -2.8033765 | 0.00505706 | 0.03681508 |
| STK16 | 163.785247 | -0.9745338 | 0.34767406 | -2.8030097 | 0.00506281 | 0.03683787 |
| MTCL1 | 18.957712 | -3.7405869 | 1.33492121 | -2.8021032 | 0.00507706 | 0.03692243 |
| TECPR2 | 157.834254 | -1.0234152 | 0.36536734 | -2.8010584 | 0.00509353 | 0.037023 |
| PAFAH1B2 | 1268.27429 | 0.59253851 | 0.21157213 | 2.80064539 | 0.00510005 | 0.03705124 |
| LINC00702 | 6.6979735 | -6.4426847 | 2.30093053 | -2.8000344 | 0.00510972 | 0.03710224 |
| MT1G | 8.78243205 | 6.32243201 | 2.25899669 | 2.79877878 | 0.00512963 | 0.03722757 |
| CNKSR3 | 39.8175536 | -2.5421507 | 0.9083914 | -2.7985192 | 0.00513375 | 0.03723827 |
| RFNG | 195.465941 | -0.8956685 | 0.32008003 | -2.7982642 | 0.00513781 | 0.03724843 |
| CCDC22 | 252.28457 | -0.7787511 | 0.27835907 | -2.7976495 | 0.00514759 | 0.03730013 |
| TSR2 | 330.284415 | 0.75492663 | 0.26991132 | 2.7969432 | 0.00515886 | 0.03736249 |
| GPR27 | 10.7012731 | 4.1494279 | 1.48479735 | 2.79460891 | 0.00519625 | 0.03761391 |
| EPRS1 | 692.852585 | -0.9145807 | 0.32729922 | -2.794326 | 0.0052008 | 0.03762744 |
| TMEM26 | 210.477238 | 2.50900224 | 0.89844549 | 2.79260375 | 0.00522857 | 0.03780886 |
| ENSG000002 | 5.85425101 | -5.2401405 | 1.87655166 | -2.7924307 | 0.00523137 | 0.03780962 |
| CDC42EP4 | 29.4129997 | -2.0441328 | 0.73209405 | -2.7921724 | 0.00523554 | 0.03782035 |
| COL9A2 | 20.61549 | -2.047426 | 0.73334944 | -2.7918833 | 0.00524023 | 0.0378347 |
| ZNF641 | 529.016014 | 1.22508101 | 0.43890384 | 2.79122874 | 0.00525083 | 0.03789182 |
| HTRA3 | 31.4578922 | -2.123662 | 0.76105858 | -2.7904054 | 0.00526421 | 0.03796881 |
| ALS2CL | 6.95845267 | -4.5171162 | 1.6193908 | -2.7893923 | 0.00528071 | 0.03806246 |
| POLR2B | 923.820753 | 0.62162271 | 0.22286172 | 2.78927536 | 0.00528261 | 0.03806246 |
| HHEX | 364.55829 | 1.54296763 | 0.55324528 | 2.78893954 | 0.00528809 | 0.03808241 |
| CCL4 | 1666.33109 | 1.39118889 | 0.49915961 | 2.78706223 | 0.00531883 | 0.0382841 |
| PAIP2 | 761.760902 | 0.8137116 | 0.29200791 | 2.78660803 | 0.00532629 | 0.03831815 |
| LINC02642 | 11.2944225 | 4.23404578 | 1.51989329 | 2.785752 | 0.00534037 | 0.03839981 |
| FMNL1 | 2614.05301 | 1.10855338 | 0.39822363 | 2.78374585 | 0.00537351 | 0.03861834 |
| PTK2B | 941.870484 | 1.3070184 | 0.46959547 | 2.78328583 | 0.00538114 | 0.03865336 |
| NDUFB6 | 143.573241 | -1.1587849 | 0.41645561 | -2.7824931 | 0.0053943 | 0.0387281 |
| FHOD1 | 1910.94712 | 1.1921305 | 0.42878209 | 2.78027118 | 0.00543135 | 0.03897418 |
| IFITM10 | 58.4436141 | 2.39080615 | 0.86027917 | 2.77910501 | 0.00545089 | 0.03909441 |
| DEDD2 | 380.255534 | 0.73608038 | 0.26494787 | 2.77820831 | 0.00546596 | 0.03918245 |
| IL11RA | 39.2528642 | -1.7865567 | 0.64317921 | -2.7776966 | 0.00547457 | 0.03922307 |
| RASGRF1 | 202.823599 | -2.618188 | 0.94262836 | -2.7775401 | 0.00547721 | 0.03922307 |
| KCTD17 | 143.070549 | 0.88506004 | 0.31882417 | 2.77601304 | 0.005503 | 0.03938768 |
| NR6A1 | 63.2191387 | -1.7007181 | 0.61279459 | -2.7753477 | 0.00551427 | 0.03944825 |
| BAZ1A | 620.668383 | 1.24550235 | 0.44883042 | 2.77499538 | 0.00552025 | 0.03947091 |
| PRRG4 | 31.64023 | 2.09016002 | 0.75331616 | 2.77461194 | 0.00552676 | 0.03949736 |
| NAGA | 3558.08709 | 0.75156238 | 0.27090937 | 2.77422072 | 0.00553341 | 0.03951041 |
| RMDN2 | 25.5329772 | -1.6724494 | 0.602864 | -2.7741735 | 0.00553422 | 0.03951041 |

| | | | | | | |
|---|---|---|---|---|---|---|
| RUVBL2 | 343.003024 | -0.796606 | 0.28722878 | -2.7734197 | 0.00554705 | 0.03958194 |
| PIK3R1 | 2004.27727 | 1.46004131 | 0.52663204 | 2.77241261 | 0.00556425 | 0.03968446 |
| NMB | 161.262639 | -2.3484438 | 0.8472595 | -2.7718117 | 0.00557453 | 0.03973761 |
| SLC22A4 | 34.5256162 | 2.09629562 | 0.75634524 | 2.77161211 | 0.00557795 | 0.0397418 |
| MYLK3 | 11.7197842 | 5.27996506 | 1.90522391 | 2.77130947 | 0.00558313 | 0.03975683 |
| DUSP13A | 91.9309805 | -8.4766304 | 3.05887571 | -2.7711588 | 0.00558572 | 0.03975683 |
| KDM5A | 524.292052 | 1.16332485 | 0.41988407 | 2.77058585 | 0.00559555 | 0.03979471 |
| SPINT1 | 584.740608 | -2.1616703 | 0.78024027 | -2.7705188 | 0.00559671 | 0.03979471 |
| SLCO4A1 | 118.951315 | -2.6013107 | 0.93902723 | -2.7702186 | 0.00560187 | 0.03981125 |
| ADNP2 | 316.264518 | 0.69100866 | 0.24947891 | 2.76980798 | 0.00560893 | 0.03982696 |
| BACH2 | 29.8842609 | 2.2331293 | 0.80625354 | 2.76976062 | 0.00560975 | 0.03982696 |
| BAIAP2 | 1108.04097 | 0.65696758 | 0.23722349 | 2.76940358 | 0.0056159 | 0.03985049 |
| TNFRSF10C | 32.4032069 | -2.1511557 | 0.77707978 | -2.7682559 | 0.00563572 | 0.03995985 |
| RET | 17.9692775 | -3.050294 | 1.10191256 | -2.7681815 | 0.00563701 | 0.03995985 |
| DSEL | 31.1762617 | -2.4319173 | 0.87858742 | -2.7679855 | 0.0056404 | 0.03996371 |
| SLC33A1 | 594.04901 | 0.66723069 | 0.24108944 | 2.76756498 | 0.00564768 | 0.03999513 |
| SERP1 | 1754.19357 | 0.61695 | 0.22306736 | 2.76575643 | 0.00567909 | 0.0401713 |
| CST6 | 13.3464585 | 3.01305157 | 1.08945773 | 2.76564338 | 0.00568106 | 0.0401713 |
| ABL1 | 738.199061 | 0.74594508 | 0.26971888 | 2.76563909 | 0.00568114 | 0.0401713 |
| NT5M | 9.96122341 | -3.6332506 | 1.31386905 | -2.7653065 | 0.00568693 | 0.04019205 |
| NUBPL | 57.418817 | -1.5979756 | 0.57799295 | -2.7646974 | 0.00569756 | 0.04024692 |
| PRUNE2 | 22.0305591 | 2.65307843 | 0.9601847 | 2.76309176 | 0.00572567 | 0.04042511 |
| PLPP6 | 46.6569214 | -1.3951248 | 0.50523717 | -2.7613266 | 0.00575671 | 0.04062384 |
| IRF5 | 763.22503 | 0.85378594 | 0.30929048 | 2.76046631 | 0.00577189 | 0.04071053 |
| R3HDM2 | 158.45554 | -1.4521233 | 0.52627922 | -2.7592259 | 0.00579385 | 0.04083093 |
| PEBP1 | 1690.94602 | -0.5356697 | 0.19414135 | -2.7591735 | 0.00579478 | 0.04083093 |
| LYNX1 | 230.870624 | -1.2277874 | 0.44515958 | -2.7580838 | 0.00581413 | 0.04094674 |
| LINC01504 | 10.1831209 | 3.40083951 | 1.23346013 | 2.75715398 | 0.00583069 | 0.04104279 |
| CYBRD1 | 322.776813 | -2.1545966 | 0.78206294 | -2.7550168 | 0.00586891 | 0.04128161 |
| TBC1D17 | 139.237967 | -1.1912368 | 0.43240209 | -2.7549285 | 0.0058705 | 0.04128161 |
| IL10RA | 2602.64202 | 1.44559977 | 0.52486735 | 2.7542193 | 0.00588323 | 0.04135047 |
| PTPRF | 29.4816453 | -3.6775343 | 1.33634224 | -2.7519405 | 0.00592433 | 0.04161849 |
| POMGNT1 | 313.406636 | -1.2283625 | 0.44639901 | -2.7517143 | 0.00592842 | 0.04162642 |
| TUG1 | 793.032001 | 0.5872862 | 0.21372027 | 2.74792003 | 0.00599746 | 0.04209015 |
| POLD4 | 702.363759 | 0.64465358 | 0.23465189 | 2.74727629 | 0.00600925 | 0.04215179 |
| MTA2 | 1253.76853 | 0.97494885 | 0.35492321 | 2.74692896 | 0.00601562 | 0.04217539 |
| SH3BGRL3 | 10297.0432 | -1.2504284 | 0.45524837 | -2.746695 | 0.00601991 | 0.04218443 |
| MTARC2 | 10.9532093 | -3.2448026 | 1.18189436 | -2.7454252 | 0.00604325 | 0.04229498 |
| PELI1 | 536.853304 | 1.17564984 | 0.42822853 | 2.74537957 | 0.00604409 | 0.04229498 |
| TSPAN17 | 433.37399 | -1.3931528 | 0.5074599 | -2.7453456 | 0.00604472 | 0.04229498 |
| CUL9 | 186.037927 | -1.1568842 | 0.42143237 | -2.7451241 | 0.0060488 | 0.04230246 |
| STXBP1 | 649.008366 | 1.12513658 | 0.41001775 | 2.74411677 | 0.0060674 | 0.04241136 |
| GPR65 | 1146.53308 | 1.33394846 | 0.48652618 | 2.74178145 | 0.0061107 | 0.04269281 |
| COBLL1 | 6.66372291 | -5.4368029 | 1.98309069 | -2.7415806 | 0.00611444 | 0.04269768 |
| MORC3 | 357.231426 | 0.79996502 | 0.29184379 | 2.74107255 | 0.0061239 | 0.0427425 |
| PPP1R12B | 98.3191221 | -2.3107816 | 0.84332887 | -2.7400718 | 0.00614258 | 0.04285157 |
| ZC3H12D | 28.4288829 | -1.7823712 | 0.650551 | -2.739787 | 0.0061479 | 0.04286742 |
| SMC5 | 160.628124 | 0.92704562 | 0.33853602 | 2.73839579 | 0.00617397 | 0.04302784 |

| | | | | | | |
|---|---|---|---|---|---|---|
| PARP12 | 310.912758 | -1.9380629 | 0.7081418 | -2.7368289 | 0.00620346 | 0.04321187 |
| MYO6 | 266.006502 | -1.8096266 | 0.66148856 | -2.7356884 | 0.00622499 | 0.0433404 |
| ENSG000002 | 29.304006 | -2.6444964 | 0.96686679 | -2.7351197 | 0.00623576 | 0.04339384 |
| AGBL5 | 157.310029 | -0.7838065 | 0.28675356 | -2.7333803 | 0.00626879 | 0.04360209 |
| MRPL15 | 454.915245 | -0.648753 | 0.23748737 | -2.7317366 | 0.00630015 | 0.0437985 |
| FNTA | 524.239444 | 0.72325672 | 0.26486069 | 2.73070616 | 0.00631988 | 0.04391393 |
| ECI1 | 160.752221 | -0.8189485 | 0.29998787 | -2.7299389 | 0.00633461 | 0.0439945 |
| PROX1 | 11.4990518 | -2.9594028 | 1.08426818 | -2.7294011 | 0.00634495 | 0.04404454 |
| ENSG000002 | 13.4347314 | 6.93518627 | 2.54220905 | 2.72801573 | 0.00637166 | 0.04420809 |
| ENSG000002 | 14.2426003 | 7.01793737 | 2.57277079 | 2.72777405 | 0.00637633 | 0.04421865 |
| KALRN | 53.7939976 | -3.2725717 | 1.19998097 | -2.7271863 | 0.0063877 | 0.04427562 |
| FNBP1 | 533.423116 | 0.75251887 | 0.27606344 | 2.72589108 | 0.00641282 | 0.04440622 |
| BABAM1 | 808.160707 | -0.7546017 | 0.27682778 | -2.7258887 | 0.00641286 | 0.04440622 |
| SYPL2 | 25.4925085 | -2.4862235 | 0.91222004 | -2.7254647 | 0.0064211 | 0.04444138 |
| ARRB2 | 5727.06823 | 0.71247006 | 0.26150843 | 2.72446312 | 0.00644061 | 0.04454815 |
| CD300LB | 374.940962 | -1.8299156 | 0.67168959 | -2.7243471 | 0.00644288 | 0.04454815 |
| RASA3 | 1281.39618 | 0.91919924 | 0.33744331 | 2.72401087 | 0.00644944 | 0.04457159 |
| STAT5B | 427.811223 | -0.657934 | 0.24155867 | -2.7237025 | 0.00645546 | 0.04459127 |
| PUM2 | 917.90993 | 0.8533464 | 0.31337078 | 2.72312054 | 0.00646685 | 0.04464794 |
| MMP12 | 99.0059801 | -3.1619473 | 1.16274685 | -2.7193772 | 0.0065405 | 0.04513425 |
| USP6NL | 285.916572 | -0.8180457 | 0.30088008 | -2.7188431 | 0.00655107 | 0.04518499 |
| RBMXL1 | 135.756355 | 1.3050012 | 0.48005473 | 2.71844255 | 0.006559 | 0.04521753 |
| SYNGAP1 | 19.3637298 | -1.8784622 | 0.69122872 | -2.7175697 | 0.00657633 | 0.04531472 |
| ENSG000002 | 34.461447 | -2.6506333 | 0.97561585 | -2.7168822 | 0.00659001 | 0.04538669 |
| KCTD10 | 1217.11518 | 0.63378028 | 0.23335568 | 2.71594103 | 0.00660877 | 0.0454828 |
| ARHGEF40 | 678.839418 | 0.78506783 | 0.28906813 | 2.71585739 | 0.00661044 | 0.0454828 |
| SMC6 | 93.5500426 | -1.1757719 | 0.43297345 | -2.7155751 | 0.00661608 | 0.04548757 |
| HCST | 240.031642 | -1.3668939 | 0.50336763 | -2.7154983 | 0.00661761 | 0.04548757 |
| ST18 | 88.8420759 | 1.15953783 | 0.42706503 | 2.71513176 | 0.00662494 | 0.04551568 |
| SEZ6L | 8.10569082 | 4.73794795 | 1.74553477 | 2.71432459 | 0.00664111 | 0.04560443 |
| IFT140 | 56.3093176 | -1.3925521 | 0.51332526 | -2.7128065 | 0.00667161 | 0.04579147 |
| SFT2D1 | 465.621504 | 0.58510719 | 0.21584007 | 2.71083671 | 0.00671137 | 0.04601856 |
| SRRT | 732.170694 | 0.84105611 | 0.3102708 | 2.71071629 | 0.0067138 | 0.04601856 |
| IFI44 | 34.1219038 | -2.5352112 | 0.93526729 | -2.7106809 | 0.00671452 | 0.04601856 |
| SLC25A23 | 392.436634 | -1.4834297 | 0.54730044 | -2.7104485 | 0.00671923 | 0.04602835 |
| INO80B | 78.8860422 | -1.4600738 | 0.5387303 | -2.7102128 | 0.00672401 | 0.04603862 |
| LSM14B | 257.043956 | 0.75408206 | 0.27835455 | 2.70907035 | 0.0067472 | 0.04617494 |
| CLUAP1 | 52.5854758 | -1.3871221 | 0.51224255 | -2.7079399 | 0.00677023 | 0.04630994 |
| PBXIP1 | 572.022602 | 0.86556418 | 0.31979864 | 2.70659124 | 0.00679779 | 0.04645684 |
| ENSG000002 | 5.38829992 | 5.61619966 | 2.07502826 | 2.70656538 | 0.00679832 | 0.04645684 |
| GPR18 | 10.1374118 | 5.0396739 | 1.86230517 | 2.70614826 | 0.00680686 | 0.04647384 |
| BEST1 | 455.702636 | 0.78017996 | 0.28831722 | 2.70597768 | 0.00681036 | 0.04647384 |
| NOTCH2NLB | 67.4646438 | -2.0578621 | 0.76049256 | -2.7059595 | 0.00681074 | 0.04647384 |
| PAPSS2 | 359.885666 | 1.38341669 | 0.51140166 | 2.70514706 | 0.00682742 | 0.04656503 |
| TMEM104 | 455.309277 | -1.8668435 | 0.69019434 | -2.7048084 | 0.00683438 | 0.0465899 |
| YES1 | 202.010599 | 1.44128538 | 0.53311875 | 2.70349781 | 0.00686139 | 0.04675133 |
| EYA3 | 544.882701 | 1.06254929 | 0.39321864 | 2.7021844 | 0.00688855 | 0.04691366 |
| CLEC4A | 183.522808 | 2.01170502 | 0.74459233 | 2.70175361 | 0.00689749 | 0.04695172 |

| | | | | | | |
|---|---|---|---|---|---|---|
| SRSF8 | 293.050863 | -0.658647 | 0.24394807 | -2.6999475 | 0.00693504 | 0.04716417 |
| DEF8 | 492.10709 | -1.3025075 | 0.48242274 | -2.6999297 | 0.00693541 | 0.04716417 |
| ANAPC1 | 377.087938 | -0.8780113 | 0.32529277 | -2.6991417 | 0.00695186 | 0.04725311 |
| DDX42 | 828.813517 | 0.71494473 | 0.26490062 | 2.69891683 | 0.00695656 | 0.04726217 |
| DARS2 | 388.59659 | -1.1588909 | 0.4295201 | -2.6981063 | 0.00697352 | 0.04735449 |
| TCN1 | 24.2158443 | 2.80970354 | 1.04160059 | 2.69748651 | 0.00698651 | 0.04741567 |
| STK40 | 3785.57529 | 1.22179924 | 0.45296203 | 2.69735466 | 0.00698928 | 0.04741567 |
| KLHL6 | 435.269392 | -1.5314905 | 0.56783768 | -2.6970568 | 0.00699553 | 0.04743519 |
| CD14 | 14160.0561 | 1.85650426 | 0.68849905 | 2.69645142 | 0.00700826 | 0.04748819 |
| ZNF277 | 170.951071 | 0.91572047 | 0.33961315 | 2.69636343 | 0.00701011 | 0.04748819 |
| KCTD11 | 112.456506 | 0.94112972 | 0.34944579 | 2.69320659 | 0.00707684 | 0.0479171 |
| TPI1 | 5002.40644 | 0.55737658 | 0.20697209 | 2.69300363 | 0.00708115 | 0.04792317 |
| LAPTM4A | 1870.94301 | 0.68430171 | 0.25412944 | 2.69272894 | 0.00708699 | 0.04793957 |
| CAPN5 | 42.803741 | -2.0233801 | 0.75166969 | -2.6918474 | 0.00710574 | 0.04800549 |
| BOLA3 | 137.081784 | -1.2313953 | 0.45752236 | -2.6914429 | 0.00711437 | 0.04800549 |
| TAMALIN | 115.057223 | 1.65657602 | 0.6155073 | 2.69139949 | 0.00711529 | 0.04800549 |
| ZFYVE16 | 920.825037 | 0.98687909 | 0.36669447 | 2.69128439 | 0.00711775 | 0.04800549 |
| FA2H | 6.51130522 | -6.4018983 | 2.37891465 | -2.6911004 | 0.00712167 | 0.04800549 |
| ALPL | 6.51130522 | -6.4018983 | 2.37891465 | -2.6911004 | 0.00712167 | 0.04800549 |
| JDP2-AS1 | 22.269783 | 1.88109216 | 0.69900822 | 2.69108731 | 0.00712196 | 0.04800549 |
| SPEG | 39.2748565 | -8.9961526 | 3.343067 | -2.6909878 | 0.00712408 | 0.04800549 |
| NUDT7 | 11.1158836 | -2.8747385 | 1.06845046 | -2.6905679 | 0.00713305 | 0.0480429 |
| CNOT6L | 453.582751 | 0.74859254 | 0.27824574 | 2.69040076 | 0.00713663 | 0.04804393 |
| EVL | 1801.3979 | -1.1928051 | 0.44341604 | -2.6900359 | 0.00714443 | 0.04807345 |
| ZNRF1 | 36.5596444 | -1.8550261 | 0.68967083 | -2.6897267 | 0.00715106 | 0.04809496 |
| LINC02910 | 20.3263173 | 2.50836913 | 0.9327079 | 2.68934049 | 0.00715934 | 0.04812759 |
| TBK1 | 268.691246 | 0.73138182 | 0.27199275 | 2.68897537 | 0.00716717 | 0.04813636 |
| CCDC28B | 15.5518732 | -2.2285003 | 0.82879449 | -2.6888455 | 0.00716996 | 0.04813636 |
| CCDC112 | 81.3684595 | 1.48622618 | 0.55274688 | 2.68880066 | 0.00717092 | 0.04813636 |
| SEC14L2 | 30.4887495 | -3.0407055 | 1.13097275 | -2.6885754 | 0.00717576 | 0.04814584 |
| ZNF440 | 46.4858377 | 1.53791904 | 0.57230762 | 2.68722448 | 0.00720485 | 0.04831792 |
| KCTD9 | 265.74553 | -0.8296984 | 0.30898162 | -2.6852678 | 0.00724717 | 0.04857258 |
| SLC10A3 | 420.51581 | 0.72417958 | 0.26969807 | 2.68514926 | 0.00724974 | 0.04857258 |
| ZNF106 | 1201.24369 | 0.67638086 | 0.25192027 | 2.68490053 | 0.00725514 | 0.04858556 |
| SORBS3 | 467.24396 | -0.7948327 | 0.29606165 | -2.6846865 | 0.00725979 | 0.04859351 |
| ENSG000002 | 30.9292279 | -1.4870598 | 0.55446467 | -2.6819739 | 0.00731892 | 0.04896313 |
| CYB5D1 | 57.4474094 | -1.2989519 | 0.4843521 | -2.6818339 | 0.00732198 | 0.04896313 |
| ENSG000002 | 14.0126987 | 6.9934531 | 2.60788853 | 2.68165339 | 0.00732593 | 0.04896624 |
| PECAM1 | 2753.13191 | 1.52738617 | 0.56973997 | 2.68084785 | 0.00734359 | 0.04906091 |
| SLC26A11 | 535.988275 | -2.1115396 | 0.78826504 | -2.6787179 | 0.00739046 | 0.04935058 |
| PRKCB | 824.395399 | -1.7704426 | 0.66100379 | -2.6784152 | 0.00739715 | 0.04937172 |
| PLEKHG4 | 22.46417 | -2.377234 | 0.88850961 | -2.6755299 | 0.00746112 | 0.04977508 |
| FRMD4A | 796.339329 | 0.84896563 | 0.31734226 | 2.6752366 | 0.00746765 | 0.04977696 |
| C1S | 10.1776554 | -4.0839744 | 1.52660591 | -2.6751989 | 0.00746849 | 0.04977696 |
| NEK9 | 520.687811 | 0.67512191 | 0.25240204 | 2.67478787 | 0.00747765 | 0.04981438 |
| ARF6 | 1706.08603 | 0.88739647 | 0.33183194 | 2.67423465 | 0.00749 | 0.04987297 |
| IL21R | 108.764826 | -1.3282341 | 0.49676104 | -2.6737888 | 0.00749997 | 0.04991566 |
| LILRB2 | 816.825544 | 1.66751362 | 0.6237462 | 2.6733848 | 0.00750901 | 0.04995214 |

2793e-74
92827e-49
45922e-49
95269e-39
13795e-38
26233e-35
25846e-35
09615e-33
58727e-31
24081e-31
36655e-31
71285e-30
07369e-30
15954e-29
95888e-29
4371e-29
27059e-27
32559e-26
22321e-26
17453e-25
63216e-25
48071e-24
08242e-24
1601e-23
92914e-23
79159e-22
74472e-22
54573e-22
57425e-22
16895e-21
20925e-21
27614e-21
06338e-20
25618e-20
28997e-20
52089e-19
54191e-19
65129e-19
19148e-19
18042e-19
74942e-19
76897e-18
47648e-18
63016e-18
96866e-17
30447e-17
25812e-17

60834e-17
3736e-17
2876e-17
85003e-17
99985e-17
25485e-17
58118e-17
6984e-17
96371e-16
48881e-16
62203e-16
62431e-16
73357e-16
59313e-16
81925e-16
61443e-16
58379e-16
2179e-16
8303e-16
8303e-16
56788e-15
44487e-15
95275e-15
59048e-15
43383e-15
61886e-15
39169e-15
89456e-15
77462e-15
91508e-15
91508e-15
88093e-15
89132e-15
28177e-15
54904e-15
8514e-14
25851e-14
56908e-14
73816e-14
23729e-14
78129e-13
68809e-13
44757e-13
47035e-13
57616e-13
38606e-13
02364e-13
45424e-13

08658e-13
07212e-13
07489e-13
82129e-13
0573e-13
77784e-13
77784e-13
57835e-13
52265e-13
96722e-13
71324e-13
8089e-13
33697e-13
35838e-13
63427e-13
83555e-12
83555e-12
72211e-12
30232e-12
91553e-12
39246e-12
74312e-12
47552e-12
63707e-12
00969e-12
72219e-12
33831e-12
68358e-12
4283e-12
03839e-12
55526e-12
19002e-12
9952e-12
66146e-12
53003e-11
21499e-11
43931e-11
40583e-11
76716e-11
06565e-11
37701e-11
86277e-11
8649e-11
07236e-11
12307e-11
63788e-11
68541e-11
01709e-11

87991e-11
43437e-11
28216e-11
93963e-11
01408e-11
36344e-10
84547e-10
30805e-10
30781e-10
15454e-10
53489e-10
71039e-10
75742e-10
69957e-10
21629e-10
96577e-10
10154e-10
58848e-10
67188e-10
0056e-10
75036e-10
69417e-10
90546e-10
34149e-10
43658e-10
64111e-10
66983e-10
0022e-10
76318e-10
95674e-10
5069e-10
08723e-10
32166e-10
63229e-10
06968e-09
37994e-09
60063e-09
8457e-09
20826e-09
20826e-09
20826e-09
39576e-09
58865e-09
84498e-09
46217e-09
51479e-09
03642e-09
03642e-09

03642e-09
47769e-09
89198e-09
89198e-09
36238e-09
27853e-09
16653e-09
23739e-09
10673e-09
37321e-09
52967e-09
25514e-09
66761e-09
38736e-09
18667e-09
62496e-09
88413e-09
22543e-09
22543e-09
22543e-09
29793e-09
66029e-09
37054e-09
02889e-09
90513e-08
84777e-08
65373e-08
46618e-08
14222e-08
02582e-08
92938e-08
41744e-08
34361e-08
27161e-08
38401e-08
45411e-08
00703e-08
00703e-08
66233e-08
19405e-08
21215e-08
75822e-08
39052e-08
0829e-08
23803e-08
17255e-08
2532e-08
64299e-08

15498e-08
52029e-08
7843e-08
46278e-08
83304e-08
83304e-08
6124e-08
61617e-08
84469e-08
83295e-08
55005e-08
79119e-08
81925e-08
07277e-08
8099e-08
95073e-08
55528e-08
49325e-08
02907e-08
12372e-08
73631e-07
2292e-07
64913e-07
95403e-07
72276e-07
45059e-07
64702e-07
64702e-07
64702e-07
00577e-07
1544e-07
88548e-07
08625e-07
08625e-07
08625e-07
77859e-07
64281e-07
60614e-07
59351e-07
35805e-07
35805e-07
99191e-07
78592e-07
71027e-07
52036e-07
62029e-07
95023e-07
95023e-07

31495e-07
08793e-07
08793e-07
75039e-07
85185e-07
23609e-07
31859e-07
04741e-07
79977e-07
45372e-07
50838e-07
50838e-07
21397e-07
79056e-07
13502e-07
84782e-07
81368e-07
85767e-07
82714e-07
11714e-07
42179e-07
97026e-07
74464e-07
61521e-07
75103e-07
76541e-07
38866e-07
90174e-07
90174e-07
28534e-07
48237e-07
06663e-07
06663e-07
00845e-07
84581e-07
40222e-07
10425e-07
52635e-07
21989e-06
57903e-06
1603e-06
92989e-06
92989e-06
84528e-06
72716e-06
48595e-06
29633e-06
64233e-06

84649e-06
86906e-06
73417e-06
86301e-06
68914e-06
06494e-06
63287e-06
38317e-06
66066e-06
5623e-06
17909e-06
40886e-06
63611e-06
6128e-06
69532e-06
03345e-06
44395e-06
78788e-06
41431e-06
4833e-06
09418e-06
04781e-06
40913e-06
77277e-06
88178e-06
88178e-06
49677e-06
8201e-06
60777e-06
60777e-06
60777e-06
80751e-06
20731e-06
67664e-06
36908e-06
36908e-06
7426e-06
83073e-06
31769e-06
77695e-06
30893e-06
64752e-06
76135e-06
43782e-06
42257e-06
89028e-06
87527e-06
86659e-06

88343e-06
85576e-06
75851e-06
75851e-06
60789e-06
60789e-06
01905e-06
24976e-06
44419e-06
26628e-06
33823e-06
33823e-06
37403e-06
68768e-06
51077e-06
9527e-06
45201e-06
6372e-06
11464e-06
01657e-06
64643e-06
78531e-06
25115e-06
08734e-06
62376e-06
12589e-06
81778e-06
79102e-06
77674e-06
57853e-06
59334e-06
28807e-06
60509e-06
39541e-06
50395e-06
63674e-06
94877e-05
88818e-05
6375e-05
6375e-05
97177e-05
73503e-05
29724e-05
29488e-05
57322e-05
65715e-05
096e-05
04506e-05

05185e-05
75484e-05
79486e-05
88604e-05
69705e-05
69705e-05
49501e-05
48172e-05
72701e-05
40888e-05
64185e-05
53207e-05
53207e-05
73498e-05
64554e-05
78036e-05
00016e-05
00016e-05
26533e-05
95183e-05
21241e-05
71392e-05
71392e-05
71392e-05
43589e-05
61705e-05
56441e-05
16215e-05
8803e-05
77437e-05
10278e-05
52555e-05
63133e-05
55474e-05
74723e-05
22669e-05
2136e-05
93825e-05
65854e-05
58029e-05
26869e-05
87818e-05
50092e-05
05187e-05
27764e-05
97493e-05
83499e-05
63409e-05

04004e-05
04004e-05
65161e-05
41033e-05
78623e-05
78623e-05
88865e-05
00273e-05
56024e-05
10338e-05
21424e-05
086e-05
65394e-05
1242e-05
8548e-05
1398e-05
07396e-05
71373e-05
62737e-05
77719e-05
12204e-05
6258e-05
28095e-05
28095e-05
03037e-05
24353e-05
79403e-05
79403e-05
60251e-05
40931e-05
70786e-05
00633e-05
91746e-05
348e-05
23548e-05
92749e-05
65201e-05
34373e-05
40324e-05
92483e-05
84908e-05
5377e-05
35459e-05
39278e-05
75592e-05
70141e-05
84733e-05
88342e-05

88342e-05
7747e-05
17244e-05
98756e-05
77593e-05
92887e-05
97546e-05
38822e-05
03582e-05
03582e-05
13226e-05
7716e-05
15009e-05
8327e-05
8327e-05
8327e-05
69818e-05
23793e-05
29654e-05
52588e-05
52927e-05
30256e-05
624e-05
40388e-05
16861e-05
16861e-05
16861e-05
67215e-05
67215e-05
22468e-05
02609e-05
02609e-05
77209e-05
77209e-05
85901e-05
85901e-05
50622e-05
