## Supplementary material for "Syngeneic human iPSC Macrophages and Microglia possess distinct neuroinflammatory profiles, responses to HIV infection, and ART treatment": Table S8

**Table 5: GC**

| Name | Term | ES | NES | NOM p-val | FDR q-val |
|---|---|---|---|---|---|
| **prerank** | adenylyltransferase activity (GO:0070566) | -0.8069334 | -1.6459907 | 0.00162866 | 0.55323379 |
| **prerank** | chemoattractant activity (GO:0042056) | -0.7663679 | -1.5925175 | 0.01034483 | 0.67471106 |
| **prerank** | glucuronosyltransferase activity (GO:0015020) | -0.7970167 | -1.5805284 | 0.00695652 | 0.54965615 |
| **prerank** | single-stranded DNA helicase activity (GO:0017116) | 0.77317136 | 1.57984015 | 0.00757576 | 1 |
| **prerank** | calcium channel regulator activity (GO:0005246) | -0.7323567 | -1.5782002 | 0.00801282 | 0.42785365 |
| **prerank** | oxidoreductase activity, acting on paired donors, with incorporation or reduction of molecular oxygen, NAD(P)H as one donor, and incorporation of one atom of oxygen (GO:0016709) | -0.7316426 | -1.5637034 | 0.01022147 | 0.42287747 |
| **prerank** | neuropeptide receptor activity (GO:0008188) | -0.7330992 | -1.5533159 | 0.01522843 | 0.41468142 |
| **prerank** | protease binding (GO:0002020) | -0.5985841 | -1.541954 | 0.00145773 | 0.41370569 |
| **prerank** | phospholipase A2 activity (GO:0004623) | -0.7554685 | -1.5408131 | 0.01367521 | 0.36796877 |
| **prerank** | exopeptidase activity (GO:0008238) | -0.6546825 | -1.5361767 | 0.01263823 | 0.34779032 |
| **prerank** | histone methyltransferase activity (H3-K4 specific) (GO:0042800) | 0.7818171 | 1.53434204 | 0.02669903 | 1 |
| **prerank** | endopeptidase activity (GO:0004175) | -0.5382045 | -1.5237137 | 0 | 0.37418899 |
| **prerank** | NAD-retinol dehydrogenase activity (GO:0004745) | -0.7680029 | -1.5035021 | 0.01715266 | 0.4420016 |
| **prerank** | translation initiation factor activity (GO:0003743) | 0.62689351 | 1.49795368 | 0.01897019 | 1 |
| **prerank** | arachidonic acid monooxygenase activity (GO:0008391) | 0.75078264 | 1.4979226 | 0.0443459 | 1 |
| **prerank** | NAD+ ADP-ribosyltransferase activity (GO:0003950) | -0.7161462 | -1.4880704 | 0.01050788 | 0.48908014 |
| **prerank** | cysteine-type endopeptidase activity (GO:0004197) | -0.5927975 | -1.4880098 | 0.00738552 | 0.45205903 |

| | | | | | |
|---|---|---|---|---|---|
| **prerank** | G protein-coupled receptor activity (GO:0004930) | 0.47444997 | 1.48125295 | 0 | 1 |
| **prerank** | heme binding (GO:0020037) | -0.590523 | -1.4786289 | 0.01659125 | 0.46590677 |
| **prerank** | cholesterol transfer activity (GO:0120020) | -0.755856 | -1.4752972 | 0.03672788 | 0.45143357 |
| **prerank** | wide pore channel activity (GO:0022829) | -0.7657992 | -1.4732588 | 0.02139037 | 0.4332811 |
| **prerank** | metalloendopeptidase activity (GO:0004222) | -0.5898184 | -1.4728155 | 0.00902256 | 0.40940104 |
| **prerank** | nucleosomal DNA binding (GO:0031492) | 0.62945069 | 1.47185928 | 0.01785714 | 1 |
| **prerank** | DNA replication origin binding (GO:0003688) | 0.66249919 | 1.47071198 | 0.05882353 | 0.89463062 |
| **prerank** | gap junction channel activity (GO:0005243) | -0.7701928 | -1.4652915 | 0.02473498 | 0.42026471 |
| **prerank** | nuclear import signal receptor activity (GO:0061608) | 0.72308713 | 1.45037146 | 0.0744186 | 0.93325565 |
| **prerank** | sterol transfer activity (GO:0120015) | -0.7308682 | -1.4454623 | 0.04746318 | 0.49248232 |
| **prerank** | olfactory receptor activity (GO:0004984) | -0.5802296 | -1.4426796 | 0.0123839 | 0.48039622 |
| **prerank** | GTP binding (GO:0005525) | -0.525864 | -1.4404944 | 0 | 0.46653402 |
| **prerank** | microtubule motor activity (GO:0003777) | 0.550953 | 1.43652012 | 0.02762431 | 0.92968991 |
| **prerank** | phosphatidylinositol monophosphate phosphatase activity (GO:0052744) | 0.70795562 | 1.43424698 | 0.07226107 | 0.85220806 |
| **prerank** | phospholipase activity (GO:0004620) | -0.5890993 | -1.4314613 | 0.03007519 | 0.48808224 |
| **prerank** | guanyl ribonucleotide binding (GO:0032561) | -0.5166952 | -1.4260579 | 0.00133869 | 0.49104219 |
| **prerank** | ferrous iron binding (GO:0008198) | -0.6978342 | -1.4250457 | 0.05932203 | 0.47542005 |

| | | | | | |
|---|---|---|---|---|---|
| **prerank** | monocarboxylic acid binding (GO:0033293) | 0.65928729 | 1.42240547 | 0.06030151 | 0.84626455 |
| **prerank** | carboxylic ester hydrolase activity (GO:0052689) | -0.5578339 | -1.4081367 | 0.02435312 | 0.53067513 |
| **prerank** | mannose binding (GO:0005537) | 0.70462766 | 1.40559146 | 0.0921659 | 0.87526552 |
| **prerank** | GTPase activity (GO:0003924) | -0.507905 | -1.3994404 | 0.00398406 | 0.54899316 |
| **prerank** | nucleoside-triphosphatase activity (GO:0017111) | -0.4930803 | -1.3982226 | 0.00254453 | 0.53447827 |
| **prerank** | phosphatidylinositol-3-phosphatase activity (GO:0004438) | 0.7141594 | 1.39393429 | 0.07962529 | 0.88085565 |
| **prerank** | 3'-5' DNA helicase activity (GO:0043138) | 0.68422291 | 1.39122178 | 0.08372093 | 0.83456787 |
| **prerank** | glucose transmembrane transporter activity (GO:0005355) | -0.6938133 | -1.3867154 | 0.05574324 | 0.56912411 |
| **prerank** | pentosyltransferase activity (GO:0016763) | -0.6262429 | -1.3834749 | 0.05025126 | 0.56477422 |
| **prerank** | double-stranded RNA binding (GO:0003725) | -0.5690649 | -1.3824362 | 0.03095975 | 0.5505343 |
| **prerank** | intracellular calcium activated chloride channel activity (GO:0005229) | -0.7225744 | -1.3811761 | 0.0658363 | 0.52091304 |
| **prerank** | intracellular chloride channel activity (GO:0061778) | -0.7225744 | -1.3811761 | 0.0658363 | 0.52091304 |
| **prerank** | metallopeptidase activity (GO:0008237) | -0.5281685 | -1.3808128 | 0.02513966 | 0.50678356 |
| **prerank** | oxidoreductase activity, acting on the CH-OH group of donors, NAD or NADP as acceptor (GO:0016616) | -0.5411787 | -1.3795859 | 0.03953148 | 0.4963837 |
| **prerank** | protein phosphatase 2A binding (GO:0051721) | -0.6757882 | -1.3730277 | 0.06112054 | 0.50927059 |
| **prerank** | calcium-release channel activity (GO:0015278) | 0.66929883 | 1.3719258 | 0.0959596 | 0.89042351 |

| | | | | | |
|---|---|---|---|---|---|
| **prerank** | hydrolase activity, acting on carbon-nitrogen (but not peptide) bonds, in cyclic amidines (GO:0016814) | -0.6914873 | -1.3684775 | 0.10544815 | 0.51398813 |
| **prerank** | cysteine-type peptidase activity (GO:0008234) | -0.5491931 | -1.3653876 | 0.03851852 | 0.51233263 |
| **prerank** | steroid dehydrogenase activity, acting on the CH-OH group of donors, NAD or NADP as acceptor (GO:0033764) | -0.6992617 | -1.358838 | 0.09185442 | 0.52563116 |
| **prerank** | DNA-directed 5'-3' RNA polymerase activity (GO:0003899) | 0.60826112 | 1.35748726 | 0.07560976 | 0.92273168 |
| **prerank** | hydrolase activity, hydrolyzing O-glycosyl compounds (GO:0004553) | -0.6234754 | -1.3561828 | 0.08390411 | 0.52413729 |
| **prerank** | growth factor activity (GO:0008083) | -0.5452391 | -1.3556711 | 0.04322767 | 0.51291212 |
| **prerank** | protein-cysteine S-palmitoyltransferase activity (GO:0019706) | -0.6417196 | -1.3514063 | 0.07986111 | 0.51722731 |
| **prerank** | cholesterol binding (GO:0015485) | -0.5805884 | -1.3499085 | 0.08210181 | 0.51088286 |
| **prerank** | G protein-coupled chemoattractant receptor activity (GO:0001637) | 0.66745935 | 1.34912373 | 0.09692671 | 0.91626597 |
| **prerank** | ATPase activity, coupled to transmembrane movement of ions, rotational mechanism (GO:0044769) | -0.7045308 | -1.3478819 | 0.11643836 | 0.49540015 |
| **prerank** | proton-transporting ATPase activity, rotational mechanism (GO:0046961) | -0.7045308 | -1.3478819 | 0.11643836 | 0.49540015 |
| **prerank** | G protein-coupled peptide receptor activity (GO:0008528) | 0.49875168 | 1.3451865 | 0.05649718 | 0.88747853 |
| **prerank** | histone acetyltransferase activity (GO:0004402) | 0.59444613 | 1.34399388 | 0.10079576 | 0.84645851 |
| **prerank** | receptor ligand activity (GO:0048018) | -0.4718629 | -1.3430478 | 0.01898734 | 0.50368885 |

| | | | | | |
|---|---|---|---|---|---|
| **prerank** | mRNA binding (GO:0003729) | 0.4215708 | 1.34219608 | 0 | 0.81276264 |
| **prerank** | myosin V binding (GO:0031489) | -0.7146945 | -1.3335486 | 0.11971831 | 0.52928621 |
| **prerank** | phosphatidylcholine transporter activity (GO:0008525) | -0.6775104 | -1.3332954 | 0.10915493 | 0.51910432 |
| **prerank** | O-acyltransferase activity (GO:0008374) | -0.6261606 | -1.3287907 | 0.09171076 | 0.52489723 |
| **prerank** | miRNA binding (GO:0035198) | 0.57291756 | 1.32193197 | 0.10843373 | 0.87824762 |
| **prerank** | GTPase regulator activity (GO:0030695) | 0.42811433 | 1.32048456 | 0.01265823 | 0.84607557 |
| **prerank** | transcription coactivator binding (GO:0001223) | -0.6761799 | -1.3195988 | 0.10720562 | 0.54905213 |
| **prerank** | tumor necrosis factor receptor superfamily binding (GO:0032813) | -0.6358043 | -1.3127452 | 0.11794872 | 0.56418138 |
| **prerank** | 1-phosphatidylinositol binding (GO:0005545) | -0.6889953 | -1.3094448 | 0.13913043 | 0.56616688 |
| **prerank** | protein phosphatase inhibitor activity (GO:0004864) | -0.6253129 | -1.3085107 | 0.09983633 | 0.55901306 |
| **prerank** | histone-lysine N-methyltransferase activity (GO:0018024) | 0.52843808 | 1.30691313 | 0.08222812 | 0.88028554 |
| **prerank** | oxidoreductase activity, acting on NAD(P)H, oxygen as acceptor (GO:0050664) | 0.65126327 | 1.30374101 | 0.13163972 | 0.86093039 |
| **prerank** | acylglycerol O-acyltransferase activity (GO:0016411) | -0.6418102 | -1.3030989 | 0.11785714 | 0.56812735 |

| | | | | | |
|---|---|---|---|---|---|
| **prerank** | actin binding (GO:0003779) | -0.4791215 | -1.3012551 | 0.03961749 | 0.56425583 |
| **prerank** | alpha-tubulin binding (GO:0043014) | 0.56426395 | 1.30019014 | 0.1031941 | 0.84412149 |
| **prerank** | cell adhesion mediator activity (GO:0098631) | -0.6230311 | -1.2998257 | 0.14802065 | 0.55933649 |
| **prerank** | 5'-3' RNA polymerase activity (GO:0034062) | 0.58589741 | 1.29677061 | 0.12339332 | 0.8287255 |
| **prerank** | N-acyltransferase activity (GO:0016410) | -0.5863549 | -1.2923046 | 0.11348684 | 0.57750485 |
| **prerank** | phosphotransferase activity, phosphate group as acceptor (GO:0016776) | -0.6278411 | -1.2908902 | 0.1559322 | 0.57322752 |
| **prerank** | neuropeptide hormone activity (GO:0005184) | -0.6587892 | -1.2901004 | 0.11992945 | 0.56618733 |
| **prerank** | palmitoyltransferase activity (GO:0016409) | -0.5993889 | -1.284928 | 0.11744387 | 0.57466658 |
| **prerank** | ubiquitin-like protein ligase activity (GO:0061659) | -0.4539858 | -1.2806827 | 0.04107831 | 0.58076539 |
| **prerank** | ATPase binding (GO:0051117) | -0.5251254 | -1.2806354 | 0.09146341 | 0.57143659 |
| **prerank** | sterol binding (GO:0032934) | -0.5397197 | -1.2793207 | 0.1136 | 0.56676799 |
| **prerank** | ubiquitin-like protein ligase binding (GO:0044389) | -0.453163 | -1.2755302 | 0.02634881 | 0.57152469 |
| **prerank** | ubiquitin protein ligase binding (GO:0031625) | -0.453682 | -1.2734969 | 0.03620474 | 0.56951614 |
| **prerank** | phosphatidylinositol kinase activity (GO:0052742) | 0.66279293 | 1.2693952 | 0.17703349 | 0.93134526 |
| **prerank** | phospholipid transfer activity (GO:0120014) | -0.6308812 | -1.2692174 | 0.15608919 | 0.57689128 |

| | | | | | |
|---|---|---|---|---|---|
| **prerank** | retinoid binding (GO:0005501) | -0.6180645 | -1.2691022 | 0.17102967 | 0.56846095 |
| **prerank** | GTPase activator activity (GO:0005096) | 0.39788578 | 1.2684842 | 0.01522843 | 0.90257465 |
| **prerank** | GDP binding (GO:0019003) | -0.5198393 | -1.2625216 | 0.125 | 0.58268018 |
| **prerank** | telomeric DNA binding (GO:0042162) | 0.54270973 | 1.25414134 | 0.15024631 | 0.94180834 |
| **prerank** | ubiquitin protein ligase activity (GO:0061630) | -0.4461139 | -1.2515412 | 0.04587156 | 0.61380024 |
| **prerank** | neuropeptide receptor binding (GO:0071855) | -0.6505002 | -1.2512202 | 0.19642857 | 0.60617726 |
| **prerank** | iron ion binding (GO:0005506) | -0.5355153 | -1.2511498 | 0.14082278 | 0.59786606 |
| **prerank** | death receptor binding (GO:0005123) | -0.6702331 | -1.2487772 | 0.17753623 | 0.59859795 |
| **prerank** | microtubule binding (GO:0008017) | 0.40211526 | 1.24825324 | 0.032 | 0.93948342 |
| **prerank** | cytokine activity (GO:0005125) | -0.4622618 | -1.244804 | 0.08423913 | 0.60399839 |
| **prerank** | chemokine binding (GO:0019956) | 0.53644967 | 1.24455975 | 0.17142857 | 0.92724973 |
| **prerank** | adrenergic receptor binding (GO:0031690) | -0.6580146 | -1.2420824 | 0.1975089 | 0.60550838 |
| **prerank** | acyltransferase activity, transferring groups other than amino-acyl groups (GO:0016747) | -0.5017387 | -1.2402516 | 0.13888889 | 0.60370758 |
| **prerank** | 3'-5' exonuclease activity (GO:0008408) | -0.5411222 | -1.2392423 | 0.15566038 | 0.59940491 |

| | | | | | |
|---|---|---|---|---|---|
| **prerank** | RNA polymerase II-specific DNA-binding transcription factor binding (GO:0061629) | -0.4495141 | -1.2367853 | 0.06451613 | 0.60008124 |
| **prerank** | phosphatase binding (GO:0019902) | -0.4767131 | -1.2367561 | 0.11160059 | 0.59231332 |
| **prerank** | nucleobase-containing compound kinase activity (GO:0019205) | -0.6451997 | -1.2362723 | 0.1956912 | 0.58652089 |
| **prerank** | regulatory RNA binding (GO:0061980) | 0.50257222 | 1.23098132 | 0.15424165 | 0.96411298 |
| **prerank** | protein serine/threonine kinase inhibitor activity (GO:0030291) | -0.5738134 | -1.2304846 | 0.16498316 | 0.59929029 |
| **prerank** | phosphatidylinositol phospholipase C activity (GO:0004435) | 0.56457357 | 1.22966431 | 0.17777778 | 0.94138584 |
| **prerank** | chemokine receptor activity (GO:0004950) | 0.54869718 | 1.2291567 | 0.17821782 | 0.91641882 |
| **prerank** | histone deacetylase binding (GO:0042826) | 0.44701959 | 1.22876462 | 0.11515152 | 0.89201726 |
| **prerank** | nucleoside monophosphate kinase activity (GO:0050145) | -0.6200311 | -1.2280781 | 0.21206897 | 0.60001962 |
| **prerank** | protein phosphatase binding (GO:0019903) | -0.465469 | -1.2273197 | 0.11637931 | 0.59512957 |
| **prerank** | ubiquitin-like protein-specific protease activity (GO:0019783) | 0.59377649 | 1.22431554 | 0.18965517 | 0.88684334 |
| **prerank** | lysophospholipase activity (GO:0004622) | -0.6129573 | -1.2223206 | 0.22259696 | 0.60476852 |
| **prerank** | sulfotransferase activity (GO:0008146) | -0.552434 | -1.2213583 | 0.16666667 | 0.6005739 |
| **prerank** | cysteine-type endopeptidase inhibitor activity (GO:0004869) | 0.50184422 | 1.21943788 | 0.147343 | 0.8857696 |
| **prerank** | RNA polymerase binding (GO:0070063) | 0.57446226 | 1.2166265 | 0.21788991 | 0.87378391 |
| **prerank** | lipase activity (GO:0016298) | -0.5263617 | -1.2144012 | 0.16694491 | 0.61725968 |
| **prerank** | methylation-dependent protein binding (GO:0140034) | 0.46607556 | 1.21395797 | 0.14730878 | 0.86299921 |

| | | | | | |
|---|---|---|---|---|---|
| **prerank** | purine ribonucleoside triphosphate binding (GO:0035639) | -0.4147045 | -1.2118235 | 0.04316547 | 0.61934176 |
| **prerank** | motor activity (GO:0003774) | 0.45496136 | 1.21150307 | 0.13896458 | 0.852312 |
| **prerank** | transcription coregulator binding (GO:0001221) | -0.5206323 | -1.2083336 | 0.19224806 | 0.62438201 |
| **prerank** | methylated histone binding (GO:0035064) | 0.46216794 | 1.20806002 | 0.15428571 | 0.84546713 |
| **prerank** | cytokine receptor binding (GO:0005126) | -0.4808806 | -1.2075682 | 0.16155989 | 0.61962768 |
| **prerank** | histone demethylase activity (GO:0032452) | 0.53712026 | 1.20650297 | 0.2044335 | 0.83172188 |
| **prerank** | guanyl-nucleotide exchange factor activity (GO:0005085) | 0.40770419 | 1.20368053 | 0.09491525 | 0.82429641 |
| **prerank** | CoA hydrolase activity (GO:0016289) | -0.6355376 | -1.1966442 | 0.25473322 | 0.65022933 |
| **prerank** | protein tyrosine/serine/threonine phosphatase activity (GO:0008138) | -0.5994582 | -1.1965255 | 0.24915254 | 0.6433948 |
| **prerank** | nuclease activity (GO:0004518) | 0.46208172 | 1.19386017 | 0.14540816 | 0.84411487 |
| **prerank** | growth factor receptor binding (GO:0070851) | -0.4664398 | -1.1877821 | 0.13931034 | 0.66623312 |
| **prerank** | acyl-CoA hydrolase activity (GO:0047617) | -0.6074409 | -1.1877541 | 0.23865546 | 0.65904055 |
| **prerank** | inorganic cation transmembrane transporter activity (GO:0022890) | 0.46796007 | 1.18224026 | 0.18592965 | 0.87351586 |
| **prerank** | signaling receptor complex adaptor activity (GO:0030159) | -0.5313175 | -1.1817187 | 0.20813008 | 0.6729187 |

| | | | | | |
|---|---|---|---|---|---|
| **prerank** | alpha-actinin binding (GO:0051393) | -0.579966 | -1.1810992 | 0.22926829 | 0.66772888 |
| **prerank** | acetylcholine receptor activity (GO:0015464) | -0.5892238 | -1.1765588 | 0.25357143 | 0.6764549 |
| **prerank** | alcohol dehydrogenase (NADP+) activity (GO:0008106) | 0.5749435 | 1.17644353 | 0.24611973 | 0.87843269 |
| **prerank** | lysophosphatidic acid acyltransferase activity (GO:0042171) | -0.6165195 | -1.1763062 | 0.28326996 | 0.67031004 |
| **prerank** | ubiquitin-protein transferase activity (GO:0004842) | -0.4038306 | -1.1724586 | 0.0973236 | 0.67675397 |
| **prerank** | delayed rectifier potassium channel activity (GO:0005251) | 0.50725681 | 1.17162198 | 0.23469388 | 0.8800442 |
| **prerank** | hydro-lyase activity (GO:0016836) | 0.45796206 | 1.17083449 | 0.20789474 | 0.86510967 |
| **prerank** | SH2 domain binding (GO:0042169) | 0.50319141 | 1.17036979 | 0.21635884 | 0.84923621 |
| **prerank** | lipoprotein particle binding (GO:0071813) | -0.5799258 | -1.1684699 | 0.25255973 | 0.68342716 |
| **prerank** | DNA-binding transcription factor binding (GO:0140297) | -0.4210934 | -1.1652353 | 0.12890625 | 0.68765429 |
| **prerank** | CCR chemokine receptor binding (GO:0048020) | -0.5425815 | -1.1622902 | 0.25680272 | 0.69071189 |
| **prerank** | chloride transmembrane transporter activity (GO:0015108) | 0.5793044 | 1.160483 | 0.27186761 | 0.87251803 |
| **prerank** | Hsp90 protein binding (GO:0051879) | -0.5245901 | -1.1601639 | 0.23657718 | 0.69140564 |
| **prerank** | ribonuclease activity (GO:0004540) | 0.44269253 | 1.15507103 | 0.20168067 | 0.87811369 |
| **prerank** | DNA-binding transcription activator activity, RNA polymerase II-specific (GO:0001228) | -0.40186 | -1.1543397 | 0.124533 | 0.70417107 |
| **prerank** | hydrolase activity, acting on carbon-nitrogen (but not peptide) bonds, in linear amides (GO:0016811) | -0.4875448 | -1.1522445 | 0.23923445 | 0.70455664 |
| **prerank** | leak channel activity (GO:0022840) | 0.56956166 | 1.14645117 | 0.27625571 | 0.89592702 |

| | | | | | |
|---|---|---|---|---|---|
| **prerank** | cytokine receptor activity (GO:0004896) | 0.41131003 | 1.14496449 | 0.17073171 | 0.88429821 |
| **prerank** | oxidoreductase activity, acting on NAD(P)H, quinone or similar compound as acceptor (GO:0016655) | -0.6023236 | -1.1431451 | 0.30427046 | 0.72931827 |
| **prerank** | ubiquitin binding (GO:0043130) | -0.4641004 | -1.1426856 | 0.22272048 | 0.72387238 |
| **prerank** | protein kinase binding (GO:0019901) | -0.3898411 | -1.1425797 | 0.11606096 | 0.71737794 |
| **prerank** | SH3 domain binding (GO:0017124) | -0.482407 | -1.1413805 | 0.24592593 | 0.71496415 |
| **prerank** | calcium ion transmembrane transporter activity (GO:0015085) | 0.4375108 | 1.14108866 | 0.2122905 | 0.88332091 |
| **prerank** | myosin binding (GO:0017022) | -0.4841202 | -1.1401288 | 0.24689441 | 0.71246781 |
| **prerank** | bHLH transcription factor binding (GO:0043425) | -0.5641448 | -1.1379912 | 0.29021559 | 0.71327914 |
| **prerank** | ion channel activity (GO:0005216) | 0.41515317 | 1.13633894 | 0.20175439 | 0.88564854 |
| **prerank** | chloride channel activity (GO:0005254) | -0.4702818 | -1.1360891 | 0.23739837 | 0.71316252 |
| **prerank** | aminopeptidase activity (GO:0004177) | -0.5395879 | -1.1353444 | 0.28397213 | 0.7090653 |
| **prerank** | cation channel activity (GO:0005261) | 0.40698369 | 1.13294164 | 0.19384615 | 0.88386536 |
| **prerank** | chemokine activity (GO:0008009) | -0.5040465 | -1.132773 | 0.27244094 | 0.71115116 |
| **prerank** | ion gated channel activity (GO:0022839) | -0.5477061 | -1.1303385 | 0.30487805 | 0.71283387 |
| **prerank** | epidermal growth factor receptor binding (GO:0005154) | -0.5466364 | -1.128745 | 0.30050083 | 0.71164598 |
| **prerank** | K63-linked polyubiquitin modification-dependent protein binding (GO:0070530) | -0.567989 | -1.1279423 | 0.31958763 | 0.70808243 |

| | | | | | |
|---|---|---|---|---|---|
| **prerank** | exoribonuclease activity, producing 5'-phosphomonoesters (GO:0016896) | -0.5246965 | -1.1277574 | 0.29752066 | 0.70243426 |
| **prerank** | voltage-gated sodium channel activity (GO:0005248) | 0.52855261 | 1.12725086 | 0.28125 | 0.89018645 |
| **prerank** | phosphatase activity (GO:0016791) | -0.4584799 | -1.1262998 | 0.23308271 | 0.70139198 |
| **prerank** | oxidoreductase activity, acting on single donors with incorporation of molecular oxygen, incorporation of two atoms of oxygen (GO:0016702) | -0.5840186 | -1.1252141 | 0.31238779 | 0.69882471 |
| **prerank** | mRNA 3'-UTR binding (GO:0003730) | 0.41131024 | 1.12517947 | 0.23809524 | 0.88311781 |
| **prerank** | tRNA methyltransferase activity (GO:0008175) | 0.56379933 | 1.12498481 | 0.31395349 | 0.86911864 |
| **prerank** | transcription regulatory region nucleic acid binding (GO:0001067) | -0.4043375 | -1.1245 | 0.17718121 | 0.69510197 |
| **prerank** | oxidoreductase activity, acting on the CH-CH group of donors, NAD or NADP as acceptor (GO:0016628) | -0.5841122 | -1.1244851 | 0.3102253 | 0.68928538 |
| **prerank** | actinin binding (GO:0042805) | -0.5648573 | -1.1234971 | 0.29084967 | 0.6867856 |
| **prerank** | acidic amino acid transmembrane transporter activity (GO:0015172) | -0.5873221 | -1.1233508 | 0.33846154 | 0.68157739 |
| **prerank** | transmembrane receptor protein kinase activity (GO:0019199) | -0.4719723 | -1.1211789 | 0.28293413 | 0.68263679 |
| **prerank** | protein serine/threonine kinase activity (GO:0004674) | 0.3471934 | 1.11874945 | 0.08988764 | 0.87703251 |
| **prerank** | protein tyrosine phosphatase activity (GO:0004725) | 0.41647759 | 1.11817159 | 0.23192771 | 0.86492117 |

| | | | | | |
|---|---|---|---|---|---|
| **prerank** | cadherin binding (GO:0045296) | 0.35157209 | 1.11599912 | 0.13333333 | 0.85942018 |
| **prerank** | triglyceride lipase activity (GO:0004806) | -0.5488346 | -1.1136002 | 0.31533101 | 0.70167824 |
| **prerank** | cadherin binding involved in cell-cell adhesion (GO:0098641) | 0.54037997 | 1.11346244 | 0.3202765 | 0.8555292 |
| **prerank** | pyrophosphatase activity (GO:0016462) | 0.43789692 | 1.11168923 | 0.24719101 | 0.84843047 |
| **prerank** | SUMO transferase activity (GO:0019789) | -0.5659974 | -1.1110843 | 0.33047945 | 0.70416367 |
| **prerank** | transmembrane receptor protein tyrosine kinase activity (GO:0004714) | -0.46632 | -1.1097083 | 0.28507463 | 0.70253472 |
| **prerank** | transition metal ion binding (GO:0046914) | -0.3787928 | -1.1095161 | 0.18779904 | 0.69746239 |
| **prerank** | transition metal ion transmembrane transporter activity (GO:0046915) | 0.4834126 | 1.10944281 | 0.28057554 | 0.84354901 |
| **prerank** | tubulin binding (GO:0015631) | 0.34817756 | 1.1090668 | 0.14418605 | 0.83215384 |
| **prerank** | endopeptidase inhibitor activity (GO:0004866) | -0.4345121 | -1.1075346 | 0.276411 | 0.69817061 |

| | | | | | |
|---|---|---|---|---|---|
| **prerank** | zinc ion binding (GO:0008270) | -0.3839245 | -1.102984 | 0.21689786 | 0.70665067 |
| **prerank** | phospholipase C activity (GO:0004629) | 0.49645035 | 1.10073728 | 0.32663317 | 0.85107092 |
| **prerank** | mitogen-activated protein kinase binding (GO:0051019) | -0.5119006 | -1.1006246 | 0.33887043 | 0.70837343 |
| **prerank** | kinase binding (GO:0019900) | -0.3760295 | -1.0997982 | 0.18544601 | 0.70569395 |
| **prerank** | transmembrane receptor protein phosphatase activity (GO:0019198) | -0.5773638 | -1.0988826 | 0.35765125 | 0.69770703 |
| **prerank** | transmembrane receptor protein tyrosine phosphatase activity (GO:0005001) | -0.5773638 | -1.0988826 | 0.35765125 | 0.69770703 |
| **prerank** | C-C chemokine receptor activity (GO:0016493) | 0.5020178 | 1.09796763 | 0.33095238 | 0.8489645 |
| **prerank** | amino acid transmembrane transporter activity (GO:0015171) | 0.44169062 | 1.09790903 | 0.29442971 | 0.83691675 |
| **prerank** | transcription corepressor binding (GO:0001222) | -0.5200546 | -1.0948725 | 0.33222037 | 0.70425046 |
| **prerank** | GABA-A receptor activity (GO:0004890) | 0.51725168 | 1.09364808 | 0.32079646 | 0.83897792 |
| **prerank** | transforming growth factor beta binding (GO:0050431) | -0.5332682 | -1.0931947 | 0.34102142 | 0.70374968 |
| **prerank** | voltage-gated cation channel activity (GO:0022843) | 0.39168827 | 1.09270399 | 0.22461538 | 0.83022098 |
| **prerank** | receptor antagonist activity (GO:0048019) | -0.5692434 | -1.0923821 | 0.37413793 | 0.70101577 |
| **prerank** | sulfuric ester hydrolase activity (GO:0008484) | -0.5716917 | -1.0921234 | 0.36983842 | 0.69657868 |
| **prerank** | DNA polymerase binding (GO:0070182) | 0.53537601 | 1.09198246 | 0.33407572 | 0.82105767 |
| **prerank** | low-density lipoprotein particle receptor binding (GO:0050750) | -0.5503337 | -1.0919012 | 0.33982301 | 0.6920853 |

| | | | | | |
|---|---|---|---|---|---|
| **prerank** | cysteine-type endopeptidase activator activity involved in apoptotic process (GO:0008656) | 0.53932908 | 1.09016868 | 0.33261803 | 0.81630302 |
| **prerank** | nuclear receptor binding (GO:0016922) | -0.4194469 | -1.0894949 | 0.29472141 | 0.69455779 |
| **prerank** | glutamate receptor binding (GO:0035254) | 0.52962544 | 1.08068426 | 0.37560976 | 0.8376901 |
| **prerank** | calcium ion binding (GO:0005509) | 0.33654252 | 1.07745188 | 0.20353982 | 0.83805136 |
| **prerank** | metal ion transmembrane transporter activity (GO:0046873) | 0.47943431 | 1.0770554 | 0.35696822 | 0.82835098 |
| **prerank** | UDP-glycosyltransferase activity (GO:0008194) | -0.4348535 | -1.075639 | 0.31852986 | 0.73037278 |
| **prerank** | 3'-5'-exoribonuclease activity (GO:0000175) | -0.4952858 | -1.0736814 | 0.35679214 | 0.73114258 |
| **prerank** | hexosyltransferase activity (GO:0016758) | -0.4130159 | -1.0727644 | 0.31470588 | 0.72875977 |
| **prerank** | calcium channel activity (GO:0005262) | 0.38743144 | 1.07251953 | 0.29498525 | 0.83334623 |
| **prerank** | C-C chemokine binding (GO:0019957) | 0.48532012 | 1.07238038 | 0.3627685 | 0.82306208 |
| **prerank** | cyclic nucleotide binding (GO:0030551) | -0.5411624 | -1.0718205 | 0.37913907 | 0.72658231 |
| **prerank** | chemokine receptor binding (GO:0042379) | -0.4719375 | -1.0701288 | 0.35634029 | 0.72674837 |
| **prerank** | cyclic-nucleotide phosphodiesterase activity (GO:0004112) | -0.5289854 | -1.0696645 | 0.3777403 | 0.7230228 |
| **prerank** | ankyrin binding (GO:0030506) | 0.50627008 | 1.0693186 | 0.35576923 | 0.8233797 |
| **prerank** | endodeoxyribonuclease activity (GO:0004520) | 0.50385603 | 1.06835034 | 0.38297872 | 0.81607574 |
| **prerank** | lysine N-methyltransferase activity (GO:0016278) | 0.49757669 | 1.06468272 | 0.34739454 | 0.81811429 |
| **prerank** | DNA binding, bending (GO:0008301) | 0.54541689 | 1.06131553 | 0.41387025 | 0.81949469 |
| **prerank** | exonuclease activity (GO:0004527) | -0.4609638 | -1.057776 | 0.36928105 | 0.75334407 |
| **prerank** | general transcription initiation factor binding (GO:0140296) | -0.5096462 | -1.0559618 | 0.39455782 | 0.75345709 |

| | | | | | |
|---|---|---|---|---|---|
| **prerank** | protein heterodimerization activity (GO:0046982) | -0.3854198 | -1.0549984 | 0.34178905 | 0.75129188 |
| **prerank** | non-membrane spanning protein tyrosine kinase activity (GO:0004715) | 0.45429861 | 1.05498966 | 0.32541568 | 0.83133605 |
| **prerank** | hormone activity (GO:0005179) | -0.4415174 | -1.0515112 | 0.37676609 | 0.7565068 |
| **prerank** | bicarbonate transmembrane transporter activity (GO:0015106) | 0.54181284 | 1.05128484 | 0.39856802 | 0.83403579 |
| **prerank** | endopeptidase regulator activity (GO:0061135) | 0.4366096 | 1.05091799 | 0.33866667 | 0.82522633 |
| **prerank** | zinc ion transmembrane transporter activity (GO:0005385) | -0.5349584 | -1.0498214 | 0.40243902 | 0.75655047 |
| **prerank** | FAD binding (GO:0071949) | -0.5105771 | -1.0482431 | 0.40508475 | 0.7562358 |
| **prerank** | nucleotidase activity (GO:0008252) | 0.53141562 | 1.04813762 | 0.3923445 | 0.82428432 |
| **prerank** | serine-type peptidase activity (GO:0008236) | -0.4035746 | -1.0477674 | 0.3694362 | 0.75266858 |
| **prerank** | MHC protein binding (GO:0042287) | 0.47428013 | 1.04118369 | 0.39671362 | 0.83748554 |
| **prerank** | metalloexopeptidase activity (GO:0008235) | -0.4393897 | -1.0404645 | 0.39749609 | 0.7687911 |
| **prerank** | NADP binding (GO:0050661) | -0.4705346 | -1.0398368 | 0.40930979 | 0.7655392 |
| **prerank** | serine-type endopeptidase activity (GO:0004252) | -0.4098639 | -1.0398182 | 0.37537538 | 0.76060618 |
| **prerank** | phosphatidylinositol-3,4-bisphosphate binding (GO:0043325) | 0.46125439 | 1.03669937 | 0.41518987 | 0.84265025 |
| **prerank** | DNA polymerase activity (GO:0034061) | 0.48823243 | 1.03570395 | 0.4028436 | 0.83648735 |
| **prerank** | lipoprotein particle receptor binding (GO:0070325) | -0.5008706 | -1.0348918 | 0.40172414 | 0.77029079 |
| **prerank** | solute:proton symporter activity (GO:0015295) | 0.51249882 | 1.03284895 | 0.44811321 | 0.8364553 |
| **prerank** | nuclear localization sequence binding (GO:0008139) | 0.48255958 | 1.03276282 | 0.42688679 | 0.82751477 |
| **prerank** | DNA-directed DNA polymerase activity (GO:0003887) | 0.4908885 | 1.03231785 | 0.40566038 | 0.819808 |
| **prerank** | RNA polymerase II complex binding (GO:0000993) | 0.44976613 | 1.03132693 | 0.4075067 | 0.81418931 |
| **prerank** | receptor tyrosine kinase binding (GO:0030971) | -0.4327889 | -1.0309215 | 0.39613527 | 0.77746198 |
| **prerank** | glutathione transferase activity (GO:0004364) | 0.46006616 | 1.03023555 | 0.37990196 | 0.80897766 |
| **prerank** | pre-mRNA binding (GO:0036002) | 0.45767244 | 1.02930748 | 0.4085213 | 0.80353655 |

| | | | | | |
|---|---|---|---|---|---|
| **prerank** | proton transmembrane transporter activity (GO:0015078) | 0.42069209 | 1.02754207 | 0.39845758 | 0.80062326 |
| **prerank** | endoribonuclease activity (GO:0004521) | -0.4744629 | -1.0243081 | 0.44781145 | 0.79229165 |
| **prerank** | flavin adenine dinucleotide binding (GO:0050660) | -0.4261062 | -1.0241467 | 0.39481707 | 0.78772795 |
| **prerank** | serine-type endopeptidase inhibitor activity (GO:0004867) | 0.40687382 | 1.02348689 | 0.41530055 | 0.80564593 |
| **prerank** | CXCR chemokine receptor binding (GO:0045236) | -0.5378188 | -1.0220342 | 0.43915344 | 0.78902689 |
| **prerank** | phosphatidylinositol-5-phosphate binding (GO:0010314) | -0.5330324 | -1.020477 | 0.45884413 | 0.78846793 |
| **prerank** | ligand-gated calcium channel activity (GO:0099604) | 0.48704918 | 1.01624777 | 0.42488263 | 0.8189874 |
| **prerank** | ATP hydrolysis activity (GO:0140603) | -0.4415903 | -1.0117252 | 0.44067797 | 0.80899996 |
| **prerank** | single-stranded RNA binding (GO:0003727) | -0.4488252 | -1.0098208 | 0.41853035 | 0.80938464 |
| **prerank** | CARD domain binding (GO:0050700) | 0.51206141 | 1.00929265 | 0.44083527 | 0.83325624 |
| **prerank** | protein kinase regulator activity (GO:0019887) | -0.3951966 | -1.0090372 | 0.45820896 | 0.80684942 |
| **prerank** | peptidase activator activity involved in apoptotic process (GO:0016505) | 0.49780557 | 1.00795299 | 0.4595843 | 0.82930997 |
| **prerank** | retinoic acid receptor binding (GO:0042974) | -0.5186298 | -1.0044526 | 0.44178082 | 0.81541121 |
| **prerank** | ligand-gated anion channel activity (GO:0099095) | 0.49302512 | 1.00316407 | 0.44897959 | 0.83611375 |
| **prerank** | androgen receptor binding (GO:0050681) | 0.45322506 | 1.00211944 | 0.46028037 | 0.83105515 |
| **prerank** | pyrophosphate hydrolysis-driven proton transmembrane transporter activity (GO:0009678) | -0.4878072 | -0.9988129 | 0.44949495 | 0.82687918 |
| **prerank** | phosphatidylinositol-3-phosphate binding (GO:0032266) | -0.4440163 | -0.9988074 | 0.47882736 | 0.82192149 |
| **prerank** | carboxypeptidase activity (GO:0004180) | -0.4475345 | -0.997593 | 0.47987118 | 0.82032427 |
| **prerank** | oxidoreductase activity, acting on paired donors, with incorporation or reduction of molecular oxygen, reduced flavin or flavoprotein as one donor, and incorporation of one atom of oxygen (GO:0016712) | 0.48510541 | 0.99543154 | 0.44772727 | 0.84319954 |
| **prerank** | phosphatidylinositol-4,5-bisphosphate binding (GO:0005546) | -0.403368 | -0.9952546 | 0.45970149 | 0.82194038 |

| | | | | | |
|---|---|---|---|---|---|
| **prerank** | DNA-binding transcription repressor activity, RNA polymerase II-specific (GO:0001227) | 0.32274134 | 0.99422595 | 0.43722944 | 0.83903959 |
| **prerank** | copper ion binding (GO:0005507) | 0.40402278 | 0.99390168 | 0.4488189 | 0.83215744 |
| **prerank** | Wnt-activated receptor activity (GO:0042813) | -0.5276527 | -0.9922893 | 0.5 | 0.82532716 |
| **prerank** | metalloaminopeptidase activity (GO:0070006) | -0.5079723 | -0.9918404 | 0.48364888 | 0.82181534 |
| **prerank** | inward rectifier potassium channel activity (GO:0005242) | -0.4879642 | -0.9913765 | 0.50259067 | 0.8182476 |
| **prerank** | deubiquitinase activity (GO:0101005) | -0.3919326 | -0.9893436 | 0.48762737 | 0.81896459 |
| **prerank** | histone methyltransferase activity (GO:0042054) | 0.43715939 | 0.98862218 | 0.47201946 | 0.84043729 |
| **prerank** | excitatory extracellular ligand-gated ion channel activity (GO:0005231) | -0.4978584 | -0.9872118 | 0.47950089 | 0.8201245 |
| **prerank** | steroid hydroxylase activity (GO:0008395) | -0.4635528 | -0.9868009 | 0.46952224 | 0.81665603 |
| **prerank** | NAD binding (GO:0051287) | -0.4597316 | -0.9832454 | 0.46959459 | 0.82219268 |
| **prerank** | active ion transmembrane transporter activity (GO:0022853) | 0.40471395 | 0.98264476 | 0.46883469 | 0.85103945 |
| **prerank** | solute:sodium symporter activity (GO:0015370) | -0.4561649 | -0.9825238 | 0.48548387 | 0.81951138 |
| **prerank** | adenyl ribonucleotide binding (GO:0032559) | -0.3434774 | -0.9795512 | 0.5258512 | 0.82276431 |
| **prerank** | thiol-dependent deubiquitinase (GO:0004843) | -0.3903812 | -0.9792476 | 0.48613139 | 0.81884368 |
| **prerank** | histone deacetylase activity (GO:0004407) | 0.46537996 | 0.97445987 | 0.49144254 | 0.86855751 |
| **prerank** | phosphatidylinositol binding (GO:0035091) | -0.3822377 | -0.9731907 | 0.5058309 | 0.83037126 |
| **prerank** | metal ion binding (GO:0046872) | -0.328174 | -0.9721466 | 0.54329775 | 0.82873945 |
| **prerank** | protein kinase A regulatory subunit binding (GO:0034237) | -0.4789842 | -0.9721197 | 0.49839744 | 0.82425242 |

| | | | | | |
|---|---|---|---|---|---|
| **prerank** | semaphorin receptor binding (GO:0030215) | 0.4469426 | 0.9690224 | 0.48019802 | 0.8782399 |
| **prerank** | RNA polymerase core enzyme binding (GO:0043175) | 0.41976461 | 0.966991 | 0.49869452 | 0.87647343 |
| **prerank** | RNA cap binding (GO:0000339) | -0.4883718 | -0.9661221 | 0.51669596 | 0.83603741 |
| **prerank** | nucleotide diphosphatase activity (GO:0004551) | -0.5099639 | -0.9658943 | 0.5353902 | 0.83207139 |
| **prerank** | transmembrane receptor protein serine/threonine kinase activity (GO:0004675) | 0.46869345 | 0.96574404 | 0.4893617 | 0.8722667 |
| **prerank** | low-density lipoprotein particle binding (GO:0030169) | -0.5167906 | -0.9656729 | 0.52659574 | 0.82809015 |
| **prerank** | mRNA 3'-UTR AU-rich region binding (GO:0035925) | 0.45061369 | 0.96556955 | 0.5025641 | 0.86489622 |
| **prerank** | neuropeptide binding (GO:0042923) | 0.48838583 | 0.96553073 | 0.50117096 | 0.85732505 |
| **prerank** | potassium ion antiporter activity (GO:0022821) | -0.5054338 | -0.9615177 | 0.53504274 | 0.83493235 |
| **prerank** | ribosomal small subunit binding (GO:0043024) | 0.47473631 | 0.95984662 | 0.50608273 | 0.86745631 |
| **prerank** | Notch binding (GO:0005112) | 0.45162411 | 0.95938044 | 0.50728155 | 0.86129607 |
| **prerank** | poly(A) binding (GO:0008143) | 0.45165312 | 0.95899419 | 0.53284672 | 0.85499115 |
| **prerank** | phosphotransferase activity, alcohol group as acceptor (GO:0016773) | -0.3912941 | -0.957991 | 0.51131222 | 0.8398597 |
| **prerank** | estrogen receptor binding (GO:0030331) | 0.40520596 | 0.95717512 | 0.4961039 | 0.852694 |
| **prerank** | single-stranded DNA binding (GO:0003697) | 0.3442942 | 0.95679633 | 0.54455446 | 0.84664812 |
| **prerank** | tau protein binding (GO:0048156) | -0.4267735 | -0.9562316 | 0.54103853 | 0.84007489 |
| **prerank** | ATPase regulator activity (GO:0060590) | -0.4299657 | -0.9561169 | 0.52842809 | 0.83596452 |
| **prerank** | postsynaptic neurotransmitter receptor activity (GO:0098960) | -0.4938663 | -0.9540968 | 0.53623188 | 0.83675043 |
| **prerank** | kinase activity (GO:0016301) | -0.3682374 | -0.9537282 | 0.53084648 | 0.83328626 |
| **prerank** | poly(U) RNA binding (GO:0008266) | -0.4772675 | -0.9527094 | 0.53571429 | 0.8315697 |
| **prerank** | GABA receptor activity (GO:0016917) | 0.44290706 | 0.95217341 | 0.503386 | 0.85278366 |
| **prerank** | voltage-gated anion channel activity (GO:0008308) | 0.47924923 | 0.95139036 | 0.52415459 | 0.84789452 |
| **prerank** | aldehyde dehydrogenase [NAD(P)+] activity (GO:0004030) | -0.5132941 | -0.9500767 | 0.56852792 | 0.8339162 |
| **prerank** | calmodulin-dependent protein kinase activity (GO:0004683) | 0.43132111 | 0.94990368 | 0.52235294 | 0.84565465 |

| | | | | | |
|---|---|---|---|---|---|
| **prerank** | calcium-dependent phospholipid binding (GO:0005544) | -0.4087582 | -0.9498671 | 0.52542373 | 0.830106 |
| **prerank** | protein tyrosine kinase activity (GO:0004713) | -0.3678879 | -0.9484809 | 0.5497076 | 0.82948354 |
| **prerank** | S-adenosylmethionine-dependent methyltransferase activity (GO:0008757) | 0.39731032 | 0.94820457 | 0.55955679 | 0.84364745 |
| **prerank** | protein kinase activator activity (GO:0030295) | -0.3973691 | -0.9477646 | 0.56528662 | 0.82712117 |
| **prerank** | organic anion transmembrane transporter activity (GO:0008514) | -0.3560578 | -0.9466822 | 0.56363636 | 0.82566405 |
| **prerank** | phosphatidylinositol 3-kinase regulator activity (GO:0035014) | 0.45023724 | 0.94544593 | 0.54066986 | 0.84466434 |
| **prerank** | phosphatase inhibitor activity (GO:0019212) | 0.44875907 | 0.94540172 | 0.53465347 | 0.83791959 |
| **prerank** | 3',5'-cyclic-nucleotide phosphodiesterase activity (GO:0004114) | -0.4738439 | -0.9453768 | 0.54697987 | 0.82489036 |
| **prerank** | phosphatidylinositol bisphosphate binding (GO:1902936) | -0.371907 | -0.9449925 | 0.57692308 | 0.82160141 |
| **prerank** | transmitter-gated ion channel activity (GO:0022824) | 0.40326053 | 0.94131237 | 0.53902439 | 0.8430405 |
| **prerank** | magnesium ion binding (GO:0000287) | 0.31961408 | 0.93993763 | 0.59726962 | 0.84048258 |
| **prerank** | polyubiquitin modification-dependent protein binding (GO:0031593) | -0.4077892 | -0.9363243 | 0.5850234 | 0.83906611 |
| **prerank** | ion channel inhibitor activity (GO:0008200) | 0.40018758 | 0.93626963 | 0.56598985 | 0.84436505 |
| **prerank** | tRNA binding (GO:0000049) | 0.37468298 | 0.9362158 | 0.56198347 | 0.8379471 |
| **prerank** | catalytic activity, acting on a tRNA (GO:0140101) | 0.39460772 | 0.93292032 | 0.53026634 | 0.84123307 |
| **prerank** | WW domain binding (GO:0050699) | -0.4350374 | -0.9325289 | 0.56185567 | 0.84407182 |
| **prerank** | kinase inhibitor activity (GO:0019210) | -0.4302423 | -0.9324133 | 0.57705479 | 0.84015401 |
| **prerank** | amyloid-beta binding (GO:0001540) | -0.3813055 | -0.9312449 | 0.59600614 | 0.83897513 |
| **prerank** | muscle alpha-actinin binding (GO:0051371) | -0.4929058 | -0.9310662 | 0.55985915 | 0.83523158 |
| **prerank** | peptidase inhibitor activity (GO:0030414) | -0.4211191 | -0.9306422 | 0.56012658 | 0.83211781 |
| **prerank** | channel activity (GO:0015267) | -0.4023198 | -0.9293616 | 0.58101473 | 0.83105725 |

| | | | | | |
|---|---|---|---|---|---|
| **prerank** | voltage-gated potassium channel activity (GO:0005249) | -0.3742188 | -0.9282868 | 0.56888889 | 0.82970386 |
| **prerank** | voltage-gated calcium channel activity (GO:0005245) | 0.39793084 | 0.92818607 | 0.56167979 | 0.84822633 |
| **prerank** | PDZ domain binding (GO:0030165) | -0.3906994 | -0.9273611 | 0.6 | 0.82777434 |
| **prerank** | ATPase activator activity (GO:0001671) | -0.4599184 | -0.9263058 | 0.56347826 | 0.82627617 |
| **prerank** | endoribonuclease activity, producing 5'-phosphomonoesters (GO:0016891) | 0.41805234 | 0.92598858 | 0.57655502 | 0.84803536 |
| **prerank** | cysteine-type endopeptidase inhibitor activity involved in apoptotic process (GO:0043027) | -0.4632672 | -0.9259472 | 0.58422939 | 0.82326107 |
| **prerank** | protein tyrosine kinase binding (GO:1990782) | -0.3712836 | -0.925325 | 0.62442748 | 0.82079043 |
| **prerank** | snoRNA binding (GO:0030515) | -0.4374455 | -0.9249315 | 0.59546926 | 0.81785765 |
| **prerank** | thyroid hormone receptor binding (GO:0046966) | -0.4375355 | -0.9163551 | 0.59228188 | 0.83414018 |
| **prerank** | kinase activator activity (GO:0019209) | -0.4340231 | -0.9156964 | 0.58053097 | 0.83190062 |
| **prerank** | protein kinase inhibitor activity (GO:0004860) | -0.4049987 | -0.9148486 | 0.6091954 | 0.83005587 |
| **prerank** | methyltransferase activity (GO:0008168) | 0.37040746 | 0.91327161 | 0.62637363 | 0.87760418 |
| **prerank** | manganese ion binding (GO:0030145) | -0.3994468 | -0.9106119 | 0.57835218 | 0.83610665 |
| **prerank** | disordered domain specific binding (GO:0097718) | -0.4523141 | -0.9077198 | 0.61435726 | 0.83889418 |
| **prerank** | mRNA 5'-UTR binding (GO:0048027) | 0.40864318 | 0.90640436 | 0.59134615 | 0.88961334 |
| **prerank** | DNA secondary structure binding (GO:0000217) | 0.38761011 | 0.90493824 | 0.62210797 | 0.88710879 |
| **prerank** | cation transmembrane transporter activity (GO:0008324) | -0.3975894 | -0.9048454 | 0.62193126 | 0.84169199 |
| **prerank** | ligand-gated cation channel activity (GO:0099094) | -0.3596199 | -0.9047055 | 0.66925466 | 0.83807271 |
| **prerank** | transforming growth factor beta receptor binding (GO:0005160) | 0.44482586 | 0.90470267 | 0.59047619 | 0.88118435 |
| **prerank** | transmitter-gated ion channel activity involved in regulation of postsynaptic membrane potential (GO:1904315) | 0.3945515 | 0.90413994 | 0.57320099 | 0.87636352 |
| **prerank** | G protein-coupled receptor binding (GO:0001664) | 0.30900542 | 0.90246934 | 0.75555556 | 0.87429124 |

| | | | | | |
|---|---|---|---|---|---|
| **prerank** | methyl-CpG binding (GO:0008327) | 0.41375793 | 0.90126609 | 0.58542714 | 0.87114179 |
| **prerank** | GTPase binding (GO:0051020) | -0.3273497 | -0.9003598 | 0.72207447 | 0.84422141 |
| **prerank** | omega peptidase activity (GO:0008242) | -0.3555396 | -0.8990133 | 0.65647744 | 0.84351078 |
| **prerank** | acetylglucosaminyltransferase activity (GO:0008375) | -0.3902215 | -0.8983707 | 0.61788618 | 0.84118186 |
| **prerank** | ligand-gated ion channel activity (GO:0015276) | 0.38749073 | 0.89763554 | 0.63549161 | 0.87407968 |
| **prerank** | four-way junction DNA binding (GO:0000400) | 0.44185377 | 0.89751585 | 0.57718121 | 0.86814807 |
| **prerank** | fibroblast growth factor binding (GO:0017134) | -0.4512953 | -0.8954888 | 0.6320132 | 0.84387998 |
| **prerank** | cyclin-dependent protein serine/threonine kinase regulator activity (GO:0016538) | 0.36212066 | 0.89539103 | 0.60526316 | 0.8676598 |
| **prerank** | carboxylic acid transmembrane transporter activity (GO:0046943) | -0.37929 | -0.8906505 | 0.66932907 | 0.85102996 |
| **prerank** | protein phosphorylated amino acid binding (GO:0045309) | 0.37084982 | 0.89016164 | 0.62280702 | 0.87538386 |
| **prerank** | ATP binding (GO:0005524) | -0.313281 | -0.8900505 | 0.77078086 | 0.84855491 |
| **prerank** | carboxy-lyase activity (GO:0016831) | -0.4365725 | -0.888949 | 0.62648557 | 0.84725972 |
| **prerank** | dicarboxylic acid transmembrane transporter activity (GO:0005310) | 0.39376928 | 0.88748162 | 0.6015625 | 0.87620377 |
| **prerank** | protein kinase C binding (GO:0005080) | -0.3890451 | -0.8872562 | 0.65814696 | 0.84727983 |
| **prerank** | inorganic anion transmembrane transporter activity (GO:0015103) | 0.4018396 | 0.88648839 | 0.63037975 | 0.87284024 |
| **prerank** | poly-pyrimidine tract binding (GO:0008187) | -0.4356653 | -0.8860221 | 0.63202726 | 0.84634983 |
| **prerank** | clathrin adaptor activity (GO:0035615) | 0.44727147 | 0.87615476 | 0.65478842 | 0.89260506 |
| **prerank** | ADP binding (GO:0043531) | -0.420215 | -0.8751598 | 0.69482759 | 0.866697 |
| **prerank** | cell-cell adhesion mediator activity (GO:0098632) | -0.3844433 | -0.8738878 | 0.66129032 | 0.86588633 |

| | | | | | |
|---|---|---|---|---|---|
| **prerank** | neurotransmitter receptor activity involved in regulation of postsynaptic membrane potential (GO:0099529) | 0.41274565 | 0.8736827 | 0.64764268 | 0.89229349 |
| **prerank** | ubiquitin-specific protease binding (GO:1990381) | -0.440233 | -0.8705013 | 0.65604027 | 0.86921891 |
| **prerank** | hydrolase activity, hydrolyzing N-glycosyl compounds (GO:0016799) | -0.4607188 | -0.8690522 | 0.6427289 | 0.86837055 |
| **prerank** | tropomyosin binding (GO:0005523) | -0.4548281 | -0.8684798 | 0.63888889 | 0.86581779 |
| **prerank** | ephrin receptor activity (GO:0005003) | 0.43759508 | 0.86748312 | 0.64634146 | 0.89513126 |
| **prerank** | transmembrane-ephrin receptor activity (GO:0005005) | 0.43759508 | 0.86748312 | 0.64634146 | 0.89513126 |
| **prerank** | anion channel activity (GO:0005253) | -0.3557303 | -0.86614 | 0.71090047 | 0.86710606 |
| **prerank** | L-amino acid transmembrane transporter activity (GO:0015179) | 0.34489198 | 0.86427831 | 0.71134021 | 0.89662273 |
| **prerank** | secondary active transmembrane transporter activity (GO:0015291) | 0.33781312 | 0.86389806 | 0.72423398 | 0.89149756 |
| **prerank** | ubiquitin-like protein conjugating enzyme binding (GO:0044390) | 0.36115079 | 0.86139543 | 0.69543147 | 0.89111415 |
| **prerank** | poly-purine tract binding (GO:0070717) | 0.37660509 | 0.85944035 | 0.68105516 | 0.89015454 |
| **prerank** | frizzled binding (GO:0005109) | 0.36692563 | 0.8590314 | 0.6953317 | 0.88538953 |
| **prerank** | phosphotyrosine residue binding (GO:0001784) | 0.36549231 | 0.85878034 | 0.70028011 | 0.88025444 |
| **prerank** | neutral amino acid transmembrane transporter activity (GO:0015175) | -0.3985378 | -0.8584134 | 0.68874172 | 0.88002527 |
| **prerank** | syntaxin binding (GO:0019905) | -0.3609379 | -0.8583419 | 0.72527473 | 0.87641334 |
| **prerank** | signal sequence binding (GO:0005048) | 0.35767539 | 0.85553015 | 0.7154047 | 0.88193168 |
| **prerank** | nucleotidyltransferase activity (GO:0016779) | -0.4285617 | -0.8546647 | 0.6592845 | 0.88062953 |
| **prerank** | E-box binding (GO:0070888) | 0.35771173 | 0.85433722 | 0.7372449 | 0.87881016 |
| **prerank** | sialyltransferase activity (GO:0008373) | -0.425634 | -0.8526927 | 0.67238422 | 0.88104722 |
| **prerank** | small GTPase binding (GO:0031267) | -0.3116885 | -0.8508531 | 0.82567568 | 0.88122773 |
| **prerank** | potassium channel activity (GO:0005267) | -0.3394569 | -0.8499719 | 0.76117103 | 0.87922258 |

| | | | | | |
|---|---|---|---|---|---|
| **prerank** | histone acetyltransferase binding (GO:0035035) | -0.4258851 | -0.8485961 | 0.68838028 | 0.87840907 |
| **prerank** | R-SMAD binding (GO:0070412) | -0.4324336 | -0.8483242 | 0.67977528 | 0.87535532 |
| **prerank** | protein-lysine N-methyltransferase activity (GO:0016279) | 0.36030276 | 0.84762351 | 0.70050761 | 0.88785699 |
| **prerank** | peptidase activator activity (GO:0016504) | -0.4177471 | -0.8463987 | 0.71 | 0.87562491 |
| **prerank** | anion transmembrane transporter activity (GO:0008509) | 0.41801593 | 0.84512229 | 0.67505721 | 0.88780307 |
| **prerank** | antiporter activity (GO:0015297) | -0.41969 | -0.8433517 | 0.69204152 | 0.87801637 |
| **prerank** | beta-tubulin binding (GO:0048487) | 0.36250541 | 0.840893 | 0.70263789 | 0.89124719 |
| **prerank** | phosphatidylinositol phosphate binding (GO:1901981) | 0.30002315 | 0.84059675 | 0.87859425 | 0.88624207 |
| **prerank** | protein phosphatase regulator activity (GO:0019888) | -0.3527678 | -0.837586 | 0.74454829 | 0.88590666 |
| **prerank** | syntaxin-1 binding (GO:0017075) | -0.4194704 | -0.837312 | 0.71177504 | 0.88280845 |
| **prerank** | voltage-gated ion channel activity (GO:0005244) | 0.41991847 | 0.83531846 | 0.69515012 | 0.89166089 |
| **prerank** | rRNA binding (GO:0019843) | 0.34532708 | 0.83393843 | 0.76030928 | 0.88906971 |
| **prerank** | oxidoreductase activity, acting on the aldehyde or oxo group of donors, NAD or NADP as acceptor (GO:0016620) | -0.4148724 | -0.8336527 | 0.70894309 | 0.886502 |
| **prerank** | calcium activated cation channel activity (GO:0005227) | -0.4283337 | -0.8330737 | 0.71119134 | 0.88407431 |
| **prerank** | oxidoreduction-driven active transmembrane transporter activity (GO:0015453) | -0.3545778 | -0.8330558 | 0.75555556 | 0.88055671 |
| **prerank** | phosphoric diester hydrolase activity (GO:0008081) | -0.3760757 | -0.8287962 | 0.73217116 | 0.88549618 |
| **prerank** | chemorepellent activity (GO:0045499) | -0.3955546 | -0.8269265 | 0.74621849 | 0.88571267 |
| **prerank** | 1-phosphatidylinositol-3-kinase regulator activity (GO:0046935) | 0.41213699 | 0.82647954 | 0.70921986 | 0.89853682 |
| **prerank** | carboxylic acid binding (GO:0031406) | -0.3714636 | -0.82575 | 0.76333333 | 0.88448135 |
| **prerank** | RNA methyltransferase activity (GO:0008173) | 0.40348936 | 0.82410863 | 0.70495495 | 0.8975578 |
| **prerank** | 2-oxoglutarate-dependent dioxygenase activity (GO:0016706) | 0.35648768 | 0.8222081 | 0.75888325 | 0.89575282 |
| **prerank** | cytochrome-c oxidase activity (GO:0004129) | -0.4312602 | -0.8194 | 0.73529412 | 0.89316414 |

| | | | | | |
|---|---|---|---|---|---|
| **prerank** | acetylgalactosaminyltransferase activity (GO:0008376) | 0.37395093 | 0.81918789 | 0.75184275 | 0.89621746 |
| **prerank** | phosphatidylinositol-4-phosphate binding (GO:0070273) | 0.37571916 | 0.81851077 | 0.7470726 | 0.89215792 |
| **prerank** | TBP-class protein binding (GO:0017025) | -0.4076555 | -0.8173452 | 0.72680412 | 0.8935753 |
| **prerank** | metallocarboxypeptidase activity (GO:0004181) | -0.3855523 | -0.8166709 | 0.74958814 | 0.89127886 |
| **prerank** | snRNA binding (GO:0017069) | -0.3646746 | -0.8116062 | 0.75919732 | 0.89707404 |
| **prerank** | protein serine/threonine phosphatase activity (GO:0004722) | -0.33931 | -0.8098432 | 0.80945122 | 0.89683622 |
| **prerank** | protein tyrosine kinase activator activity (GO:0030296) | 0.38435124 | 0.80941065 | 0.75268817 | 0.90463489 |
| **prerank** | phosphatidylinositol-3,4,5-trisphosphate binding (GO:0005547) | 0.34418919 | 0.80895274 | 0.82793017 | 0.90014922 |
| **prerank** | phosphatidic acid binding (GO:0070300) | -0.4065115 | -0.8065453 | 0.74450085 | 0.89941732 |
| **prerank** | core promoter sequence-specific DNA binding (GO:0001046) | -0.3616076 | -0.8053902 | 0.78537361 | 0.89801123 |
| **prerank** | lysine-acetylated histone binding (GO:0070577) | -0.4031532 | -0.7977358 | 0.75935829 | 0.90421939 |
| **prerank** | acetylation-dependent protein binding (GO:0140033) | -0.4031532 | -0.7977358 | 0.75935829 | 0.90421939 |
| **prerank** | aminoacyl-tRNA ligase activity (GO:0004812) | 0.3355633 | 0.79765411 | 0.83333333 | 0.91569735 |
| **prerank** | phosphoric ester hydrolase activity (GO:0042578) | -0.3440368 | -0.7962806 | 0.80640244 | 0.90318491 |
| **prerank** | potassium channel regulator activity (GO:0015459) | -0.3417785 | -0.7886065 | 0.82472613 | 0.91252256 |
| **prerank** | potassium ion transmembrane transporter activity (GO:0015079) | 0.35091188 | 0.78725842 | 0.82524272 | 0.92812777 |
| **prerank** | N-acetyltransferase activity (GO:0008080) | -0.4027103 | -0.7865391 | 0.74526678 | 0.91219523 |
| **prerank** | phosphatidylinositol-3,5-bisphosphate binding (GO:0080025) | -0.3785654 | -0.7856827 | 0.80806452 | 0.91009265 |
| **prerank** | sodium ion transmembrane transporter activity (GO:0015081) | -0.3732357 | -0.7846608 | 0.80369128 | 0.908341 |
| **prerank** | damaged DNA binding (GO:0003684) | -0.3382539 | -0.7839543 | 0.82922824 | 0.90602662 |
| **prerank** | monocarboxylic acid transmembrane transporter activity (GO:0008028) | 0.35251028 | 0.77887866 | 0.83123426 | 0.93614121 |
| **prerank** | kinesin binding (GO:0019894) | 0.34499485 | 0.77772682 | 0.83682984 | 0.93254444 |
| **prerank** | acetyltransferase activity (GO:0016407) | -0.3658708 | -0.7764211 | 0.82166667 | 0.91419574 |
| **prerank** | MHC class II protein complex binding (GO:0023026) | -0.4063158 | -0.7716607 | 0.796875 | 0.91784398 |

| | | | | | |
|---|---|---|---|---|---|
| **prerank** | cyclin-dependent protein serine/threonine kinase activity (GO:0004693) | 0.35282141 | 0.77027196 | 0.81173594 | 0.93354 |
| **prerank** | cyclin-dependent protein kinase activity (GO:0097472) | 0.35282141 | 0.77027196 | 0.81173594 | 0.93354 |
| **prerank** | ligand-gated channel activity (GO:0022834) | 0.3340659 | 0.76747299 | 0.8501171 | 0.93219853 |
| **prerank** | mannosyltransferase activity (GO:0000030) | 0.35589385 | 0.76357753 | 0.83293556 | 0.93253996 |
| **prerank** | protein serine/threonine/tyrosine kinase activity (GO:0004712) | -0.3563195 | -0.7596704 | 0.82623509 | 0.93165031 |
| **prerank** | sodium channel regulator activity (GO:0017080) | -0.3504587 | -0.7588817 | 0.85113269 | 0.92929473 |
| **prerank** | gamma-tubulin binding (GO:0043015) | -0.3659933 | -0.758485 | 0.82271945 | 0.92648847 |
| **prerank** | amino acid binding (GO:0016597) | -0.3565744 | -0.7584668 | 0.83044983 | 0.92311455 |
| **prerank** | ubiquitin-like protein conjugating enzyme activity (GO:0061650) | -0.3421296 | -0.7580444 | 0.84677419 | 0.92036579 |
| **prerank** | NADH dehydrogenase (ubiquinone) activity (GO:0008137) | 0.31807212 | 0.75687492 | 0.8617284 | 0.93149258 |
| **prerank** | NADH dehydrogenase (quinone) activity (GO:0050136) | 0.31807212 | 0.75687492 | 0.8617284 | 0.93149258 |
| **prerank** | endonuclease activity (GO:0004519) | -0.320699 | -0.7532715 | 0.88114105 | 0.92332405 |
| **prerank** | microtubule plus-end binding (GO:0051010) | 0.35124977 | 0.75132296 | 0.82673267 | 0.93390528 |
| **prerank** | ephrin receptor binding (GO:0046875) | -0.3789514 | -0.7486343 | 0.84417808 | 0.92612952 |
| **prerank** | MAP kinase kinase activity (GO:0004708) | -0.4041284 | -0.7473444 | 0.84341637 | 0.92441078 |
| **prerank** | Hsp70 protein binding (GO:0030544) | 0.32871704 | 0.74671898 | 0.89583333 | 0.93476379 |
| **prerank** | C4-dicarboxylate transmembrane transporter activity (GO:0015556) | -0.3959413 | -0.7449673 | 0.82562278 | 0.92407117 |
| **prerank** | basal RNA polymerase II transcription machinery binding (GO:0001099) | 0.32213129 | 0.73914502 | 0.87257618 | 0.93919939 |
| **prerank** | ribosome binding (GO:0043022) | -0.3165939 | -0.739072 | 0.89448819 | 0.92813528 |
| **prerank** | ubiquitin conjugating enzyme activity (GO:0061631) | -0.3396896 | -0.7385967 | 0.85809683 | 0.92535419 |
| **prerank** | actin monomer binding (GO:0003785) | -0.3585313 | -0.7373996 | 0.86394558 | 0.92348157 |
| **prerank** | telomerase RNA binding (GO:0070034) | -0.3660936 | -0.7368547 | 0.83838384 | 0.92090351 |
| **prerank** | pyridoxal phosphate binding (GO:0030170) | 0.3532814 | 0.73534225 | 0.8296837 | 0.9384974 |

| | | | | | |
|---|---|---|---|---|---|
| **prerank** | ubiquitin conjugating enzyme binding (GO:0031624) | 0.3204373 | 0.73310187 | 0.8943299 | 0.93576943 |
| **prerank** | solute:anion antiporter activity (GO:0140323) | 0.36922106 | 0.72733992 | 0.85514019 | 0.93698697 |
| **prerank** | glutamate receptor activity (GO:0008066) | -0.3600778 | -0.7241605 | 0.87666667 | 0.93193463 |
| **prerank** | G protein-coupled amine receptor activity (GO:0008227) | -0.3492516 | -0.7138171 | 0.8812709 | 0.94006062 |
| **prerank** | N-methyltransferase activity (GO:0008170) | -0.3542586 | -0.7137041 | 0.90016103 | 0.93690515 |
| **prerank** | sodium channel activity (GO:0005272) | 0.29812558 | 0.70286132 | 0.94960212 | 0.95622454 |
| **prerank** | NF-kappaB binding (GO:0051059) | 0.31593517 | 0.69819972 | 0.92270531 | 0.95522419 |
| **prerank** | serotonin receptor activity (GO:0099589) | -0.3498262 | -0.6934917 | 0.89761092 | 0.94913429 |
| **prerank** | G protein-coupled serotonin receptor activity (GO:0004993) | -0.3498262 | -0.6934917 | 0.89761092 | 0.94913429 |
| **prerank** | protein serine/threonine kinase activator activity (GO:0043539) | -0.3160564 | -0.693172 | 0.94551282 | 0.94607878 |
| **prerank** | nuclear receptor coactivator activity (GO:0030374) | -0.2931897 | -0.6915967 | 0.97583082 | 0.94419299 |
| **prerank** | protein phosphatase 1 binding (GO:0008157) | 0.32364446 | 0.68888594 | 0.90888383 | 0.95775077 |
| **prerank** | fibroblast growth factor receptor binding (GO:0005104) | 0.33094873 | 0.68203735 | 0.90547264 | 0.95781984 |
| **prerank** | Lys48-specific deubiquitinase activity (GO:1990380) | 0.33940602 | 0.67163911 | 0.8974359 | 0.95993963 |
| **prerank** | anion:anion antiporter activity (GO:0015301) | -0.3422139 | -0.6668143 | 0.90459966 | 0.95949835 |
| **prerank** | tau-protein kinase activity (GO:0050321) | 0.30522286 | 0.64361451 | 0.94651163 | 0.97025996 |
| **prerank** | ionotropic glutamate receptor activity (GO:0004970) | -0.3118621 | -0.603036 | 0.95621716 | 0.98636643 |
| **prerank** | hormone receptor binding (GO:0051427) | -0.2934367 | -0.5458511 | 0.9699115 | 0.99443403 |

### )_Molecular Function

| FWER p-val | Tag % | Gene % |
|---|---|---|
| 0.381 | 22-Jun | 3.12% |
| 0.692 | 25-Aug | 8.62% |
| 0.764 | 20-Jul | 6.50% |
| 0.831 | 13/19 | 13.45% |
| 0.776 | Nov-34 | 10.21% |
| 0.85 | Sep-33 | 7.82% |
| 0.89 | 26-Jun | 9.16% |
| 0.921 | 32/106 | 7.84% |
| 0.924 | 25-Dec | 11.69% |
| 0.938 | 18/53 | 11.48% |
| 0.93 | 15-Dec | 20.30% |
| 0.97 | 65/280 | 10.84% |
| 0.988 | 19-Feb | 2.95% |
| 0.976 | 21/41 | 17.18% |
| 0.976 | 15-Jan | 1.16% |
| 0.994 | 14/25 | 18.03% |
| 0.994 | 19/83 | 7.11% |

| | | |
|---|---|---|
| 0.987 | 35/240 | 6.31% |
| 0.995 | 19/83 | 7.63% |
| 0.995 | 16-May | 10.41% |
| 0.996 | 18-Jun | 13.00% |
| 0.996 | 21/81 | 10.52% |
| 0.989 | 13/32 | 11.57% |
| 0.989 | 16/23 | 17.73% |
| 0.999 | 17-Jun | 13.00% |
| 0.995 | 16-Sep | 16.84% |
| 1 | 17-May | 10.41% |
| 1 | 21/80 | 31.65% |
| 1 | 59/185 | 14.35% |
| 0.996 | 20/55 | 10.23% |
| 0.996 | 17-Sep | 15.75% |
| 1 | 27/69 | 11.69% |
| 1 | 63/209 | 14.35% |
| 1 | 22-Oct | 11.60% |

| | | |
|---|---|---|
| 0.996 | 23-Feb | 2.03% |
| 1 | 26/91 | 11.69% |
| 0.999 | 15-Mar | 2.31% |
| 1 | 66/215 | 14.08% |
| 1 | 79/276 | 13.03% |
| 0.999 | 15-Aug | 15.75% |
| 0.999 | 16-Oct | 16.24% |
| 1 | 20-Aug | 13.37% |
| 1 | Oct-37 | 6.03% |
| 1 | 13/72 | 4.81% |
| 1 | 17-Feb | 0.91% |
| 1 | 17-Feb | 0.91% |
| 1 | 28/119 | 11.28% |
| 1 | 25/83 | 17.08% |
| 1 | 23-Aug | 12.29% |
| 0.999 | 17-Aug | 9.95% |

| | | |
|---|---|---|
| 1 | 19-Jul | 2.44% |
| 1 | 17/78 | 7.11% |
| 1 | 18-Apr | 7.72% |
| 1 | 14/27 | 16.17% |
| 1 | Oct-34 | 9.68% |
| 1 | 19/76 | 8.62% |
| 1 | 25-Oct | 12.35% |
| 1 | 14/46 | 12.72% |
| 1 | 17-Apr | 2.41% |
| 1 | 16-Jul | 13.10% |
| 1 | 16-Jul | 13.10% |
| 1 | Oct-70 | 6.24% |
| 1 | 19/31 | 18.45% |
| 1 | 49/256 | 8.87% |

| | | |
|---|---|---|
| 1 | 122/252 | 18.95% |
| 1 | 15-May | 7.87% |
| 1 | 18-Jul | 6.48% |
| 1 | 29-Oct | 13.05% |
| 1 | 15/29 | 15.59% |
| 1 | 81/225 | 11.90% |
| 1 | 20-May | 4.25% |
| 1 | 14/27 | 15.58% |
| 1 | 15-Jun | 12.64% |
| 1 | 27-May | 7.11% |
| 1 | 25/44 | 20.30% |
| 1 | 15-May | 5.84% |
| 1 | 20-Jul | 13.05% |

| | | |
|---|---|---|
| 1 | 48/174 | 11.76% |
| 1 | 14/30 | 15.62% |
| 1 | 24-Sep | 10.87% |
| 1 | 13/27 | 16.17% |
| 1 | Aug-37 | 8.68% |
| 1 | 24-Sep | 13.82% |
| 1 | 21-Mar | 10.29% |
| 1 | 29-Oct | 12.35% |
| 1 | 47/249 | 9.49% |
| 1 | 28/69 | 17.60% |
| 1 | 16/55 | 12.72% |
| 1 | 72/275 | 12.65% |
| 1 | 68/258 | 13.12% |
| 1 | 15-Jul | 17.56% |
| 1 | 22-May | 3.24% |

| | | |
|---|---|---|
| 1 | 25-Feb | 0.06% |
| 1 | 108/327 | 11.90% |
| 1 | 16/67 | 10.54% |
| 1 | 18/32 | 16.92% |
| 1 | 46/243 | 9.49% |
| 1 | 16-May | 15.43% |
| 1 | 17/52 | 11.60% |
| 1 | 15-Jul | 14.81% |
| 1 | 70/230 | 11.48% |
| 1 | 30/148 | 8.87% |
| 1 | Nov-32 | 8.76% |
| 1 | 16-May | 6.15% |
| 1 | 18/76 | 13.05% |
| 1 | Jul-47 | 7.42% |

| | | |
|---|---|---|
| 1 | 41/187 | 11.16% |
| 1 | 27/109 | 8.44% |
| 1 | 17-Jun | 13.82% |
| 1 | 16/38 | 13.64% |
| 1 | 30-Dec | 10.72% |
| 1 | 23-Apr | 5.07% |
| 1 | 26-Aug | 8.76% |
| 1 | 32/86 | 15.28% |
| 1 | 20-Jun | 6.48% |
| 1 | 36/119 | 14.48% |
| 1 | 20-Jul | 14.91% |
| 1 | 20-Sep | 15.51% |
| 1 | Nov-38 | 10.35% |
| 1 | Aug-36 | 4.54% |
| 1 | 18-Oct | 13.57% |
| 1 | 16/47 | 15.29% |
| 1 | 33/62 | 17.81% |

| | | |
|---|---|---|
| 1 | 109/446 | 13.88% |
| 1 | 22/65 | 10.23% |
| 1 | Dec-53 | 11.16% |
| 1 | 35/67 | 17.81% |
| 1 | 24/93 | 9.03% |
| 1 | 18/27 | 18.17% |
| 1 | 66/148 | 17.72% |
| 1 | 16-Jul | 6.44% |
| 1 | 21-Jun | 5.23% |
| 1 | 24/56 | 16.84% |
| 1 | 26/96 | 8.62% |
| 1 | 18-Aug | 10.92% |
| 1 | 16/51 | 12.43% |
| 1 | Dec-39 | 11.91% |

| | | |
|---|---|---|
| 1 | 23-Jul | 5.51% |
| 1 | 21-Mar | 11.21% |
| 1 | 17-Mar | 8.01% |
| 1 | 15-May | 7.18% |
| 1 | 65/367 | 9.70% |
| 1 | 31-Jul | 12.89% |
| 1 | Dec-55 | 9.16% |
| 1 | Dec-32 | 9.48% |
| 1 | 21-Jul | 5.39% |
| 1 | 57/203 | 15.80% |
| 1 | Jun-32 | 3.43% |
| 1 | 16-May | 17.20% |
| 1 | May-34 | 6.37% |
| 1 | 19/62 | 16.84% |
| 1 | 60/315 | 11.89% |
| 1 | 13/55 | 7.78% |
| 1 | 17-Feb | 3.61% |

| | | |
|---|---|---|
| 1 | 20/84 | 9.40% |
| 1 | 16-Mar | 2.78% |
| 1 | 26/73 | 16.77% |
| 1 | 136/499 | 13.54% |
| 1 | 13/60 | 4.20% |
| 1 | 14/54 | 10.75% |
| 1 | 15/54 | 13.97% |
| 1 | 21-Jul | 16.80% |
| 1 | 16/81 | 9.96% |
| 1 | Dec-62 | 12.13% |
| 1 | 27-Oct | 11.20% |
| 1 | 20/95 | 10.56% |
| 1 | Jul-41 | 3.43% |
| 1 | 26-Mar | 4.53% |
| 1 | 26-Aug | 6.16% |
| 1 | 22-Mar | 2.12% |

| | | |
|---|---|---|
| 1 | May-33 | 7.42% |
| 1 | 21-Mar | 3.81% |
| 1 | 23/75 | 14.61% |
| 1 | 18-Oct | 20.87% |
| 1 | 37/82 | 18.31% |
| 1 | 16-Jul | 15.10% |
| 1 | 45/204 | 11.38% |
| 1 | 18-Jun | 6.91% |
| 1 | 20-Aug | 11.38% |
| 1 | 15-Jun | 17.23% |
| 1 | Dec-59 | 7.70% |
| 1 | 104/334 | 14.52% |
| 1 | 26/74 | 12.73% |

| | | |
|---|---|---|
| 1 | 121/317 | 15.86% |
| 1 | 21-Jun | 14.79% |
| 1 | 18-Jun | 10.75% |
| 1 | 17/53 | 14.54% |
| 1 | 18-Feb | 2.14% |
| 1 | Nov-59 | 4.27% |
| 1 | 99/432 | 12.99% |
| 1 | 29-Jul | 11.76% |
| 1 | 100/303 | 14.70% |
| 1 | 19/100 | 7.08% |

| | | |
|---|---|---|
| 1 | 76/326 | 13.97% |
| 1 | 25-Apr | 5.07% |
| 1 | 28-Nov | 16.33% |
| 1 | 111/457 | 12.20% |
| 1 | 16-Jun | 11.06% |
| 1 | 16-Jun | 11.06% |
| 1 | 23-Apr | 2.41% |
| 1 | Oct-46 | 10.82% |
| 1 | 29-Aug | 10.32% |
| 1 | 18-Mar | 2.48% |
| 1 | 23-Aug | 6.40% |
| 1 | Nov-91 | 4.09% |
| 1 | 16-Apr | 7.63% |
| 1 | 17-Jul | 14.91% |
| 1 | 18-Nov | 15.02% |
| 1 | 21-Aug | 11.97% |

| | | |
|---|---|---|
| 1 | 15-Aug | 8.32% |
| 1 | 29/117 | 13.48% |
| 1 | 17-Mar | 3.73% |
| 1 | 69/314 | 11.08% |
| 1 | 27-Nov | 15.87% |
| 1 | 20/72 | 10.92% |
| 1 | Apr-34 | 7.42% |
| 1 | 37/106 | 16.65% |
| 1 | Nov-81 | 4.66% |
| 1 | 24-Apr | 2.41% |
| 1 | 22-Sep | 20.32% |
| 1 | Aug-45 | 3.43% |
| 1 | 23-Nov | 10.60% |
| 1 | 19-Mar | 0.76% |
| 1 | 20-Jul | 13.45% |
| 1 | 21-Oct | 21.73% |
| 1 | 15-May | 6.98% |
| 1 | 19/48 | 20.28% |
| 1 | 26-Aug | 11.25% |

| | | |
|---|---|---|
| 1 | 52/183 | 16.77% |
| 1 | Nov-35 | 13.32% |
| 1 | Aug-58 | 12.01% |
| 1 | 15-May | 19.51% |
| 1 | Jun-42 | 5.43% |
| 1 | 19-Aug | 22.75% |
| 1 | 25-Sep | 9.65% |
| 1 | 16-May | 10.82% |
| 1 | 19/111 | 9.13% |
| 1 | 23-Jun | 8.23% |
| 1 | Nov-54 | 11.20% |
| 1 | Jul-36 | 7.72% |
| 1 | 16/92 | 9.13% |
| 1 | 26-Aug | 10.65% |
| 1 | 22-Dec | 22.46% |
| 1 | 25-Aug | 11.97% |
| 1 | 16-May | 6.26% |
| 1 | 23-Dec | 20.77% |
| 1 | 21-Dec | 22.46% |
| 1 | 15/32 | 17.87% |
| 1 | 15/56 | 7.21% |
| 1 | 27-Mar | 1.63% |
| 1 | 27-Dec | 17.51% |

| | | |
|---|---|---|
| 1 | 17/40 | 17.93% |
| 1 | 13/34 | 18.21% |
| 1 | 16/54 | 13.94% |
| 1 | Jun-51 | 3.26% |
| 1 | 16-Feb | 0.11% |
| 1 | 16-Mar | 3.13% |
| 1 | 19-Jun | 12.43% |
| 1 | Nov-44 | 10.92% |
| 1 | Jul-43 | 6.66% |
| 1 | 16-Jun | 8.32% |
| 1 | 19/97 | 10.72% |
| 1 | 17-Aug | 8.32% |
| 1 | 17-Mar | 4.95% |
| 1 | 17-Mar | 2.48% |
| 1 | 16/26 | 21.35% |
| 1 | 23-Sep | 15.32% |
| 1 | 14/39 | 16.28% |
| 1 | 14/37 | 22.67% |
| 1 | 17-Feb | 1.16% |
| 1 | 17/74 | 12.11% |

| | | |
|---|---|---|
| 1 | 58/243 | 11.40% |
| 1 | May-44 | 6.16% |
| 1 | 15-Apr | 3.30% |
| 1 | 17-Jul | 11.20% |
| 1 | 23-Mar | 2.59% |
| 1 | 23/91 | 16.25% |
| 1 | 30-Dec | 15.78% |
| 1 | 19-May | 14.56% |
| 1 | Feb-29 | 1.18% |
| 1 | 26-Jun | 6.91% |
| 1 | Oct-41 | 10.37% |
| 1 | 29-Apr | 6.47% |
| 1 | 54/296 | 11.93% |
| 1 | 21/85 | 16.25% |
| 1 | 18-Sep | 16.63% |
| 1 | 21/92 | 14.01% |
| 1 | 72/475 | 8.30% |
| 1 | 22-Feb | 2.00% |

| | | |
|---|---|---|
| 1 | 23-May | 7.22% |
| 1 | 15/33 | 17.87% |
| 1 | 19-Feb | 1.63% |
| 1 | 15-Jun | 11.84% |
| 1 | 18-Apr | 8.29% |
| 1 | 15-May | 5.39% |
| 1 | 22-Apr | 9.40% |
| 1 | 15-Apr | 19.36% |
| 1 | 16-Jun | 18.97% |
| 1 | 17-Nov | 17.77% |
| 1 | 22-May | 4.03% |
| 1 | 22-Dec | 17.85% |
| 1 | 14/74 | 7.31% |
| 1 | 15/35 | 15.03% |
| 1 | 36/96 | 15.55% |
| 1 | Oct-40 | 11.94% |
| 1 | Dec-37 | 18.76% |
| 1 | 18-Jan | 3.17% |
| 1 | 26/109 | 12.22% |
| 1 | 22-Jul | 13.99% |
| 1 | 20-Mar | 2.48% |
| 1 | 16-Jul | 23.03% |
| 1 | 15-May | 10.51% |
| 1 | 24-Jul | 10.13% |

| | | |
|---|---|---|
| 1 | Dec-46 | 12.40% |
| 1 | 11/106 | 3.18% |
| 1 | 13/40 | 12.00% |
| 1 | 14/62 | 11.02% |
| 1 | 29/140 | 9.35% |
| 1 | 18-Sep | 13.35% |
| 1 | 21-Jul | 13.06% |
| 1 | 21-Sep | 10.60% |
| 1 | 19/93 | 12.11% |
| 1 | Apr-33 | 2.48% |
| 1 | 32/140 | 10.99% |
| 1 | Sep-51 | 12.12% |
| 1 | Oct-35 | 12.84% |
| 1 | 23/48 | 18.06% |
| 1 | 16/35 | 17.68% |
| 1 | 13/29 | 19.30% |
| 1 | 13/33 | 15.43% |
| 1 | 22/76 | 11.84% |
| 1 | 15-Apr | 5.51% |
| 1 | Jul-36 | 5.31% |
| 1 | Aug-42 | 11.59% |

| | | |
|---|---|---|
| 1 | Oct-75 | 6.40% |
| 1 | May-35 | 3.81% |
| 1 | Dec-61 | 9.16% |
| 1 | 20-Apr | 6.33% |
| 1 | 27-Oct | 16.36% |
| 1 | 20-Jun | 9.92% |
| 1 | 23/72 | 12.67% |
| 1 | 29-Apr | 9.03% |
| 1 | 27-Feb | 3.79% |
| 1 | 29-Jun | 8.16% |
| 1 | 15/42 | 15.43% |
| 1 | 20/47 | 19.57% |
| 1 | 18/47 | 17.42% |
| 1 | 22-Apr | 4.41% |
| 1 | 16/25 | 18.65% |
| 1 | 13/33 | 15.11% |
| 1 | 16/46 | 13.96% |
| 1 | 20/76 | 19.13% |
| 1 | 18-May | 3.82% |
| 1 | 28-Feb | 0.99% |
| 1 | 14/134 | 6.03% |

| | | |
|---|---|---|
| 1 | 24-Aug | 17.92% |
| 1 | 54/199 | 15.22% |
| 1 | 21/91 | 16.25% |
| 1 | 13/46 | 8.50% |
| 1 | 29-Mar | 3.63% |
| 1 | 17-Aug | 14.72% |
| 1 | 21-Apr | 5.30% |
| 1 | 16/44 | 15.51% |
| 1 | Dec-55 | 9.06% |
| 1 | 18/42 | 12.82% |
| 1 | 58/268 | 14.84% |
| 1 | 23-Jul | 18.59% |
| 1 | 29-Sep | 17.54% |
| 1 | Sep-43 | 7.92% |
| 1 | 28-Apr | 10.02% |
| 1 | 25-Jul | 13.99% |
| 1 | 15-Jun | 11.91% |
| 1 | 25-Jul | 13.49% |
| 1 | Oct-42 | 10.87% |

| | | |
|---|---|---|
| 1 | 20-Feb | 0.99% |
| 1 | 20-Jun | 19.07% |
| 1 | 16-Jun | 7.65% |
| 1 | 16-Jul | 21.70% |
| 1 | 16-Feb | 4.77% |
| 1 | 16-Feb | 4.77% |
| 1 | Nov-66 | 12.13% |
| 1 | Aug-52 | 7.08% |
| 1 | Nov-55 | 10.02% |
| 1 | Dec-35 | 14.24% |
| 1 | 14/28 | 17.85% |
| 1 | Mar-33 | 2.58% |
| 1 | 16/37 | 12.82% |
| 1 | 31-Aug | 9.15% |
| 1 | 23/60 | 17.05% |
| 1 | 18/41 | 21.75% |
| 1 | 19-Feb | 1.99% |
| 1 | Sep-37 | 11.14% |
| 1 | 20-Jun | 15.87% |
| 1 | 48/173 | 15.22% |
| 1 | Nov-85 | 6.61% |

| | | |
|---|---|---|
| 1 | 20-Mar | 2.19% |
| 1 | 19-Feb | 0.16% |
| 1 | Sep-34 | 15.05% |
| 1 | 22-Jul | 15.33% |
| 1 | 16-Mar | 8.24% |
| 1 | 20-May | 16.11% |
| 1 | 13/31 | 15.62% |
| 1 | 35/101 | 16.55% |
| 1 | 22/54 | 19.39% |
| 1 | 22-Jun | 6.87% |
| 1 | 16-Feb | 3.81% |
| 1 | 21/42 | 16.84% |
| 1 | 20-May | 10.02% |
| 1 | 19-Apr | 6.61% |
| 1 | 14/58 | 17.68% |
| 1 | Oct-36 | 11.37% |
| 1 | 25-May | 7.68% |
| 1 | 15-Aug | 13.35% |
| 1 | Jun-38 | 8.85% |
| 1 | 18-Sep | 21.93% |
| 1 | 16/33 | 18.17% |
| 1 | 15-Jan | 0.80% |

| | | |
|---|---|---|
| 1 | 26-Sep | 17.14% |
| 1 | 25-Aug | 16.55% |
| 1 | 22-May | 11.25% |
| 1 | 28-Mar | 8.79% |
| 1 | Jan-40 | 0.07% |
| 1 | Jun-61 | 6.91% |
| 1 | 19-Jun | 12.28% |
| 1 | Nov-35 | 11.48% |
| 1 | 18-Aug | 19.30% |
| 1 | Oct-37 | 15.08% |
| 1 | 21-Jun | 19.40% |
| 1 | 21-Jun | 19.40% |
| 1 | 13/38 | 17.86% |
| 1 | Sep-51 | 6.91% |
| 1 | Sep-46 | 12.19% |
| 1 | 30-Jul | 11.06% |
| 1 | 18-Apr | 8.68% |
| 1 | 24-Aug | 16.28% |
| 1 | 13/28 | 19.88% |
| 1 | Nov-44 | 9.53% |
| 1 | 27-Jun | 8.41% |
| 1 | 27-Aug | 11.92% |
| 1 | 30-Aug | 11.76% |
| 1 | 15-Mar | 12.72% |

| | | |
|---|---|---|
| 1 | 22-Jul | 13.26% |
| 1 | 22-Jul | 13.26% |
| 1 | 28-Feb | 2.12% |
| 1 | 22-Apr | 6.15% |
| 1 | 28-May | 6.07% |
| 1 | Oct-36 | 13.42% |
| 1 | 22-Jul | 15.20% |
| 1 | Sep-31 | 17.67% |
| 1 | 16/37 | 20.75% |
| 1 | 16/35 | 24.25% |
| 1 | 16/35 | 24.25% |
| 1 | 15/51 | 18.21% |
| 1 | 20-Aug | 17.76% |
| 1 | 21-Jul | 13.75% |
| 1 | 15-Mar | 6.07% |
| 1 | 31-Dec | 15.82% |
| 1 | 16-May | 16.11% |
| 1 | 15/33 | 17.87% |
| 1 | 16/48 | 18.65% |
| 1 | Sep-34 | 14.06% |
| 1 | 24-Jul | 13.37% |
| 1 | 22-Feb | 4.23% |
| 1 | 20-Apr | 10.03% |

| | | |
|---|---|---|
| 1 | 29-Sep | 13.44% |
| 1 | 15-Jan | 0.92% |
| 1 | 19-Feb | 9.29% |
| 1 | 25-Mar | 11.21% |
| 1 | 22-Aug | 20.82% |
| 1 | May-37 | 9.96% |
| 1 | 25-Sep | 14.33% |
| 1 | 22-Mar | 11.21% |
| 1 | 22-Mar | 11.21% |
| 1 | Oct-37 | 16.93% |
| 1 | Oct-53 | 13.06% |
| 1 | 21-Apr | 8.75% |
| 1 | 19-Jan | 0.94% |
| 1 | 15-Aug | 25.06% |
| 1 | 18-May | 16.11% |
| 1 | 20-Nov | 22.62% |
| 1 | 17-Jan | 1.39% |
| 1 | 15-Feb | 12.54% |

Lead_genes

OAS2;OAS1;OASL;OAS3;TENT4B;PAPOLG

CXCL10;GPNMB;LGALS3;PDGFB;VEGFB;PGF;VEGFC;FGF2

LARGE1;B4GAT1;CSGALNACT1;CHPF;CHSY1;EXT1;B3GAT3

MCM7;RFC2;DHX9;POLQ;RFC3;RAD51;DSCC1;RFC4;MCM2;HELQ;WRNIP1;HELB;DNA2

GEM;PRKG1;STIMATE;NRXN1;SGK1;SRI;RRAD;CACNG8;FKBP1A;CACNB3;MCUB

CYP27A1;CH25H;POR;MSMO1;KMO;CYP27B1;MICAL3;FAXDC2;MICAL2

GAL;HCRTR2;SORCS1;GALR2;GPR143;SSTR2

TIMP3;DPP4;ATP9A;ITGA3;PZP;SERPINE1;CSTB;ECM1;SELL;PRKN;FN1;VWF;BCL2;CD70;NOL3;COL1A2;RIPK2;ITGB3;CRADD;A2M;F3;TIMP2;LDLR;RHBDD2;SRI;BRCA2;TYSND1;TIMP4;TNFRSF10A;UBC;ITGB1;FAP

PLAAT3;PLAAT4;PLA2G7;PNPLA3;PLA2G4C;PLA2G2D;PLAAT1;PNPLA8;PLA2G6;PLA2G12A;PRDX6;PLA2G15

MME;DPP4;CPE;ERAP2;LAP3;ENPEP;PRCP;CTSH;CTSZ;XPNPEP2;DPP7;ERAP1;DPEP3;RCE1;FAP;TRHDE;ANPEP;DPP6

SETD3;WDR82;KMT2C;KMT2D;SETD1A;CXXC1;SETD1B;ASH2L;KMT2B;RBBP5;SETD4;WDR5

MMP7;MME;UCHL1;MMP19;DPP4;USP2;ADAMTS15;USP18;ADAMDEC1;ADAMTS6;CTSW;MMP9;HTRA4;BACE2;CASP7;PIK3IP1;PSEN2;CTSV;LGMN;CTSF;CASP9;MMP12;CTSL;USP11;OMA1;ADAMTS2;ECE1;PAPLN;PCSK6;ADAMTS9;USP36;F3;USP49;RHBDF2;CTSH;CTSZ;HTRA2;MMP17;PSMB2;MST1;PRSS21;PHEX;PSMA5;TYSND1;CTSB;HTRA3;MMP2;USP37;RCE1;RHBDF1;PSMA4;FAP;PSMB5;PSMB10;ADAMTS19;ATG4B;C1S;KLK1;ADAM28;BACE1;MEP1A;MIPEP;ADAMTSL1;PSMA2;CASP6

DHRS9;RDH10

EIF4E3;EIF3L;EIF2S3;EIF3A;EIF3J;EIF3B;EIF4G2;EIF3H;EIF1;EIF3E;EIF3F;EIF5B;EIF3M;EIF2B3;EIF2S3B;EIF2S2;EIF2A;EIF2B5;EIF3C;EIF4G3;EIF4G1

CYP2S1

PARP9;PARP14;PARP15;PARP10;ZC3HAV1;PARP3;TIPARP;PARP4;SIRT5;PARP16;SIRT1;ART4;SIRT4;PARP1

UCHL1;USP2;USP18;CTSW;CLC;CASP7;CAPN5;CTSV;LGMN;CTSF;CASP9;CTSL;USP11;USP36;USP49;CTSH;CTSZ;CTSB;USP37

CX3CR1;ADGRG1;FFAR4;FPR1;CNR2;ADGRE1;CALCRL;ADGRG6;CMKLR1;GPBAR1;GPR84;GPR141;P2RY13;ADGRG3;P2RY6;ADGRE3;GPR34;LPAR1;PTAFR;ADGRG5;CXCR2;CYSLTR1;GPR18;GPR183;P2RY10;CALCR;P2RY12;GIPR;ADRB1;FFAR2;GPR35;S1PR2;FPR2;GPR27;TAPT1

HBE1;CYP27A1;CYP19A1;HBA1;CYB5A;HBG2;HBG1;HBD;GUCY1B1;ABCB6;STEAP4;HBZ;SLC48A1;FLVCR2;STC2;HEBP1;CYP2U1;PGRMC2;PXDN

PLTP;ABCG1;STARD5;ABCA1;STARD4

GJA1;GJC1;PANX1;PANX2;GJC2;GJA5

MMP7;MME;MMP19;ADAMTS15;ADAMDEC1;ADAMTS6;MMP9;MMP12;OMA1;ADAMTS2;ECE1;PAPLN;ADAMTS9;MMP17;PHEX;MMP2;RCE1;ADAMTS19;MEP1A;MIPEP;ADAMTSL1

H1-5;H1-3;H1-4;H1-1;HDAC1;H1-2;GATAD2B;MTA2;ACTL6A;RBBP4;RCC1;HDAC2;HNRNPC

CDC6;CDC45;MCM10;MCM7;MCM4;DHX9;ORC1;ORC5;HSPD1;MCM8;MCM2;KAT7;MCM9;MCMDC2;ORC2;MCM3

GJA1;GJC1;PANX1;PANX2;GJC2;GJA5

KPNB1;IPO9;IPO5;KPNA6;KPNA3;TNPO3;TNPO1;TNPO2;KPNA4

PLTP;ABCG1;STARD5;ABCA1;STARD4

OR51B5;OR8G5;OR1J2;OR8G1;OR10D3;OR8B2;OR1F1;OR1B1;OR8D1;OR6J1;OR7D2;OR8B8;OR8A1;OR1L8;OR1L6;OR8B3;OR9Q1;OR51M1;OR51B2;OR52A5;OR7C1

GBP1;GEM;MX2;GBP4;ATL1;GBP3;RAB38;RAB13;GBP2;TUBA8;RRAS2;RHOB;GBP5;MRAS;RHOC;TUBB3;RHEBL1;RRAS;TUBA4A;TUBE1;TGM2;RAC2;DIRAS1;MFHAS1;ARL4A;GBP7;RRAGD;RND3;TUBA1A;TUBG2;DIRAS2;RHOBTB2;RRAD;RAB42;RAB8A;ARL4D;SEPTIN6;ARL2;ARL5B;ARL8A;RASEF;RHEB;SEPTIN4;RAP2B;RALA;RND1;INSR;ARL6;ARL1;TUBG1;RRAGA;GCH1;NRAS;GTPBP10;TUBB1;RAP1B;RAB11B;RABL6;EIF2B2

KIF11;KIF2C;KIF21B;KIF14;KIF4A;KIF23;KIF18B;KIF20A;STARD9;DNAH17;CENPE;KIF22;KIF7;KIF18A;KIF3C;KIFC2;KIF1C;KIF20B;DNAH8;KIF24

MTMR4;SACM1L;MTMR14;SYNJ1;MTMR10;MTMR8;PTEN;MTMR3;MTM1

PLAAT3;PLAAT4;CLC;MGLL;PLCL1;SMPD1;SMPDL3A;PNPLA3;PLA2G4C;ABHD4;LPL;ABHD6;PNPLA7;FAM83B;LIPG;PLA2G2D;GPLD1;PLAAT1;PNPLA8;PLD6;PLA2G6;PLA2G12A;PRDX6;DDHD1;SMPD3;ENPP2;PLA2G15

GBP1;GEM;MX2;GBP4;ATL1;GBP3;RAB38;RAB13;GBP2;TUBA8;RRAS2;RHOB;GBP5;MRAS;RHOC;TUBB3;RHEBL1;RRAS;TUBA4A;TUBE1;TGM2;RAC2;DIRAS1;MFHAS1;ARL4A;GBP7;RRAGD;RND3;TUBA1A;TUBG2;DIRAS2;RHOBTB2;RRAD;MPPED2;RAB17;RAB42;RAB8A;ARL4D;SEPTIN6;ARL2;ARL5B;ARL8A;RASEF;RHEB;PDE2A;SEPTIN4;RAP2B;RALA;PDE10A;RND1;INSR;ARL6;ARL1;TUBG1;RRAGA;GCH1;NRAS;GTPBP10;TUBB1;RAP1B;RAB11B;RABL6;EIF2B2

ACP5;HAAO;PHYH;ISCU;FECH;CDO1;FTL;FTH1;EGLN3;ALKBH2

RXRA;PYGL

PLAAT3;PLAAT4;CLC;MGLL;PLA2G7;PNPLA3;PLA2G4C;ABHD4;CA2;H6PD;LIPA;LPL;ABHD6;PNPLA7;LIPG;PLA2G2D;ABHD5;ACHE;PLAAT1;PLA2G6;PLA2G12A;PRDX6;ESD;ENPP2;PTRH1;PLA2G15

CLEC4A;CD207;CLEC17A

MX1;GBP1;GEM;MX2;GBP4;ATL1;GBP3;RAB38;GBP2;RRAS2;GNG11;GNA11;RHOB;GBP5;MRAS;RHOC;RHEBL1;IFT22;RAB3A;RRAS;GNAL;RAC2;RAB20;DIRAS1;DNM1;GBP7;RRAGD;RAB6B;SEPTIN8;RND3;DIRAS2;RAB3C;DNM3;RHOBTB2;GTPBP1;RAB17;RAB42;RAB8A;RAP1GAP;SEPTIN6;RAB11A;RAB1A;GNA13;ARL2;GNA12;ARL8A;RHEB;SEPTIN4;RAP2B;RALA;RAB24;RND1;ARL1;IRGM;MFN1;TBCC;RRAGA;GCH1;RABL2B;SEPTIN11;NRAS;GTPBP10;SEPTIN9;RAP1B;RAB11B;RASD2

MX1;GBP1;GEM;MX2;GBP4;ATL1;GBP3;RAB38;TDRD9;GBP2;RRAS2;GNG11;GNA11;MYO1E;DYNC2H1;DYNC1I1;TOR1B;RHOB;GBP5;MRAS;RHOC;RNF213;RHEBL1;IFT22;RAB3A;RRAS;GNAL;ABCB6;MYO6;ABCA3;RAC2;TOR3A;RAB20;DIRAS1;DNM1;GBP7;RRAGD;RAB6B;SEPTIN8;RND3;THTPA;DIRAS2;RAB3C;DNM3;RHOBTB2;GTPBP1;RAB17;RAB42;RAB8A;RAP1GAP;SEPTIN6;RAB11A;RAB1A;ABCB10;GNA13;ARL2;SMCHD1;MYL6B;GNA12;TOR1A;PIN1;ARL8A;RHEB;SEPTIN4;RAP2B;RALA;ABCD1;RAB24;RND1;ARL1;IRGM;MFN1;TBCC;RRAGA;GCH1;RABL2B;SEPTIN11;NRAS;MYL6

MTMR4;MTMR14;SYNJ1;MTMR10;MTMR8;PTEN;MTMR3;MTM1

MCM7;RECQL4;DHX9;FBH1;BLM;ERCC3;MCM2;HELQ;RECQL;RECQL5

SLC5A3;SLC5A4;SLC2A5;SLC2A10;SLC2A14;MFSD4A;SLC2A8;SLC2A3

PARP9;PARP14;PARP15;PARP10;LARGE1;ZC3HAV1;TYMP;UPP1;PARP3;TIPARP

OAS2;OAS1;OASL;OAS3;IFIH1;DDX60;DHX58;TFRC;EIF2AK2;ADARB1;ZBP1;ADAR;RFTN1

ANO5;TTYH2

ANO5;TTYH2

MMP7;MME;MMP19;ADAMTS15;ADAMDEC1;ADAMTS6;ERAP2;MMP9;LAP3;MMP12;OMA1;ADAMTS2;ECE1;PAPLN;ADAMTS9;MMP17;PHEX;ERAP1;MMP2;RCE1;ADAMTS19;ADAM28;MEP1A;MIPEP;PAPPA2;ADAMTSL1;ANPEP;MPND

HSD3B7;DHRS9;ME3;HPGD;DHDH;CBR1;CBR3;RDH10;H6PD;G6PD;HADH;BDH2;MDH1;LDHC;HSD17B7;ADH5;CBR4;HSD17B13;PTGR1;KDSR;DHRS3;DCXR;HIBADH;HMGCR;KCNAB1

ADCY8;FOXO1;MASTL;MFHAS1;SPHK1;DAB2IP;PTPA;PPME1

TRPM2;TPCN1;TRPV1;PKD2;MCOLN2;FKBP1B;ITPR1;RYR1

APOBEC3G;APOBEC3H;APOBEC3F;APOBEC3A;APOBEC3C;ADA;APOBEC3D

UCHL1;USP2;USP18;CTSW;CASP7;CTSV;LGMN;CTSF;CASP9;CTSL;USP11;USP36;USP49;CTSH;CTSZ;CTSB;USP37

HSD3B7;DHRS9;HSD17B14;HSD17B1

POLR3B;POLR2B;CRCP;POLR2J;POLR2L;POLR3A;POLR1A;POLR1B;POLR2D;POLR1D;POLR2G;PRIM1;POLR1C;POLR3H

CHIT1;CHI3L1;GUSB;MAN1A1;HYAL2;ADPRS;PGGHG;ACER2;GLA;HPSE2

MDK;INHBA;AREG;CSF1;TGFA;CLCF1;EGF;FGF9;BDNF;PDGFB;IL7;EREG;DKK1;PDGFA;VEGFB;IL6;PGF;VEGFC;FGF2

GOLGA7B;ZDHHC24;GLUL;ZDHHC9;ZDHHC1;ZDHHC19;ZDHHC11B;ZDHHC4;ZDHHC12;ZDHHC8

NR1H3;ABCG1;STARD5;SOAT1;OSBP2;SCARB2;SYP;ABCA1;STARD3NL;OSBPL1A;STARD4;STARD3;SIDT1;CD81

CX3CR1;CXCR3;CMKLR1;CCR2

ATP6V1C2;TCIRG1;ATP6V0E2;ATP6V0C;ATP6V1D;ATP6V0B;ATP6V0A2

ATP6V1C2;TCIRG1;ATP6V0E2;ATP6V0C;ATP6V1D;ATP6V0B;ATP6V0A2

CX3CR1;EDNRA;FPR1;CALCRL;GPR84;CYSLTR1;CALCR;GIPR;S1PR2;FPR2

ARRB1;CLOCK;KAT14;TADA3;MCM3AP;KAT6A;SRCAP;TAF1;ATF2;USP22;KAT7;KAT8;EP300;NAA40;EPC1;NAA60;NAA50;CREBBP;EPC2

CXCL10;MDK;GDF15;DPP4;BMP2;SEMA5A;TNFSF13B;INHBA;AREG;IFNB1;BMP7;BMP6;IFNA2;CSF1;TGFA;GPNMB;SEMA3D;LGALS3;IL15;CLCF1;TGFB2;EGF;CD70;FGF9;BMP8A;IFNA8;LRPAP1;POMC;BDNF;IFNA14;NPPC;PDGFB;IL7;EREG;DKK1;PDGFA;SEMA3C;IFNA1;TAFA2;WNT1;IFNW1;VEGFB;IL6;SEMA3A;PGF;VEGFC;KITLG;FGF2;CXCL9

SLPI;PCBP3;ELAVL4;DHFR;SRSF1;SRRM3;CPEB3;NCL;RPS3;TARDBP;CSTF3;HNRNPM;RPL5;TYMS;RPL7;RBM4;NCBP1;QKI;HNRNPAB;RPS7;DHX9;RBM14;RPL13A;RPS13;CARHSP1;RBMX;RBMS2;LUC7L3;FUBP3;HNRNPA2B1;EIF3A;HDLBP;CELF2;PCF11;RPL26;RC3H1;EIF4G2;DDX5;RPS26;LSM14B;GRSF1;PABPC1;CCT5;FUBP1;THOC5;YTHDF1;RBM15;LARP4;LUC7L2;KHDRBS1;RBM4B;IREB2;SYNCRIP;IGF2BP1;ZFP36L1;RC3H2;HNRNPK;PTBP1;PUM3;AQR;PUM2;RBMS1;PUM1;SF1;LSM14A;UTP23;LARP4B;METTL3;SRSF6;RPS14;HNRNPC;RPL35;SF3B1;DAZAP1;RBM25;NCBP3;EIF2S2;PABPC4;EIF2A;RBFOX1;UPF3B;RSL1D1;SNRNP70;SCAF8;RBM5;FYTTD1;ETF1;RPS5;CPSF6;DDX3X;SERBP1;ANGEL2;FXR1;CSTF2;EDC3;SUPT5H;XPO5;NCBP2;GEMIN5;TRA2B;CLUH;SRSF4;HNRNPU;EIF4G3;EIF4G1;TPR;LUC7L;PARK7;RNF40;CELF1;SECISBP2;HNRNPLL;SHMT2;LARP1;RBPMS;YTHDF2;HNRNPA0;GNL3;RPS3A;PURB;SAMD4B;DHFR2

RAB3A;RAB6B;RAB3C;RAB8A;RAB11A

PLTP;PITPNM2;ABCG1;ABCB1;ABCA3;ABCA1;ATP10A

DGAT2;LPCAT1;PLA2G4C;SOAT1;MBOAT1;PLAAT1;LPCAT3;PRDX6;PLA2G15;MOGAT1

TUT4;HNRNPA1;RBM4;ZC3H7A;HNRNPA2B1;TUT7;RC3H1;PUM2;AGO3;PUM1;ZNF346;RBM10;DDX21;MATR3;ZC3H10

PREX2;RGS6;NRP1;PREX1;ARHGAP31;ARHGAP22;RHOH;LRRK2;RGS16;RGS10;AGAP2;ARHGAP9;RGS12;ARHGAP35;GARNL3;TBC1D12;ARHGEF6;SH3BP4;VAV3;ADAP1;TBC1D16;SIPA1L1;ARRB1;OPHN1;TBC1D4;ARHGAP18;DEPDC5;RIMS1;ARAP3;ARHGAP15;RACGAP1;RASA4;ARHGAP45;CHN1;TAGAP;ARHGAP26;TBC1D10A;IPO7;EVI5L;RIC8B;SH3BP1;DOCK1;RGS3;SIPA1L3;ABR;EVI5;BNIP2;GNAQ;ARHGEF10L;NCKAP1L;SIPA1;TBC1D13;WAS;ARHGAP19;TBC1D22A;ALS2;LARS1;NF1;ELMOD3;TBC1D14;CDC42SE1;RGS14;RALGAPA2;TBCK;GNB5;RGS7;STARD8;ARHGAP1;RP2;RALGDS;ARHGAP33;DNM1L;RASGRP3;IPO5;KRIT1;ACAP2;RGS8;SMAP1;ECT2;RABGAP1;RALGAPB

SMAD3;THRB;RORA;AR;ESR1

STAT1;TRAF5;TNFSF9;TNFSF18;TRAF1;MYD88;TMBIM1;FEM1B;TRAF4;DAB2IP;RIPK1;TRAF3;PIDD1;TRAP1

SCIN;SNX9;SNX10;SCARB1;WDFY1;ZFYVE16

PPP1R1A;MYOZ1;LGALS3;PPP1R17;LMTK2

SETD3;WDR82;KMT5B;SETD5;ASH1L;EED;KMT2C;KMT5A;SETDB1;NSD1;SETD2;SETD7;KMT2D;SETD1A;EHMT1;CXXC1;SETD1B;SMYD2;ASH2L;KMT2B;RBBP5;MECOM;SUV39H2;SETD4;WDR5

NCF2;CYBB;NCF1;NCF4;MICAL1

DGAT2;LPCAT1;PNPLA3;AGPAT4;ABHD5;PLA2G15;MOGAT1

SCIN;GBP1;ADCY8;MYOZ1;TPM2;NRAP;MLPH;KCNMA1;PRKCE;PDLIM4;V
CL;PLEKHH2;CXCR4;PRKN;SORBS1;ABLIM1;CALD1;IQGAP2;MYO6;SPTB
N4;TMSB10;TLN2;SYNE2;DAG1;MICAL3;SYNPO2;PACRG;BCL7B;FXYD5;C
EACAM1;PAWR;ITGB1;MICAL2;LIMA1;MAP1A;EPS8;FERMT2;ABL2;MYH10
;MAP1B;NCALD;RUSC1;EMD;DMD;LCP1;COBL;TPM1;PFN4
TRPV4;TRIM36;RACGAP1;WASHC1;FNTA;CDK5R1;SETD2;TBCEL;SNCA;AR
L4C;TAOK1;OFD1;TRAPPC14;SPAST

PALLD;BSG;CD47;JAM3;VSTM2L;NEXN;BCAM;ROBO3;LRRC4C
POLR3B;POLR2B;CRCP;POLR2J;POLR2L;POLR3A;POLR1A;POLR1B;POL
R2D;POLR1D;POLR2G;POLR1C;POLR3H

PLAAT3;PLAAT4;NAT14;TGM2;NAA80;PLAAT1;NAA10;SAT1

CMPK2;ITPKB;AK8;AK3;NME4;ITPKC;PMVK;DTYMK;AK5

GAL;ASIP;PNOC
GOLGA7B;ZDHHC24;GLUL;ZDHHC9;ZDHHC1;ZDHHC19;ZDHHC11B;ZDHH
C4;ZDHHC12;ZDHHC8
HERC5;RNF128;SH3RF2;BIRC3;FBXO6;RNF144A;TRIM2;EGR2;PELI2;TRI
M22;NEURL3;DTX4;CBLB;TRIM5;PRKN;MDM2;ZNRF1;RNF150;RNF122;D
TX3L;TRIM14;DTX1;TRIM55;MARCHF8;TRIM9;TRIM54;TRIM72;TRIM21;BI
RC7;MARCHF2;SH3RF3;UBE2D1;FBXO27;MYLIP;IRF2BPL;PELI3;HECTD1;
TRAF3IP2;TRIM56;NHLRC1;TRIM25;NEURL2;RNF13;TRIM35;RNF19A;STU
B1;UBE2D3

BBC3;AR;TMEM106B;TCIRG1;ANK1;TRPC1;NCSTN;ESR1;ATP1B3;ATP1B1
;ABCA1;SNX10;S100A1;PEX19;WFS1;ATP6V1G1;RALA;USP25;DERL1;ATP6
V0A2;CLPP;ATP6V1E1;ANK2;GABARAPL2;TMTC4;NR1H2;PDE4D;SELENOS
NR1H3;ABCG1;RORA;INSIG1;STARD5;SOAT1;OSBP2;SCARB2;SYP;ABCA1;
STARD3NL;OSBPL1A;STARD4;STARD3;SIDT1;CD81
ISG15;UCHL1;SMAD3;UBE2L6;USP2;FOXO1;PARP9;STAT1;EGR2;TRAF5;F
ZD6;ABCB1;VCL;STAT2;CXCR4;TRAF1;PML;PRKACB;PRKN;MDM2;PER1;B
CL2;SMC6;SPART;DTX3L;GABARAPL1;CD40;MID1;MFHAS1;TRIB3;DTX1;C
KB;PYHIN1;TRIB2;ATP6V0C;DDRGK1;JUN;CDKN1A;SNX9;BLZF1;PACRG;B
AG1;UBE2D1;UBE2D4;NFKBIA;SYT11;UBC;UBE2J1;HM13;TRAF4;PPARGC
1A;WFS1;NAE1;PRR7;STUB1;FBXO7;ANKRA2;PRKAR1A;PRR5L;FHIT;PRDX
6;RALA;CAMLG;RELA;PINK1;ARRDC5;YWHAZ;RIPK1;USP25;RRAGA;CALR;
DERL1
ISG15;UCHL1;SMAD3;UBE2L6;USP2;FOXO1;EGR2;TRAF5;FZD6;ABCB1;V
CL;CXCR4;TRAF1;PML;PRKACB;PRKN;MDM2;PER1;BCL2;SMC6;SPART;G
ABARAPL1;CD40;MID1;MFHAS1;TRIB3;DTX1;CKB;PYHIN1;TRIB2;ATP6V0C
;JUN;CDKN1A;SNX9;BLZF1;PACRG;BAG1;UBE2D1;UBE2D4;NFKBIA;SYT11
;UBC;UBE2J1;HM13;TRAF4;PPARGC1A;WFS1;NAE1;STUB1;FBXO7;ANKRA
2;PRKAR1A;PRR5L;FHIT;PRDX6;RALA;CAMLG;RELA;PINK1;ARRDC5;YWHA
Z;RIPK1;USP25;RRAGA;CALR;DERL1;RNF8;TRAF3

PIK3C2B;PIK3CD;PIK3C2A;PI4KB;PIK3C3;PIK3CG;PI4KA

PLTP;PITPNM2;PRELID2;GLTPD2;ABCA3

ALDH1A2;CRABP2

FGD5;PREX2;RGS6;NRP1;PREX1;ARHGAP31;RCBTB2;ARHGAP22;LRRK2;RGS16;RASGRP2;RGS10;DOCK10;RASGRP4;DOCK2;RAB3IL1;AGAP2;ARHGAP9;ARHGAP35;GARNL3;TBC1D12;ARHGEF6;VAV3;ELMO1;ADAP1;VAV1;TBC1D16;SIPA1L1;ARRB1;OPHN1;TBC1D4;ARHGAP18;FGD3;DEPDC5;RIN3;ARAP3;ARHGAP15;RACGAP1;RASA4;FGD4;ARHGAP45;DOCK4;ADRB1;IQSEC2;CHN1;TAGAP;ARHGAP26;TBC1D10A;EVI5L;RIC8B;SH3BP1;DOCK1;RGS3;SIPA1L3;ABR;DENND2B;HPS1;EVI5;BNIP2;RAP1A;GNAQ;ARHGEF10L;NCKAP1L;SIPA1;GAPVD1;TBC1D13;ARHGAP19;DENND4B;TBC1D22A;ALS2;LARS1;NF1;ELMOD3;RIC1;TBC1D14;RGS14;ANKRD27;DOCK11;RALGAPA2;TBCK;GNB5;NUCB2;TIAM1;DENND1A;RGS7;STARD8;ARHGEF16;ARHGAP1;RP2;MYCBP2;RAPGEF6;ARHGAP33;RCC1;DNM1L;DIS3;MADD;RASGRP3;RAPGEF4;DENND1C;ACAP2;RGS8;EIF2B3;ARHGEF2;CYTH3;SMAP1;ECT2;RABGAP1;RALGAPB

GBP1;GEM;RRAS2;MRAS;RHEBL1;RRAS;DIRAS1;RRAGD;DIRAS2;RAB17;RAB42;RAB8A;RASEF;RHEB;RAP2B;RALA

NCL;HNRNPD;HNRNPA2B1;XRCC5;SMG7;UPF2;SMG5;UPF1;BLM;ZBTB48;SMG6;RPA1;SMG1;HMBOX1;TERF1;UPF3A;KDM1A;XRCC6

HERC5;RNF128;SH3RF2;BIRC3;FBXO6;RNF144A;TRIM2;PELI2;TRIM22;NEURL3;DTX4;CBLB;TRIM5;PRKN;MDM2;ZNRF1;RNF150;RNF122;DTX3L;TRIM14;DTX1;TRIM55;MARCHF8;TRIM9;TRIM54;TRIM72;TRIM21;BIRC7;MARCHF2;SH3RF3;UBE2D1;FBXO27;MYLIP;IRF2BPL;PELI3;HECTD1;TRAF3IP2;TRIM56;NHLRC1;TRIM25;NEURL2;RNF13;TRIM35;RNF19A;STUB1;UBE2D3

GAL;ASIP;SHANK1;APLN;NTS

ACP5;HAAO;PHYH;ISCU;LTF;ALOX15;NFU1;FDX1;FECH;CDO1;FTL;FTH1;RIOX1;MELTF;EGLN3;ISCA2;ALKBH2

MYD88;TMBIM1;BDNF;FEM1B;DAB2IP;RIPK1;PIDD1

S100A9;S100A8;TRPV4;DLGAP5;MTUS2;BIRC5;REEP1;FMN1;NUSAP1;KIF2C;LRRK2;KIF21B;GTSE1;PRC1;TPX2;PLK1;KIF14;KIF23;KIF18B;KIF20A;STARD9;PSRC1;FHDC1;FAM83D;RAE1;CENPE;RACGAP1;RPS3;CAMSAP2;KIF22;KIF7;CKAP5;KIF18A;GAS2L3;SKA2;DNM2;REEP4;KIF3C;FNTA;KIFC2;FBXW11;KIF1C;HOOK3;SPAG5;EML2;LRPPRC;TUBGCP3;MARK4;EML1;MAP1S;RGS14;TUBGCP4;OPA1;RP1;HAUS6;MAP7D3;KIF20B;MACF1;NUMA1;HAUS7;PRNP;ZNF207;EML3;DNM1L;SVBP;CCDC66;KRIT1;IRAG2;CCDC170;ARHGEF2

CXCL10;CXCL11;GDF15;CCL8;BMP2;INHBA;IFNB1;CCL22;BMP7;BMP6;IFNA2;CXCL16;CSF1;IL15;CLCF1;CCL7;TGFB2;CD70;BMP8A;IFNA8;CCL1;IFNA14;IL7;IFNA1;WNT1;IFNW1;IL6;KITLG;FGF2;CXCL9

CX3CR1;PDPN;CXCR3;ITGA4;CXCR2;ITGAV;CCR2;PLP2;HMGB1;ACKR2;ACKR3

UCHL1;MAGI2;AKAP5;ARRDC3;SAV1

PLAAT3;PLAAT4;DGAT2;PLA2G4C;SOAT1;SPHK1;ZDHHC9;ZDHHC1;PLAAT1;NAA10;MCAT;LPCAT3;ZDHHC12;GTF2B;NAA16;PLA2G15;ZDHHC8;MOGAT1

ISG20;PNPT1;APLF;NOCT;REV3L;RAD9A;ERI1

FBP1;SMAD3;PPARG;ZFPM2;PARP9;STAT1;EGR2;HSPB1;JUP;TACC2;CDK
N2A;AR;BBS5;NFATC1;DTX3L;GABARAPL1;HES1;BBS2;DDRGK1;CRY1;JU
N;HEY2;TAF4B;MDFIC;STAT3;NFKBIA;TTC8;PROX1;DMAP1;MAD2L2;KLF4
;TBX6;PPARGC1A;ANXA4;RBBP8;SMARCD3;LDB2;HES6;GATA1;IFI27;REL
A

SMAD3;SH3RF2;JUP;KCNN4;CD22;JAK3;MAGI2;PDLIM4;LGALS3;SKAP1;K
IF3A;EIF2AK3;ANK1;SPTBN4;SFI1;CRY2;FBXL2;CRY1;BAD;STX17;SYTL2;
MVP;STAT3;CEACAM1;DZIP3;KIFAP3;CDKN1B

CMPK2;AK8;AK3;NME4;DTYMK;AK5

TLR7;TUT4;HNRNPA1;RBM4;ZC3H7A;DHX9;HNRNPA2B1;TUT7;RC3H1;PU
M2;AGO3;PUM1;ZNF346;RBM10;DDX21;MATR3

HSPB1;PKIA;PRKAG2;CDKN2A;INKA1;PRKAR1B;PKIG;CDKN2B;CDKN1A;C
DKN1B;PRKAR1A;HEXIM1

EDNRA;PLCB4;PLCG2;PLCB3

CX3CR1;CXCR3;CMKLR1;CXCR2;CCR2;GPR35;ACKR2;ACKR3

H1-
5;TOP2A;TAL1;MEF2A;NCOR2;SFPQ;HDAC1;TOP2B;SATB2;HDAC3;MTA1;
MTA2;PRDM1;NIPBL;MEF2C;YWHAB;MIER2;RBBP4;SMG5;BCOR;GMNN;K
CTD21;HSP90AB1;SP2;HDAC2;MIER1;SUDS3;IKZF3;CAMTA2;SPI1;AKAP8;
CCND1

CMPK2;AK8;AK3;MAGI3;GUK1;DLG2

ADCY8;FOXO1;SH3RF2;JUP;KCNN4;CD22;JAK3;PDLIM4;MASTL;AKAP5;L
GALS3;SKAP1;KIF3A;EIF2AK3;ANK1;MFHAS1;FBXL2;BAD;SPHK1;STX17;M
VP;STAT3;CEACAM1;KIFAP3;CDKN1B;DAB2IP;PTPA;PPP1R15A;PPP1R16A;
PPME1;SMTNL1;PHACTR4;TRAF3;MET;PPP1R3F;MYO16

OTULINL;USP7;EIF3H;EIF3F;USP9X;SENP5;YOD1

CLC;MGLL;PLA2G4C;ABHD4;PNPLA7;PLA2G6;ENPP2;LYPLA2;GDE1

HS3ST2;HS3ST3A1;SULT1C2;CHST7;HS6ST1;CHST13;UST;CHST12;CHST1
4;HS2ST1;TPST1

CST7;SPOCK1;CSTA;CST3;BIRC5;RPS6KA3;PTTG1;GAS6

DHX9;YTHDC2;TCERG1;BRCA1;NEDD4;BIN1;PPIB;SCAF8;CCNT2;ANP32B

MGLL;PNPLA3;LIPA;LPL;ABHD6;LIPG;ABHD5;PNPLA8;DDHD1;SMPD3;PLA
2G15;PLA1A;PLBD2;PITPNM3;PNPLA4;ABHD2

UHRF1;SPIN3;KDM7A;SPIN4;ZCWPW2;MBTD1;SUZ12;PHF2;NCAPG2;ATRX
;SPIN1;MTF2;KDM5A;ING5;L3MBTL1;KDM4A;GLYR1;MORC3;NCAPD3;PH
F8;BPTF;ZMYND8;KAT8;CHD8;SPIN2B;MSH6;CXXC1;SPIN2A;CBX2;CBX5;Z
MYND11;RBBP5;MSL3

GBP1;OAS2;GEM;OAS1;MX2;OAS3;GBP4;ATL1;GBP3;RAB38;HSPA4L;RAB13;GBP2;SYN3;NUAK2;TUBA8;RRAS2;ABCG1;RHOB;GBP5;PRPS1;MRAS;ABCB1;RHOC;TUBB3;PRKACB;RHEBL1;KCNJ1;PRKAG2;RRAS;PFKP;TUBA4A;ABCB6;TUBE1;TGM2;DYRK3;RAC2;PIM1;DIRAS1;MAP4K3;MFHAS1;ARL4A;GBP7;DAPK2;RRAGD;HSPA2;RND3;PRAG1;TUBA1A;TUBG2;DIRAS2;P2RX4;GUK1;ABCD4;RHOBTB2;SPHK1;GALK1;P2RX7;RRAD;ABCA1;DAPK3;RAB42;RAB8A;ARL4D;LMTK2;NEK11;ABCB8;SEPTIN6;SIK3;PNPLA8;ARL2;CDKL5;MYH10;ARL5B;SIK1;ARL8A;RASEF;RHEB;SEPTIN4;SYN2;SYN1;RAP2B;RALA;STK38;ABCD1;PTPA;RND1;LATS2;PINK1;INSR;PDK2;MARK1;ARL6;ARL1;PMVK;TUBG1;RRAGA;GCH1;NRAS;RPS6KA4;PFKL;AKT3;GTPBP10;CARS2;MSH2;TUBB1;ATP1A3;RAP1B;RAB11B

KIF11;KIF2C;KIF21B;KIF14;KIF4A;KIF23;KIF18B;KIF20A;STARD9;DNAH17;CENPE;KIF22;KIF7;KIF18A;KLC1;KIF3C;KIFC2;KIF1C;KIF20B;MYH9;DNAH8;KIF24

SMAD3;THRB;RORA;AR;PER1;USP11;ZBTB16;ESR1;THAP7;PER3;NR1D1;RELA

UHRF1;SPIN3;KDM7A;SPIN4;ZCWPW2;MBTD1;SUZ12;PHF2;NCAPG2;ATRX;SPIN1;MTF2;KDM5A;ING5;L3MBTL1;KDM4A;GLYR1;TAF1;MORC3;NCAPD3;PHF8;TAF7;BPTF;ZMYND8;KAT8;CHD8;SPIN2B;MSH6;CXXC1;SPIN2A;CBX2;CBX5;ZMYND11;RBBP5;MSL3

SMAD3;IL1RN;CCRL2;TNFSF9;CSF1;TNFSF18;CLCF1;TGFB2;IFNA8;IL12RB1;TRIP6;TGFBR3;IFNA14;ERAP1;LIFR;IFNW1;VEGFB;IL6;PXDN;PRLR;PGF;VEGFC;KITLG;FKBP1A

KDM3A;KDM7A;KDM2A;HR;PHF2;KDM5C;KDM5A;KDM4A;JMJD6;PHF8;KDM5D;KDM4D;FBXL19;KDM3B;KDM1A;KDM2B;KDM4B;RIOX2

FGD5;PREX2;PREX1;RCBTB2;RASGRP2;DOCK10;RASGRP4;DOCK2;RAB3IL1;ARHGEF6;VAV3;ELMO1;VAV1;FGD3;RIN3;FGD4;DOCK4;ADRB1;IQSEC2;RIC8B;DOCK1;ABR;DENND2B;HPS1;RAP1A;GAPVD1;DENND4B;ALS2;RIC1;ANKRD27;DOCK11;NUCB2;TIAM1;DENND1A;ARHGEF16;MYCBP2;RAPGEF6;RCC1;DIS3;MADD;RAPGEF4;DENND1C;EIF2B3;ARHGEF2;CYTH3;ECT2;VAV2;RASGEF1A;RAPGEF1;ARFGEF1;RALGPS1;FGD2;DOCK9;RAB3GAP1;EIF2B5;LAMTOR1;RGL1;EPS8L1;RAB3GAP2;DENND6B;ARHGEF11;SERGEF;DENND1B;DENND10;LAMTOR4;RGP1

ACOT4;ACOT2;ACOT13;ACAA2;NUDT7;ACOT1;ACOT9

DUSP5;DUSP1;DUSP2;DUSP18;DUSP15;DUSP19

RNASE2;RNASE1;RNASE6;RNASE3;FEN1;SND1;DFFB;MAP1S;N4BP2;ENPP1;NME1;EXO1;DIS3;OGG1;RNASET2;SAMHD1;ISG20L2;RNASE4;DNA2;XRN2;REXO1;DIS3L2;DDX1;ANG

AREG;ERBB4;PDGFC;PDGFRA;TGFA;IL1R1;PLSCR1;EGF;FGF9;ITGB3;IL12RB1;TRIP6;NCSTN;FAM83B;PDGFD;PDGFB;LINGO1;EREG;ERAP1;PDGFA;VEGFB;IL6;PDGFRB;PGF;VEGFC;FGF2

ACOT4;ACOT2;ACOT13;ACAA2;ACOT1;ACOT9;PLA2G6;ABCD1

TRPV4;TRPM2;SLC46A1;PKD1L3;PKD2;TRPC6;PIEZO2;DMAC2L;CALHM2;PIEZO1;PKD2L1;UNC80;CALHM6;TMEM165;SLC12A2;TRPM4

STAP1;LAT;MAGI2;SORBS1;SHANK3;TRADD;SAV1;IRS1;DLG5;LDLRAP1;SHANK1;SHC1

PPARG;NRAP;MAGI1;PALLD;PDLIM4;DAG1;SYNPO2

CHRNE;CHRM3;CHRM4

AKR1B1;AKR1E2;DHRS13

LPCAT1;PNPLA3;ABHD4;AGPAT4;ABHD5

HERC5;RNF128;UBE2L6;TRIM47;UBA7;SH3RF2;BIRC3;FBXO6;RNF144A;TRIM2;PELI2;TRAF5;TRIM22;NEURL3;DTX4;CBLB;TRIM5;RNF213;TRAF1;PML;PRKN;RNF168;MDM2;ZNRF1;RNF150;RNF122;DTX3L;TRIM14;DTX1;TRIM55;MARCHF8;TRIM9;TRIM54;TRIM72;DDB2;TRIM21;UBE2R2;BIRC7;MARCHF2;SH3RF3;UBE2D1;UBE2D4;FBXO27;MYLIP;IRF2BPL;PELI3;HECTD1;TRAF3IP2;TRIM56;DZIP3;RFPL2;UBE2J1;FBXO4;RBCK1;NHLRC1;TRIM25;NEURL2;RNF13;TRIM35;RNF19A;TSPAN17;RMND5B;STUB1;UBE2D3;FBXO7

KCNQ3;KCNC3;KCNQ1;KCNE3;KCNA1;KCNG1;KCNE5

TBXAS1;HACD4;CYP2S1;CA13;CA11;ENO3;ALAD;ECHDC3;HACD1;ENO1;HACD2;IREB2

SIGLEC10;TRPV4;SH3PXD2B;PAG1;ABL1;SIT1;INPPL1;RACK1;KHDRBS1;CRK;BLNK;LAT2

PLTP;APOL2;THBS1;LPL;CD36;MSR1;LDLR

FBP1;SMAD3;PPARG;ZFPM2;TLE1;STAT1;EGR2;PRDM5;HSPB1;TLE5;AR;BBS5;BCL2;HDAC9;NFATC1;HES1;BBS2;DDIT3;USF2;DDRGK1;JUN;HEY2;TWIST1;DAPK3;STAT3;NBN;TTC8;DMAP1;MAD2L2;KLF4;TBX6;ZNF503;RBBP8;SIK1;LDB2;HES6;GATA1;YAP1;IFI27;RELA;CREM;YWHAZ;ZNF703;USF1;TP53BP1;FOS;NKX3-1;EIF4E;TLE3;NFE2L2;SIRT1;PITX1;PSMD9;PPARA;GFI1B;CTNNB1;GSC

CCL8;NES;CCRL2;CCL22;CCL7;CCL1

SLC26A1;SLC12A5;SLC26A2;SLC26A3;CLCN6

GBP1;KDR;PTGES3L;NPAS2;PACRG

RNASE2;RNASE1;RNASE6;RNASE3;SND1;CNOT6;ERN1;CPSF3;AGO3;SMG6;EXOSC3;RNASET2;SAMHD1;RNASE4;EXOSC5;ELAC2;DIS3L2;YBEY;ANG

SMAD3;ATF3;FOXO1;ZNF395;EGR2;SOX4;FOXO3;NR4A3;FOXF1;MAFF;IRF1;MDM2;GRHL1;AR;RUNX1;HOXB2;PLSCR1;CASZ1;ATF5;NFATC1;SOX9;ZNF521;NR4A2;NFIB;NFKB2;ZBTB16;REL;CREB3;ESR1;LEF1;HOXB7;DDIT3;ZNF43;FOSL2;JUN;NFIA;KLF10;MEIS1;CREBRF;FOXA1;TFDP2;FOXK2;STAT3;ONECUT2;HOXA5;KLF6;KLF4;ALX1;MITF;FOXC1;MEIS2;SMARCD3;HNF4G;GATA1;RELA;RFX5;NFKB1;LITAF;NFIC;NR4A1

VNN2;VNN1;AMDHD2;HDAC9;TGM2;NGLY1;NIT1;ASAH1;HINT3;AGA;NTAQ1;ASAH2B;ACER2

KCNK6;KCNK13

CX3CR1;IL7R;CXCR3;CMKLR1;IL1RAP;IFNGR2;IL17RA;CSF3R;CCR2;CD74;IL22RA2;IL18R1;GFRA2;IL10RA;ACKR2;IL1RL2;CD4;IL13RA1;CSF2RB;IL4R

CBR1;TP53I3;AIFM2

UCHL1;SMAD3;UBE2L6;HSPB1;CXCR4;PRKN;RNF168;MDM2;DNAJB2;MVB12B;RBCK1;UBXN2A;FBXO7;NSFL1C;SHARPIN;RAD23A;USP25;PLAA;WDR48;AUP1;MVB12A;UBXN11;ADRM1;OTUB1;NBR1;IKBKG

SMAD3;RGCC;ACTA2;STAP1;CAVIN3;IL15RA;CSPG4;CEP68;SOCS1;HSPB1;TRIM22;TEX14;TFRC;DSP;SPDYA;FOXO3;NR4A3;PKIA;LAT;DUSP5;RHOB;TRIM5;RHOC;PRKN;PRKAG2;CDKN2A;NBEAL2;DUSP1;CASP9;SKAP1;WARS1;HDAC9;TUBA4A;INKA1;SOX9;CALM3;PRKAR1B;CDKN2B;DUSP2;EPHA4;PLEK;PARD6A;RAC2;PTPN1;TRIM14;GSTP1;DIRAS1;TRIB3;CASS4;UNC5C;PRKCB;DNM1;ESR1;TRIB2;FAM83B;TBL2;PDE3B;RND3;MAP2K3;ITGB2;DUSP19;BAD;PLCG1;RHOBTB2;CDKN1A;TRIM21;NBEA;NEDD9;FGR;IRS1;PEBP1;MVP;PPP1CB;USP37;STAT3;CEACAM1;RAD9A;ITGB1;PDGFRB;STX1B;CIR1;GRK5;GATA6;ACSL3;GPRC5B;MAD2L2;RNF13;FERMT2;CDKN1B;PGAM1;RELB;FRMD5;CD24;PRR7;CYLD;LYST;MAPK7;FBXO7;ANKRA2;LRBA;SIK1;SHC2;PRDX3;PIN1;TPCN2;PRKAR1A;ZFP36;RICTOR;TNIP2;SYN1;STK38;CAMK2N2;DAB2IP;PTPRR;RELA;RND1;PINK1;DUSP3;YWHAZ;PPP1R15A;SHC1;CNTLN;TRIM6;NEFH;IRGM;GCH1;USF1;CD247;PTN;GSKIP;TRIM68;RYR2;PRKD2;ADIPOR1;MSH2;ILK

AFAP1L1;DPYSL3;ENAH;SGIP1;CBLB;SYNJ2;AKAP5;PRKN;CASP9;SKAP1;PLSCR1;AFAP1;SHANK3

CACNA1D;TRPV4;TRPM2;ANXA6;PKD1L3;ATP2A3;TRPV1;PKD2;ATP2A2;ITPR1;ATP2C1;PKD2L1;RYR1;TMEM165

MLPH;CXCR4;RAB3A;CALD1;SPATA6;RAB6B;RAB3C;RAB8A;LMTK2;RAB11A;DMD;RALA;STXBP5L;RAB11B;MYRIP

SMAD3;USF2;TWIST1;USF1;SIRT1;PSMD9;LMO2

TRPV4;TRPM2;TMC8;ANXA6;PKD1L3;GRIN2D;TRPV1;PKD2;TRPC6;PIEZO2;CALHM2;PIEZO1;PKD2L1;UNC80;CALHM6;GRIK1

ANO5;GABRB2;TTYH2;APOL1;SLC1A4;CLIC4;GABRG2;GABRR2;GLRA1;PACC1;FXYD1;GLRB

ERAP2;LAP3;ENPEP;CTSH;XPNPEP2;MMP17;ERAP1;RNPEPL1;TRHDE;ANPEP

CACNA1D;TRPV4;TRPM2;ANXA6;HVCN1;PKD1L3;TMEM63C;TRPV1;PKD2;TRPC6;PIEZO2;CALHM2;PIEZO1;PKD2L1;UNC80;CALHM6;KCNA1;RYR1;TMEM38A;KCNG1

CXCL10;CXCL11;CCL8;CCL22;CXCL16;CCL7;CCL1

ANO5;TTYH2;TMEM63A

AREG;ERBB4;TGFA;PLSCR1;EGF;FAM83B;LINGO1;EREG

OPTN;PARP10;RNF168

ISG20;PNPT1;XRN1;NOCT;ERI1

SCN1B;SCN9A;PKD2

ACP5;PLPP3;DUSP5;PLPP4;ACP2;DUSP18;DUSP15;NOCT;PLPP1;NT5M;PXYLP1;PPP1CB;PLPP6;ALPL;INPP5K;PLPPR4;CILP2;PLPP5;DUSP3;LPIN1;HDDC2;DUSP4;DUSP12

HAAO;IDO1;ALOX15;CDO1;PIR;HGD;ALOX15B;IDO2;ADI1;ALOXE3

ELAVL4;CPEB3;TARDBP;RPL5;RBM4;RPS7;ILF3;CARHSP1;RBMS2;HNRNPA2B1;CELF2;RC3H1;DDX5;PABPC1;CCT5;LARP4;IGF2BP1;ZFP36L1;PUM2;RBMS1;PUM1;UTP23;LARP4B;HNRNPC;DAZAP1;PABPC4;RSL1D1;SERBP1;ANGEL2;FXR1;GEMIN5;HNRNPU;RNF40;CELF1;SECISBP2;LARP1;HNRNPA0

TARBP1;TRDMT1;TRMT13;METTL2B;TRMT44;THUMPD3;TRMT12

SMAD3;PPARG;ATF3;NR1H3;EGR2;PRDM5;ERBB4;SOX4;FOXO3;HMGA2;THRA;IRF1;KLF17;GRHL1;AR;RUNX1;PER1;PAX8;ATF5;ARID5B;MESP1;CRY2;DDIT3;JUN;MACROH2A2;FOXA1;FOXK2;SOX6;SOX13;SOX5;STAT3;PROX1;GATA6;KLF4;TBL1X;FOXC1;PER3;NR1D1;GATA1;BASP1;CALCOCO1;YAP1;MAFK;RELA;NFKB1

TM7SF2;BLVRB;SRD5A3;BLVRA;DECR1;BDH2

PPARG;MAGI1;PDLIM4;DAG1;SYNPO2;CSRP2;RELA;NFKB1

SLC1A3;SLC1A4;SLC25A18;SLC1A7;SLC25A22;SLC1A2

FLT1;FGFR1;TIE1;ERBB4;ROR1;KDR;PDGFRA;RET;EPHA4;ACVR2A;LMTK2;PDGFRB

NEK10;TGFBR1;EGFR;CDK1;LRRK2;CAMK1D;PLK1;STK39;NEK2;DAPK1;BMP2K;SYK;PAK1;CAMK1;PRKCZ;MKNK1;DMPK;HIPK2;PRKCA;PBK;MYLK4;CDK14;MAST3;MAP3K14;PRKDC;STK32C;TTK;RPS6KA3;CIT;AAK1;PKMYT1;TGFBR2;ULK2;MINK1;MAP2K1;PTK2B;WNK1;AURKB;TOP1;MAP3K7;MARK2;STK38L;MAP3K13;TTN;MAP3K1;IKBKB;STK4;GRK2;MELK;AURKA;STK17B;ERN1;CHEK1;CHEK2;CDK8;TAOK3;LATS1;MAP3K2;CDK13;RIOK1;MARK4;CSNK2A2;CHUK;MAPK14;NRBP2;CSNK2A1;PAK4;CSNK1G3;IRAK3;PRKAA1;STK36;MYLK3;PAK2;UHMK1;TAF1;MAP3K6;MTOR;WNK3;CDK12;HIPK1;MARK3;CDK6;PDIK1L;MAPK1;TNIK;MAPK9;SNRK;GSK3B;MAP4K2;MAPKAPK5;HIPK3;TAOK1;CDK19;GAK;NUAK1;STK17A;CLK4;VRK1;CSNK1D;PRKCI;SMG1;BRD4;VRK2;CSNK1A1

PALD1;PTPN14;EYA2;CDKN3;PTPRO;PTPRJ;PTPN13;CDC25C;PTPN22;EYA3;PTPN9;DUSP22;PTPN11;CDC25A;HACD2;PTPDC1;PTPRE;PTPN18;PTPN12;PTPN2;DUSP23;PTPN7;CDC14A;PTPRD;UBASH3B;PTEN

EGFR;PLEC;ANLN;CDH6;PDLIM1;NIBAN2;TMPO;PTPRO;CTNND1;TES;CBL
;SWAP70;PTPRJ;ITGA6;ARHGAP18;RPL6;EPS15;STXBP6;CAPZB;CLINT1;CI
P2A;DIAPH3;EEF1G;FASN;BAIAP2;SEPTIN2;TBC1D10A;BZW1;CDH8;MARK
2;SND1;CKAP5;PTPN11;EPHA2;RPL24;PLCB3;PICALM;CDH26;EIF2S3;PDLI
M5;MB21D2;PFN1;USP8;CAPG;EHD1;HDLBP;CDH2;GAPVD1;RPL15;CDK5
R1;EIF4G2;HSPA5;RPL34;GIPC1;RPL29;RPS26;TJP1;ENO1;RACK1;ZC3H1
5;NOTCH3;YWHAB;RPL23A;CCT8;VASP;PAK4;ARHGEF16;ARHGAP1;MYH9;
PCMT1;HNRNPK;EEF2;PACSIN2;EIF3E;PAK2;MACF1;SNX2;ATXN2L;HSP90
AB1;DBNL;CRKL;ERC1;EXOC3;GIGYF2;RAB10;HCFC1;PAICS;GOLGA3;KT
N1;EIF2A;PKP4;DHX29;GCN1;RSL1D1;SEPTIN7;TJP2;RUVBL1;CSNK1D;RA
RS1;ARGLU1;DDX3X;DOCK9;SH3GL1;SERBP1;RPL14;CDH20;DNAJB1;PA
RVA;AHSA1;CCNB2;BZW2;YKT6;PPP1R13L;RANBP1;GPRC5A;EPS15L1;CA
PZA1;KRT18;OLA1;CDH5;EPS8L1

PNPLA3;LPL;LIPG;ABHD5;PLA2G15;PNPLA4

PDLIM1;STXBP6;BAIAP2;PDLIM5;PAK4;RAB10

PRUNE2;TRPM2;NUDT3;ENTPD1;DCP2;DHX9;BNIP2;NUDT15;ENPP1;ENTP
D7;NUDT16L1;DHX29;ENTPD4;NUDT14;LHPP;DDX3X;PHOSPHO1

EGR2;MDM2

FLT1;FGFR1;TIE1;ERBB4;ROR1;KDR;PDGFRA;RET;NRP2;EPHA4;IGF2R

MME;SMAD3;PPARG;TNFSF10;ZFPM2;MT2A;IFIH1;APOBEC3G;ZNF331;D
HX58;TRIM2;ADAMDEC1;ERAP2;APOBEC3F;MMP9;PAM;CBLB;ADA;RASG
RP1;LARGE1;PRKN;MDM2;SMPD1;ISCU;SMPDL3A;MMP12;PLSCR1;LTF;Z
NF90;MTARC1;ARG2;MTARC2;CA2;ENPEP;QPCT;PTPN1;ECE1;PIM1;ATOX
1;SHANK3;ZNF93;TRIM54;ALOX15;MGAT1;TIMP2;NFU1;ZNF184;P2RX4;S
100A13;FDX1;HMGCL;PHEX;ERAP1;MPPED2;SCO2;POMGNT1;TDP2;MT1E
;MOXD1;FTL;TMEM163;FTH1;KLF4;HPF1;CZIB;GLRA1;ABL2;SOD2;RIOX1
;MELTF;CYLD;TRHDE;QPCTL;CDA;PAPPA2;PDE2A;ADPRM;B4GALT1;ISCA2
;GPC1;ZFPL1;LANCL1;ANPEP;GTF2B;RAG1;ENPP2;LITAF;PLA2G15;ZCCH
C4;WDFY1;MNAT1;ADH5;IKZF5;GCH1;CALR;GYG1;DPYS;RNF8;BHMT2

SLC40A1;SLC11A1;SLC39A8;ATP7A;TMEM165;SLC11A2;SLC25A37

S100A9;S100A8;TRPV4;DLGAP5;MTUS2;BIRC5;REEP1;FMN1;TRIM36;NUS
AP1;KIF2C;LRRK2;KIF21B;GTSE1;PRC1;TPX2;PLK1;KIF14;KIF23;KIF18B;
KIF20A;PAK1;STARD9;PSRC1;FHDC1;FAM83D;RAE1;CENPE;RACGAP1;WA
SHC1;RPS3;KIF22;KIF7;CKAP5;KIF18A;GAS2L3;SKA2;DNM2;REEP4;KIF3C
;FNTA;STMN1;AGBL4;KIFC2;KIF1C;HOOK3;CDK5R1;SPAG5;EML2;LRPPR
C;TUBGCP3;MARK4;EML1;TPPP2;CCT5;MAP1S;RGS14;TUBGCP4;OPA1;C
ENPJ;RP1;HAUS6;MAP7D3;KIF20B;SETD2;NUMA1;BRCA1;PRNP;ZNF207;E
ML3;DNM1L;TBCEL;GABARAP;TTLL11;SVBP;CCDC66;SNCA;KRIT1;IRAG2;
CCDC170;ARHGEF2;RABGAP1;CLIP2;ARL4C;GAPDH;PDCD5;CETN3;CHP1;
TAOK1;NME8;CDK5RAP2;KIF3B;OFD1;TUBGCP2;SMC3;MAP1LC3B2;TRA
PPC14;DDX3X;TBCD;CEP295NL

TIMP3;SERPING1;PZP;SERPINE1;BST2;RARRES1;SERPINI1;CSTB;LXN;TFP
I;SERPINE2;LTF;A2M;SPINK5;TIMP2;CAST;BIRC7;TIMP4;TFPI2

MME;SMAD3;PPARG;TNFSF10;ZFPM2;MT2A;IFIH1;APOBEC3G;ZNF331;DHX58;TRIM2;ADAMDEC1;ERAP2;APOBEC3F;MMP9;PAM;CBLB;ADA;RASGRP1;PRKN;MDM2;SMPD1;SMPDL3A;MMP12;PLSCR1;ZNF90;CA2;ENPEP;QPCT;PTPN1;ECE1;SHANK3;ZNF93;TRIM54;TIMP2;ZNF184;P2RX4;S100A13;PHEX;ERAP1;MT1E;TMEM163;KLF4;HPF1;CZIB;GLRA1;CYLD;TRHDE;QPCTL;CDA;PAPPA2;PDE2A;ZFPL1;LANCL1;ANPEP;GTF2B;RAG1;ENPP2;LITAF;PLA2G15;ZCCHC4;WDFY1;MNAT1;ADH5;IKZF5;GCH1;CALR;DPYS;RNF8;BHMT2;TET1;PAPPA;SIRT5;TIMM8B;THAP11;MYRIP

EDNRA;PLCB4;PLCG2;PLCB3

DUSP5;DUSP1;NFATC1;DUSP2;DIRAS1;MAPK7;GCH1;DUSP4;SIRT1;MAPKAPK3;NBR1

SMAD3;RGCC;ACTA2;STAP1;IL15RA;CSPG4;CEP68;SOCS1;FAS;HSPB1;TRIM22;TEX14;TFRC;BCAR3;SPDYA;FOXO3;NR4A3;LAT;TTC28;RHOB;PIK3IP1;TRIM5;RHOC;PRKN;PRKAG2;CDKN2A;NBEAL2;PER1;CASP9;SKAP1;WARS1;WWC1;TUBA4A;INKA1;ATF5;CALM3;PDE8A;CDKN2B;RAC2;PTPN1;TRIM14;BTG1;TRIP6;TRIB3;UNC5C;DNM1;ESR1;FAM83B;TBL2;RND3;MAP2K3;GADD45A;ITGB2;BAD;PLCG1;RHOBTB2;CDKN1A;TRIM21;NBEA;TRADD;FGR;DLG2;PEBP1;MVP;PPP1CB;USP37;STAT3;CEACAM1;RAD9A;PDGFRB;STX1B;CIR1;GATA6;ACSL3;GPRC5B;FERMT2;CDKN1B;PGAM1;RELB;FRMD5;CD24;CARD16;CYLD;STUB1;LYST;FBXO7;ANKRA2;LRBA;SIK1;SHC2;PRDX3;TPCN2;ZFP36;RICTOR;TNIP2;SYN1;RAD23A;CAMK2N2;DAB2IP;PTPRR;RELA;RND1;DUSP3;YWHAZ;PPP1R15A;MIDN;SHC1;CNTLN;TRIM6;NEFH;IRGM

PTPRN2;PTPRG;PTPRF;PTPRB;PTPRA;PTPRR

PTPRN2;PTPRG;PTPRF;PTPRB;PTPRA;PTPRR

CX3CR1;CXCR3;CXCR2;CCR2

PDPN;SLC43A2;SLC36A1;SLC7A5;SLC7A6;SLC7A7;SLC7A1;SLC6A6;SLC38A7;SLC38A10

RORA;PER1;USP11;ZBTB16;ESR1;THAP7;PER3;NR1D1

GABRA3;GABRE;GABRB3

CD109;THBS1;LTBP1;CD36;HYAL2;TGFBR3;TSKU;LRRC32

KCNQ3;CACNA2D3;CACNA1D;KCNC3;HVCN1;ITGAV;KCNQ1;GRIN2D;CATSPER1;PKD2;PTK2B

IL1RN;LRPAP1;DKK1;PXDN

GALNS;ARSA;SULF2;GNS;ARSK;ARSD;ARSF

CDT1;FANCI;HMGB1;RAD51;HSP90AB1;PTGES3;FANCD2;SMG6;POLG2;NAT10;SMARCA4

APOE;MESD;LRPAP1;DKK1;LDLRAP1;ANKRA2;LANCL1;SYT1

ATP2A3;NLRP1;CASP1;APAF1;PYCARD;NLRP12;RACK1;NOD1

SMAD3;OASL;STAT1;NR4A3;JUP;TACC2;NR4A2;TRIP6;PRKCB;ESR1;LEF1;
CRY1;WBP2;PROX1;PPARGC1A;STAT5B;SMARCD3;CCDC62;DNAAF4;GTF
2B;TRERF1;NR4A1;TAF10;CALR;NCOA2;TRIM68;HMGA1;HMGN3;MED16

SYNDIG1;HIP1;DLG4
SYT6;S100A9;SPOCK1;S100A10;SLIT3;S100A8;S100B;TESC;CLEC4A;SLC
8A1;GALNT3;ADAM8;S100Z;TRPM2;S100A12;CDH6;CLEC3B;TNNC1;S100
A4;SYT4;ANXA6;RASGRP2;AOAH;TPT1;TKT;S100A5;SPARC;PKD2;SLC25A2
4;CDH8;LRP1;ATP2A2;TPD52;TTN;CANX;BNIP2;RCN3;CDH26;TCHH;THBS
4;MELK;HSP90B1;CDH2;CDK5R1;ACER3;ITPR1;HSPA5;EML1;MCTP1;ESY
T1;PKD2L1;LTBP4;HPGDS;ENPP1;NUCB2;CCDC47;PPP3CA;PPP3R1;PRNP;
ANXA5;RYR1;PPP3CB;PLA2G4A;IDS;RASGRP3;CALML4;MEX3B;SNCA;SLC
25A13
SLC11A1;ATP2A3;ITPR1;KCNA1;CLDN16;TMEM165;SLC11A2;SLC12A2;C
NNM2;ATP2B1;ZDHHC13
GCNT4;LARGE1;B4GAT1;CSGALNACT1;B3GLCT;RFNG;CHPF;CHSY1;EXT
1;HEXB;B3GAT3;B3GNT2;POMGNT1;POGLUT3;MGAT4C;POGLUT1;POM
GNT2;CHSY3;B4GALT1;B3GALT1

ISG20;PNPT1;NOCT;ERI1
GCNT4;FUT4;LARGE1;B4GAT1;B4GALT5;DPY19L2;CSGALNACT1;B3GLC
T;RFNG;CHPF;CHSY1;EXT1;B4GALT2;HEXB;B3GAT3;B3GNT2;POMGNT1;
POGLUT3;MGAT4C;POGLUT1;POMGNT2;CHSY3;GBE1;ALG1;B4GALT1;B
3GALT1;PIGV;B4GALT6;GYG1;TMTC3;EXT2;ALG9;B4GALT3;B4GALT7;TM
TC4;POMT2;LFNG
CACNA2D3;CACNA1D;TRPV4;TRPM2;ANXA6;ITGAV;PKD1L3;CATSPER1;TR
PV1;PKD2;TRPC6

CX3CR1;CXCR3;CXCR2;CCR2

POPDC3;PDE4B;PDE2A;PDE10A;CNGA4;HCN1;PDE4D;RAPGEF2;KCNH1

CXCL10;CXCL11;CCL8;CCRL2;CCL22;CXCL16;CCL7;CCL1

PDE6B;CNP;PDE8A;PDE3B;PDE8B;PDE7A;PDE4B;PDE4C;PDE2A;ADPRM;P
DE10A
CACNA1D;SLC8A1;PLEC

MBD4;RPS3;FEN1;DCLRE1C;EXO1;ERCC4;DNA2
SETD3;KMT5A;SETD2;ATPSCKMT;SETD7;EHMT1;SMYD2;EEF1AKMT2;SET
D4;SETD6

TOP2A;HMGB3;HHEX;HMGB2;HMGB1
ISG20;APLF;REV3L;RAD9A;ERI1;AEN;ENPP2;EXOSC9;EXOSC6;EXOSC4;RE
XO2;ZC3H12A;EXOSC1;WRN;APEX2;EXOSC2;EXOSC8;EXD3;RAD1

THRA;AR;ESR1;DRAP1;THAP7;RELA;BRF2;GTF2B

ITGA3;ATF3;ADCY8;TPM2;BMP6;ABCG1;MLXIPL;EPAS1;BCL2A1;ZHX2;BC
L2;TENM1;NR4A2;RRAGD;DDIT3;USF2;NPAS3;GADD45A;EXT1;SRI;PDGFB
;ATP1B1;BAK1;PDGFA;RALGAPA1;ITGB1;PEF1;NAE1;FBXO7;MCL1;PPP2R
1A;TPM1;BAX;NR4A1;ADRA2C;RRAGA;USF1;TRMT112;SUPT4H1;EXT2;FM
R1;KATNB1;GCA;TENM3;TPM4;QTRT1;ADD2;FZD4;MLX;PAFAH1B3;ZBTB
1;IKBKG

SLA;TXK;SYK;HCK;PTK2B;MELK;TEC;ABL1;CSK;FER;PEAK1
GAL;INHBA;POMC;ASIP;NPPC;PNOC;EDN1;APLN

SLC26A1;SLC39A8;SLC26A2;SLC26A3;SLC4A3

CST7;SLPI;CST3;VSIR;NLRC4;SPINT2
SLC30A3;SLC39A3;SLC39A6;SLC39A13;SLC39A10;SLC39A14;SLC30A2;S
LC39A11
SQOR;STEAP4;KMO;CRY2;DDO;CRY1;MICAL3;MICAL2;CYB5R3

NT5DC2;ACP3;NT5E;NT5C2;NT5DC4
DPP4;KLK4;MMP9;PIK3IP1;PCSK6;F3;RHBDF2;CTSH;HTRA2;MST1;PRSS2
1;TYSND1;HTRA3;MMP2;KLK10;RHBDF1;FAP;C1S;KLK1

CD244;MARCHF1;CD74;CLEC7A;PILRA;CD4
CPE;ERAP2;LAP3;ENPEP;MMP17;PEPD;ERAP1;RNPEPL1;CPXM1;TRHDE;A
NPEP
ME3;TP53I3;TM7SF2;CBR3;G6PD;DECR1;HSD17B1
DPP4;MMP9;PIK3IP1;PCSK6;F3;RHBDF2;CTSH;HTRA2;MST1;PRSS21;TYS
ND1;MMP2;RHBDF1;FAP;C1S;KLK1

MYO1G;HIP1;ARAP3;NCF1;FCHSD2;DAPP1;MAPKAP1;ADAP2
POLE4;POLQ;POLE;REV1;POLL;POLD1;POLE2;POLE3;POLN;POLA1;POL
M;POLK

APOE;MESD;LRPAP1;DKK1;LDLRAP1;ANKRA2;LANCL1;SYT1

SLC36A1;SLC2A9;SLC2A13;SLC39A8;SLC15A2
KPNB1;IPO5;BRAP;KPNA6;KPNA3;TNPO1;TNPO2;KPNA4;POM121C;NUP1
53;NUP214;RANBP6
POLE4;POLQ;POLE;REV1;POLL;POLD1;POLE2;POLE3;POLN;POLA1;POL
M;POLK
DHX9;WAC;PCF11;CTR9;URI1;SCAF8;ELP4;BRD4;SMYD2;RPRD1B;CCAR2;
RECQL5;HNRNPU;WDR43;ZNF326
STAP1;PITPNM2;CBLB;TOB1;FLT3LG;PTPN1;HYAL2;ALKAL1;ITGAX;ANGP
T1;TRADD;MST1;IRS1;ANGPT4;PCNA

ALOX5AP;GSTM2;GSTM3
HNRNPA1;RBM4;RBM41;CELF2;DDX5;RNPC3;PTBP1;SRSF6;TRA2B;HNRN
PU;U2AF2;CELF1

SLC46A1;COX6B1;SLC9A1;DMAC2L;NDUFA4;UQCRH;UQCR10;COX4I1;COX10;SLC11A2;UQCRQ;NNT;SLC33A1;NDUFA4L2;CLCN4;COX5A;COX8A

PIWIL4;ZC3H12D;ZC3H12C;RCL1;RIDA;PIWIL2;ZC3H12B;AGO2;LACTB2;ZC3H12A;SLFN13;TSNAX;RNASEL

POR;AIFM2;SQOR;STEAP4;KMO;CRY2;DDO;CRY1;MICAL3;GCDH;MICAL2;CYB5R3;ACOX2;ACADVL;CYB5R1;PPOX

SERPINF1;SPOCK1;SLPI;AGT;SERPINA1;SERPINB8

CXCL10;CXCL11

PLEKHF1;PHLDA3;VEPH1

TRPM2;GRIN2D;TRPV1;ITPR1;RYR1;TRPM4

TDRD9;MYO1E;TOR1B;RNF213;ABCB6;ABCA3;TOR3A;ABCB10;SMCHD1;TOR1A;ABCD1

IFIH1;DDX60;DHX58;IFIT5;RBM11;DDX11;ADARB2

CARD9;CARD11;CARD19;CASP1;MAVS;NOD1

CXCL10;RGCC;SOCS1;CCNA1;SPDYA;SH3BP5;PRKAG2;PRKAR1B;SOCS3;DUSP19;FGF13;CDKN1A;ITPRIP;CCNB3;GPRC5B;CDKN1B;MAPK8IP1;PRKAR1A;HEXIM1

ATP2A3;NLRP1;CASP1;APAF1;PYCARD;NLRP12;RACK1;NOD1

PPARG;NR4A2;RARG

GABRA3;GABRE;GABRB3

KDM3A;DDX5;ARID5A;NSD1;PRMT2;KDM5D;RNF6;EP300;SMARCA4;KDM1A;PARK7;FOXH1;SNW1;TMF1;DAXX;ZBTB7A

ATP6V1C2;TCIRG1;ATP6V0E2;ATP6V0C;ATP6V1D;ATP6V0B;ATP6V0A2;ATP6V1E1;ATP2C2

PARD3;PLEKHA4;PLEKHF1;BBS5;PHLDA3;ATG2A;SNX20;ZFYVE26;DAB2IP;WIPI2;KIF16B;RUFY4;WIPI1;WDR45B

CPE;MINDY1;CTSZ;PEPD;SCPEP1;CPXM1;MINDY2;CTSA;CPB1;AGBL1;CPD;AGBL3;CPO;CPB2

CYP1B1;CYP2S1

SCIN;PARD3;PLEKHA4;KCNJ1;SVIL;PHLDA3;FZD7;ALOX15;PLCD1;PLCB1;SYTL2;SNX20;LDLRAP1;GSDME;HIP1R;MARK1;KCNJ2

KLF12;SKI;SKIL;ETS2;GZF1;MXD1;E2F8;FOXP4;TBX18;JDP2;ZNF217;HHE
X;MYC;SNAI1;ZNF124;IRF8;ZNF749;ZNF169;KLF8;ZNF175;ELK3;ZNF785;
TBX3;ZNF764;TFAP2C;ZNF180;ZNF587B;NFATC3;REST;POU6F1;ETV6;ZB
TB26;ZBTB34;PPARD;ZNF148;PRDM1;ERF;TCFL5;TRPS1;ZFHX3;ENO1;ZB
TB39;MYPOP;ZBTB37;ZNF281;MAX;ZGPAT;ZBTB18;BATF3;ZNF3;ZNF133;
ZNF614;SP2;ZNF589;TCF3;ZNF248;ZNF75A;FOXK1

IL1A;LACC1;S100A12;S100A5;ATP7A

ROR1;FZD6;TSPAN12;FZD7

ERAP2;ENPEP;MMP17;ERAP1;RNPEPL1;TRHDE;ANPEP

KCNJ5;KCNJ1;KCNJ15

UCHL1;USP2;USP18;USP11;USP36;USP49;MINDY1;USP51;USP37;CYLD;O
TUD4;USP31;MPND;OTUD1;USP25;MINDY2;JOSD1;USP28;USP44;ZC3H12
A;OTUB1;USP50;BAP1

KMT5B;SUZ12;EED;KMT2C;KMT5A;SETDB1;SETD2;PRMT2;SETD7;EHMT1
;SMYD2;PRMT7

P2RX4;P2RX7;P2RX1;GLRB;P2RX6

CYP27A1;CH25H

PARP9;HPGD;PARP15;ZC3HAV1;HADH;BDH2

SLCO2B1;SLC16A7;SLCO4C1;COX6B1;NDUFA4;UQCRH;UQCR10;SLCO3
A1;COX4I1;COX10

SLC28A3;SLC5A3;SLC5A4;SLC5A6

FBP1;OAS2;OAS1;OAS3;HSPA4L;SYN3;NUAK2;ABCG1;POPDC3;PRPS1;AB
CB1;PRKACB;KCNJ1;PRKAG2;PFKP;ABCB6;MYO6;DYRK3;PIM1;MAP4K3;
DAPK2;HSPA2;PRAG1;P2RX4;GUK1;ABCD4;SPHK1;GALK1;P2RX7;ABCA1;
PDE4B;MPPED2;DAPK3;LMTK2;NEK11;ABCB8;SIK3;PNPLA8;COQ8A;CDK
L5;MYH10;SIK1;PDE2A;SYN2;SYN1;PDE10A;STK38;ABCD1;PTPA;LATS2;PI
NK1;INSR;PDK2;MARK1

UCHL1;USP2;USP18;USP11;USP36;USP49;USP51;USP37;CYLD;OTUD4;US
P31;MPND;OTUD1;USP25;JOSD1;USP28;USP44;ZC3H12A;OTUB1;USP50;
BAP1

HDAC1;HDAC3;MIER2;SIRT6;HDAC2;MIER1;SUDS3;HDAC8;HDAC4

SCIN;PITPNM2;PARD3;MYO1E;MREG;PLEKHF1;ESYT2;SNX9;SNX10;AIDA;
SNX31;RUBCN;SCARB1;ING2;PIGK;HIP1R;WDFY1;ZFYVE16;PIGU;SNX16;S
NX8

SCIN;FBP1;GEM;MT2A;MCTP2;DYSF;S100A16;APOE;SCUBE1;NUAK2;PITP
NM2;RCN1;NT5C3A;SELL;PAM;THBS1;CDH23;THBS3;ANXA11;AGRN;KCNI
P1;RASGRP1;PRKACB;ESYT2;ADGRV1;RET;ASTN2;PLA2G4C;MMP12;S100
P;PLSCR1;NOL3;CALM3;NCS1;PAPOLG;TGM2;MAN1C1;DYRK3;LPL;PRTF
DC1;ARSA;PLSCR2;ANXA10;CSGALNACT1;NRXN1;MAN1B1;THTPA;PLA2G
2D;S100A13;SPHK1;NUCB1;HMGCL;ADPRS;SRI;NUDT18;FBN1;RCN2;CIB
2;OTOF;FOXK2;CALU;SYT11;TDP2;SYT12;MT1E;ADPRH;S100A1;PLSCR4;I
TSN1;SIK3;PCLO;SDF4

AKAP7;AKAP5

SEMA4C;SEMA4B;SEMA4D;SH3BP1;SEMA6B
DHX9;WAC;PCF11;CTR9;SPTY2D1;URI1;SCAF8;ELP4;SMYD2;RPRD1B;CCAR2;RECQL5;HNRNPU;WDR43;ZNF326
IFIT5;CYFIP2

ENPP4;NUDT17;NUDT4;FHIT;CILP2;NUDT6

TGFBR1;ENG;TGFBR2;LTBP4

PLTP;THBS1;CD36;MSR1;LDLR

ELAVL4;CPEB3;ILF3;ZFP36L1

SORL1;GPR37;OPRD1;OPRM1

SLC24A2;SLC9A9;SLC9C2;SLC9A8;SLC9A3;SLC9A2
EIF4B;NPM1;ABCE1;PTCD3;NME1;EIF1;DHX29;DDX3X;MTIF2;EIF4H;LARP1
JAG2;NCOR2;SNED1;AAK1;KCTD10
ELAVL4;RBMS2;PABPC1;LARP4;KHDRBS1;RBMS1;PABPC4;DDX3X;DDX1;ZC3H14;HNRNPU;RBPMS

NMRK2;NT5C3A;IDNK;FAM20A;PKLR;CHKB;NADK2;GK;NADK;PGM2L1;DGKG;SPHK1;GALK1;FGGY
ZNF366;PPARGC1B;CNOT1;LATS1;ARID5A;NSD1;PRMT2;TACC1;DCAF13;NRIP1;MED1;STRN;NCOA6;TRIP4;DCAF1
NEIL3;CDC45;MCM10;MCM7;HNRNPA1;MCM4;JCHAIN;TOP1;SUB1;DHX9;FBH1;HMGB2;FUBP3;SPRTN;HMGB1;HSPD1;RAD51;MCM8;TWNK;FUBP1;SSBP1;BLM;MSH3;RBMS1;MCM2;ERCC4;SAMHD1;RPA1;POLR2D;MCM9;PMS2;POLR2G;SHOC1;RAD51AP1;FAM111A;TDP1
MAP2;APOE;LGMN;CDK5;MAP1A;STUB1;ROCK2;PIN1;MARK1;BAG2
FNIP2;DNAJB2;ATP1B3;ATP1B1;DNAJC1;BAG1;BAG2;KATNB1;BAG3;SIL1;DNAJC19;HSPBP1

CHRNE
NMRK2;FN3K;ITPKB;MASTL;IDNK;EPHA4;CHKB;NADK2;TRIB3;CDK5;CKB;FN3KRP;GK;NADK;PGM2L1;DGKG;SPHK1;NEK1;PSTK;COQ8A;CDKL5;ITPKC;PRKD1;PINK1;DGKE;PMVK

IFIT5;PNPT1;RBM11;SSB;PABPC5;RBMS3;FMR1

GABRA3;GABRE;GABRB3

CLCNKA;CLCN6;CLCN4;VDAC1;CLCN2;GPR89A;GPR89B

ALDH1A2;ALDH9A1;ALDH3B1;ALDH1A1;ALDH4A1

CAMK1D;DAPK1;CAMK1;MKNK1;PTK2B;MYLK3;PHKA2

MCTP2;DYSF;ESYT2;PLA2G4C;CPNE2;SYT11;SYT12;PCLO;ANXA4;SYT17;SYT1;CPNE8

FLT1;FGFR1;TIE1;ERBB4;ROR1;MATK;KDR;PDGFRA;JAK3;RET;EPHA4

ARMT1;SETD3;TARBP1;DNMT3B;CARNMT1;DNMT1;HNMT;TPMT;DPH5;PCMT1;DNMT3A;METTL4;METTL2B

RGCC;SPDYA;PRKAG2;FAM20A;LTF;CALM3;DUSP19;CAB39L;FGF13;CDKN1A;GPRC5B;CD24;RICTOR;MLST8

GJA1;SLC1A3;SLC7A2;MFSD12;SLC6A7;SLC19A2;MPC2;CD36;SLCO6A1;SLC38A5;SLC35D2;SLC25A21;MPC1;SLC38A1;SLCO4A1;SLC3A1;SLC1A4;SLC5A6;SLC2A10;SLC36A4;SLC17A9;SLC38A6;MFSD2A;SLCO5A1;SLC2A14;LRRC8A;SLC25A1;SLC13A3;SLC25A18

AGAP2;PIK3R5;PIK3R6;SOCS6;SLA2;SOCS4;CISH;PIK3R1;SOCS5

TESC;MGAT5;STYX;BOD1L1;PPP1R9B;URI1;CABIN1

PDE6B;PDE8A;PDE3B;PDE8B;PDE7A;PDE4B;PDE4C;PDE2A;PDE10A

SCIN;PARD3;PLEKHA4;KCNJ1;SVIL;PHLDA3;FZD7;ALOX15;PLCD1;PLCB1;SYTL2;SNX20;LDLRAP1;GSDME;WIPI2;CLVS1;HIP1R;MARK1;KCNJ2

GABRA3;GABRE;GRIN2D;GABRB3

TOP2A;TESC;GNAI1;LRRK2;EYA2;NUDT3;PLK1;RAP2A;NLRC4;TKT;ATP10D;FEN1;MARK2;STK38L;STK4;ERN1;ABL1;XXYLT1;LATS1;TDG;CERK;OPA1;HPGDS;PDXP;IDH2;NME1;RP2;IRAK3;PAPOLA;MAP3K6;DUT;SNCA

OPTN;PARP10;RNF168;MINDY1;DZIP3;TNIP2;SHARPIN;RAD23A;MINDY2

SCN1B;NEDD4L;WNK1;FKBP1B;ITPR1;RACK1;WNK3;NEDD4;STX8;STX1A

XPOT;EIF2S3;EEF1A1;HSD17B10;NSUN2;AARS2;NSUN6;NEMF;SLFN11;EIF2S3B;ALKBH1;EIF2A;DARS2;TRNT1;TRMT10B;ELP1;NAT10;MRPS27;PUS1;AARS1;FARS2;MTRES1;TRMT10C

MTFMT;TRIT1;TRDMT1;TRMT13;AARS2;QTRT2;ALKBH1;QRSL1;PTRH2;TRPT1;FTO;DTWD1;TRMT12;ELAC2;AARS1;GATC

TP63;ENAH;TCEAL1;NDFIP2;TRAF4;TCEAL9;LITAF;NDFIP1;WBP11;TCEAL8;TNK2;NFE2;RAPGEF2

SOCS1;SH3BP5;WARS1;SOCS3;DUSP19;CDKN1A;ITPRIP;CDKN1B;MAPK8IP1;HEXIM1;GSKIP;MBIP;PRKRIP1

APOE;CACNA1A;FZD6;APBB2;TGFB2;EPHA4;ITGA2;CD36;LRPAP1;MSR1;ITGB2;LDLR;APBA3;ITM2A;ACHE;LDLRAP1;COL25A1;BACE1;PFDN4;CLSTN1;INSR;PGRMC1

NRAP;PALLD;PDLIM4;SYNPO2

GPC3;PZP;CSTB;TFPI;A2M;TIMP2;CAST

AQP9;TRPV2;BCL2;PANX1;TMEM120A;MCUB;PANX2;BAX

KCNJ5;KCNMA1;KCNJ1;KCNG2;KCNJ15;KCNH4;KCNE1;KCNQ5;KCNF1;KCNH8

CACNA2D3;CACNA1D;ITGAV;CATSPER1;PKD2

ATP2B2;CCDC88C;SYNJ2;PRKN;F11R;FZD7;TGFBR3;TMEM88;PLEKHA1;GNG12;CRIPT;SSTR2

DNAJB2;ATP1B3;ATP1B1;DNAJC1

FEN1;PRORP;POP1;EXO1;AGO3;NUDT16L1;DROSHA;EDC3;ELAC2;POP5

BIRC3;NOL3;TNFSF14;BIRC7;PRDX5;PRDX3

PITPNM2;CBLB;TOB1;EPHA4;FLT3LG;PTPN1;CASS4;HYAL2;ALKAL1;ITGAX;ANGPT1;NEDD9;MST1;ANGPT4;PCNA;CEACAM1;ITGB1;PRR7;SHC2;DUSP3;SHC1;TRIM6;CD247

ISG20;NUDT7;IMP4;NUDT4

OASL;TRIP6

RGCC;SPDYA;PRKAG2;DUSP19;FGF13;GPRC5B

SOCS1;SH3BP5;INKA1;SOCS3;DUSP19;CDKN1A;ITPRIP;CDKN1B;MAPK8IP1;HEXIM1;CAMK2N2;GSKIP;MBIP;CIB1;PRKRIP1

ARMT1;DNMT3B;CARNMT1;DNMT1;NSUN2;FTSJ3;PRMT3;COQ3;AMT;METTL3;METTL4;EHMT1;MGMT;NSUN3;PRMT7;LRTOMT;TRMT11;PRMT1;EEF1AKMT2;BUD23

LARGE1;ARG2;PIM1;MGAT1;HMGCL;MPPED2;POMGNT1;TDP2;ABL2;SOD2;ADPRM;B4GALT1;GYG1;ARG1;B4GALT7;GALNT2;WRN;FAM20C

MDM2;FN1;CALM3;HSPA2

NCL;RPL5;RPS7;RPS13;RPL26;CCT5;SYNCRIP;IGF2BP1;UTP23;RPS14;RSL1D1;DDX3X;SHMT2;LARP1;GNL3;RPS3A

HMGB3;NEIL3;CLSPN;RECQL4;XRCC2;HMGB2;ABL1;HMGB1;MEF2C;BLM;MSH6;XRCC3;RAD51AP1

SLC7A2;MFSD12;SLC6A7;SLC22A16;CHRNE;SLC38A5;SLC29A4;SLC38A1;SLC1A4;SLC36A4;SLC38A6;SLC22A1;SLC8B1;SFXN2;SLC22A3;SLC41A1

KCNJ5;GRIK4;KCNJ1;KCNJ15;P2RX4;P2RX7;P2RX1;TRPM8;MCOLN1;KCNJ2;CCDC51;P2RX6;SHROOM2;GRIN2A;ASIC2;GRIN1;GRIN3B;KCNJ10;ASIC3;KCNN1

SMAD6;TGFBR1;SMAD7;ENG;TGFBR2

GABRA3;GABRE

WNT5A;FPR1;GNAI1;REEP1;USP4;ADA2;WNT2B;FZD1;ARRB1;GNA15;S1PR2;GNAQ;GRK2;GHRL

UHRF1;TET3;CHTOP;ZBTB33;MBD1;LHX4;MBD2;PRMT1

OPTN;RAB34;MLPH;RAB11FIP4;NCKAP1;SPTBN1;KIF3A;CDC42EP2;YBX3;
RTKN;IQGAP2;ATP6AP1;PARD6A;CDC42EP4;RRAGD;RASA1;USP6NL;BNIP
3;RUSC2;PIH1D2;MTSS2;RABGAP1L;ABCA1;RABGEF1;DAPK3;C9ORF72;R
AB8A;UNC13B;SGSM1;RAP1GAP;EXPH5;EPS8;RAB11FIP3;CDKL5;AKAP13;
DMXL2;PDE6D;MICALL1;NOX1;LCP1;RANBP2;WHAMM;PEX5L;CHML;RGP
D8;RALBP1;BICD1;FGD6;CIB1;AFDN;SIKE1;STOML2;NDRG1;MYO5B

UCHL1;USP2;USP18;USP11;USP36;USP49;USP51;USP37;CYLD;OTUD4;US
P31;MPND;OTUD1;USP25;JOSD1;USP28;USP44;ZC3H12A;OTUB1;USP50;
BAP1

GCNT4;LARGE1;B4GAT1;B3GLCT;RFNG;MGAT1;EXT1;HEXB;B3GNT2;PO
MGNT1;MGAT4C;PIGP;POMGNT2

ANXA6;GRIN2D;TRPV1

HMGB3;XRCC2;HMGB2;ABL1;HMGB1;BLM;MSH6;XRCC3

FGFR1;THBS1;ITGB3;S100A13

CCNA2;CDKN2C;CCNB1;CCNE2;CCNF;CCNJL;CDK5R1;CCNE1;CCNI;CCN
C;CCNH;CCNY;CCNT2;CCNB2;CCND1;CCNYL1

AQP9;SLC7A2;SLC19A2;ABCB1;SLC16A8;SLC6A12;SLC38A5;SLC38A1;SL
C3A1;SLC36A4;SLC38A6;SLC13A3

SHC3;SYK;VAV1;HCK;SAMSN1;PLCG2;PTPN11;PFN1;NCK2;ABL1;SH3BP2;
CRK;YWHAB;CRKL;PIK3R1;NEDD4;GRB2;VAV2

OAS2;OAS1;OAS3;HSPA4L;SYN3;NUAK2;ABCG1;PRPS1;ABCB1;PRKACB;K
CNJ1;PRKAG2;PFKP;ABCB6;DYRK3;PIM1;MAP4K3;DAPK2;HSPA2;PRAG1;
P2RX4;GUK1;ABCD4;SPHK1;GALK1;P2RX7;ABCA1;DAPK3;LMTK2;NEK11;
ABCB8;SIK3;PNPLA8;CDKL5;MYH10;SIK1;SYN2;SYN1;STK38;ABCD1;PTPA
;LATS2;PINK1;INSR;PDK2;MARK1;PMVK;RPS6KA4;PFKL;AKT3;CARS2;MS
H2;ATP1A3;EIF2B2;MVK;P2RX6;BMPR1B;BCKDK

ECHDC1;BCKDHA;GAD1;CSAD;PISD;GGCX;UROD

SLC26A1;SLC46A1;SLC25A10;SLC38A7;SLC25A13;SLC26A2;SLC25A12;S
LC26A3;SLC1A5

CAVIN3;HSPB1;DSP;HDAC9;PLEK;PARD6A;PRKCB;IRS1;GRK5

SLC26A1;ANKH;SLC39A8;SLC25A10

IFIT5;PNPT1;RBM11;SSB;PABPC5;RBMS3;FMR1

HIP1;DAB2;AP2A2;AP2A1;AP2S1;AP1G1

ABCG1;PRKAG2;MYO6;COQ8A;MYH10;ABCD1;MSH2

PALLD;BSG;CD47;JAM3;VSTM2L;NEXN;CDC42EP1;EPCAM;ROBO3;LRRC4
C

GABRA3;GABRE

PRKN;RHBDD2;RAD23A;DERL1;SELENOS;SUMO1

MACROD1;MACROD2;DNPH1;NTHL1;ADPRS;ADPRH

CALD1;TNNT1;SMTNL1;SMTNL2;TMOD1;TMOD2;NEBL

EPHB3;EPHA2

EPHB3;EPHA2

ANO5;GABRB2;APOL1;SLC1A4;CLIC4;GABRG2;GABRR2;LRRC8A;GLRA1;FXYD1;GLRB

OCA2;SLC43A2;SLC36A1;SLC7A5;SLC7A6;SLC7A7;SLC7A1;SFXN1

ANKH;SLCO2B1;SLC7A5;SLC16A10;SLCO4C1;SLC20A1;SLC20A2;SLCO3A1;SLC4A4;SLC35A1;SLC25A10

RNF125;RNF144B;RNF19B;RPS3;ARIH2;DCUN1D4;TCERG1;DCUN1D1;MARCHF7;DCUN1D2;ZMYM2;RNF185

ELAVL4;RBMS2;ATXN1;PABPC1;LARP4;KHDRBS1;RBMS1;DAZAP1;PABPC4;DDX3X;DDX1;ZC3H14;HNRNPU;RBPMS

WNT5A;WNT2B;FZD1

SHC3;SYK;VAV1;HCK;SAMSN1;PLCG2;PTPN11;PFN1;NCK2;ABL1;SH3BP2;CRK;CRKL;PIK3R1;GRB2;VAV2

MFSD12;SLC6A7;SLC38A5;SLC38A1;SLC1A4;SLC36A4;SLC38A6;SLC43A1

CACNA1A;SNAP25;SYPL2;NAPA;VAMP2;SYP;TMED10;ABCA1;VAMP3;UNC13B;STX10;STX16;SYT12;RAB11A;SYT17;SYT1;SNPH;STXBP5L;SNAP47;TMED9;SYPL1;STXBP4;STXBP5

CAT;KPNB1;TOMM70;IPO5;BRAP;KPNA6;KPNA3;TNPO1;TNPO2;SEC61A1;KPNA4;SEC61A2;POM121C;NUP153;NUP214;BABAM2;RANBP6;TOMM20

TENT4B;PNPT1

ATOH8;TAL1;MYC;SNAI1;AHR;CLOCK;MAX;SCX;TCF3

ST6GALNAC3;ST6GALNAC2;ST3GAL4;ST3GAL5;ST8SIA5;ST6GAL1

OPTN;RAB34;MLPH;RAB11FIP4;NCKAP1;KIF3A;CDC42EP2;YBX3;RTKN;IQGAP2;ATP6AP1;PARD6A;CDC42EP4;USP6NL;RUSC2;PIH1D2;MTSS2;RABGAP1L;ABCA1;RABGEF1;DAPK3;C9ORF72;RAB8A;UNC13B;SGSM1;RAP1GAP;EXPH5;EPS8;RAB11FIP3;CDKL5;AKAP13;DMXL2;PDE6D;MICALL1;NOX1;RANBP2;WHAMM;PEX5L;CHML;RGPD8;RALBP1;BICD1;FGD6;CIB1;AFDN;SIKE1;NDRG1;MYO5B

KCNMA1;KCNN4;GRIK4;TMEM38B;KCNG2;KCNH4;KCNE1;KCNQ5;KCNF1;KCNH8;KCNMB2

STAT1;CITED2;EPAS1
SMAD3;RGCC

SETD3;KMT5B;KMT5A;SETDB1;SETD2;ATPSCKMT;SETD7;EHMT1;SMYD2

PCOLCE2;EBAG9;APH1B;APH1A;PINK1;PSME2;ADRM1

SLCO2B1;SLC39A8;ABCG2
SLC35D2;SLC35E3;SLC35E4;ATP1A3;SLC25A11
TRPV4;RACGAP1;CDK5R1;LRPPRC;CCT5;MAP1S;GABARAP;PDCD5;TAOK1;SMC3;TBCD;HTT;SPAST

MYO1G;HIP1;ARHGAP9;GSDMA;ARAP3;FCHSD2;SH3PXD2B;RACGAP1;ARFIP1;NCF4;SNX24;DAPP1;RUBCNL;PLEKHA5;MAPKAP1;DENND1A;SNX13;PLA2G4A;TECPR1;GBF1;SNX27;ADAP2;DENND1C;OGT;CYTH3;SNX11;PLEKHA8;ANKFY1;SNX12;OSBP;BTK;ZFYVE1;IQGAP1;ZFYVE28;GOLPH3
EIF2AK2;LGALS3;RCAN1;CALM3;PPP2R3A;B3GAT3;LMTK2;PPP2R1A;PTPA;PPP1R15A;PPP1R16A;PPME1;PTN;PHACTR4;PPP2R5B;PPP1R7;PPP1R15B;PPP2R5C;BOD1;PPP2R2C;RCAN2;PPP2R5A
SNAP25;SYPL2;VAMP2;SYP;VAMP3;UNC13B

HVCN1;PKD2
CAVIN1;RPS3;RPL5;RPL23;RPLP0;PPAN;RPS13;FASTKD5;PTCD3;NOP53;RPL11;UTP23;RPS14;MTERF4;DDX21;RRS1;RPF2;RPS5;MRPS27;RPL12;ANG

ALDH1A2;ALDH9A1;ALDH3B1;FAR2;ALDH1A1

KCNMA1;KCNN4;TMEM63A;KCNMB2

CYB5A;CYBRD1;CYB561A3;NDUFV1;NDUFB6;NDUFC1;NDUFS8;UQCRC1;NDUFA7;UQCRFS1;NDUFA5;NDUFC2;CYB561D2;NDUFB4
PLCL1;SMPD1;SMPDL3A;FAM83B;MPPED2;GPLD1;TDP2;PLD6;SMPD3;ENPP2

DPP4;SEMA5A;SEMA3D;SEMA3C;SEMA3A

PIK3R5;PIK3R6;SOCS6;SLA2;SOCS4;CISH;PIK3R1;SOCS5

RTN4R;PHYH;EPDR1;SESN1;THNSL2;GLRA1

TRDMT1;TRMT13;FTSJ3;METTL3;METTL4;TRMT12;TGS1;TYW3;FBLL1

TET3;KDM7A;HSPBAP1;KDM2A;TET2;KDM4A;ALKBH3;PHF8;HIF1AN;ALKBH1;TMLHE;FTO;ALKBH4;KDM2B;KDM4B;RIOX2

CYB5A

B3GNT7;B3GNT5;B3GALNT1;GBGT1;B4GALNT3;B3GNT9;B3GNT8;EXTL2;B3GALT9

GSDMA;ARFIP1;RUBCNL;PLEKHA5;PLA2G4A;PLEKHA8;OSBP;GOLPH3

THRA;ESR1;DRAP1;BRF2;GTF2B

CPE;PEPD;CPXM1

ISG20

PPM1E;LRRC39;CTDSPL;DUSP1;PPM1H;PPP1CB

DGKQ;NRG1;GAS6;GHRL;PAK2;ERCC6

MYO1G;ARHGAP9;ARAP3;FCHSD2;RACGAP1;DAPP1;MAPKAP1;GBF1;ADAP2;OGT;CYTH3

PLTP;MICALL1;SMPD3;MARK1;PLEKHN1;COMMD1;PITPNM1;RAPGEF2

STAT1;MMP12;DRAP1;KLF10;RELA;GTF2B;FOS;AGO2;RRN3;TAF1C

MLLT3;THAP7;DPF2;BRD7;YEATS4;CARM1

MLLT3;THAP7;DPF2;BRD7;YEATS4;CARM1

TARS1;NARS1;MARS1;LARS1;AARS2;QARS1;YARS1;DARS2;RARS1;FARSB;AARS1;FARS2;HARS2

DUSP5;SMPD1;SMPDL3A;ACP2;DUSP18;DUSP15;MPPED2;PXYLP1;PPP1CB

SLC5A3;KCNIP1;RASA1;SGK1;KCNE1;KCNMB2;DPP6;SGK2;AMIGO1

SLC12A9;SLC12A5;PKD2;KCNA1;TMEM38A;KCNG1;SLC12A2

NAT14;NAA80;NAA10;SAT1

PLEKHA4;PHLDA3;WIPI2;CLVS1;HIP1R;KIF16B;WIPI1;WDR45B

SLC9A9;SLC8A2;SLC9C2;SLC8B1;SLC9A8;SLC8A3;SLC41A1;SLC9B2;SLC9A3;SLC4A8;ASIC3;SLC9A2;SLC4A11

SMC6;MPG;CRY2;NTHL1;DDB2;XPC;PCNA;NBN;UNG;RBBP8;SDE2

SLC16A7;SLC6A11;SLC27A1;SLC16A3;ABCG2;SLC6A6

PRC1;KIF18B;FAM83D;RAB29;IFT88;SNCA;AP1G1;MAPK8IP3

NAT14;SPHK1;NAA80;NAA10;SAT1;GTF2B;NAA16;ACAT1

ANXA11;ATP1B1;CD81

CDK1;CDK14;CDK8;CDK13;CDK12;CDK6;CDK19

CDK1;CDK14;CDK8;CDK13;CDK12;CDK6;CDK19

ANXA6;GRIN2D

TMTC1;ALG2;DPY19L3;DPY19L1

MAPK10;SGK1;MAP2K3;RPS6KA2;MAP3K9

FXYD6;SGK1;FGF13;GPLD1;FXYD5;CAMK2D;FXYD1;FGF14;SGK2;PRSS41

DIXDC1;BRCA2;GIT1;BLOC1S2;RAB11FIP5;RAD51D;NDRG1

ASS1;DDAH1;THNSL2;GLRA1;GLRB;GRIN1;GRIN3B;PADI4;DDAH2

UBE2L6;UBE2R2;UBE2D1;UBE2D4;UBE2J1;UBE2D3;UBE2F;UBE2Z;CDC34;AKTIP;UFC1;UBE2B;UBE2O;UBE2Q1;UBE2I;UBE2L3

NDUFA4;NDUFA1;NDUFB1;NDUFA12;NDUFB3;NDUFB2;NDUFA2;NDUFS7;NDUFV3;NDUFS6;NDUFB8;NDUFA13;NDUFS5;NDUFB5;NDUFA10;NDUFA8

NDUFA4;NDUFA1;NDUFB1;NDUFA12;NDUFB3;NDUFB2;NDUFA2;NDUFS7;NDUFV3;NDUFS6;NDUFB8;NDUFA13;NDUFS5;NDUFB5;NDUFA10;NDUFA8

PIWIL4;ZC3H12D;APLF;ZC3H12C;NTHL1;RCL1;RAG1;PIWIL2;ZC3H12B;AGO2;LACTB2;ZC3H12A;SLFN13;TSNAX;RNASEL

KIF2C;KIF18A;FBXW11;NUMA1;CLIP2;KNSTRN;APC;PAFAH1B1

EFNB2;PTPN1;EFNA1;EFNA5;EFNB1;SHC1;EFNA2

MAPK10;MAP2K3;MAP3K9

RPS3;ERN1;SACS;DNAJC2;SNCA;GPR37;DNAJB14;SPN;HDAC8;TSC1;DNAJB1;DNAJC10

SLC1A4;SLC13A3;SLC25A18;SLC25A22;SLC25A11

DHX9;WAC;PCF11;CTR9;URI1;SCAF8;ELP4;LLPH;SMYD2;RPRD1B;CCAR2;RECQL5;HNRNPU;WDR43;ZNF326

SHFL;TMEM147;NCLN;ERI1;SBDS;NAA10;MAIP1;NAA16;CPEB2;FMR1;ZC3H12A;EIF6;IGHMBP2;EFL1;PYM1;YTHDF3

UBE2L6;UBE2R2;UBE2D1;UBE2D4;UBE2J1;UBE2D3;UBE2Z;CDC34;AKTIP

PRKCE;TMSB10;LIMA1;ABL2;COBL;PFN4;TMSB4Y

TENT4B;XRN1

PYGL;ABAT;OAT;PNPO

RNF125;RNF144B;RNF19B;ARIH2;DCUN1D4;DCUN1D1;MARCHF7;DCUN1D2;ZMYM2

SLC26A1

GRIK4;GRM8

HTR2B;CHRM3;CHRM4

PEMT;N6AMT1;RNMT;ZCCHC4;CSKMT;TRMT1L;FDXACB1;TRMO

SCN1B;SCN9A;PKD2;PKD2L1;GRIK1
HDAC1;HDAC3;NPM1;MTDH;HDAC2;HIF1AN;GSK3B;EP300;TP53BP2

HTR2B;CHRM3;CHRM4

HTR2B;CHRM3;CHRM4
FAM20A;LTF;CALM3;CAB39L;RICTOR;MLST8;IRGM;PARP16;CCNT1;DBF4B
HELZ2;PRKCB;FGF2;PPARGC1A;CCDC62;CALCOCO1;TRERF1;SFR1;NCOA2;HMGA1

PPP1R16B;KCNQ1;AKAP11;PPP1R9B

KL

USP7;USP13;USP9X;YOD1;MINDY3;USP27X;DESI2;USP15

SLC25A4;SLC26A11;SLC4A2;SLC26A6;SLC25A11
MARK2;MARK4;PRKAA1;MARK3;GSK3B;TAOK1;TAOK2;PHKG2;ROCK1;DYRK1A;BRSK1

GRIK4

NPPC;SOCS2
